# Mapping the genetic evolutionary timeline of human neural and cognitive traits

**DOI:** 10.1101/2023.02.05.525539

**Authors:** Ilan Libedinsky, Yongbin Wei, Christiaan de Leeuw, James K. Rilling, Danielle Posthuma, Martijn P. van den Heuvel

**Author notes:** **Corresponding author:** Martijn P. van den Heuvel, PhD, De Boelelaan 1085, Amsterdam, 1081 HV, the Netherlands. **E-mail:**.

## Abstract

Human evolution is characterised by extensive changes of body and brain, with perhaps one of the core developments being the fast increase in cranial capacity and brain volume. Paleontological records are the most direct method to study such changes, but they can unfortunately provide a limited view of how ‘soft traits’ such as brain function and cognitive abilities have evolved in humans. A potential complementary approach is to identify when particular genetic variants associated with human phenotypes (such as height, body mass index, intelligence, and also disease) have emerged in the 6-7 million years since we diverged from chimpanzees. In this study, we combine data from genome-wide association studies on human brain and cognitive traits with estimates of human genome dating. We systematically analyse the temporal emergence of genetic variants associated with modern-day human brain and cognitive phenotypes over the last five million years. Our analysis provides evidence that genetic variants related to neocortex structure (e.g., area, thickness; median evolutionary age = 400,170 years old), cognition (e.g., fluid intelligence; median age = 459,465), education (median age = 637,646), and psychiatric disorders (median age = 412,639) have emerged more recently in human evolution than expected by chance. In contrast, variants related to other physical traits, such as height (median age = 811,305) and body mass index (median age = 794,265), emerged relatively later. We further show that genes containing recent evolutionary modifications (from around 54,000 to 4,000 years ago) are linked to intelligence (*P* = 2 × 10^−6^) and neocortical surface area (*P* = 6.7 × 10^−4^), and that these genes tend to be highly expressed in cortical areas involved in language and speech (pars triangularis, *P* = 6.2 × 10^−4^). Elucidating the temporal dynamics of genetic variants associated with brain and cognition is another source of evidence to advance our understanding of human evolution.

## Main

The human genome contains a footprint of our evolutionary history. Mapping the timeline of when specific genetic variants associated with modern-day phenotypes have emerged in our genome provides a unique window to examine the evolution of human features. This approach is particularly valuable for studying the evolutionary timeline of neural and cognitive traits^1^, which leave no direct physical traces in the fossil record.

Human evolution involved dramatic changes to the brain and human cognitive abilities. Cranial capacity tripled over the past two million years of human evolution, a pace of encephalization that is unparalleled in other mammals^2^, with the neocortex being among the most expanded brain regions^3^. This expansion is widely believed to be one of the strongest catalysers for the development of complex behaviour and advanced cognition in the human lineage^3–5^. Human encephalization is thought to be the outcome of a wide range of natural (e.g. climate), nutritional (e.g. diet), and social (e.g. group size, parental care)^6^ selection pressures. These pressures have synergistically interacted with, and led to, modifications in the human genome at different periods across history^7^. In recent years, a rapidly growing number of Genome-Wide Association Studies (GWAS) have begun to unravel the genetic basis of modern human phenotypes^8^, and in this study we combined dating estimates for the emergence of variants in the human genome^9^ with these GWAS findings^8^ to map the genetic evolutionary timeline of human brain and cognitive abilities. We show that the emergence of genetic variants associated with core human attributes, such as brain morphology, cognition, and neuropsychiatric conditions, follows a distinctive temporal pattern, revealing recent genetic modifications in human evolution.

## Results

### Genetic timelines of human phenotypes run parallel to evolutionary milestones

We began by examining when single nucleotide polymorphisms (SNPs) associated with a wide range of human phenotypes appeared in the human genome. Data from the Human Dating Genome project (HDG)^9^ were used for this purpose. The HDG is a database that, by tracing back alleles to a mutation event in the genome of a common ancestor, describes estimates of the emergence date of ∼13.5 million genetic variants across 22 chromosomes of the human genome covering ∼200,000 generations. This database provides dating information from ∼5.1M to ∼100 years ago (Methods, Supplementary Methods). We first examined the genetic timeline of a total of 67,119 variants (36,506 unique lead SNPs; referred to as human-phenotypic SNPs), combined from the summary statistics of 2,538 GWAS^8^ associated with a broad range of human phenotypes (e.g. eye shape, cancer, height, brain, cognition, behaviour). Using the HDG we computed the dates when human-phenotypic SNPs appeared and compared them to the dates for all other SNPs in the human genome. The combined genetic timeline of the 36,506 human-phenotypic SNPs (ranging from ∼4.5M to ∼2,000 years ago, median evolutionary age = 673,520 years old) displayed a bimodal distribution with two distinct peaks (Figure 1A): an ‘old peak’ of genetic variants arising from ∼2.4M until ∼280,000 years ago reaching a maximum at ∼1.1M years ago, comprising ∼85% out of all variants with high minor allele frequency (MAF >= 0.4; Supplementary Methods); and a second ‘young peak’ of genetic variants arising at ∼280,000 until ∼2,000 years ago reaching a maximum at ∼55,000 years ago, a period including ∼62% of variants with low MAF (MAF <= 0.1; Figure 1B).

**Figure 1.**
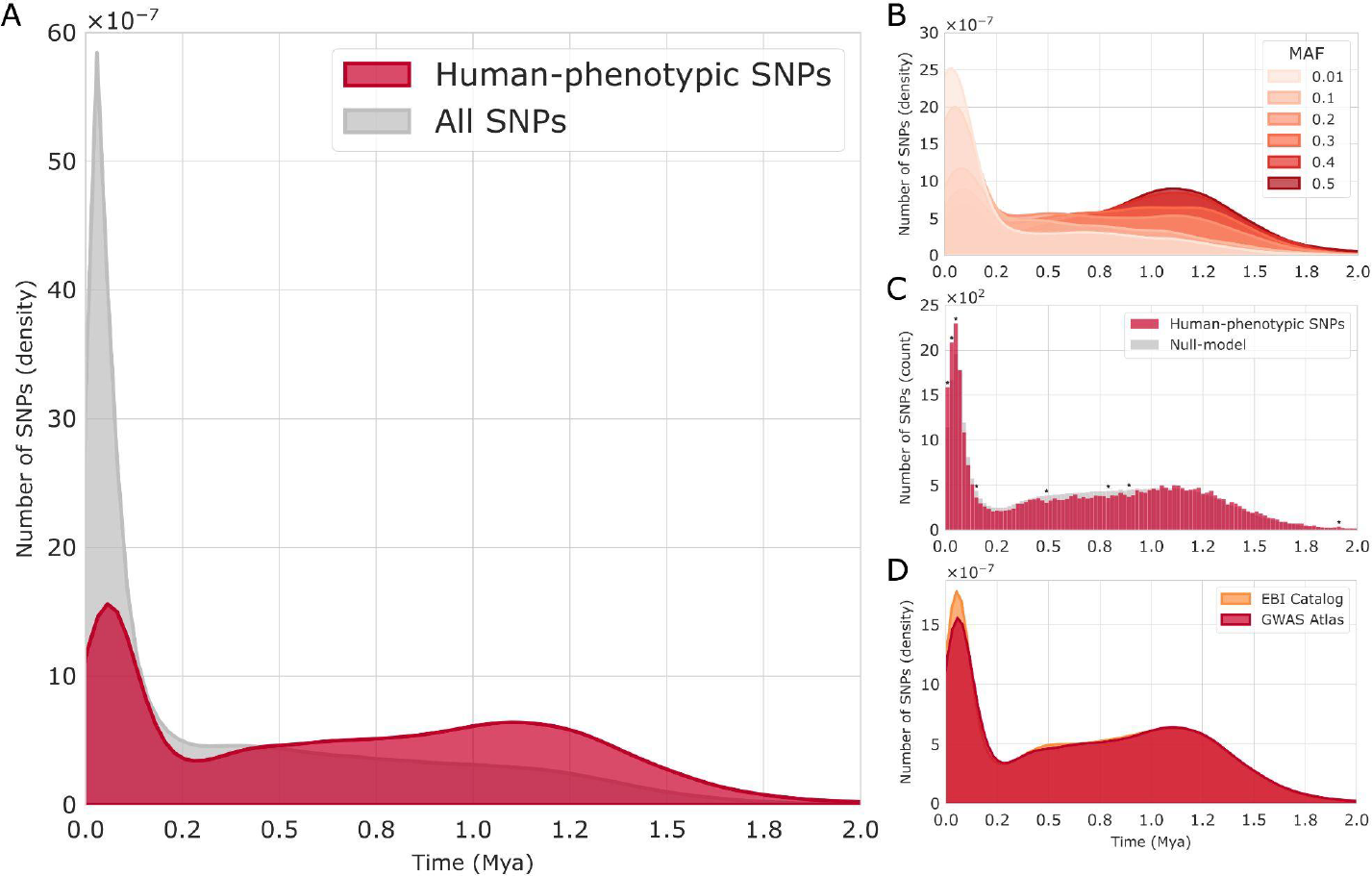
Genetic timeline of human-phenotypic SNPs. *(a)* Timeline of the density (normalised count; y-axis) of all significant human-phenotypic SNPs (red) associated with human traits extracted from the GWAS Atlas repository (2,538 GWAS studies, 36,506 unique SNPs) and the total set of SNPs (grey) extracted from the Human Dating Genome (HDG; ∼13.5 million SNPs) across the last 2M years (x-axis). *(b)* Timeline of the density of human-phenotypic SNPs per minor allele frequency category (MAF; six bins ranges: 0 - 0.01, 0.01 - 0.1, 0.1 - 0.2, 0.2 - 0.3, 0.3 - 0.4, 0.4 - 0.5). *(c)* Absolute count of number of SNPs per time-bin (100 bins of ∼20,000 years old) of human-phenotypic SNPs (red). Asterisks (*) denote bins where the number of human-phenotypic SNPs per age bin was significantly higher or lower than the null-model of random equally sized sets of SNPs selected from all SNPs from the HDG (MAF-controlled, 100 tests, Bonferroni *P* value threshold < 5 × 10^−4^). *(d)* Validation: timeline of the density of human-phenotypic SNPs (GWAS Atlas) compared to the alternative EBI Catalog of GWAS results of human traits showing high consistency between both distributions of genetic variation (*rho* = 0.96, *P* = 5.5 × 10^−59^). Mya, million years ago; MAF, Minor Allele Frequency.

Permutation testing comparing the timeline of human-phenotypic SNPs to that of all SNPs allowed to identify periods of time when there were more (or less) genetic variants than expected by chance (MAF-controlled, 10,000 permutations; see Supplementary Methods). This analysis revealed a significant increase in human-phenotypic SNPs at the beginning of the old peak (from ∼1,920,000 to 1,900,000 years ago, *P* = 2.5 × 10^−8^). A high concentration of human-phenotypic variants was also found at the highest point of the young peak, ranging from ∼61,000 to ∼2,000 years ago (three consecutive significant bins, *P* = 4.2 × 10^−16^, *P* = 6.2 × 10^−27^, and *P* = 2.3 × 10^−40^; Figure 1C). After the highest point of the old peak, three separate time periods show a slightly less human-phenotypic SNPs than expected (∼900,000 to ∼880,000 years ago, *P* = 1.7 × 10^−4^; ∼800,000 to ∼780,000 years ago, *P* = 3.1 × 10^−4^; ∼500,000 to ∼480,000 years ago, *P* = 5.0 × 10^−5^), and once after the beginning of the young peak (∼160,000 to ∼140,000 years ago, *P* = 4.2 × 10^−4^; Figure 1C). We observed this bimodal distribution again by selecting human-phenotypic SNPs from the EBI Catalog^10^ as an alternative resource (including unique non-overlapping variants with GWAS Atlas, *n* = 88,362), resulting in a highly similar distribution (*rho* = 0.96, *P* = 5.5 × 10^−59^; Figure 1D, Supplementary Results). Figure S1 further shows the genetic timeline of human-phenotypic SNPs grouped by their functional genetic consequences (e.g. intergenic, missense).

Plotting the bimodal genetic timeline side-by-side with the emergence of evolutionary milestones in the human lineage identified several remarkable points of overlap: the old peak coincides with (among others, see also Discussion) the origin of *Homo erectus* (∼1.8M years ago)^11^ and the divergence of *Homo sapiens* and Neanderthals (∼800,000 years ago)^12^. The beginning of the young peak co-occurs with the origin of modern humans in Africa (∼200,000 years ago)^13,14^ and the highest point of the young peak (∼55,000 years ago) matches with the transition to the Upper Paleolithic (∼45,000 years ago), a period considered as the origin of modern human behavior^15^.

### Psychiatric phenotypes influenced by evolutionarily recent genetic modifications

We utilised data from the GWAS Atlas^8^ to gain insight into the genetic timeline of specific classes of human characteristics. This procedure involved grouping identified genetic variants according to a nested hierarchical organisation, ranging from global categories to specific phenotypes according to domain, chapter, subchapter, and trait levels. Using permutation testing, we assessed the median evolutionary age for each phenotype to determine whether it was younger or older than expected by random chance (null-distribution of equally sized random sets out of all human-phenotypic SNPs, details in Methods). At the domain level (*n* = 28 domains, Bonferroni *P* value threshold < 1.7 × 10^−3^; Figure 2, Figure S2), genetic variants related to ‘Psychiatric’ phenotypes presented an evolutionary age younger than expected by chance (median evolutionary age = 412,639 years old), a class of traits suggested to be linked to human brain evolution^5,16–20^. ‘Ophthalmological’ and ‘Nutritional’ phenotypes as well displayed a significantly younger evolutionary age (median age = 30,484 and 429,993, respectively). In contrast, genetic variants associated with ‘Metabolic’, ‘Skeletal’ and ‘Immunological’ phenotypes overall displayed the relatively oldest age estimates (median age = 780,323, 759,056, and 747,780, respectively; all *P* values listed in Table S1).

**Figure 2.**
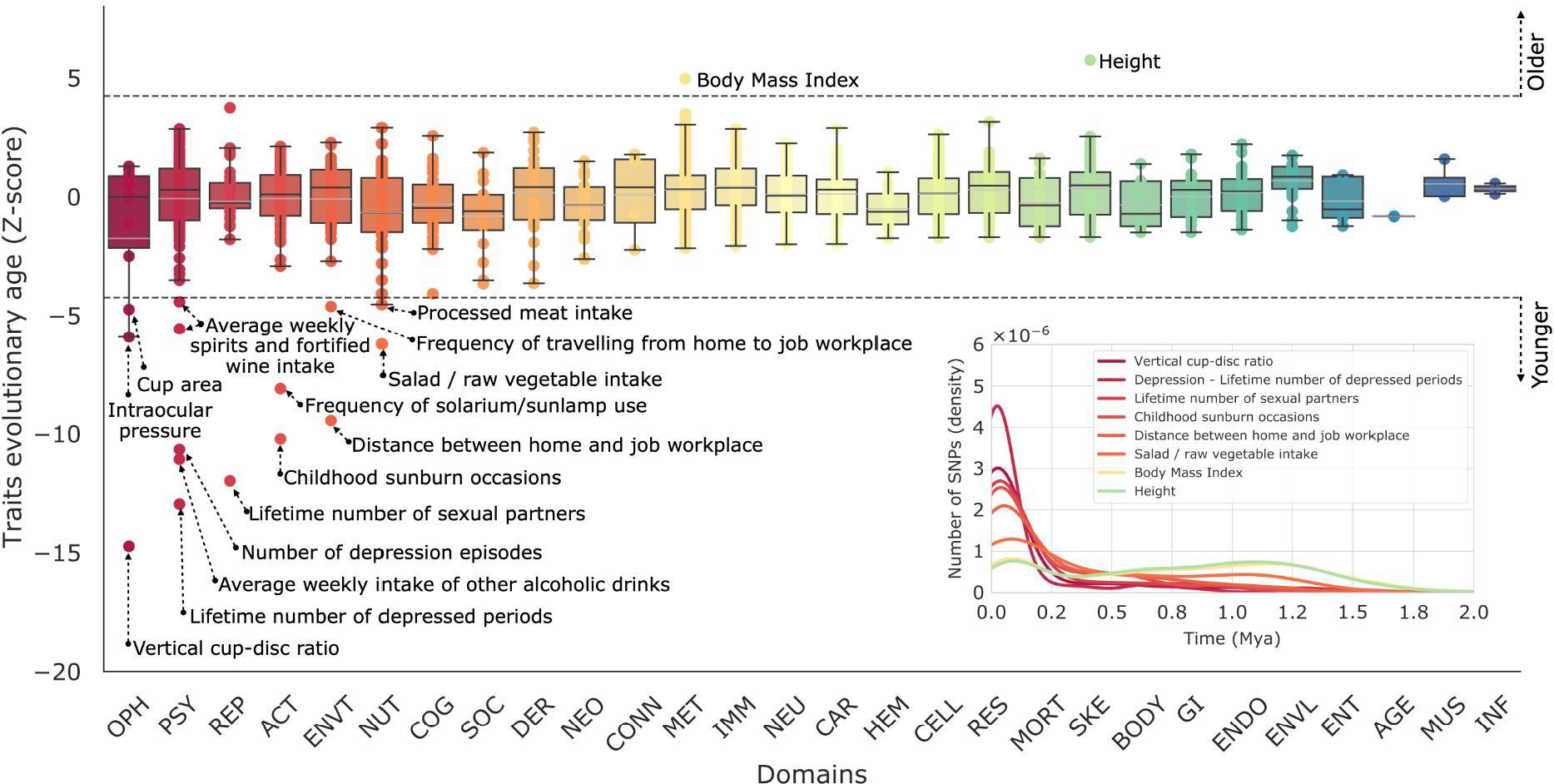
Genetic timeline of human traits. Expected age of traits (dots) under a null-model (see *Statistical inference* in Methods). Z-scores beyond the dotted line indicate traits (*n* = 2,251) with a median age of SNPs significantly younger (negative z-score values) or older (positive z-score values) than expected by chance (Bonferroni *P* value threshold < 2.2 × 10^−5^). Black and grey lines within boxes denote domain median and mean z-scores, respectively. Exemplary genetic timelines of SNPs from significant traits are shown in the insert. Colours denote the different domains in GWAS Atlas. OPH, ophthalmological; PSY, psychiatric; REP, reproduction; ACT, activities; ENVT, environment; NUT, nutritional; COG, cognitive; SOC, social interactions; DER, dermatological; NEO, neoplasms; CONN, connective tissue; MET, metabolic; IMM, immunological; NEU, neurological; CAR, cardiovascular; HEM, haematological; CELL, cell; RES, respiratory; MORT, mortality; SKE, skeletal; BODY, body structures; GI, gastrointestinal; ENDO, endocrine; ENVL, environmental; ENT, ear, nose, throat; AGE, ageing; MUS, muscular; INF, infection.

### Sensitivity analyses

We performed a series of sensitivity analyses to test the robustness of the presented findings. We first confirmed our results by examining also the more fine-grained chapter (Figure S3), subchapter (Figure S4), and trait level (Figure S5), which all showed similar results with the findings reported above (for all results see Table S1). We further examined whether our findings were robust when controlled for differences in the number of SNPs per phenotype, the sample size of the included GWAS, and when taking into account variants with the most accurate date estimates (Supplementary Results). We also considered only independent SNPs in Linkage Disequilibrium (LD) blocks^21^ (R^2^ < 0.1) and outside the MHC (Supplementary Results). All these analyses confirmed our main findings.

We also specifically examined the effect of an expected positive association between the MAF and the age of emergence of a genetic variant (*r* = 0.38, *P* < 1 × 10^−100^). We investigated whether the median evolutionary age of variants related to a phenotype deviated from the expected values when taking into account both the number of variants and their corresponding MAF (Supplementary Methods; 33,621 unique SNPs age and MAF estimates were available; 26 domains, Bonferroni *P* value threshold < 1.9 × 10^−3^). The results of this sensitivity analysis suggest that selection pressure may have influenced the variants linked to significant phenotypes^22^. When controlling both polygenicity and MAF, SNPs related to ‘Psychiatric’ phenotypes presented the most recent evolutionary age (median evolutionary age = 475,833 years old), followed by ‘Activities’ (e.g. looking after one’s health, use of communication devices; median age = 644,450), and ‘Environment’ phenotypes (e.g. education, work; median age = 626,965). In contrast, SNPs associated with ‘Neoplasm’ and ‘Metabolic’ phenotypes exhibited the relatively oldest evolutionary age (median age = 591,690 and 785,572, respectively; all *P* values listed in Table S2). The effects observed in the ‘Psychiatric’, ‘Activities’, and ‘Neoplasm’ domains further remained statistically significant after also controlling for LD (by including SNPs with an R^2^ below 0.1 and excluding SNPs within the MHC; Supplementary Results).

### Neocortical variation linked to evolutionarily recent genetic modifications

Further examination was conducted on the genetic timeline of several brain structures by analysing the date estimates of 6,284 genetic variants (2,476 unique BRAIN-SNPs) collated from extensive GWAS on 1,744 neuroimaging-derived phenotypes from the UK Biobank (e.g. cortical surface, myelin, volume of brain structures; Table S3)^23^. The genetic timeline of BRAIN-SNPs ranged from ∼3.6M to ∼4,000 years ago (median evolutionary age = 716,318 years old; Figure S7). Neocortex SNPs (a subset of 130 SNPs out of the 2,476 BRAIN-SNPs; Supplementary Methods) exhibited a median evolutionary age significantly younger than expected by chance when controlling for the number of variants (median age = 400,170 years old, *P* = 4.4 × 10^−4^; Figure 3A; complete results in Table S4) and a trend towards significance when controlling for MAF as well (*P* = 0.06). The neocortex is believed to be the most expanded brain structure in the *Homo* lineage^24^, now vastly exceeding the volume and size found in chimpanzees and bonobos^5,25^, and substrate of complex cognitive abilities including multisensory processing and sociability^26,27^. No significant results were found for date estimates of SNPs related to white matter (*n* = 676), cerebellum (*n* = 144), amygdala (*n* = 39), hippocampus (*n* = 30), or thalamus (*n* = 19) among others (Bonferroni *P* value threshold was applied < 5.5 × 10^−3^ to all tests performed; Supplementary Results).

**Figure 3.**
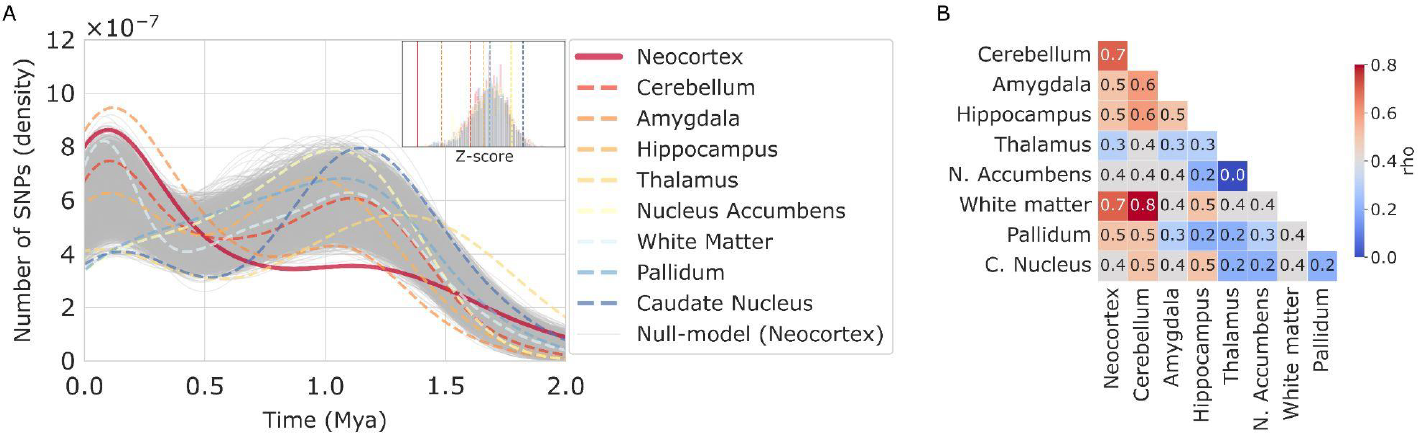
Genetic timeline of brain-imaging phenotypes. *(a)* Histogram showing the density (y-axis) of the number of SNPs per age bin (x-axis). Solid lines denote phenotypes (neocortex, red) significantly younger than expected by chance (see insert for the null-model of each brain phenotype). *(b)* Heatmap of the co-fluctuation between the timelines of SNPs related to brain structures (Spearman’s correlation). All correlations are significant (Bonferroni *P* value threshold < 1.4 × 10^−3^) except for the thalamus which only presents a significant correlation with white matter and cerebellum; and for pallidum which has a significant correlation with neocortex, cerebellum, nucleus accumbens, white matter, and caudate nucleus. Mya, million years ago.

### Genes containing recent mutations are enriched for genes related to cortical surface area, intelligence, and neuropsychiatric disorders

An estimate of the evolutionary age of each gene was derived by computation of the median age of the variants located within the start and stop position of each gene, resulting in date estimates for 18,328 human genes (out of 26,836 genes, GRCh37 human assembly^28^; Methods). Computed gene evolutionary age varied between ∼3M to ∼4,000 years ago with a peak density ∼70,000 years ago (Figure 4A). We evaluated whether the resilience of genes against mutations is related to their evolutionary age, by ranking genes according to intolerance to Loss-of-Function (LoF; Supplementary Methods)^29^. The top 2,000 most intolerant LoF genes were found to be on average younger than the top tolerant LoF genes (*t* = -4.89, *P* = 1.0 × 10^−6^; Figure 4B), indicating that genes intolerant to LoF contain genetic variants relatively recent in evolution compared to tolerant LoF genes.

**Figure 4.**
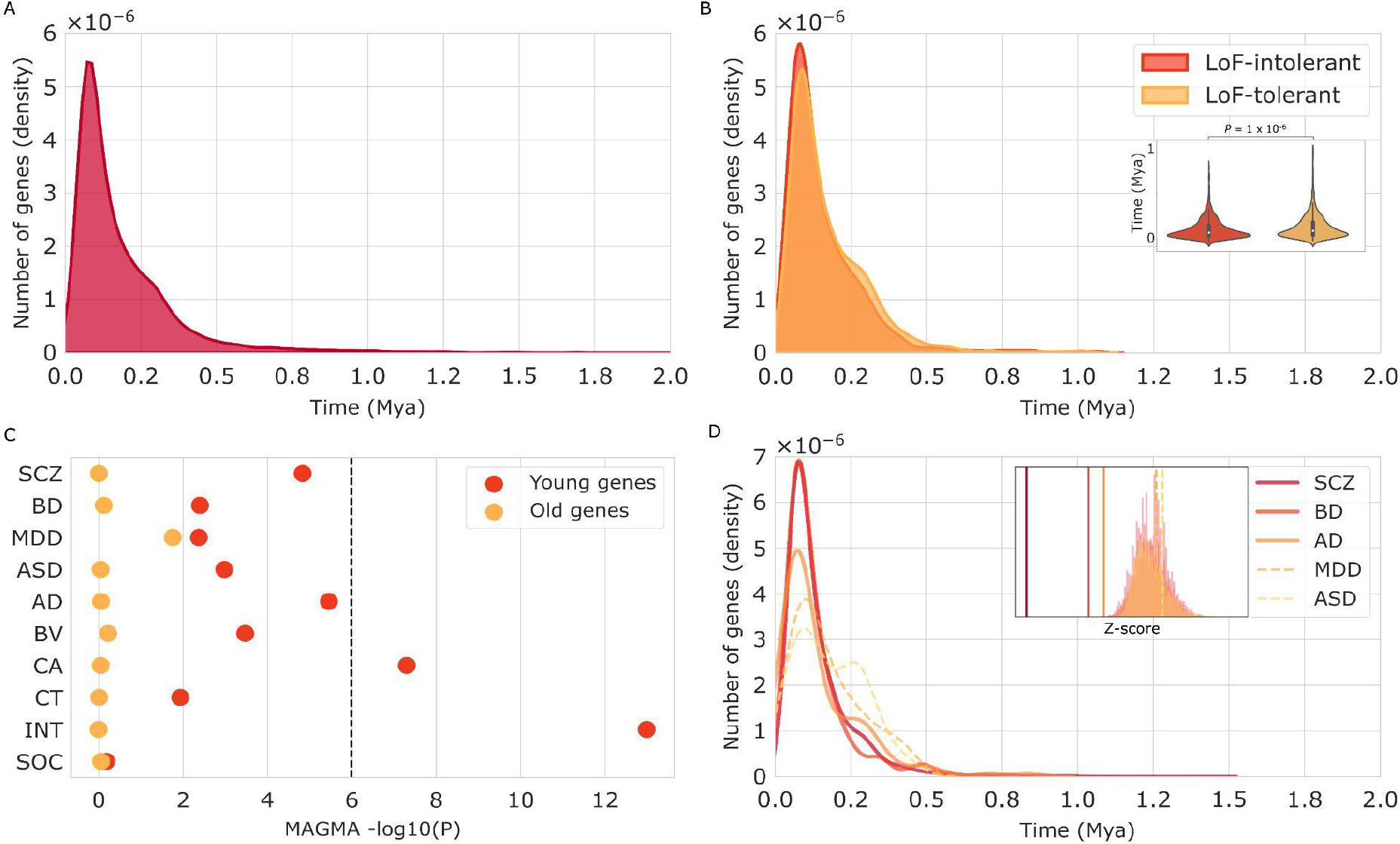
Genetic timeline of genes. *(a)* Timeline of genes (*n* = 18,328) with date estimates ranging from 2,965,600 to 3,803 years ago. Histogram depicts the density (y-axis) of the number of genes per age bin (x-axis; until 2M years ago is shown). *(b)* Timeline of the set of genes most intolerant to loss-of-function (LoF; orange) and the most tolerant to LoF (yellow); insertion (y-axis genes-age) shows LoF-intolerant genes (red) to be significantly younger (pair-sample t-test, *P* = 1 × 10^−6^) than LoF-tolerant genes (yellow). *(c)* Plots show *P* values (x-axis) of MAGMA gene-set analysis of the top oldest (yellow dots) and youngest (red dots) genes for enrichment of genes related to neuropsychiatric and brain phenotypes (y-axis). Dotted line indicates Bonferroni *P* value threshold < 2.5 × 10^−3^. Figure shows strong enrichment of the young genes for intelligence, and cortical area (also, AD and SCZ nominally significant). In contrast, the old genes are not found to be significantly enriched in disorders and/or brain aspects. *(d)* Timeline of genes associated with five major brain disorders, including SCZ, BD, AD, MDD, and ASD; solid lines denote phenotypes significantly different from the null model (Bonferroni *P* value threshold < 5 × 10^−3^, SCZ, BD, AD; see insert for the null model distribution of each condition). Mya, million years ago; LoF, loss-of-function; SCZ, schizophrenia; BD, bipolar disorder; AD, Alzheimer’s disease; MDD, major depressive disorder; ASD, autism spectrum disorder; BV, brain volume; CA, cortical area; CT, cortical thickness; INT, intelligence; SOC, sociability.

We next examined whether genes containing recent evolutionary modifications of the human genome are enriched for genes related to variation in brain (e.g. cortical area, thickness, and volume), cognition (e.g. intelligence, sociability), and neuropsychiatric disorders (Supplementary Methods; included studies in Table S6). Gene-set analysis^30^ (Methods) was used to examine the top 2,000 genes with the youngest evolutionary age (∼10% from all genes; ranging from ∼54,000 to ∼4,000 years ago) and their association with genes-sets related to variation in these ten phenotypes (using Bonferroni *P* value threshold < 2.5 × 10^−3^; Table S7). Genes containing recent mutations were found to be significantly enriched for genes involved in intelligence (*b* = 0.13, *P* = 2 × 10^−6^; using a lower or higher number of genes revealed similar findings, Supplementary Results) and cortical area (*b* = 0.08, *P* = 6.7 × 10^−4^). ‘Young genes’ were possibly also associated with Alzheimer’s disease (AD; *b* = 0.06, *P* = 4.2 × 10^−3^) and schizophrenia (SCZ; *b* = 0.06, *P* = 7.9 × 10^−3^; Figure 4C), but these effects did not survive our stringent Bonferroni correction. As a control condition, we also examined the group of genes with oldest evolutionary age estimates (ranging from ∼3M to ∼350,000 years ago) and no significant enrichment could be found (*P* > 0.05; Figure 4C, Table S7). A direct comparison of enrichment effect sizes confirmed that ‘young genes’ have a greater enrichment than ‘old genes’ for these phenotypes (Supplementary Results).

It was further tested the median evolutionary age of the set of genes associated with the same brain, cognitive, and neuropsychiatric phenotypes (*q* > 0.05, FDR; Supplementary Methods) compared to a null-model of randomly selected genes (ten phenotypes, Bonferroni *P* value threshold < 5 × 10^−3^; Methods). Genes related to intelligence were indeed found to display a younger evolutionary age than expected by chance (*n* = 2,102 genes; *P* = 6.4 × 10^−17^), as well genes related to SCZ (*n* = 2,275; *P* = 8.1 × 10^−10^), bipolar disorder (BD; *n* = 357; *P* = 5.5 × 10^−5^), sociability (*n* = 250; *P* = 1.4 × 10^−4^), brain volume (*n* = 170; *P* = 1.6 × 10^−4^), and AD (*n* = 143; *P* = 2.6 × 10^−3^), suggesting that genes linked to these phenotypes contain relatively more recent mutations in the human genome than compared to all genes (Figure 4D, Figure S8; sensitivity analysis in Supplementary Results). These findings corroborate theories that link neuropsychiatric conditions to recent human evolution^16–20^.

### Genes containing recent mutations are highly expressed in language areas

We further used gene expression data from the Allen Human Brain Atlas^31^ to investigate the transcriptomic pattern of the youngest genes. The top 1% genes containing the evolutionary most recent mutations (we opted for a more confined set of genes than the previous analysis to generate a null-model that evenly samples across the whole distribution; analyses with other thresholds in Supplementary Results) were found to be highly expressed in the gyrus pars triangularis *(P* = 6.2 × 10^−4^; Figure 5, Table S10; sensitivity analyses in Supplementary Results), a region recognised as Broca’s area and well known to be involved in speech and language, a human unique trait^32^. These genes were found to be more expressed in multimodal regions^33^ than primary regions (two-sided t-test, *t* = 2.5, *P* = 1.2 × 10^−2^), and a trend towards higher expression compared to unimodal regions (*t* = 2.1, *P* = 3.1 × 10^−2^; Bonferroni *P* value threshold 2.5 × 10^−2^). Expression levels in cortical regions of the genes containing the oldest variants were found to not be different from chance level (*P* > 0.05; Bonferroni *P* value threshold < 1.5 × 10^−3^).

**Figure 5.**
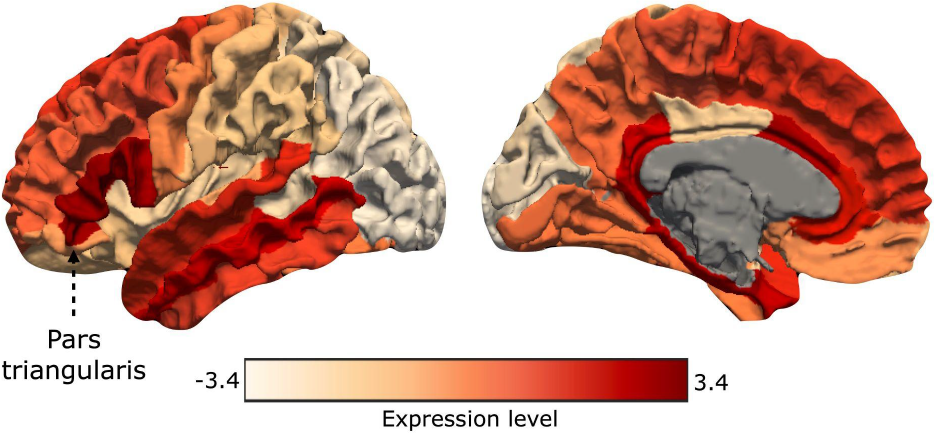
Transcriptomic brain map of evolutionarily recent genes. Brain plot of the normative expression levels of the top 1% genes with the youngest evolutionary age across brain areas (*n* = 164, ranging from 26,552 to 6,717 years ago), ranging from relatively low (white) to high (red) expression levels (z-scores). Figure shows high expression in Broca’s area (pars triangularis, a region part of Brodmann Area 45) a key brain area for language (*P* = 6.2 × 10^−4^, showing expression levels significantly exceeding the null-model of random genes expression, see Methods).

## Discussion

Mapping the evolutionary genetic timeline during human evolution reveals distinct timelines for traits related to human brain organisation and cognition. Traits related to neocortex, intelligence and psychiatric disorders are found to be relatively evolutionarily more recent, with particularly enrichment of variants related to these traits around a ‘young peak’ starting from ∼280,000 and continuing until ∼2,000 years ago (reaching the maximum point at ∼55,000 years ago). It is important to acknowledge that the analyses conducted in this study made use of all available GWAS data, of which the majority (91%) focused on European populations. Consequently, using these analyses to draw conclusions about the genetic timelines of phenotypes among different populations is not feasible.

The two observed ‘old’ and ‘young’ peaks line up with important paleontological observations. The old peak highlights the origin of *Homo erectus*^11^ and the divergence of *Homo sapiens* and Neanderthals^12^, a period displaying an enrichment for genetic variants associated with skeletal and metabolic phenotypes. The start of the young peak is consistent with studies pinpointing the African origin of modern humans ∼300,000 to ∼200,000 years ago^13,14^, marking a critical period of increased genetic diversification in the human genome reaching the maximum at ∼55,000 years ago, overlapping with the dispersion of *Homo sapiens* out of Africa (∼60,000 years ago)^34^, followed by the Upper Paleolithic period (∼45,000 years ago)^15^, and the beginning of agriculture (∼10,000 years ago)^35^. Comparative genetic analyses of archaic human ancestry revealed that a large number of adaptive changes in the human genome occurred around ∼600,000 and ∼200,000 years ago^36^, coinciding with the observed genetic timeline of the old and young peak from the human-phenotypic SNPs.

The Upper Paleolithic is considered to be a period in which important revolutions in cognition occurred^15^, characterised by the first consistent evidence of symbolic behaviour (i.e. art carvings, ritual artefacts, musical instruments), technological advancement (i.e. microlithic stone tools and bone artefacts, upgraded hunting tools), and ecological expansion (i.e. long-distance trading of materials)^37^. Our findings suggest an enrichment around this period for genetic variants associated with brain and neuropsychiatric phenotypes of modern-day humans. Consistent with these results, a previous study reported an enrichment for biological processes related to postsynaptic membrane in genes associated with modern-specific human variants arising within the last ∼300,000 years ago^38^, which also aligns with our observation of an enrichment of neuronal processes in genes exhibiting evolutionary recent age (Table S8).

The genetic timeline of neocortical organisation of modern humans presents an enrichment of variants that emerged recently in evolution (Figure 3A). Genetic modifications are widely believed to have played a major role in the growth and remodelling of the neocortex in humans^7^. The human brain has undergone major changes in size and structure since the last common ancestor shared by modern humans and other primates^39–41^, with the neocortex being the most expanded region^3^. These genetic and neocortical modifications potentially set humans apart from other primates^3–5^, enabling higher cognitive abilities such as complex language, social cognition, and higher-order thinking^40,42^. Important to mention in this context is that the studied SNPs are linked to the variation of brain size in modern-day humans and thus not necessarily to an expansion of the brain per se. Indeed, human brain size has not expanded continuously in history, with evidence it has even been decreasing in the last ∼3,000 years ago^43^.

One of the arguably most distinctive features between humans and other apes is our linguistic capacity^32^, with the human neocortex particularly expanded around language-related areas (such as Brodmann areas 44 and 45) as well as an expanded arcuate fasciculus pathway interconnecting language-related areas^32,44^. Our findings suggest that genes containing evolutionary recent variants are expressed in frontal cortical areas (Figure 5), most prominently in the gyrus pars triangularis - a key region within Broca’s area and central to language processing and speech production^32^. These findings align with comparative transcriptomic studies indicating that multimodal and language areas of the human brain are enriched for genes that have undergone accelerated divergence during the human lineage (known as HAR genes)^5^. Our findings also converge with evidence of selection pressures acting on alleles involved in variation of surface area of the inferior frontal cortex^45^.

Genetic modifications have driven the development of advanced cognitive skills^4,46^ through adaptations in brain areas and circuits^47^. These genetic and neuronal changes may have also rendered the human brain vulnerable to dysfunction^19,20,48^, potentially shaping certain brain disorders to be unique or specific to humans^16,18,20^. For instance, we observe that genetic variants related to the nervous system emerged before than variants underlying cognition (two-sided t-test, *t* = 3.95, *P* = 7.9 × 10^−5^) and psychiatric disorders (*t* = 8.91, *P* = 6.9 × 10^−19^; Figure S6). Previous studies have found that genes in human accelerated areas of the genome have been noted to be enriched for neurodevelopmental disorders^19^; and the strong genetic basis and human-specific character of schizophrenia have led to evolutionary theories that genetic adaptations of the human brain (related to the emergence of language) have contributed to vulnerability to the disorder^16,48^. Our findings provide several lines of supporting evidence for such theories. First, recent genetic modifications in the modern-day human genome reveal a particular enrichment for psychiatric traits (e.g. depression, alcohol intake; see Figure 2 and Table S1), in line with recent evidence showing that introgressed variants from Neanderthals (variants relatively recent in human evolution, ∼50,000 years ago) are linked to smoking, alcohol consumption, and mood-related traits^49^; and with findings of an overrepresentation of alleles conferring risk for mood-related traits specifically in the genomes of ancient farmers (∼11,000 years old), but not in the genomes of evolutionarily earlier hunter-gatherer^50^. Second, genetic variants associated with intelligence, educational attainment, hippocampal volume, and psychiatric disorders appear to have a more recent evolutionary age than expected considering their polygenicity and MAF, suggesting the potential action of selection pressures on these phenotypes (Supplementary Results, Table S2). Third, genes intolerant to LoF variants tend to encompass more recently evolved mutations than genes tolerant to LoF (Figure 4B). Fourth, genes containing relatively recent modifications are particularly associated with genes involved in intelligence and neocortical surface area (Figure 4C), aspects exhibiting recent genetic selection^45^ and often reported to be affected in the context of psychiatric and neurological conditions^51^. Fifth, genes related to neuropsychiatric conditions (SCZ, BD, and AD) contain more recent genetic modifications compared to other genes (Figure 4D).

Mapping the evolutionary genetic timeline of human-characteristic traits has inherent methodological limitations. To our knowledge, there is scarce available data on comprehensive dating estimates of the emergence of SNPs in the human genome, constraining the replication of the presented results with alternative resources. In this study, we utilised all available SNPs date estimates. Dating precision varies among genetic variants, with certain SNPs demonstrating higher precision than others (e.g., lower confidence interval). We carried out sensitivity analyses focusing on genetic variants with the highest dating precision, which supported our primary findings (see Supplementary Results). One of our strategies involved matching SNP dating estimates with GWAS results. It is important to point out that GWAS results largely target common variants. This introduces a bias into our evolutionary age estimation of human phenotypes because it neglects rare variants, which generally appear more recently in evolution, and entirely fixed variants, which tend to be older. Consequently, the median evolutionary age of traits should be approached with caution, and also take into account the lower and upper bounds of the evolutionary age (see Table S1 and Table S2). Our null models are designed by permuting human-phenotypic SNPs, ensuring that our statistical analyses account for this potential bias. In light of this, the current study focuses on human phenotypes that are still evolving, as opposed to traits that are fully evolved and have become fixed in the human genome. This is particularly relevant as human-phenotypic SNPs are associated with modern human trait variation, and it remains an open question whether these same SNPs were associated with similar traits in ancient hominins or other animals. Furthermore, the human evolutionary milestones were set side-by-side with the time distribution of genetic variants, which does not constitute causal evidence of forces shaping the genome.

## Methods

### Human Dating Genome Data

Human Dating Genome^9^ (HDG) data was downloaded from https://human.genome.dating/. HDG was developed by inferring the time of the most recent common ancestor between individual genomes, using two nonparametric approaches to estimate the date of origin of genetic variants by means of a recombination clock and mutation clock^9^. The HDG consisted of 13,689,983 SNPs of the human genome mapped across 22 chromosomes covering ∼200,000 generations with dates ranging from 5,140,625 to 87.5 years ago (one generation was defined as 25 years), with no assumptions about the demographic or selective processes that shaped the underlying genealogy. The present study used the median age estimates from both clocks combined on the two large-scale sequencing datasets.

### Human-phenotypic SNPs

Significant lead SNPs were selected from a wide range of human GWAS (GWAS Atlas^8^, https://atlas.ctglab.nl, access date June 2021). In total, date estimates of unique 36,506 SNPs across 2,538 GWAS from human phenotypes were obtained (ranging from 4,556,425 to 1,681 years ago; Supplementary Methods). The genetic timeline was computed by counting the number of human-phenotypic SNPs per time-bin; SNP density per bin was computed by dividing SNP count per bin by the total number across all time-bins.

### BRAIN-SNPs

A set of 6,284 SNPs (2,476 unique SNPs; BRAIN-SNPs) significantly associated across 1,744 brain phenotypes was derived based on the extensive UK Biobank BIG40^23^ GWAS (*P* < 5 × 10^−8^, discovery sample; https://open.win.ox.ac.uk/ukbiobank/big40/ SNPs were grouped according to brain structure (see Supplementary Methods and Table S3).

### Statistical inference

Permutation testing was used to statistically test the median evolutionary age of a set of SNPs or genes associated with a phenotype. A null-model was built by computing the median evolutionary age of equally sized random SNPs (or genes) at 10,000 iterations, assigning a z-score across iterations and a matching *P* value to the effect of interest (description of other permutation tests in Supplementary Methods). Sensitivity analysis was conducted controlling for both the number of SNPs and their corresponding MAF (Supplementary Methods).

### Genes evolutionary age estimation

The genetic timeline of genes was computed by taking the median evolutionary age of the SNPs located within a gene, defined by the SNPs coordinate that is within the start and stop position of the gene (GRCh37; sensitivity analysis of 1-kb window both sides resulted in similar results). SNPs located outside of genes or assigned to two or more genes were excluded. Median evolutionary age estimates were obtained for 18,328 genes (out of 26,836 genes) with at least one SNP with a date estimate (median of ∼100 SNPs within genes) ranging from 2,965,600 to 3,800 years ago. The length of genes exhibited a weak correlation with the median evolutionary age of genes (*r* = -0.058, *P* = 2.3 × 10^−15^), suggesting that the gene length was not a confounder in the estimation of the evolutionary age.

### Gene-set analysis

MAGMA competitive gene-set analysis^30^ was performed (one-sided positive direction) on the set of the top 2,000 genes (∼10% of all genes) with youngest/oldest median evolutionary age, testing for the enrichment of effects on the gene-based statistics (details of gene analysis in Supplementary Methods) for five brain disorders [SCZ, BD, AD, major depressive disorder (MDD), and autism spectrum disorder (ASD)], three brain (brain volume, thickness, and area) and two cognitive phenotypes (intelligence and sociability; ten phenotypes tested twice for each young/old gene-set, Bonferroni *P* value threshold < 2.5 × 10^−3^). Gene-set analysis was conditioned for gene size, LD levels between SNPs in genes, mean minor allele count in genes, and log values of these three variables.

## Data availability

HDG is available at https://human.genome.dating. GWAS Atlas is available at https://atlas.ctglab.nl. EBI Catalog is available at https://www.ebi.ac.uk/gwas. UK Biobank BIG40 is available at https://open.win.ox.ac.uk/ukbiobank/big40. Summary statistics of GWAS from brain disorders, intelligence, and brain volume are available at https://www.med.unc.edu/pgc/download-results; from cortical area and thickness are available at https://enigma.ini.usc.edu/research/download-enigma-gwas-results; and from sociability is available at https://www.repository.cam.ac.uk/handle/1810/277812. Cortical gene microarray transcriptome expression data from the Allen Human Brain Atlas is available at http://human.brain-map.org/static/download.

## Acknowledgments

M.P.v.d.H. is supported by a VIDI (452-16-015) grant from the Netherlands Organization for Scientific Research (NWO) and a European Research Council consolidator grant (ID 101001062). D.P. is supported by an NWO Gravitation project BRAINSCAPES: A Roadmap from Neurogenetics to Neurobiology (024.004.012) and a European Research Council advanced grant (ID 834057).

## Author Contributions

M.P.v.d.H. and I.L. conceived the project. I.L. analysed the data. M.P.v.d.H, and I.L. wrote the manuscript. Y.W., C.d.L., J.R., and D.P. provided expertise and feedback on the text and analyses.

## Notes

### Competing Interest Statement

The authors have declared no competing interest.

### Summary of Updates

New sensitivity analyses included; Figure 1 and 4 revised; Supplementary files updated.

