## Supplementary Information for "Mapping the genetic evolutionary timeline of human neural and cognitive traits"

**This file includes:**

Supplementary Methods and Supplementary Results

Supplementary Figure 1 to 8

Supplementary Tables 1 to 10 (Supplementary Table 1-3 in separate files)

### Supplementary Methods

#### Human-phenotypic SNPs

***Human Dating Genome Data.*** Human Dating Genome[1] (HDG) data was downloaded from https://human.genome.dating/. HDG was developed by inferring the time of the most recent common ancestor (TMRCA) between individual genomes, using two nonparametric approaches to estimate the date of origin of genetic variants by means of a recombination clock and mutation clock[1]. Combining information from the 1000 Genomes Project[2] (TGP) and the Simons Genome Diversity Project[3] (SGDP) resulted in 13,689,983 SNPs of the human genome mapped across 22 chromosomes with dates ranging from 5,140,625 to 87.5 years ago (90th/10th percentile = 1,108,530/17,739 years ago). This procedure makes no assumptions about the demographic or selective processes that shaped the underlying genealogy. The present study used the median evolutionary age estimates from both clocks combined on the two large-scale sequencing datasets.

***Human-phenotypic SNPs.*** The most significant lead SNPs within each genomic risk loci were selected from the summary statistics of GWAS from the GWAS Atlas[4] (access date June 2021) matching the HDG SNPs ID (GRCh37[5]). Genomic risk loci were defined as merged LD blocks of independent significant variants that belong to the same lead SNPs. Lead SNPs were identified by first clumping genome-wide significant SNPs ($P < 5 \times 10^{-8}$) with independent SNPs at $R^2 < 0.6$, and then clumping significant independent SNPs at $R^2 < 0.1$. In total, 67,119 SNPs (36,506 unique SNPs) related to modern human traits variation were selected from 2,538 GWAS, referred to in the main text as human-phenotypic SNPs (ranging from 4,556,425 to 1,681 years ago). Phenotypes in the GWAS Atlas are hierarchically grouped in domains ($n$ = 28), chapters ($n$ = 50), subchapters ($n$ = 172), and traits ($n$ = 2,251) levels.

***Human-phenotypic peaks identification.*** Peaks in the distribution of the normalised count (density) of human-phenotypic SNPs that appeared across time were identified, defined as any time-bin whose two direct neighbours have a smaller increment in SNPs (python package Scipy[6]). We retained the two peaks with the highest prominence (e.g. vertical distance between the peak and its lowest contour line). The peak with the second-highest prominence ('old peak') ranged from 2,440,467 to 280,006 years ago, reaching the maximum point at 1,099,491 years ago. The peak with the highest prominence ('young peak') ranged from 280,006 to 1,681 years ago, reaching the maximum point at 56,510 years ago.

**Sensitivity analyses**

***EBI human-phenotypic timeline.*** The genetic timeline of human-phenotypic SNPs was also computed for GWAS results of the EBI Catalog[7] (https://www.ebi.ac.uk/gwas) consisting of 4,237 GWAS detecting 88,362 non-duplicated SNPs. From the EBI Catalog it was selected SNPs not present in the GWAS Atlas to ensure that the similarity between the distribution of both databases was not driven by overlapping SNPs. The time dimension was discretized ($n$ = 100 bins) with equal time ranges (~45,500 years) and the number of SNPs per time-bin was counted for each dataset separately (Figure 1D). Spearman's correlation was computed to assess the similarity of the time distribution of emerging SNPs between GWAS Atlas and EBI Catalog resources. The analysis was repeated including SNPs more recent than 2M years ago (100 bins of ~20,000 years each) to accommodate the notion that the majority of human-phenotypic SNPs have an evolutionary age younger than 2M years ago (99.5%).

***MAF evolutionary age analysis.*** Minor allele frequencies (MAF) were calculated based on the Haplotype Reference Consortium panel[8] (GRCh37). MAF was rounded and divided into high MAF (MAF >= 0.4, n = 7,628 SNPs) and low MAF variants (MAF <= 0.1, n = 8,927 SNPs; Figure 1B). Permutation testing was used to evaluate whether the percentage of low or high MAF SNPs within a peak was significantly higher or lower. By randomly selecting the same number of high/low MAF SNPs out of the total human-phenotypic SNPs pool, the proportion of SNPs with an evolutionary age within the old/young peak was estimated. This procedure was repeated 10,000 times and assigned a z-score with a matching $P$ value to the effect of interest based on the obtained null-distribution of random effects.

***LD correction.*** Analyses were validated correcting for LD across all human-phenotypic SNPs by including only independent SNPs within LD blocks detected using PLINK[9] (v1.9) pruning function using --indep-pairwise (window size = 50, step size = 10, $R^2 < 0.1$) on the reference data Phase 3 of 1,000 Genomes[2] (European population), and by excluding SNPs within the MHC (chromosome 6, base pairs between 28477797 - 33448354, GRCh37).

**Genetic timeline of brain-imaging phenotypes**

***BRAIN-SNPs.*** GWAS summary statistics of the UK Biobank BIG40 study[10] were used to identify significant SNPs ($P < 5 \times 10^{-8}$, discovery sample) from 3,935 imaging-derived phenotypes. SNP ID was matched to the HDG[1] database resulting in 6,284 SNPs (BRAIN-SNPs) with date estimates (2,476 unique SNPs) from 1,744 imaging-derived phenotypes. Variants were grouped according to brain structures (neocortex, cerebellum, hippocampus, amygdala, thalamus, nucleus accumbens, caudate nucleus, pallidum, and white matter; see Table S3 for a complete list of phenotypes).

***BRAIN-SNPs timeline co-fluctuation.*** The temporal co-fluctuations of SNPs related to brain structures were assessed by discretizing the time dimension into 100 bins (~36,500 years each bin) and counting the number of SNPs that appeared within bins for each brain structure. Spearman's correlation evaluated the similarity between the genetic timeline of the SNPs related to brain structures (36 tests, Bonferroni $P$ value threshold < $1.4 \times 10^{-3}$; Figure 3B). Overlapping SNPs between two structures were excluded from the correlation. Because of the low frequency of SNPs appearing before 2M years ago, we repeated the same analysis including only SNPs younger than 2M years ago (see *BRAIN-SNPs timeline co-fluctuation validation* in Supplementary Results).

**Gene-level analysis**

***Genes evolutionary age estimation.*** The genetic timeline of genes was computed by taking the median evolutionary age of the SNPs located within a gene, defined by the SNPs coordinate that is within the start and stop position of the gene (GRCh37; sensitivity analysis of 1-kb window both sides resulted in similar results). We opted for this strategy to map SNPs to genes but we acknowledge that distant SNPs can still have an influence on genes[11]. SNPs located outside of genes or assigned to two or more genes were excluded (58% of all SNPs). Median evolutionary age estimates could be obtained for 18,328 genes (out of 26,836 genes) with at least one SNP with a date estimate (median of 100 SNPs within genes) ranging from 2,965,600 to 3,800 years ago (Figure 4A).

***LoF genes.*** The top 2,000 genes with respectively the highest and lowest Loss-of-Function (LoF) scores were extracted[12] and an independent two-sided t-test was used to compare the median evolutionary age estimates between the two groups of genes (Figure 4B). The same test was conducted with a varying number of top genes ($n$ = 500, 1,000 and 3,000 genes; see *Validation LoF gene analysis* in Supplementary Results)

***Gene-analysis.*** Gene-analysis (MAGMA[13] v1.09; Phase 3 of 1,000 Genomes, european population; GRCh37) was conducted on the raw GWAS summary statistics from schizophrenia (SCZ)[14], bipolar disorder (BD)[14], major depressive disorder (MDD)[15], autism spectrum disorder (ASD)[16], and Alzheimer's disease (AD)[17]; plus five major brain and cognitive phenotypes (brain volume[18], cortical area[19], cortical thickness[19], intelligence[20], and social behavior[21]) to identify the genes involved with these phenotypes (more details regarding included GWAS in Table S6). FDR-significant ($q < 0.05$) genes were included for further analysis.

***Functional gene annotation.*** Functional annotation of the 2,000 genes with the youngest median evolutionary age was conducted using FUMA[22], an integrative web-based platform for functional gene annotation based on multiple biological resources to facilitate functional annotation of GWAS results. FUMA provides gene-based, pathway, and tissue enrichment results. FUMA compares the prioritised genes against the genes-set obtained from MsigDB and Wikipathways, using hypergeometric tests to evaluate the overrepresentation of biological functions ($q < 0.05$, FDR).

***Cortical gene expression.*** Cortical gene microarray transcriptome expression data was taken from the Allen Human Brain Atlas (AHBA; http://human.brain-map.org/static/download), consisting of gene expression profiles from brain samples of six human donors (five males, one female) without a history of neuropsychiatric or neuropathological conditions. Data consisted of expression levels of 20,734 genes measured by 58,692 probes for each cortical region of the left hemisphere[23]. Tissue samples were spatially mapped to each of the cortical areas of the FreeSurfer Desikan Killiany atlas[24] (DK, $n$ = 34 left cortical areas) based on their distance to the nearest voxel within the cortical ribbon of MNI 152 template. Samples were z-normalised and averaged across regions, and then averaged across subjects obtaining a group-level gene expression matrix of 34 × 20,734 (for more details in the preprocessing pipeline, see[25]). For validation purposes, AHBA samples were also mapped using von Economo-Koskinas cortical type atlas[26] (EK, $n$ = 15 left cortical areas), Brodmann Atlas[26] (BA, $n$ = 39 left cortical areas), and 114-region DK subdivision[27] (DK-114, $n$ = 57 left cortical areas).

**Statistical testing**

***Genetic timeline.*** Permutation testing was used to statistically assess to what extent more (or alternatively, less) SNPs appeared within specific time periods. The genetic timeline of human-phenotypic SNPs (or BRAIN-SNPs) was discretized into 100 bins of equal periods and the count of the number of SNPs within each bin was extracted. The distribution of MAF scores differed between the total pool of SNPs from HDG and human-phenotypic SNPs (or BRAIN-SNPs). To control for this, it was randomly selected SNPs from the total HDG pool matching the MAF proportion from the human-phenotypic SNPs (or BRAIN-SNPs; eight MAF bins: 0 - 0.00001, 0.00001- 0.0001, 0.0001 - 0.001, 0.001 - 0.01, 0.01 - 0.1, 0.1 - 0.2, 0.2 - 0.3, 0.3 - 0.4, 0.4 - 0.5) resulting in ~3.2M SNPs (MAF-matched total pool). A null-model was built by randomly selecting 10,000 random sets of SNPs from the MAF-matched total pool of

SNPs. The null-distribution was used to assign each effect of interest a z-score and matching *P* value, indicating in which time-bin the trait of interest presented a significantly higher (or lower) number of SNPs (Bonferroni correction for the number of tests equal to the number of time-bins, *n* = 100). Because the majority of SNPs are younger than 2M years ago (99.5% of all human-phenotypic SNPs; 99.1% of all BRAIN-SNPs) for this analysis only SNPs younger than 2M years ago were included.

***Phenotype evolutionary age.*** The median evolutionary age of a group of SNPs (or genes) of interest was statistically assessed by repeating the computation of median evolutionary age for sets of equally sized random SNPs (same for gene analysis) at each iteration (*n* = 10,000) from the total human-phenotypic SNPs (or BRAIN-SNPs) pool. Sensitivity analysis controlling for both the number of SNPs and their corresponding MAF was conducted by randomly drawing the same number of SNPs within each MAF bin (0 - 0.1, 0.1 - 0.2, 0.2 - 0.3, 0.3 - 0.4, 0.4 - 0.5). The results were validated by correcting for LD across all SNPs of interest by only including independent SNPs within LD blocks (see *LD correction* above), including SNPs with a date quality score above 0.7, and GWAS with more than 50,000 subjects (for human-phenotypic SNPs).

***Phenotype within peak.*** Permutation testing was used to test whether the proportion of a group of variants that appeared within a peak was significantly higher (or lower). The same number of SNPs as the phenotype of interest was randomly drawn from the total pool of all human-phenotypic SNPs (10,000 times). The null-distribution was based on the percentage of random SNPs appearing within the old or young peak and assigned a z-score with a matching *P* value to the effect of interest, in this case the percentage of SNPs of the phenotype of interest appearing within old/young peak (Bonferroni correction for the number of phenotypes tested).

***Gene-set analysis permutation.*** It was assessed whether the difference in magnitude of enrichment of brain, cognitive, and neuropsychiatric phenotypes between the 2,000 genes with the evolutionary youngest and oldest age. The genes between the old and young genes-sets were shuffled (1,000 times), gene-set analysis was conducted for both shuffled genes-set testing for the enrichment of the mentioned phenotypes, estimated the absolute difference of their beta enrichment (effect size) between the shuffled genes-set, and assigned a z-score with a matching *P* value to the original beta enrichment difference (between old and young genes) based on the obtained null-distribution of random effects for each phenotype.

***Gene expression analysis.*** Permutation testing was used to statistically assess the median expression of the youngest 1% genes (*n* = 164) using GAMBA[25]. We opted for selecting the top 1% of genes (instead of the earlier used ~10% genes) with the lowest evolutionary age to generate a null-model that evenly samples across the whole distribution, still confined enough to be tested against the baseline (here, 16,344 genes expressed in brain tissue, see[25] for details). GAMBA null-random-gene model[25] was generated by randomly selecting 10,000 times the same number of genes as the original set (*n* = 164 out of 16,344 genes with both expression and dating estimates, mapped from 26,552 to 6,717 years ago), estimating the median genetic expression (extracted from AHBA) of the random genes for each brain region, z-normalised the null distribution of each brain region across iterations including the ground truth, and assigning a *P* value to the effects of interest. We used strict Bonferroni correction for the number of brain regions tested (*n* = 34 in the main analysis).

### Supplementary Results

#### Human-phenotypic SNPs

***Human-phenotypic timeline replication.*** To validate the distinctiveness of the bimodal distribution of human-phenotypic SNPs extracted from GWAS Atlas[4], this genetic timeline was compared with the distribution of the evolutionary age of SNPs associated with human phenotypes from the EBI Catalog[7] (88,362 unique SNPs non-overlapping with GWAS Atlas). The time dimension of both datasets was binned (*n* = 100) and counted the number of SNPs that appeared within each time-bin, then assessed the similarity between the count SNPs distribution of both datasets by means of a Spearman's correlation. The bimodal distribution of the GWAS Atlas and EBI Catalog was almost identical (*rho* = 0.96, $P = 8.9 \times 10^{-59}$; Figure 1D). Because of the low frequency of SNPs appearing before 2M years ago, we conducted the same analysis, including SNPs younger than 2M years ago (*rho* = 0.98, $P = 1.5 \times 10^{-72}$).

***MAF evolutionary age analysis.*** We specifically examined the genetic timeline of high and low MAF variants because of their dominant involvement in human traits and human disorders[4]. The MAF of human-phenotypic SNPs score was divided into two SNP groups, high MAF (MAF >= 0.4, *n* = 7,628 SNPs) and low MAF variants (MAF <= 0.1, *n* = 8,927 SNPs), and evaluated the proportion of high/low MAF variants appearing within old/young peaks. The majority of high MAF variants of human-phenotypic SNPs were found to be present in the old peak (84% of all high MAF variants; $P = 7.1 \times 10^{-259}$), while low MAF variants were found to be overrepresented in the young peak (63% of all low MAF variants, $P = 10^{-324}$; Figure 1B).

**Phenotype evolutionary age**

***Chapter level.*** We continued by examining the human-phenotypic SNPs linked to modern-day human traits in more detail (Supplementary Methods). We assessed whether the evolutionary age of phenotypes at the chapter level were statistically older or younger (for the domain level, see *Psychiatric phenotypes influenced by evolutionary recent mutations* in Results). At the chapter level ($n$ = 50 chapters, Bonferroni P value threshold < 1 x $10^{-3}$; Figure S3), SNPs related to 'Mental and Behavioral Disorders' ($n$ = 2,331 SNPs, median evolutionary age = 111,560 years old, $P$ = 3.3 x $10^{-156}$), followed by 'Eye, Ear, and Related Structures' ($n$ = 756, median age = 26,744, $P$ = 2.8 x $10^{-70}$), and 'Self-Care' ($n$ = 2,612, median age= 487,959, $P$ = 4.2 x $10^{-22}$) were significantly younger. The top three oldest chapters were 'Functions of the Digestive, Metabolic and Endocrine Systems' ($n$ = 10,300, median age = 780,269, $P$ = 1.9 x $10^{-36}$), 'Functions of the Cardiovascular, Hematological, Immunological and Respiratory Systems' ($n$ = 6,844, median age = 753,774, P = 2.8 x $10^{-13}$) and 'Structures Related to Movement' ($n$ = 4,302, median age = 761,628, $P$ = 7.7 x $10^{-10}$; see Table S1 for complete chapter results).

We repeated this analysis controlling for both the number of genetic variants and their corresponding MAF. SNPs involved with 'Mental and Behavioral Disorders' ($n$ = 1,893 SNPs, median evolutionary age = 83,250 years old, median MAF = 0.047, $P$ = 5.7 x $10^{-67}$) and 'Major Life Areas' ($n$ = 817, median age = 617,780, median MAF = 0.281, $P$ = 7.3 x $10^{-5}$) phenotypes were significantly younger. The oldest chapters were 'Malignant Neoplasms' ($n$ = 363, median age = 590,657, median MAF = 0.042, $P$ = 6.4 x $10^{-6}$), 'Functions of the Digestive, Metabolic and Endocrine Systems' ($n$ = 10,147, median age =785,400, median MAF = 0.290, $P$ = 1.6 x $10^{-4}$; see Table S2 for complete chapter results).

***Subchapter level.*** At the subchapter level ($n$ = 172 subchapters, Bonferroni P value threshold < 2.9 x $10^{-4}$; Figure S4), the class 'Structure of Eyeball' ($n$ = 756 SNPs, median evolutionary age = 26,744 years old, $P$ = 2.1 x $10^{-70}$), 'Depressive Episode' ($n$ = 745, median age = 27,526, P = 3.6 x $10^{-70}$), and 'Mental and Behavioral Disorders Due to Use of Alcohol' ($n$ = 715, median age = 72,959, P = 5.3 x $10^{-61}$) ranked as the top 3 youngest subchapters. Subchapters 'Weight Maintenance Functions' ($n$ = 7,813, median age = 802,540, $P$ = 1.6 x $10^{-37}$), 'Height' ($n$ = 3,284, median age = 766,455, $P$ = 4.3 x $10^{-8}$), and Immunological System Functions ($n$ = 4,197, median age = 747,780, $P$ = 5.5 x $10^{-7}$) were ranked as the top 3 oldest subchapters (see Table S1 for complete subchapters results).

When controlling for both number of genetic variants and their corresponding MAF, the class 'Mental and Behavioural Disorders Due to Use of Alcohol' ($n$ = 481 SNPs, median evolutionary age = 39,548 years old, median MAF = 0.010, $P$ = 5.2 x $10^{-10}$), 'Sexual Functions' ($n$ = 349, median age = 76,814, median MAF = 0.016, $P$ = 7.1 x $10^{-10}$), 'Education' (*n* = 578, median age = 637,646, median MAF = 0.306, $P = 2 \times 10^{-5}$), 'Looking After One's Health' (*n* = 1,359, median age = 646,077, median MAF = 0.294, $P = 2.2 \times 10^{-5}$), and 'Depressive Episode' (*n* = 658, median age = 24,470, median MAF = 0.004, $P = 1.7 \times 10^{-4}$) ranked as the top youngest subchapters. Subchapters 'Malignant Neoplasms of Breast' (*n* = 215, median age = 487,202, median MAF = 0.026, $P = 2 \times 10^{-16}$), and 'General Metabolic Functions' (*n* = 1043, median age = 667,462, median MAF = 0.140, $P = 2.1 \times 10^{-4}$) were ranked as the oldest subchapters (see Table S2 for complete subchapters results).

***Trait level.*** Zooming into the individual trait level (*n* = 2,251 traits, Bonferroni *P* value threshold < $2.2 \times 10^{-5}$; Figure 2 and Figure S5) revealed that the eye-related trait 'Vertical cup-disc ratio' (*n* = 515 SNPs, median evolutionary age = 23,661 years old, $P = 3.9 \times 10^{-49}$) have a significantly younger age, followed by 'Depression-lifetime number of depressed periods' (*n* = 395, median age = 21,076, $P = 2.1 \times 10^{-38}$), 'Lifetime number of sexual partners' (*n* = 337, median age = 32,349, $P = 4.4 \times 10^{-33}$), and 'Average weekly intake of other alcoholic drinks' (*n* = 281, median age = 20,258, $P = 1.7 \times 10^{-28}$). In contrast, the oldest traits included 'Height' (*n* = 1,693, median age = 811,305, $P = 9.1 \times 10^{-9}$) and 'Body Mass Index' (*n* = 1,671, median age= 794,265, $P = 6.9 \times 10^{-7}$; see Table S1 for complete traits results).

The same analysis at the trait level was repeated controlling for both the number of genetic variants and their corresponding MAF. We found that only 'Use of sun/uv protection' was significantly younger after Bonferroni correction (*n* = 28 SNPs, median evolutionary age = 185,084 years old, median MAF = 0.323, $P = 1.9 \times 10^{-5}$). We also observed a trend toward a younger evolutionary age (nominally significant) in phenotypes related to brain, cognition and psychiatric disorders such as 'Educational attainment' (*n* = 465, median age = 637,522, median MAF = 0.306, $P = 1.1 \times 10^{-4}$), 'Fluid intelligence score' (*n* = 47, median age = 459,465, median MAF = 0.386, $P = 1.7 \times 10^{-4}$), 'Schizophrenia/Bipolar disorder' (*n* = 86, median age = 499,765, median MAF = 0.328, $P = 2.9 \times 10^{-4}$), 'Depression - Lifetime number of depressed periods' (*n* = 352, median age = 19,938, median MAF = 0.004, $P = 1.1 \times 10^{-3}$), 'Left hippocampus' (*n* = 8, median age = 76,249, median MAF = 0.348, $P = 9.4 \times 10^{-3}$), 'Average weekly intake of other alcoholic drinks' (*n* = 252, median age = 19,506, median MAF = 0.005, $P = 9.6 \times 10^{-3}$), within others. Sex-specific metabolic traits as well presented nominally significant effects towards younger evolutionary age, including 'Legs-leg fat ratio (female)' (*n* = 143, median age = 506,280, median MAF = 0.284, $P = 1.1 \times 10^{-3}$), 'Hip circumference (female)' (*n* = 33, median age = 382,215, median MAF = 0.354, $P = 1.9 \times 10^{-3}$), and 'Body Mass Index (male)' (*n* = 362, median age = 675,207, median MAF = 0.317, $P = 7.8 \times 10^{-3}$). The top traits with the (nominally significant) oldest age were 'Age at last live birth (female)' (*n* = 1, median age = 2,660,000, median MAF = 0.448, $P = 3.4 \times 10^{-4}$), 'Ulcerative colitis' (*n* = 213, median age = 595,307, median MAF = 0.024, $P = 5.7 \times 10^{-4}$), and

'Ease of skin tanning' (*n* = 319, median age = 628,330, median MAF = 0.086, $P = 2.6 \times 10^{-3}$; see Table S2 for complete traits results).

***Validation phenotype evolutionary age.*** We performed the same analysis at the trait level now controlling for SNPs group size differences. By resampling (10,000 times) a fixed number of SNPs (*n* = 50) from each trait and computing the median evolutionary age across the iterations, it was compared with a null-distribution of the median evolutionary age of (10,000 times) randomly selected 50 SNPs from the total human-phenotypic SNPs pool (only traits with more than 50 SNPs were included, *n* = 263 traits; Bonferroni *P* value threshold < $1.9 \times 10^{-4}$). The traits with a significantly younger age were 'Average weekly intake of other alcoholic drinks' (median evolutionary age = 20,257 years old, $P = 2.1 \times 10^{-6}$), followed by 'Depression - Lifetime number of depressed periods' (median age = 21,076, $P = 2.1 \times 10^{-6}$), and 'Vertical cup-disc ratio' (median age = 23,660, $P = 2.6 \times 10^{-6}$). No trait had a median evolutionary age significantly older. These results confirmed that the youngest traits are robust for controlling the number of SNPs.

To ensure that the main results were robust against differences across included studies or SNPs, we conducted the same analysis at the domain and trait level now applying three filtering procedures simultaneously: 1) including GWAS with a sample size above 50,000 subjects; 2) including only independent SNPs in LD blocks ($R^2 < 0.1$) and SNPs outside the MHC (see *LD correction* in Supplementary Methods); 3) including SNPs with a date quality score above 0.7, a metric ranging from 0 (low) to 1 (high), provided by the HDG database, indicating the accuracy of the SNPs date estimate. We found that out of all included domains (*n* = 22 domains, Bonferroni *P* value threshold < $2.2 \times 10^{-3}$), only 'Psychiatric' had a median age significantly younger (*n* = 581 SNPs, median evolutionary age = 24,116 years old, $P = 2 \times 10^{-7}$). The domains with significantly older SNPs were 'Metabolic' (*n* = 475, median age = 79,258, $P = 5 \times 10^{-19}$), 'Endocrine' (*n* = 39, median age = 220,504, $P = 1.4 \times 10^{-18}$), 'Skeletal' (*n* = 208, median age = 84,103, $P = 3.3 \times 10^{-10}$), and 'Immunological' (*n* = 218, median age = 63,561, $P = 1.9 \times 10^{-3}$). At the trait level (*n* = 492 traits, Bonferroni *P* value threshold < $1 \times 10^{-4}$), there was a trend towards a younger evolutionary age for 'Depression - Lifetime number of depressed periods' (*n* = 220, median age = 18,582, $P = 1.1 \times 10^{-4}$), followed by 'Average weekly intake of other alcoholic drinks' (*n* = 165, median age = 18,078, $P = 6.9 \times 10^{-4}$), and 'Lifetime number of sexual partners' (*n* = 125, median age = 18,378, $P = 3.1 \times 10^{-3}$) but none of these effects were significant after Bonferroni correction. 'Height' (*n* = 60, median age = 440,853, $P = 5 \times 10^{-188}$) was significantly older, followed by 'Waist-hip ratio' (*n* = 17, median age = 1,013,918, $P = 1.8 \times 10^{-49}$), and 'Type 2 Diabetes' (*n* = 25, median age = 582,173, $P = 8.4 \times 10^{-47}$).

***Phenotype within peak.*** We validated to what extent the top three 'oldest' and 'youngest' traits were potentially over- or underrepresented within the old or young peak (six phenotypes tested twice for each peak, Bonferroni $P$ value threshold < 8 x $10^{-3}$; see *Phenotype within peak* in Supplementary Methods). From all the SNPs linked to 'Vertical cup-disc ratio', 18% were present within the old peak ($P$ = 5.7 x $10^{-120}$) and 82% within the young peak ($P$ = 6.3 x $10^{-120}$). Similarly, SNPs related to 'Depression - Lifetime number of depressed periods' were underrepresented in the old peak (11% out of all SNPs, $P$ = 1.6 x $10^{-117}$) and overrepresented in the young peak (89% of all SNPs, $P$ = 2.3 x $10^{-121}$). SNPs related to 'Average weekly intake of other alcoholic drinks' (13% out of all SNPs, $P$ = 5.9 x $10^{-77}$) and 'Lifetime number of sexual partners' (20% out of all SNPs, $P$ = 3.7 x $10^{-73}$) were also underrepresented in the old peak and overrepresented in the young peak (87% and 80% out of all SNPs, respectively; $P$ = 7.6 x $10^{-81}$ and 3 x $10^{-74}$, respectively). In contrast, SNPs related to 'Height' (76% out of all SNPs, $P$ = 6.1 x $10^{-15}$) and 'Body Mass Index' (73% out of all SNPs, $P$ = 6.4 x $10^{-9}$) were overrepresented in the old peak and underrepresented in the young peak (24% and 27% out of all SNPs, respectively; $P$ = 1.1 x $10^{-14}$ and 5.3 x $10^{-9}$, respectively).

**Genetic timeline of brain-imaging phenotypes**

***BRAIN-SNPs.*** We continued by examining the genetic timeline of SNPs related to brain phenotypes. We started with a total subset of 6,284 SNPs (2,476 unique SNPs; referred to as BRAIN-SNPs), collated from a recently performed extensive GWAS on 1,744 neuroimaging-derived phenotypes from the UK Biobank BIG40[10] (e.g. brain size, cortical surface, white matter volume, myelin, volume of specific brain structures; see Table S3 for a complete list of phenotypes). The genetic timeline of BRAIN-SNPs ranged from 3,621,625 until 4,457 years ago (median evolutionary age = 28,652 years old) and again showed a bimodal distribution with an old peak from 2,645,319 until 375,469 years ago and a young peak from 375,469 until 4,457 years ago (Figure S7).

***BRAIN-SNPs timeline.*** The genetic timeline of BRAIN-SNPs ($n$ = 100 bins of ~20,000 years, Bonferroni $P$ value threshold < 5 x $10^{-4}$; see *Genetic timeline* in Supplementary Methods) presented four time periods with a significantly higher number of SNPs. Two time-bins had significantly more SNPs at the beginning of the old peak from 1,980,075 to 1,960,150 ($P$ = 2.9 x $10^{-6}$), and 1,880,475 to 1,860,550 ($P$ = 7.4 x $10^{-5}$) years ago; and another two time-bins matching the highest point of the young peak from 47,625 to 27,700 years ago ($P$ = 1.4 x $10^{-8}$), and from 27,700 to 7,775 ($P$ = 2.1 x $10^{-21}$; Figure S7).

***BRAIN-SNPs evolutionary age.*** We examined genetic variants associated with key structures of the brain, such as the neocortex (*n* = 130 SNPs), white matter (*n* = 676), cerebellum (*n* = 144), amygdala (*n* = 39), hippocampus (*n* = 30) and thalamus (*n* = 19) among others (nine brain structures listed in Table S3, Bonferroni *P* value threshold < 5.5 x $10^{-3}$; see *Phenotype evolutionary age* in Supplementary Methods). SNPs related to the variation of amygdala and neocortex phenotypes present the youngest median evolutionary age across all examined brain structures (median evolutionary age of 332,998 and 400,170 years old, respectively; Figure 3A), while SNPs related to caudate nucleus, thalamus, and nucleus accumbens showed the oldest age (median age of 1,098,163, 815,658 and 965,115 years old, respectively). Only Neocortex SNPs display a median evolutionary age significantly younger (*P* = 4.4 x $10^{-4}$; see Table S4 for all *P* values). We find similar results controlling for LD dependence between SNPs (see *Validation BRAIN-SNPs evolutionary age* below).

***Validation BRAIN-SNPs evolutionary age.*** We analysed the expected median evolutionary age of BRAIN-SNP under a null-model controlling for LD by including only independent SNPs within LD blocks ($R^2$ < 0.1) and SNPs outside MHC. Only neocortex SNPs (*n* = 32, median evolutionary age = 46,070 years old) showed a trend towards a younger age (*P* = 3.2 x $10^{-2}$) however this effect was no longer significant after Bonferroni correction (*P* < 5.5 x $10^{-3}$).

***BRAIN-SNPs timeline co-fluctuation.*** We further asked whether the BRAIN-SNPs associated with the neocortex variation presented a similar or distinct genetic timeline compared to SNPs associated with other brain structures[28,29]. The genetic timeline of BRAIN-SNPs related to neocortex variation (*n* = 100 bins; 36 tests, Bonferroni *P* value threshold < 1.4 x $10^{-3}$; excluded SNPs overlapping between structures; see *BRAIN-SNPs timeline co-fluctuation* in Supplementary Methods) showed the strongest correlation with the genetic timeline of the white matter (Spearman's *rho* = 0.72, *P* = 3.9 x $10^{-17}$; Figure 3B). The genetic timeline of neocortical SNPs also presented significant overlap with the genetic timeline of the cerebellum (*rho* = 0.70, *P* = 7.1 x $10^{-16}$) and hippocampus (*rho* = 0.52, *P* = 3.7 x $10^{-8}$). Neocortical SNPs exhibited an independent pattern from the genetic timeline of SNPs related to the thalamus, as expressed by a relatively non-significant low correlation (*rho* = 0.28, *P* = 4 x $10^{-3}$), supportive of evidence of a distinct evolutionary pattern between cortical and subcortical areas[28,30]. The SNPs linked to cerebellum variation significantly co-fluctuated with all other brain structures[29] with correlations ranging from 0.37 (thalamus, *P* = 1.4 x $10^{-4}$) to 0.80 (white matter, *P* = 5.3 x $10^{-24}$; see Table S5 for all correlations).

***Validation BRAIN-SNPs timeline co-fluctuation.*** We evaluated the co-fluctuation of the number of SNPs appearing across time between each brain structure (bins = 100; 36 tests, Bonferroni *P* value threshold < $1.4 \times 10^{-3}$). To validate the main results, we conducted the same analysis including SNPs younger than 2M years ago. SNPs related to neocortex correlated the highest with cerebellum (*rho* = 0.36, *P* = $2.7 \times 10^{-4}$) and white matter (*rho* = 0.32, *P* = $1.2 \times 10^{-3}$), and the lowest non-significant correlations with pallidum (*rho* = 0.04, *P* > 0.05), nucleus accumbens (*rho* = 0.16, *P* > 0.05), and thalamus (*rho* = 0.19, *P* > 0.05). The genetic timeline of cerebellum SNPs was significantly correlated with white matter (*rho* = 0.54, *P* = $8.8 \times 10^{-9}$) and hippocampus (*rho* = 0.36, *P* = $1.9 \times 10^{-4}$); amygdala correlated with white matter (*rho* = 0.33, *P* = $9.1 \times 10^{-4}$); and nucleus accumbens correlated with pallidum (*rho* = 0.37, *P* = $1.8 \times 10^{-4}$).

**Gene-level analysis**

***Validation LoF gene analysis***. We conducted an independent t-test to assess the median evolutionary age difference between the most intolerant and tolerant LoF genes[12] across varying sizes of gene-sets. The top 500 intolerant LoF genes were younger than the top 500 tolerant LoF (*t* = -3.30, *P* = $9.9 \times 10^{-4}$), same for the top 1,000 (*t* = -3.96, *P* = $7.7 \times 10^{-5}$) and 3,000 LoF genes (*t* = -5.14, *P* = $2.8 \times 10^{-7}$; Bonferroni *P* value threshold < $1.6 \times 10^{-2}$).

***Validation gene-set analysis.*** We repeated the gene-set analysis (see *Gene-set analysis* in Methods and Table S6 for included GWAS) with MAGMA[13] varying the size of the oldest and youngest genes-set and test for enrichment. We found that the youngest 1,000 genes were enriched for genes related to intelligence (*b* = 0.14, *P* = $1.9 \times 10^{-4}$) and SCZ (*b* = 0.11, *P* = $3.2 \times 10^{-3}$). No enrichment was found for genes within the old peak for any phenotype (*P* > 0.05).

We conducted another gene-set analysis with the top 3,000 oldest and youngest genes. The youngest genes showed again an enrichment for genes related to intelligence (*b* = 0.14, *P* = $1.4 \times 10^{-10}$), SCZ (*b* = 0.08, *P* = $3 \times 10^{-4}$), and cortical thickness (*b* = 0.06, *P* = $2.1 \times 10^{-3}$). No enrichment was found for genes within the old peak for any phenotype (*P* > 0.05).

***Gene-set analysis permutation.*** Using a permutation test, we statistically assessed the difference in magnitude between the enrichment of brain, cognitive, and neuropsychiatric phenotypes in the 2,000 genes with the evolutionary youngest and oldest age (ten tests, Bonferroni *P* value threshold < $5 \times 10^{-3}$; see *Gene-set analysis permutation* in Supplementary Methods). The top young genes showed a significantly larger beta than old genes for the enrichment of genes related to intelligence (*P* = $5.9 \times 10^{-18}$), cortical area (*P* = $1 \times 10^{-5}$), SCZ (*P* = $4.3 \times 10^{-6}$), and AD (*P* = $1.5 \times 10^{-3}$).

***Validation phenotype evolutionary genes age.*** To control for the differences in the number of genes associated with a phenotype across different GWAS (identified by gene analysis), we further validated the main results including the 200 genes with the largest z-stat resulting from the MAGMA[13] gene-analysis for each GWAS (ten tests, Bonferroni *P* value threshold < $5 \times 10^{-3}$). The top genes associated with brain, cognitive and neuropsychiatric phenotypes that presented a evolutionary age significantly younger were intelligence (median evolutionary age = 85,690 years old, $P = 2.0 \times 10^{-4}$), brain volume ($P = 1.7 \times 10^{-4}$), sociability (median age = 86,019, $P = 2.4 \times 10^{-4}$), SCZ (median age = 88,656, $P = 1.2 \times 10^{-3}$) and AD (median age = 91,552, $P = 2 \times 10^{-3}$).

We further validated the main analysis by including genes with more than ten SNPs and a median date quality score above 0.7 ($n$ = 14,523 genes). Once again, the sets of genes with a significantly younger evolutionary age were intelligence (median evolutionary age = 99,160 years old, $P = 1.1 \times 10^{-17}$), SCZ (median age = 101,513, $P = 1.6 \times 10^{-15}$), BD ($P = 6.5 \times 10^{-5}$), brain volume (median age = 83,882, $P = 9.1 \times 10^{-5}$), sociability (median age = 89,978, $P = 1.5 \times 10^{-4}$), and AD (median age = 88,650, $P = 2.5 \times 10^{-3}$).

***Functional gene annotation.*** We performed functional gene annotation on the set of 2,000 youngest genes by means of FUMA[22] to identify biological pathways and tissues related to this set of genes ($q < 0.05$, FDR). Gene-set analysis for GO biological terms revealed highest enrichment for these young genes in positive regulation of gene expression ($P = 6.6 \times 10^{-28}$), neurogenesis ($P = 7.1 \times 10^{-25}$), neuron differentiation ($P = 5.4 \times 10^{-24}$), central nervous system development ($P = 5.9 \times 10^{-21}$), regulation of cell differentiation ($P = 4.7 \times 10^{-17}$), and central nervous system development ($P = 2.3 \times 10^{-15}$) among other pathways (Table S8). We found that the set of youngest genes was differentially expressed (down-regulated) in the amygdala ($P = 6 \times 10^{-13}$), putamen ($P = 1.3 \times 10^{-12}$), hippocampus ($P = 1.6 \times 10^{-12}$), anterior cingulate cortex ($P = 2.5 \times 10^{-12}$), and basal ganglia ($P = 5.5 \times 10^{-13}$) within others tissues (Table S9).

***Validation gene expression permutation test (1).*** We evaluated whether the top 1% youngest genes are significantly highly or lowly expressed in certain brain areas using different atlases. The EK atlas[26] provided information about brain cytoarchitectonics (15 brain regions, Bonferroni *P* value threshold < $3.3 \times 10^{-3}$), revealing a significantly higher expression in the agranular medial orbitofrontal and inferior frontal cortex ($P = 1.3 \times 10^{-5}$ and $P = 4.3 \times 10^{-4}$, respectively), and a lower expression in the polar occipital cortex ($P = 2.5 \times 10^{-4}$). Using BA atlas[26] (39 brain regions, Bonferroni *P* value threshold < $1.2 \times 10^{-3}$), we found a trend for higher expression in BA45 (nominally significant, $P = 5 \times 10^{-3}$) and lower expression in BA18 ($P$ = 8.2 x $10^{-4}$) of the top 1% youngest genes. Using DK-144 atlas[27] (57 brain regions, Bonferroni $P$ value threshold < 8.7 x $10^{-4}$), we found significantly higher expression in pars triangularis ($P$ = 6.7 x $10^{-4}$), and lower expression in lateral occipital ($P$ = 2.8 x $10^{-4}$) and superior parietal ($P$ = 4.1 x $10^{-4}$).

***Validation gene expression permutation test (2).*** To assess the robustness of the results we conducted the same main analysis varying the number of young genes included. We found that the 0.5% youngest genes ($n$ = 82, ranging from 22,173 to 6,717 years ago) show a trend for overexpression in the pars triangularis (nominally significant, $P$ = 7.1 x $10^{-3}$). The 5% youngest genes ($n$ = 818 genes, ranging from 44,157 to 6,717 years ago) presented a trend towards an overexpression in language-related areas, namely pars opercularis ($P$ = 3.5 x $10^{-3}$), banks of the superior temporal sulcus ($P$ = 7.6 x $10^{-3}$), and pars triangularis ($P$ = 1.3 x $10^{-2}$), however these effects were not longer significant after Bonferroni correction. On the other hand, the 5% youngest genes were significantly underexpressed in the lateral occipital cortex ($P$ = 4 x $10^{-5}$). We further assessed the gene expression of the top 10% youngest genes ($n$ = 1,635 genes, ranging from 54,656 to 6,717 years ago). Again, the youngest genes were significantly overexpressed in the pars triangularis ($P$ = 7.3 x $10^{-4}$) and underexpressed in the lateral occipital cortex ($P$ = 1.1 x $10^{-3}$).

***Validation gene expression permutation test (3).*** We repeated the main analysis (with DK atlas, 34 brain regions, Bonferroni $P$ value threshold < 1.4 x $10^{-3}$) including genes containing more than ten SNPs and with a median quality score above 0.7 (13,852 genes with gene expression and evolutionary age information). The strongest effect of the top 1% youngest genes ($n$ = 139) was overexpressed in the pars triangularis but was not significant after Bonferroni correction ($P$ = 7.7 x $10^{-3}$).

### Supplementary Figures

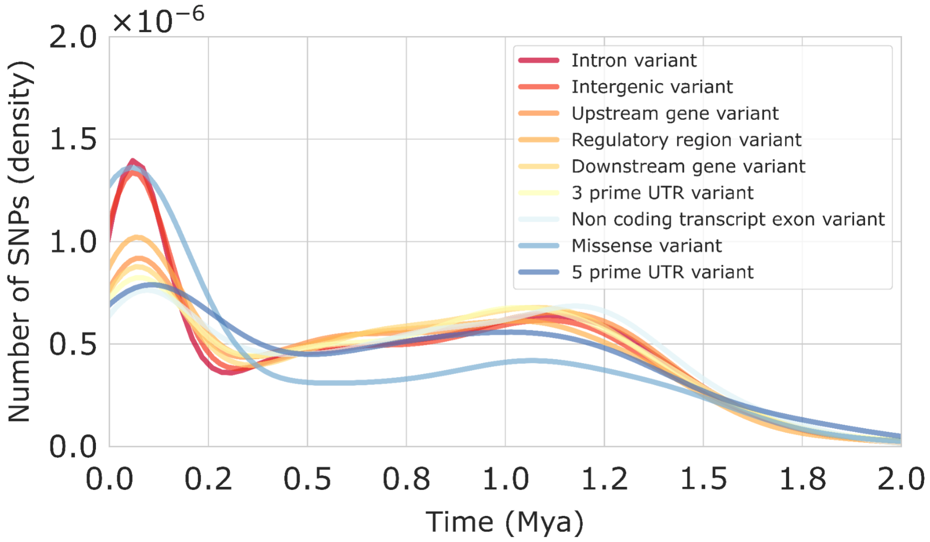


**Supplementary Figure 1. Genetic timeline of human-phenotypic SNPs grouped by their functional consequences.**

Timeline of SNPs related to human phenotypes grouped by the variant functional consequences (unique human-phenotypic SNPs, *n* = 36,506). Histogram of the density (y-axis) of the number of SNPs emerging across time (shown until 2M years ago; x-axis). Mya, million years ago.

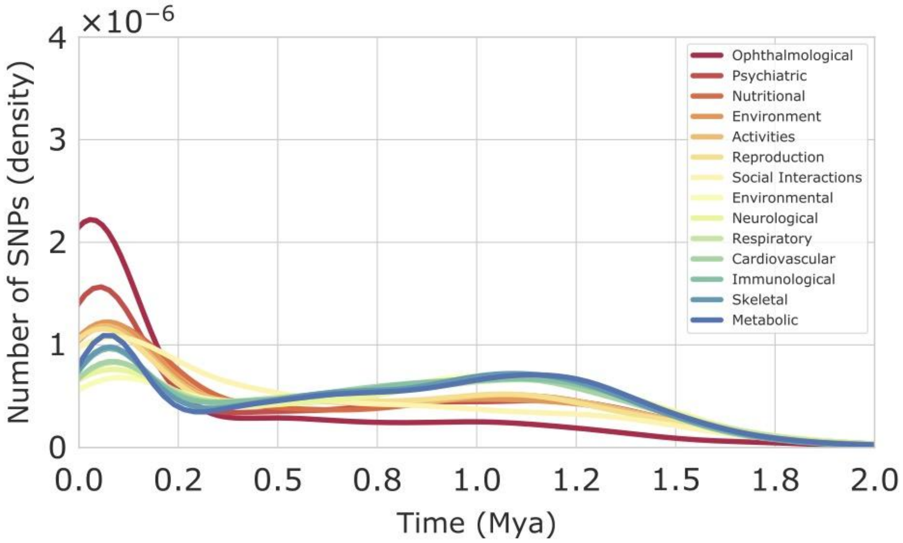


**Supplementary Figure 2. Genetic timeline of human-phenotypic SNPs (domain level).** Timeline of SNPs related to human phenotypes at the domain level. Histogram of the density (y-axis) of the number of SNPs emerging across time (shown until 2M years ago; x-axis). Figure shows the domains with a median evolutionary age younger (following legend order, from 'Ophthalmological' to 'Social Interactions') and older (from 'Environmental' to 'Metabolic') than expected by the null model ($n$ = 28 domains, Bonferroni $P$ value threshold < $1.7 \times 10^{-3}$). Mya, million years ago.

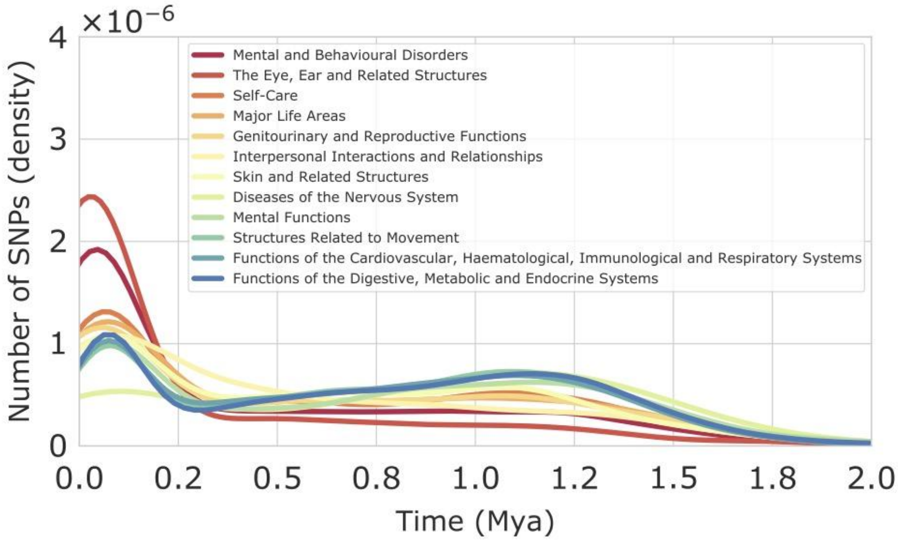


**Supplementary Figure 3. Genetic timeline of human-phenotypic SNPs (chapter level).** Timeline of SNPs related to human phenotypes at the chapter level from the GWAS Atlas. Histogram of the density (y-axis) of the number of SNPs emerging across time (shown until 2M years ago; x-axis). Figure shows the chapters with a median evolutionary age younger (following legend order, from 'Mental and Behavioural Disorders' to 'Skin and Related Structures') and older (from 'Disease of the Nervous System' to 'Functions of the Digestive, Metabolic and Endocrine Systems') than expected by the null model (*n* = 50 chapters, Bonferroni *P* value threshold < $1 \times 10^{-3}$). Mya, million years ago.

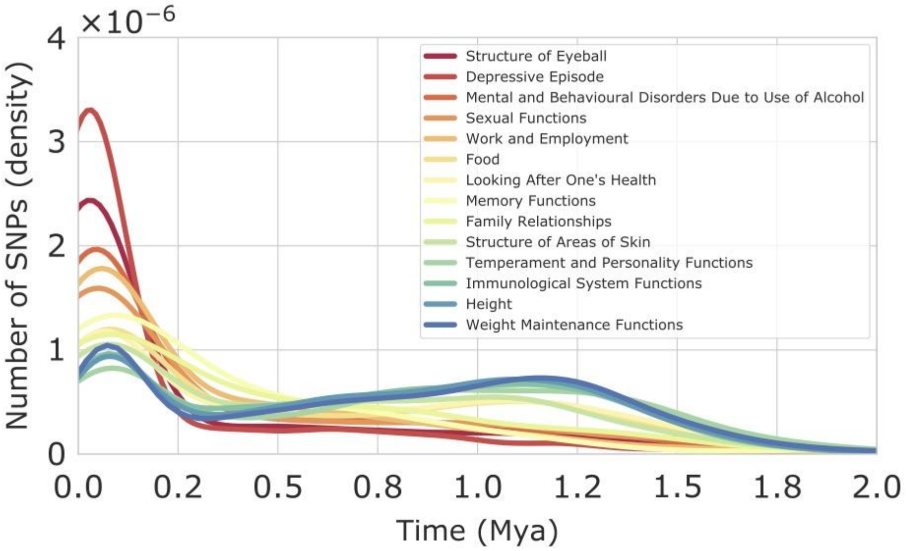


**Supplementary Figure 4. Genetic timeline of human-phenotypic SNPs (subchapter level).**

Timeline of SNPs related to human phenotypes at the subchapter level from the GWAS Atlas. Histogram of the density (y-axis) of the number of SNPs emerging across time (shown until 2M years ago; x-axis). Figure shows the subchapters with a median evolutionary age younger (following legend order, from 'Structure of Eyeball' to 'Structures of Areas of Skin') and older (from 'Temperament and Personality Function' to 'Weight Maintenance Functions') than expected by the null model ($n$ = 172 subchapters, Bonferroni $P$ value threshold < $2.9 \times 10^{-4}$). Mya, million years ago.

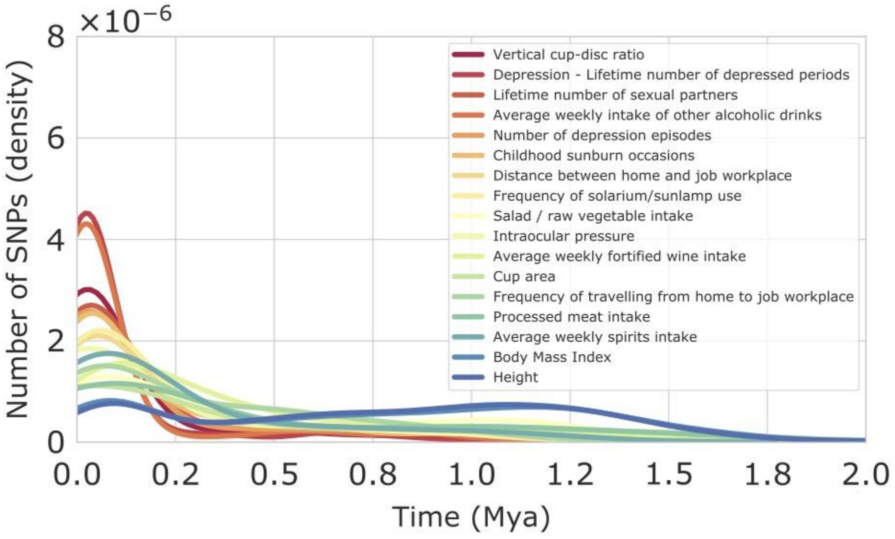


**Supplementary Figure 5. Genetic timeline of human-phenotypic SNPs (trait level).** Timeline of SNPs related to human phenotypes at the trait level from the GWAS Atlas. Histogram of the density (y-axis) of the number of SNPs emerging across time (shown until 2M years ago; x-axis). Figure shows the traits with a median evolutionary age younger (following legend order, from 'Vertical cup-disc ratio' to 'Average weekly spirits intake') and older (from 'Body Mass Index' to 'Height') than expected by the null model (*n* = 2,251 traits, Bonferroni *P* value threshold < $2.2 \times 10^{-5}$). Mya, million years ago.

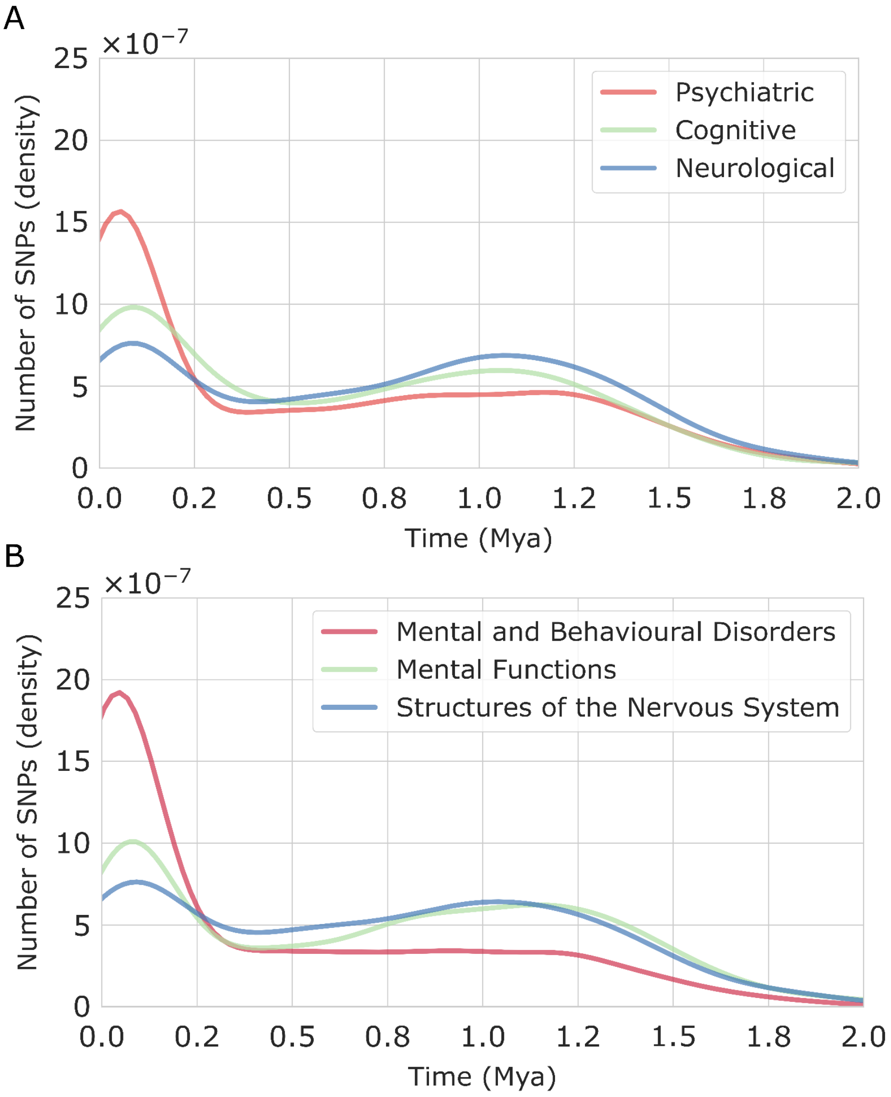


**Supplementary Figure 6. Genetic timeline of brain, cognitive and psychiatric traits.** Timeline of the density (normalised count; y-axis) of the number of SNPs across the last 2M years (x-axis) in relation to brain-related traits (blue), cognition (green) and psychiatric disorders (red). The phenotypes are organised at the (*a*) domain and (*b*) more detailed chapter levels. Genetic variants related to the neurological domain (median evolutionary age = 800,795 years old) emerged earlier in evolution than variants related to cognition ($t$ = 3.95, $P$ = 7.9 x $10^{-5}$; median age = 611,820) and psychiatric disorders ($t$ = 8.91, $P$ = 6.9 x $10^{-19}$; median age = 412,639). Mya, million years ago.

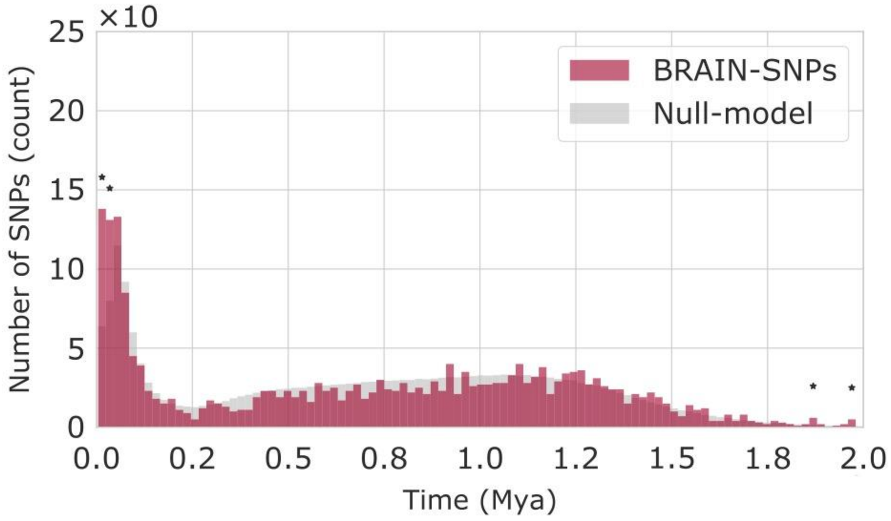


**Supplementary Figure 7. Genetic timeline of BRAIN-SNPs.**

Absolute count (y-axis) of number of SNPs per time-bin (100 bins of ~20,000 years old; shown until 2M years ago; x-axis) of all significant BRAIN-SNPs (red) associated with brain phenotypes extracted from the UK Biobank BIG40 (1,744 GWAS studies, 2,476 unique SNPs). BRAIN-SNPs date estimates ranged from 3,621,625 - 4,457 years ago. Asterisks (*) denote bins where the number of BRAIN-SNPs per time-bin significantly exceeds the null-model of random equally sized sets of SNPs selected from all SNPs from the HDG (100 tests, Bonferroni $P$ value threshold $< 5 \times 10^{-4}$). Mya, million years ago.

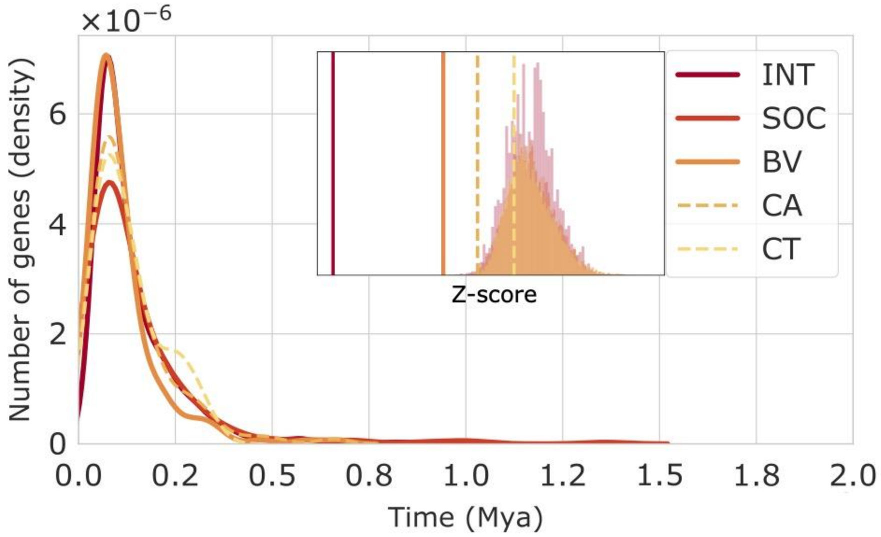


**Supplementary Figure 8. Genetic timeline of genes linked to brain and cognitive phenotypes.**

Timeline of genes associated with five brain and cognitive phenotypes, including intelligence, sociability, brain volume, cortical area, and cortical thickness from previous GWAS. Histogram of the density (y-axis) of the number of genes emerging across time (shown until 2M years ago; x-axis). Solid lines denote phenotypes significantly younger than the null model (Bonferroni *P* value threshold < $5 \times 10^{-3}$; see insert for the null model distribution of each condition). Mya, million years ago; INT, intelligence; SOC, sociability; BV, brain volume; CA, cortical area; CT, cortical thickness.

### Supplementary Tables

**Supplementary Table 1. Human-phenotypic SNPs permutation test.** Complete results of the permutation test assessing whether the median evolutionary age (third column) of SNPs related to human phenotypes (first column) at the domain/chapter/subchapter/trait level is older (sixth column; positive z-score) or younger (negative z-score) than expected by chance (seventh column) given the number of SNPs (second column). The 75th and 25th percentiles of evolutionary age for the SNPs related to each phenotype are also reported (fourth and fifth columns, respectively). *Table is presented as an online file.*

**Supplementary Table 2. Human-phenotypic SNPs permutation test (MAF-controlled).** Complete results of the permutation test assessing whether the median evolutionary age (third column) of SNPs related to human phenotypes (first column) at the domain/chapter/subchapter/trait level is older (seventh column; positive z-score) or younger (negative z-score) than expected by chance (eight column) given the number of SNPs (second column) and their corresponding MAF (sixth column). The 75th and 25th percentiles of evolutionary age for the SNPs related to each phenotype are also reported (fourth and fifth columns, respectively). *Table is presented as an online file.*

**Supplementary Table 3. Included phenotypes in BRAIN-SNPs.** List of phenotypes (second column) and number of SNPs (third column) organised by brain structures (first column) extracted from the UK Biobank BIG40 dataset. *Table is presented as an online file.*

**Supplementary Table 4. BRAIN-SNPs permutation test.** Results of the permutation test assessing whether the median evolutionary age (third column) of BRAIN-SNPs grouped based on brain structures (first column) are older (fourth column; positive z-score) or younger (negative z-score) than chance (fifth column). Bonferroni *P* value threshold < $5.5 \times 10^{-3}$ (sixth column).

| Brain phenotype | N SNPs | Median age | Z-score | P value | Bonferroni |
|---|---|---|---|---|---|
| Neocortex | 130 | 400170 | -3.51 | 4.4E-04 | 1 |
| Amygdala | 39 | 332998 | -2.35 | 1.9E-02 | 0 |
| Cerebellum | 144 | 659803 | -0.67 | 5.0E-01 | 0 |
| Hippocampus | 30 | 653218 | -0.14 | 8.9E-01 | 0 |
| Pallidum | 21 | 781750 | 0.18 | 8.6E-01 | 0 |
| White matter | 676 | 729170 | 0.87 | 3.8E-01 | 0 |
| Thalamus | 19 | 965115 | 1.30 | 1.9E-01 | 0 |
| Nucleus accumbens | 11 | 815658 | 1.33 | 1.8E-01 | 0 |
| Caudate nucleus | 25 | 1098163 | 1.96 | 5.0E-02 | 0 |

**Supplementary Table 5. BRAIN-SNPs timeline co-fluctuation.** Spearman's correlation (third column) and *P* value (fourth column) between the genetic timeline of BRAIN-SNPs associated with different brain structures (first and second columns). Bonferroni *P* value threshold < $1.4 \times 10^{-3}$ (fifth column).

| Brain structure 1 | Brain structure 2 | rho | P value | Bonferroni |
|---|---|---|---|---|
| Cerebellum | White matter | 0.81 | 5.3E-24 | 1 |
| Neocortex | White matter | 0.72 | 3.9E-17 | 1 |
| Neocortex | Cerebellum | 0.70 | 7.1E-16 | 1 |
| Cerebellum | Hippocampus | 0.63 | 1.4E-12 | 1 |
| Cerebellum | Amygdala | 0.55 | 2.2E-09 | 1 |
| Neocortex | Hippocampus | 0.52 | 3.7E-08 | 1 |
| Hippocampus | White matter | 0.51 | 4.6E-08 | 1 |
| Neocortex | Amygdala | 0.51 | 4.9E-08 | 1 |
| Cerebellum | Pallidum | 0.48 | 3.6E-07 | 1 |
| Cerebellum | Caudate nucleus | 0.48 | 4.0E-07 | 1 |
| Hippocampus | Caudate nucleus | 0.48 | 5.8E-07 | 1 |
| Neocortex | Pallidum | 0.47 | 6.3E-07 | 1 |
| Amygdala | Hippocampus | 0.45 | 2.2E-06 | 1 |
| White matter | Caudate nucleus | 0.45 | 3.4E-06 | 1 |
| Cerebellum | Nucleus accumbens | 0.43 | 6.8E-06 | 1 |
| Neocortex | Caudate nucleus | 0.43 | 7.0E-06 | 1 |
| Amygdala | Caudate nucleus | 0.42 | 1.6E-05 | 1 |
| White matter | Pallidum | 0.41 | 2.8E-05 | 1 |
| Amygdala | White matter | 0.40 | 3.0E-05 | 1 |
| Thalamus | White matter | 0.38 | 1.1E-04 | 1 |
| Amygdala | Nucleus accumbens | 0.37 | 1.4E-04 | 1 |
| Cerebellum | Thalamus | 0.37 | 1.4E-04 | 1 |
| Nucleus accumbens | White matter | 0.36 | 1.9E-04 | 1 |
| Neocortex | Nucleus accumbens | 0.36 | 2.1E-04 | 1 |
| Nucleus accumbens | Pallidum | 0.34 | 4.6E-04 | 1 |
| Amygdala | Thalamus | 0.30 | 2.8E-03 | 0 |
| Neocortex | Thalamus | 0.29 | 4.0E-03 | 0 |
| Hippocampus | Thalamus | 0.28 | 5.1E-03 | 0 |
| Amygdala | Pallidum | 0.26 | 9.0E-03 | 0 |

| | | | | |
|---|---|---|---|---|
| Pallidum | Caudate nucleus | 0.25 | 1.2E-02 | 0 |
| Hippocampus | Pallidum | 0.22 | 2.9E-02 | 0 |
| Thalamus | Pallidum | 0.20 | 4.2E-02 | 0 |
| Hippocampus | Nucleus accumbens | 0.20 | 4.4E-02 | 0 |
| Nucleus accumbens | Caudate nucleus | 0.17 | 8.5E-02 | 0 |
| Thalamus | Caudate nucleus | 0.17 | 8.9E-02 | 0 |
| Thalamus | Nucleus accumbens | 0.01 | 9.3E-01 | 0 |

**Supplementary Table 6. GWAS included in the present study.** Details of the GWAS included in the gene and gene-set analysis conducted with MAGMA.

| Phenotype | N case | N control | N total | Population | Sumstats download link |
|---|---|---|---|---|---|
| Schizophrenia | 33426 | 54065 | 87491 | EU | https://figshare.com/articles/dataset/cdg2018-bip-scz/14672019 |
| Bipolar disorder | 20129 | 54065 | 74194 | EU | https://figshare.com/articles/dataset/cdg2018-bip-scz/14672019 |
| ASD | 18381 | 27969 | 46350 | EU | https://figshare.com/articles/dataset/asd2019/14671989 |
| MDD | 59851 | 113154 | 173005 | EU | https://figshare.com/articles/dataset/mdd2018/14672085 |
| Alzheimer's disease | 71880 | 383378 | 455258 | EU | https://ctg.cncr.nl/software/summary_statistics |
| Brain volume | | | 47316 | EU | https://ctg.cncr.nl/software/summary_statistics |
| Cortical area | | | 51665 | EU | https://enigma.ini.usc.edu/research/download-enigma-gwas-results/ |
| Cortical thickness | | | 51665 | EU | https://enigma.ini.usc.edu/research/download-enigma-gwas-results/ |
| Intelligence | | | 269867 | EU | https://ctg.cncr.nl/software/summary_statistics |
| Sociability | | | 452302 | EU | https://www.repository.cam.ac.uk/handle/1810/277812 |

**Supplementary Table 7. Gene-set analysis.** Results of the MAGMA gene-set analysis (third and fourth columns) of the top 2,000 oldest and youngest genes (first column) testing for enrichment of genes related to brain, cognitive and neuropsychiatric brain phenotypes (second column). Bonferroni *P* value threshold < $2.5 \times 10^{-3}$ (fifth column).

| **Gene-set** | **Phenotype** | **Beta** | **P value** | **Bonferroni** |
|---|---|---|---|---|
| Old genes | Alzheimer's disease | -0.04 | 9.5E-01 | 0 |
| Young genes | Alzheimer's disease | 0.06 | 4.3E-03 | 0 |
| Old genes | Autism spectrum disorder | -0.04 | 9.6E-01 | 0 |
| Young genes | Autism spectrum disorder | 0.04 | 5.1E-02 | 0 |
| Old genes | Bipolar disorder | -0.03 | 8.9E-01 | 0 |
| Young genes | Bipolar disorder | 0.03 | 9.1E-02 | 0 |
| Old genes | Brain volume | -0.03 | 8.0E-01 | 0 |
| Young genes | Brain volume | 0.05 | 3.1E-02 | 0 |
| Old genes | Cortical area | -0.04 | 9.5E-01 | 0 |
| Young genes | Cortical area | 0.08 | 6.7E-04 | 1 |
| Old genes | Cortical thickness | -0.06 | 9.9E-01 | 0 |
| Young genes | Cortical thickness | 0.02 | 1.4E-01 | 0 |
| Old genes | Intelligence | -0.11 | 1.0E+00 | 0 |
| Young genes | Intelligence | 0.13 | 2.3E-06 | 1 |
| Old genes | Major depressive disorder | 0.02 | 1.7E-01 | 0 |
| Young genes | Major depressive disorder | 0.03 | 9.3E-02 | 0 |
| Old genes | Schizophrenia | -0.07 | 9.9E-01 | 0 |
| Young genes | Schizophrenia | 0.07 | 8.0E-03 | 0 |
| Old genes | Sociability | -0.04 | 9.5E-01 | 0 |
| Young genes | Sociability | -0.02 | 8.4E-01 | 0 |

**Supplementary Table 8. GO biological functional annotations.** Biological functional annotation performed by FUMA of the 2,000 genes with the youngest median evolutionary age. The top 10 genes-sets (first column) are shown with the corresponding number of genes in each set (second column), the number of overlapping genes (third column), and the enrichment probability FDR-adjusted (fourth column).

| **Gene-set** | **N genes** | **N overlap** | **P-adjusted (FDR)** |
|---|---|---|---|
| GO POSITIVE REGULATION OF BIOSYNTHETIC PROCESS | 1955 | 213 | 2.4E-24 |
| GO POSITIVE REGULATION OF GENE EXPRESSION | 1945 | 212 | 2.4E-24 |
| GO POSITIVE REGULATION OF RNA BIOSYNTHETIC PROCESS | 1585 | 180 | 2.4E-22 |
| GO NEUROGENESIS | 1586 | 178 | 1.3E-21 |
| GO NEURON DIFFERENTIATION | 1337 | 157 | 8.0E-21 |
| GO POSITIVE_REGULATION OF TRANSCRIPTION BY RNA POLYMERASE II | 1172 | 138 | 3.4E-18 |
| GO CENTRAL NERVOUS SYSTEM DEVELOPMENT | 966 | 121 | 6.3E-18 |
| GO REGULATION OF CELL DIFFERENTIATION | 1829 | 184 | 2.2E-17 |
| GO LOCOMOTION | 1906 | 188 | 6.1E-17 |
| GO APOPTOTIC PROCESS | 1945 | 188 | 4.7E-16 |

**Supplementary Table 9. Differential gene expression analysis.** Differential gene expression analysis across human tissue performed by FUMA of the 2,000 genes with the youngest median evolutionary age. The top ten differentially expressed (first column) genes-sets (second column) are shown with the corresponding number of genes in each set (third column), the number of overlapping genes (fourth column), and the enrichment probability FDR-adjusted (fifth column).

| Category | Gene-set | N genes | N overlap | P-adjusted (FDR) |
|---|---|---|---|---|
| DEG.down | Pancreas | 9586 | 705 | 2.0E-14 |
| DEG.down | Heart Left Ventricle | 9443 | 689 | 3.9E-13 |
| DEG.down | Brain Amygdala | 7678 | 569 | 3.2E-11 |
| DEG.down | Brain Putamen basal ganglia | 7776 | 573 | 7.3E-11 |
| DEG.down | Brain Hippocampus | 7493 | 555 | 8.5E-11 |
| DEG.down | Brain Anterior cingulate cortex BA24 | 6651 | 501 | 1.3E-10 |
| DEG.down | Whole Blood | 6908 | 516 | 2.1E-10 |
| DEG.down | Brain Caudate basal ganglia | 6858 | 512 | 3.0E-10 |
| DEG.down | Brain Substantia nigra | 7162 | 530 | 4.4E-10 |
| DEG.down | Liver | 7985 | 579 | 9.7E-10 |

**Supplementary Table 10. Normative gene expression analysis.** Normative gene expression levels (extracted from AHBA) of the genes with the youngest evolutionary age across brain areas using DK atlas (first column). A null-random-gene model was applied to assess whether the youngest genes are more expressed (second column; positive z-score) or less expressed (negative z-score) in each brain area than random genes (third column). Bonferroni *P* value threshold < 1.5 x $10^{-3}$ (fifth column).

| Brain region | Z-score | P value | Bonferroni |
|---|---|---|---|
| Left pars triangularis | 3.42 | 6.2E-04 | 1 |
| Left pars opercularis | 2.10 | 3.6E-02 | 0 |
| Left caudal anterior cingulate | 1.94 | 5.2E-02 | 0 |
| Left entorhinal | 1.76 | 7.8E-02 | 0 |
| Left temporal pole | 1.66 | 9.6E-02 | 0 |
| Left parahippocampal | 1.58 | 1.1E-01 | 0 |
| Left middle temporal | 1.52 | 1.3E-01 | 0 |
| Left isthmus cingulate | 1.32 | 1.9E-01 | 0 |
| Left rostral anterior cingulate | 1.28 | 2.0E-01 | 0 |
| Left superior temporal | 1.22 | 2.2E-01 | 0 |
| Left inferior temporal | 1.06 | 2.9E-01 | 0 |
| Left superior frontal | 1.04 | 3.0E-01 | 0 |
| Left caudal middle frontal | 0.89 | 3.7E-01 | 0 |
| Left paracentral | 0.53 | 6.0E-01 | 0 |
| Left lingual | 0.39 | 7.0E-01 | 0 |
| Left precuneus | 0.37 | 7.1E-01 | 0 |
| Left rostral middle frontal | 0.33 | 7.4E-01 | 0 |
| Left fusiform | 0.30 | 7.6E-01 | 0 |
| Left medial orbitofrontal | 0.19 | 8.5E-01 | 0 |
| Left precentral | 0.18 | 8.6E-01 | 0 |
| Left pars orbitalis | 0.13 | 9.0E-01 | 0 |
| Left transverse temporal | 0.10 | 9.2E-01 | 0 |
| Left frontal pole | 0.03 | 9.7E-01 | 0 |
| Left postcentral | -0.15 | 8.8E-01 | 0 |
| Left lateral orbitofrontal | -0.18 | 8.6E-01 | 0 |
| Left supramarginal | -0.20 | 8.4E-01 | 0 |

| | | | |
|---|---|---|---|
| Left posterior cingulate | -0.39 | 6.9E-01 | 0 |
| Left insula | -1.19 | 2.4E-01 | 0 |
| Left superior parietal | -1.67 | 9.4E-02 | 0 |
| Left cuneus | -1.90 | 5.7E-02 | 0 |
| Left bankssts | -1.94 | 5.2E-02 | 0 |
| Left pericalcarine | -1.97 | 4.8E-02 | 0 |
| Left inferior parietal | -2.22 | 2.7E-02 | 0 |
| Left lateral occipital | -3.38 | 7.3E-04 | 1 |
