## Supplementary Table 1 for "Mapping the genetic evolutionary timeline of human neural and cognitive traits"

| Domain | N SNPs | Median age | Age 75% upper bound | Age 25% lower bound | Z-score | P value | Bonferroni |
|---|---|---|---|---|---|---|---|
| Ophthalmological | 823 | 30484 | 501546 | 19323 | -18.37 | 2.1E-75 | 1 |
| Psychiatric | 4022 | 412639 | 1034993 | 49571 | -17.00 | 7.8E-65 | 1 |
| Nutritional | 929 | 429993 | 1050270 | 67491 | -7.55 | 4.2E-14 | 1 |
| Environment | 1161 | 473478 | 1024142 | 70585 | -6.89 | 5.7E-12 | 1 |
| Activities | 1784 | 514085 | 1051021 | 68298 | -6.84 | 7.8E-12 | 1 |
| Reproduction | 1226 | 525596 | 1052098 | 60377 | -5.21 | 1.8E-07 | 1 |
| Social Interactions | 252 | 393951 | 858820 | 62358 | -4.49 | 7.1E-06 | 1 |
| Neoplasms | 378 | 593145 | 990794 | 126391 | -1.61 | 1.1E-01 | 0 |
| Cognitive | 600 | 611820 | 1083358 | 84270 | -1.54 | 1.2E-01 | 0 |
| Dermatological | 1833 | 643843 | 1075185 | 114835 | -1.30 | 1.9E-01 | 0 |
| Gastrointestinal | 694 | 635919 | 1060403 | 164244 | -1.01 | 3.1E-01 | 0 |
| Aging | 2 | 373150 | 415107 | 331193 | -0.82 | 4.1E-01 | 0 |
| Mortality | 147 | 617010 | 1045174 | 103406 | -0.69 | 4.9E-01 | 0 |
| Hematological | 57 | 595340 | 1052398 | 214394 | -0.57 | 5.7E-01 | 0 |
| Ear, Nose, Throat | 52 | 642079 | 1078489 | 96134 | -0.22 | 8.2E-01 | 0 |
| Body Structures | 91 | 697705 | 1094841 | 139461 | 0.26 | 7.9E-01 | 0 |
| Infection | 6 | 752978 | 1142349 | 462069 | 0.32 | 7.5E-01 | 0 |
| Muscular | 3 | 782185 | 1153615 | 691033 | 0.33 | 7.4E-01 | 0 |
| Cell | 289 | 736593 | 1116948 | 213623 | 1.08 | 2.8E-01 | 0 |
| Connective Tissue | 95 | 824043 | 1177130 | 320071 | 1.51 | 1.3E-01 | 0 |
| Endocrine | 873 | 757165 | 1092063 | 359748 | 2.49 | 1.3E-02 | 0 |
| Environmental | 599 | 803743 | 1150583 | 266960 | 3.21 | 1.3E-03 | 1 |
| Neurological | 836 | 800795 | 1174863 | 170011 | 3.69 | 2.2E-04 | 1 |
| Respiratory | 1767 | 760510 | 1154128 | 208657 | 3.75 | 1.7E-04 | 1 |

| | | | | | | | |
|---|---|---|---|---|---|---|---|
| Cardiovascular | 2010 | 756169 | 1134757 | 225276 | 3.79 | 1.5E-04 | 1 |
| Immunological | 4197 | 747780 | 1134575 | 207559 | 5.05 | 4.3E-07 | 1 |
| Skeletal | 4452 | 759056 | 1142788 | 205903 | 6.05 | 1.5E-09 | 1 |
| Metabolic | 10287 | 780323 | 1170134 | 196880 | 12.52 | 5.9E-36 | 1 |

| Chapter | N SNPs | Median age | Age 75% upper bound | Age 25% lower bound | Z-score | P value | Bonferroni |
|---|---|---|---|---|---|---|---|
| Mental and Behavioural Disorders | 2331 | 111560 | 803943 | 25013 | -26.63 | 3.3E-156 | 1 |
| The Eye, Ear and Related Structures | 756 | 26744 | 381293 | 18411 | -17.72 | 2.8E-70 | 1 |
| Self-Care | 2612 | 487959 | 1049054 | 67786 | -9.67 | 4.2E-22 | 1 |
| Major Life Areas | 1099 | 473045 | 1018917 | 70947 | -6.73 | 1.7E-11 | 1 |
| Genitourinary and Reproductive Functions | 1108 | 501423 | 1048990 | 57223 | -5.86 | 4.6E-09 | 1 |
| Interpersonal Interactions and Relationships | 252 | 393951 | 858820 | 62358 | -4.50 | 6.6E-06 | 1 |
| Skin and Related Structures | 1066 | 548430 | 1046899 | 79302 | -4.08 | 4.5E-05 | 1 |
| Injuries to the Wrist or Hand | 12 | 89668 | 637349 | 43850 | -2.22 | 2.6E-02 | 0 |
| Domestic Life | 9 | 87761 | 1197513 | 64125 | -1.92 | 5.5E-02 | 0 |
| Pregnancy, Childbirth and the Puerperium | 11 | 109982 | 790909 | 37530 | -1.92 | 5.5E-02 | 0 |
| Benign Neoplasms | 13 | 196128 | 895060 | 112635 | -1.77 | 7.7E-02 | 0 |
| Mortality | 126 | 534155 | 1013774 | 102262 | -1.61 | 1.1E-01 | 0 |
| Malignant Neoplasms | 366 | 593145 | 990794 | 137758 | -1.57 | 1.2E-01 | 0 |
| Certain Conditions Originating in the Perinatal Period | 1 | 40478 | 40478 | 40478 | -1.22 | 2.2E-01 | 0 |
| Voice and Speech Functions | 1 | 69269 | 69269 | 69269 | -1.15 | 2.5E-01 | 0 |
| Diseases of the Digestive System | 693 | 637523 | 1060418 | 166442 | -0.97 | 3.3E-01 | 0 |
| External Causes of Morbidity and Mortality | 1 | 230508 | 230508 | 230508 | -0.86 | 3.9E-01 | 0 |
| Structures of the Cardiovascular, Immunological and Respiratory Systems | 14 | 530470 | 774106 | 230354 | -0.53 | 6.0E-01 | 0 |
| Persons Encountering Health Services in Circumstances related to Reproduction | 2 | 495450 | 726366 | 264534 | -0.49 | 6.2E-01 | 0 |
| Mobility | 15 | 572478 | 791724 | 344053 | -0.37 | 7.1E-01 | 0 |
| Communication | 62 | 633110 | 1099613 | 95419 | -0.31 | 7.6E-01 | 0 |
| Diseases of the Genitourinary System | 92 | 642649 | 1082311 | 169460 | -0.30 | 7.6E-01 | 0 |
| Neuromusculoskeletal and Movement-Related Functions | 5 | 599880 | 782185 | 370965 | -0.15 | 8.8E-01 | 0 |
| Symptoms, Signs and Abnormal Clinical and Laboratory Findings, Not Elsewhere C | 29 | 651273 | 1186478 | 258348 | -0.08 | 9.4E-01 | 0 |
| Factors Influencing Health Status and Contact with Health Services | 12 | 656330 | 1023916 | 166671 | 0.01 | 9.9E-01 | 0 |
| Diseases of the Eye and Adnexa | 72 | 680315 | 1062517 | 140268 | 0.08 | 9.4E-01 | 0 |
| Structures Related to Genitourinary and Reproductive Functions | 2 | 721569 | 763804 | 679333 | 0.10 | 9.2E-01 | 0 |
| Certain Infectious and Parasitic Diseases | 2 | 730364 | 870164 | 590563 | 0.13 | 8.9E-01 | 0 |
| Structure Involved in Voice and Speech | 61 | 697030 | 1082750 | 113557 | 0.19 | 8.5E-01 | 0 |
| Disease of the Respiratory System | 189 | 701663 | 1147733 | 142038 | 0.38 | 7.0E-01 | 0 |
| Diseases of the Skin and Subcutaneous Tissue | 61 | 750523 | 1250975 | 157010 | 0.64 | 5.2E-01 | 0 |
| Injury, Poisoning and Certain Other Consequences of External Causes | 32 | 804538 | 1073765 | 208062 | 0.80 | 4.2E-01 | 0 |
| Certain conditions originating in the perinatal period | 7 | 924228 | 1220385 | 660623 | 0.82 | 4.1E-01 | 0 |
| Congenital Malformations, Deformations and Chromosomal Abnormalities | 5 | 1031778 | 1061213 | 78554 | 1.05 | 2.9E-01 | 0 |
| Persons with Potential Health Hazards Related to Family and Personal History and | 21 | 890533 | 1077720 | 223191 | 1.08 | 2.8E-01 | 0 |
| Function of DNA | 289 | 736593 | 1116948 | 213623 | 1.10 | 2.7E-01 | 0 |
| Diseases of the Respiratory System | 243 | 749068 | 1103885 | 241583 | 1.19 | 2.3E-01 | 0 |
| Diseases of the Musculoskeletal System and Connective Tissue | 234 | 755908 | 1178558 | 313889 | 1.28 | 2.0E-01 | 0 |
| Structures of the Nervous System | 574 | 728523 | 1149092 | 160080 | 1.31 | 1.9E-01 | 0 |

| | | | | | | | |
|---|---|---|---|---|---|---|---|
| Persons Encountering Health Status and Contact with Health Services | 1 | 1529998 | 1529998 | 1529998 | 1.56 | 1.2E-01 | 0 |
| Sensory Functions and Pain | 116 | 848390 | 1136323 | 168780 | 1.92 | 5.5E-02 | 0 |
| Diseases of the Circulatory System | 829 | 755865 | 1131780 | 301905 | 2.39 | 1.7E-02 | 0 |
| Endocrine, Nutritional and Metabolic Diseases | 912 | 753879 | 1112101 | 314569 | 2.45 | 1.4E-02 | 0 |
| Functions of the Skin and Related Structures | 718 | 774485 | 1109228 | 299151 | 2.73 | 6.3E-03 | 0 |
| Products and Technology | 651 | 789108 | 1147636 | 201948 | 3.02 | 2.6E-03 | 0 |
| Diseases of the Nervous System | 207 | 932355 | 1242859 | 276185 | 3.79 | 1.5E-04 | 1 |
| Mental Functions | 2315 | 755888 | 1169583 | 114127 | 4.02 | 5.8E-05 | 1 |
| Structures Related to Movement | 4302 | 761628 | 1142566 | 203376 | 6.15 | 7.7E-10 | 1 |
| Functions of the Cardiovascular, Haematological, Immunological and Respiratory S | 6844 | 753774 | 1141570 | 207351 | 7.31 | 2.8E-13 | 1 |
| Functions of the Digestive, Metabolic and Endocrine Systems | 10300 | 780269 | 1168384 | 200705 | 12.61 | 1.9E-36 | 1 |

| Subchapter | N SNPs | Median age | Age 75% upper bound | Age 25% lower bound | Z-score | P value | Bonferroni |
|---|---|---|---|---|---|---|---|
| Structure of Eyeball | 756 | 26744 | 381293 | 18411 | -17.74 | 2.1E-70 | 1 |
| Depressive Episode | 745 | 27526 | 123864 | 15392 | -17.71 | 3.6E-70 | 1 |
| Mental and Behavioural Disorders Due to Use of Alcohol | 715 | 72959 | 581134 | 21394 | -16.48 | 5.3E-61 | 1 |
| Sexual Functions | 500 | 79911 | 696678 | 21077 | -13.51 | 1.4E-41 | 1 |
| Work and Employment | 346 | 99706 | 531414 | 25710 | -10.75 | 5.8E-27 | 1 |
| Food | 929 | 429993 | 1050270 | 67491 | -7.48 | 7.6E-14 | 1 |
| Looking After One's Health | 1702 | 510761 | 1049033 | 67837 | -6.86 | 6.7E-12 | 1 |
| Memory Functions | 85 | 155176 | 550950 | 47118 | -4.85 | 1.2E-06 | 1 |
| Family Relationships | 125 | 293003 | 765840 | 47668 | -4.28 | 1.9E-05 | 1 |
| Structure of Areas of Skin | 567 | 516270 | 1018909 | 69258 | -3.85 | 1.2E-04 | 1 |
| Parent-Child Relationships | 41 | 124491 | 467298 | 61860 | -3.53 | 4.2E-04 | 0 |
| Assets | 52 | 242926 | 1078069 | 57849 | -3.16 | 1.6E-03 | 0 |
| Malignant Neoplasms of Breast | 216 | 493833 | 975567 | 101913 | -2.68 | 7.3E-03 | 0 |
| Carpal Tunnel Syndrome | 12 | 89668 | 637349 | 43850 | -2.20 | 2.7E-02 | 0 |
| Mental Functions, Unspecified | 4 | 48181 | 350559 | 36350 | -1.82 | 6.9E-02 | 0 |
| Benign Neoplasm of Colon, Rectum, Anus and Anal Canal | 13 | 196128 | 895060 | 112635 | -1.80 | 7.1E-02 | 0 |
| Preterm Labour and Delivery | 2 | 21325 | 24569 | 18081 | -1.76 | 7.9E-02 | 0 |
| Acquisition of Necessities | 4 | 71768 | 392898 | 51683 | -1.71 | 8.7E-02 | 0 |
| Structure of Hair | 516 | 600458 | 1056409 | 93271 | -1.68 | 9.2E-02 | 0 |
| All-Cause Mortality | 126 | 534155 | 1013774 | 102262 | -1.59 | 1.1E-01 | 0 |
| Education | 600 | 609431 | 1126788 | 95024 | -1.57 | 1.2E-01 | 0 |
| Mental and Behavioural Disorders Due to Use of Cannabinoid | 9 | 204782 | 1218098 | 50761 | -1.52 | 1.3E-01 | 0 |
| Unspecified | 5 | 123446 | 1197513 | 64740 | -1.48 | 1.4E-01 | 0 |
| Noninfective entertitis and Colitis | 511 | 615780 | 1045638 | 198361 | -1.33 | 1.8E-01 | 0 |
| Presence of Other Functional Implants | 1 | 23028 | 23028 | 23028 | -1.25 | 2.1E-01 | 0 |
| Dislocation, Sprain and Strain of Joints and Ligaments of Knee | 1 | 26829 | 26829 | 26829 | -1.24 | 2.1E-01 | 0 |
| Disorders of eyelid, lacrimal system and orbit | 2 | 219257 | 228872 | 209641 | -1.24 | 2.1E-01 | 0 |
| Injury of Muscle and Tendon at Shoulder and Upper Arm Level | 1 | 34759 | 34759 | 34759 | -1.24 | 2.2E-01 | 0 |
| Genitourinary and Reproductive Functions, Unspecified | 1 | 26754 | 26754 | 26754 | -1.23 | 2.2E-01 | 0 |
| Problems Related to Upbringing | 14 | 364589 | 708594 | 50867 | -1.22 | 2.2E-01 | 0 |
| Digestive System Disorders of Fetus and Newborn | 1 | 40478 | 40478 | 40478 | -1.21 | 2.3E-01 | 0 |
| Structure of Nose | 1 | 69269 | 69269 | 69269 | -1.17 | 2.4E-01 | 0 |
| Gingivitis and Periodontal Diseases | 1 | 62660 | 62660 | 62660 | -1.17 | 2.4E-01 | 0 |
| Bipolar Affective Disorder/Depressive Episode | 1 | 66277 | 66277 | 66277 | -1.16 | 2.5E-01 | 0 |
| Abdominal and Pelvic Pain | 1 | 68013 | 68013 | 68013 | -1.16 | 2.5E-01 | 0 |
| Cardiomiopathy | 1 | 76263 | 76263 | 76263 | -1.15 | 2.5E-01 | 0 |
| Coxarthrosis [Anthrosis of Hip] | 10 | 343796 | 693370 | 106550 | -1.12 | 2.6E-01 | 0 |
| Mental Functions of Language | 5 | 252948 | 270908 | 247992 | -1.12 | 2.6E-01 | 0 |
| Haematological System Functions | 26 | 452849 | 863486 | 86642 | -1.11 | 2.7E-01 | 0 |
| Disorders of Lens | 2 | 277362 | 357695 | 197029 | -1.07 | 2.8E-01 | 0 |

| | | | | | | | |
|---|---|---|---|---|---|---|---|
| Psoriasis | 5 | 269828 | 367203 | 157010 | -1.05 | 2.9E-01 | 0 |
| Gout | 2 | 288515 | 415423 | 161608 | -1.05 | 3.0E-01 | 0 |
| Personal History of Certain Other Diseases | 3 | 225979 | 1377815 | 131869 | -1.04 | 3.0E-01 | 0 |
| Mental and Behavioural Disorders Due to Use of Tobacco | 540 | 632559 | 1152929 | 96316 | -0.98 | 3.3E-01 | 0 |
| Functions of Joints and Bones | 2 | 314849 | 342907 | 286791 | -0.97 | 3.3E-01 | 0 |
| Acquired Absence of Organs, Not Elsewhere Classified | 6 | 351934 | 726293 | 99518 | -0.97 | 3.3E-01 | 0 |
| Disorders of Gallbladder, Biliary Tract and Pancreas | 1 | 160339 | 160339 | 160339 | -0.97 | 3.3E-01 | 0 |
| General Metabolic Functions | 1084 | 648105 | 1075349 | 149713 | -0.86 | 3.9E-01 | 0 |
| Intestinal Malabsorption | 38 | 533051 | 1057677 | 126359 | -0.86 | 3.9E-01 | 0 |
| Fibrosis and Cirrhosis of Liver | 17 | 471018 | 837185 | 72818 | -0.82 | 4.1E-01 | 0 |
| Malignant Neoplasms of Ovary | 2 | 385889 | 404051 | 367727 | -0.78 | 4.3E-01 | 0 |
| Malignant Neoplasms of Skin | 47 | 564945 | 902261 | 132566 | -0.72 | 4.7E-01 | 0 |
| Structure of Pelvic Region | 41 | 559165 | 997142 | 108174 | -0.72 | 4.7E-01 | 0 |
| Other disorders of pigmentation | 5 | 433813 | 626938 | 424865 | -0.61 | 5.4E-01 | 0 |
| Maligrant Neoplasms of Pancreas | 3 | 415423 | 739820 | 277350 | -0.58 | 5.6E-01 | 0 |
| Pain in Chest | 4 | 460186 | 939378 | 75222 | -0.58 | 5.6E-01 | 0 |
| Malignant Neoplasms of Male Genital Organs | 26 | 563086 | 1018012 | 155701 | -0.56 | 5.8E-01 | 0 |
| Gastro-Oesophageal Reflux Disease | 2 | 492105 | 716252 | 267957 | -0.51 | 6.1E-01 | 0 |
| Structure of Cardiovascular System | 14 | 530470 | 774106 | 230354 | -0.51 | 6.1E-01 | 0 |
| Outcome of Delivery | 2 | 495450 | 726366 | 264534 | -0.50 | 6.2E-01 | 0 |
| Dementia in Alzheimer Disease | 2 | 497100 | 735271 | 258929 | -0.48 | 6.3E-01 | 0 |
| Failure of Genital Response | 1 | 453883 | 453883 | 453883 | -0.43 | 6.6E-01 | 0 |
| Conversation and Use of Communication Devices and Techniques | 61 | 614263 | 1101180 | 90711 | -0.42 | 6.7E-01 | 0 |
| Procreation Functions | 64 | 621131 | 1114773 | 79091 | -0.42 | 6.7E-01 | 0 |
| Obesity | 38 | 602585 | 1028234 | 283569 | -0.41 | 6.9E-01 | 0 |
| Diseases of Pulp and Periapical Tissues | 10 | 551380 | 865095 | 379969 | -0.38 | 7.0E-01 | 0 |
| Walking and Moving | 9 | 552143 | 836723 | 398870 | -0.35 | 7.3E-01 | 0 |
| Glaucoma | 65 | 633208 | 1048700 | 143303 | -0.30 | 7.6E-01 | 0 |
| Renal Failure | 92 | 642649 | 1082311 | 169460 | -0.29 | 7.7E-01 | 0 |
| Psychomotor Functions | 39 | 627113 | 991665 | 187568 | -0.26 | 8.0E-01 | 0 |
| Hearing Functions | 52 | 642079 | 1078489 | 96134 | -0.20 | 8.4E-01 | 0 |
| Maligrant Neoplasms of Colon | 10 | 616168 | 981138 | 374662 | -0.16 | 8.7E-01 | 0 |
| Personal History of Malignant Neoplasm | 1 | 609465 | 609465 | 609465 | -0.14 | 8.9E-01 | 0 |
| Gonarthrosis [Arthrosis of Knee] | 3 | 614298 | 1249150 | 476524 | -0.11 | 9.1E-01 | 0 |
| Multiple Sclerosis | 18 | 636436 | 1019228 | 117572 | -0.10 | 9.2E-01 | 0 |
| Abnormalities of Breathing | 24 | 651598 | 1131414 | 293286 | -0.07 | 9.5E-01 | 0 |
| Presence of Other Device | 3 | 626783 | 945589 | 324752 | -0.06 | 9.5E-01 | 0 |
| Informal Social Relationships | 87 | 667720 | 1213893 | 117543 | -0.06 | 9.5E-01 | 0 |
| Other Hypothyroidism | 47 | 665375 | 1118659 | 398794 | -0.05 | 9.6E-01 | 0 |
| Moving Around Using Transportation | 6 | 637646 | 734011 | 210343 | -0.05 | 9.6E-01 | 0 |
| Muscle Power Functions | 2 | 691033 | 736609 | 645456 | 0.03 | 9.7E-01 | 0 |

| | | | | | | | |
|---|---|---|---|---|---|---|---|
| Breast and Nipple | 2 | 721569 | 763804 | 679333 | 0.08 | 9.3E-01 | 0 |
| Disorders of Puberty, Not Elsewhere Classified | 95 | 680763 | 1100234 | 148938 | 0.09 | 9.3E-01 | 0 |
| Maligrant Neoplasms of Brain | 23 | 689958 | 1039025 | 281480 | 0.13 | 9.0E-01 | 0 |
| Other Bacterial Diseases | 2 | 730364 | 870164 | 590563 | 0.13 | 9.0E-01 | 0 |
| Other degenerative diseases of the nervous system | 3 | 719763 | 804408 | 387647 | 0.17 | 8.7E-01 | 0 |
| Structure of Mouth | 61 | 697030 | 1082750 | 113557 | 0.19 | 8.5E-01 | 0 |
| Diabetes Mellitus | 71 | 698965 | 1085338 | 127300 | 0.25 | 8.0E-01 | 0 |
| Discomfort associated with menopause | 8 | 744573 | 1367268 | 434703 | 0.31 | 7.6E-01 | 0 |
| Sensations Associated with Genital and Reproductive Functions | 4 | 762909 | 983202 | 424111 | 0.31 | 7.6E-01 | 0 |
| Thyrotoxicosis | 10 | 740721 | 1123445 | 424717 | 0.31 | 7.5E-01 | 0 |
| Pain in Abdomen and Pelvic | 1 | 845510 | 845510 | 845510 | 0.32 | 7.5E-01 | 0 |
| Schizophrenia/Bipolar Affective Disorder | 152 | 699801 | 1168273 | 96495 | 0.33 | 7.4E-01 | 0 |
| Malignant Neoplasms | 48 | 723851 | 1024151 | 322100 | 0.36 | 7.2E-01 | 0 |
| Vitiligo | 34 | 731529 | 1131053 | 99137 | 0.38 | 7.0E-01 | 0 |
| Conduct Disorder | 2 | 830614 | 1216454 | 444773 | 0.39 | 6.9E-01 | 0 |
| Other Arthrosis | 84 | 712899 | 1059573 | 205613 | 0.40 | 6.9E-01 | 0 |
| Vasomotor and Allergic Rhinitis | 203 | 701663 | 1156256 | 141397 | 0.42 | 6.8E-01 | 0 |
| Vascular Syndromes of Brain in Cerebrovascular Diseases | 1 | 928078 | 928078 | 928078 | 0.46 | 6.5E-01 | 0 |
| Endocrine Gland Functions | 115 | 719743 | 928501 | 412455 | 0.51 | 6.1E-01 | 0 |
| Other Congenital Malformations of the Dygestive System | 6 | 818603 | 1039886 | 334516 | 0.53 | 5.9E-01 | 0 |
| Migraine | 13 | 800740 | 1061773 | 406448 | 0.54 | 5.9E-01 | 0 |
| Mild Mental Retardation | 37 | 755425 | 1081700 | 274398 | 0.55 | 5.8E-01 | 0 |
| Unspecified Haematuria | 4 | 841234 | 1251904 | 394397 | 0.56 | 5.7E-01 | 0 |
| Inguinal Hernia | 14 | 820809 | 1125520 | 73491 | 0.67 | 5.0E-01 | 0 |
| Systemic Lupus Erythematosus | 61 | 756663 | 1285880 | 293270 | 0.68 | 5.0E-01 | 0 |
| Producing Body Lagnuage | 1 | 1052485 | 1052485 | 1052485 | 0.71 | 4.8E-01 | 0 |
| Disorders of Lipoprotein Metabolism and Other lipidaemias | 21 | 818555 | 1079695 | 214394 | 0.73 | 4.6E-01 | 0 |
| Other Diseases of Respiratory System | 6 | 873483 | 1013531 | 497613 | 0.74 | 4.6E-01 | 0 |
| Asthma | 218 | 724161 | 1100698 | 243253 | 0.79 | 4.3E-01 | 0 |
| Presence of Cardiac and Vascular Implants and Grafts | 9 | 890533 | 1034870 | 678693 | 0.80 | 4.3E-01 | 0 |
| Hyperkinetic Disorders | 11 | 892855 | 1212943 | 497249 | 0.84 | 4.0E-01 | 0 |
| Disorders related to length of gestation and fetal growth | 7 | 924228 | 1220385 | 660623 | 0.85 | 4.0E-01 | 0 |
| Persons with Potential Health Hazards Related to Socioeconomic and Psychosocial Ci | 3 | 1023008 | 1167565 | 854443 | 0.88 | 3.8E-01 | 0 |
| Structure of hair | 2 | 1008821 | 1495008 | 522634 | 0.89 | 3.8E-01 | 0 |
| Cholelithiasis | 15 | 881008 | 1127658 | 80472 | 0.89 | 3.7E-01 | 0 |
| Malignant Neoplasms of Lung | 3 | 1026643 | 1160283 | 980650 | 0.90 | 3.7E-01 | 0 |
| Superficial Injuries Involving Multiple Body Regions | 30 | 832468 | 1106505 | 287249 | 0.93 | 3.5E-01 | 0 |
| Personal History of Medical Treatment | 3 | 1053010 | 1791330 | 1049563 | 0.96 | 3.4E-01 | 0 |
| Thyroiditis | 75 | 783140 | 1139713 | 327709 | 0.98 | 3.3E-01 | 0 |
| Family History of Certain Disabilities and Chronic Diseases Leading to Disablement | 6 | 964495 | 1123575 | 380211 | 1.01 | 3.1E-01 | 0 |
| Undescended Testicle | 5 | 1031778 | 1061213 | 78554 | 1.06 | 2.9E-01 | 0 |

| | | | | | | | |
|---|---|---|---|---|---|---|---|
| Protein Level | 289 | 736593 | 1116948 | 213623 | 1.09 | 2.8E-01 | 0 |
| Maligrant Neoplasms of Stomach | 4 | 1026389 | 1326258 | 716506 | 1.11 | 2.7E-01 | 0 |
| Schizophrenia | 196 | 751410 | 1219578 | 126049 | 1.12 | 2.6E-01 | 0 |
| Diverticular Disease of Intestine | 76 | 797933 | 1140625 | 157939 | 1.13 | 2.6E-01 | 0 |
| Angina Pectoris | 26 | 881300 | 1172282 | 321236 | 1.14 | 2.5E-01 | 0 |
| Hypertrichosis | 3 | 1145635 | 1332681 | 912813 | 1.19 | 2.3E-01 | 0 |
| Atrial Fibrillation and Flutter | 186 | 759971 | 1150789 | 337274 | 1.21 | 2.3E-01 | 0 |
| Pervasive Developmental Disorders | 3 | 1164070 | 1190443 | 970190 | 1.22 | 2.2E-01 | 0 |
| Bipolar Affective Disorder | 12 | 984339 | 1212962 | 495684 | 1.27 | 2.0E-01 | 0 |
| Other Cataract | 6 | 1048938 | 1397449 | 342022 | 1.27 | 2.0E-01 | 0 |
| Functions of the Cardiovascular System | 6 | 1052319 | 1121252 | 894963 | 1.28 | 2.0E-01 | 0 |
| Heart Functions | 530 | 730185 | 1152666 | 204447 | 1.31 | 1.9E-01 | 0 |
| Structure of Brain | 574 | 728523 | 1149092 | 160080 | 1.33 | 1.8E-01 | 0 |
| Higher-Level Cognitive Functions | 443 | 738595 | 1133618 | 94991 | 1.40 | 1.6E-01 | 0 |
| Chronic Ischaemic Heart Disease | 425 | 740605 | 1120763 | 322275 | 1.40 | 1.6E-01 | 0 |
| Structure of Head and Neck Region | 59 | 853965 | 1291146 | 404181 | 1.44 | 1.5E-01 | 0 |
| Hematological System Functions | 32 | 917845 | 1122026 | 347349 | 1.45 | 1.5E-01 | 0 |
| Dorsopathies | 16 | 1009141 | 1215242 | 412496 | 1.53 | 1.3E-01 | 0 |
| Control of Voluntary Movement Functions | 1 | 1525045 | 1525045 | 1525045 | 1.58 | 1.1E-01 | 0 |
| Alzheimer disease | 39 | 918405 | 1219690 | 124168 | 1.58 | 1.1E-01 | 0 |
| Problems Related to Lifestyle | 1 | 1529998 | 1529998 | 1529998 | 1.60 | 1.1E-01 | 0 |
| Obstruction of blie duct | 3 | 1330175 | 1602635 | 1157248 | 1.62 | 1.0E-01 | 0 |
| Epilepsy | 21 | 1021760 | 1437625 | 722640 | 1.70 | 9.0E-02 | 0 |
| Structures Related to the Genitourinary and Reproductive Systems, Other Specified | 1 | 1613123 | 1613123 | 1613123 | 1.76 | 7.9E-02 | 0 |
| Other Rheumatoid Arthritis | 66 | 886414 | 1181998 | 439329 | 1.77 | 7.7E-02 | 0 |
| Essential (Primary) Hypertension | 210 | 796706 | 1132637 | 155420 | 1.78 | 7.5E-02 | 0 |
| Diaphragmatic Hernia | 3 | 1398910 | 1434606 | 1106865 | 1.80 | 7.1E-02 | 0 |
| Structure of Trunk | 37 | 955958 | 1314745 | 427510 | 1.81 | 7.1E-02 | 0 |
| Type 2 Diabetes Mellitus | 534 | 755668 | 1098538 | 363788 | 1.95 | 5.1E-02 | 0 |
| Parkinson Disease | 28 | 1028789 | 1293166 | 481049 | 1.99 | 4.7E-02 | 0 |
| Spinal Muscular Strophy and Related Syndrome | 10 | 1199350 | 1369743 | 217373 | 2.03 | 4.2E-02 | 0 |
| Atopic Dermatitis | 22 | 1084311 | 1379780 | 553161 | 2.08 | 3.7E-02 | 0 |
| Sensation of Pain | 59 | 948753 | 1173568 | 241821 | 2.13 | 3.3E-02 | 0 |
| Type 1 Diabetes Mellitus | 79 | 922743 | 1205271 | 390338 | 2.26 | 2.4E-02 | 0 |
| Internal Derangement of Knee | 1 | 1884003 | 1884003 | 1884003 | 2.28 | 2.3E-02 | 0 |
| Menstruation Functions | 545 | 770950 | 1157570 | 269260 | 2.29 | 2.2E-02 | 0 |
| Structures related to movement, unspecified | 933 | 746725 | 1142685 | 178645 | 2.29 | 2.2E-02 | 0 |
| Sleep Functions | 699 | 768070 | 1191784 | 126304 | 2.50 | 1.2E-02 | 0 |
| Sleep Disorders | 74 | 970969 | 1183357 | 116859 | 2.64 | 8.2E-03 | 0 |
| Hyperhidrosis, unspecified | 2 | 1686329 | 1737813 | 1634844 | 2.68 | 7.4E-03 | 0 |
| Functions of Hair | 718 | 774485 | 1109228 | 299151 | 2.70 | 6.9E-03 | 0 |

| | | | | | | | |
|---|---|---|---|---|---|---|---|
| Water, Mineral and Electrolyte Balance Functions | 1438 | 750085 | 1124753 | 260158 | 2.96 | 3.1E-03 | 0 |
| Blood Pressure Functions | 766 | 781438 | 1136306 | 176599 | 3.06 | 2.2E-03 | 0 |
| Products or Substances for Personal Consumption | 599 | 803743 | 1150583 | 266960 | 3.25 | 1.1E-03 | 0 |
| Exercise Tolerance Functions | 1355 | 763848 | 1164004 | 205416 | 3.36 | 7.9E-04 | 0 |
| Potential Health Hazards Related to Socioeconomic and Psychosocial Circumstances | 142 | 953724 | 1197395 | 307947 | 3.46 | 5.5E-04 | 0 |
| Temperament and Personality Functions | 1068 | 804389 | 1196450 | 134519 | 4.34 | 1.4E-05 | 1 |
| Immunological System Functions | 4197 | 747780 | 1134575 | 207559 | 5.01 | 5.5E-07 | 1 |
| Height | 3284 | 766455 | 1142876 | 207088 | 5.48 | 4.3E-08 | 1 |
| Weight Maintenance Functions | 7813 | 802540 | 1184605 | 198782 | 12.80 | 1.6E-37 | 1 |

| Trait | N SNPs | Median age | Age 75% upper bound | Age 25% lower bound | Z-score | P value | Bonferroni |
|---|---|---|---|---|---|---|---|
| Vertical cup-disc ratio | 515 | 23661 | 80315 | 17915 | -14.73 | 3.9E-49 | 1 |
| Depression - Lifetime number of depressed periods | 395 | 21076 | 43485 | 13709 | -12.96 | 2.1E-38 | 1 |
| Lifetime number of sexual partners | 337 | 32349 | 164566 | 15909 | -11.98 | 4.4E-33 | 1 |
| Average weekly intake of other alcoholic drinks | 281 | 20258 | 42061 | 14154 | -11.07 | 1.7E-28 | 1 |
| Number of depression episodes | 270 | 30529 | 194139 | 16894 | -10.66 | 1.5E-26 | 1 |
| Childhood sunburn occasions | 271 | 51971 | 322210 | 18685 | -10.22 | 1.6E-24 | 1 |
| Distance between home and job workplace | 249 | 83490 | 427368 | 22529 | -9.45 | 3.5E-21 | 1 |
| Frequency of solarium/sunlamp use | 184 | 80130 | 334445 | 29208 | -8.10 | 5.4E-16 | 1 |
| Salad / raw vegetable intake | 162 | 194781 | 777221 | 51721 | -6.22 | 5.0E-10 | 1 |
| Intraocular pressure | 83 | 25426 | 312584 | 16388 | -5.92 | 3.3E-09 | 1 |
| Average weekly fortified wine intake | 132 | 201850 | 506902 | 73456 | -5.57 | 2.5E-08 | 1 |
| Cup area | 57 | 52302 | 742885 | 20433 | -4.75 | 2.1E-06 | 1 |
| Frequency of travelling from home to job workplace | 71 | 134420 | 503480 | 33335 | -4.62 | 3.8E-06 | 1 |
| Processed meat intake | 76 | 160045 | 758022 | 54394 | -4.54 | 5.7E-06 | 1 |
| Average weekly spirits intake | 56 | 95610 | 352450 | 35048 | -4.42 | 9.7E-06 | 1 |
| Prospective memory test - Number of attempts | 68 | 190118 | 553660 | 45600 | -4.08 | 4.5E-05 | 0 |
| Pork intake | 56 | 134221 | 580550 | 43214 | -4.08 | 4.6E-05 | 0 |
| Number in household | 52 | 168502 | 536184 | 60810 | -3.66 | 2.5E-04 | 0 |
| Ease of skin tanning | 445 | 503970 | 1007923 | 66422 | -3.64 | 2.8E-04 | 0 |
| Number of children fathered (male) | 41 | 124491 | 467298 | 61860 | -3.51 | 4.4E-04 | 0 |
| Frequency of needing morning drink of alcohol after heavy drinking session in last year | 33 | 60270 | 333940 | 28723 | -3.51 | 4.5E-04 | 0 |
| Dried fruit intake | 49 | 178220 | 554673 | 51945 | -3.50 | 4.6E-04 | 0 |
| Exposure to tobacco smoke outside home | 58 | 251797 | 480483 | 65161 | -3.25 | 1.1E-03 | 0 |
| Number of unsuccessful stop-smoking attempts | 22 | 51853 | 399298 | 25193 | -3.05 | 2.3E-03 | 0 |
| Time spent using computer | 30 | 162893 | 532663 | 65235 | -2.92 | 3.5E-03 | 0 |
| Hair colour (natural, before greying): Dark brown | 224 | 481623 | 953399 | 77350 | -2.90 | 3.7E-03 | 0 |
| Cooked vegetable intake | 40 | 229193 | 545094 | 92228 | -2.81 | 5.0E-03 | 0 |
| Number of full brothers | 33 | 191690 | 968785 | 38344 | -2.76 | 5.8E-03 | 0 |
| Average total household income before tax | 47 | 284860 | 1091773 | 57845 | -2.71 | 6.7E-03 | 0 |
| Use of sun/uv protection | 28 | 185084 | 995476 | 56749 | -2.64 | 8.3E-03 | 0 |
| Breast cancer (GEE, adjusted for age) | 178 | 482129 | 987751 | 102244 | -2.62 | 8.8E-03 | 0 |
| Exposure to tobacco smoke at home | 45 | 292468 | 551775 | 78580 | -2.58 | 9.9E-03 | 0 |
| Breast cancer (GEE, adjusted for age and BMI) | 186 | 486744 | 941669 | 101316 | -2.58 | 1.0E-02 | 0 |
| Disc area | 45 | 303970 | 1027155 | 28035 | -2.50 | 1.3E-02 | 0 |
| Carpal tunnel syndrome | 12 | 89668 | 637349 | 43850 | -2.23 | 2.6E-02 | 0 |
| Fluid intelligence test - Fluid intelligence score | 13 | 82009 | 877208 | 57133 | -2.20 | 2.8E-02 | 0 |
| Glycine level (male) | 12 | 102899 | 563625 | 27396 | -2.16 | 3.1E-02 | 0 |
| Fresh fruit intake | 59 | 396183 | 974729 | 71794 | -2.14 | 3.2E-02 | 0 |
| Longest period of depression | 9 | 27979 | 79346 | 14239 | -2.09 | 3.6E-02 | 0 |
| Neutrophil percentage of white cells (three-way meta) | 95 | 460268 | 1110643 | 77813 | -2.06 | 3.9E-02 | 0 |
| Legs-leg fat ratio (female) | 147 | 506280 | 1151456 | 67821 | -2.02 | 4.3E-02 | 0 |
| Left hippocampus | 8 | 76249 | 633694 | 42060 | -2.00 | 4.6E-02 | 0 |

| | | | | | | | |
|---|---|---|---|---|---|---|---|
| Pulse pressure | 26 | 290495 | 916129 | 71718 | -1.98 | 4.7E-02 | 0 |
| Number of full sisters | 37 | 348723 | 761838 | 20181 | -1.97 | 4.9E-02 | 0 |
| Prospective memory test - Duration screen displayed | 8 | 86876 | 150747 | 47791 | -1.95 | 5.1E-02 | 0 |
| Cereal type: Muesli | 9 | 73237 | 222987 | 41485 | -1.93 | 5.4E-02 | 0 |
| Hair colour (natural, before greying): Light brown | 77 | 457560 | 761985 | 73528 | -1.92 | 5.5E-02 | 0 |
| Other sociodemographic factors - Private healthcare | 4 | 17123 | 29635 | 10233 | -1.90 | 5.7E-02 | 0 |
| Self-rated health | 6 | 71608 | 174189 | 56422 | -1.86 | 6.3E-02 | 0 |
| Intracranial Volume | 6 | 86245 | 470171 | 44418 | -1.84 | 6.5E-02 | 0 |
| Insomnia (male) | 4 | 33202 | 35982 | 30368 | -1.83 | 6.7E-02 | 0 |
| Sleep efficiency | 4 | 34179 | 44624 | 29291 | -1.83 | 6.7E-02 | 0 |
| L5 timing | 4 | 46067 | 359654 | 40801 | -1.83 | 6.8E-02 | 0 |
| Cholesterol esters in large VLDL | 11 | 154010 | 792240 | 67378 | -1.82 | 6.9E-02 | 0 |
| Trail making test - Duration to complete numeric path (trail #1) | 4 | 54729 | 368738 | 28919 | -1.80 | 7.2E-02 | 0 |
| Number of stillbirths (female) | 2 | 21325 | 24569 | 18081 | -1.79 | 7.3E-02 | 0 |
| Ever taken oral contraceptive pill (female) | 2 | 15392 | 16507 | 14276 | -1.79 | 7.4E-02 | 0 |
| Diagnoses - main ICD10: D12 Benign neoplasm of colon, rectum, anus and anal canal | 13 | 196128 | 895060 | 112635 | -1.78 | 7.4E-02 | 0 |
| Illnesses of father: Lung cancer | 4 | 51982 | 283413 | 43957 | -1.78 | 7.5E-02 | 0 |
| Brain stem | 16 | 249259 | 719940 | 68144 | -1.75 | 7.9E-02 | 0 |
| High-density lipoprotein 2 cholesterol (at exam 5, GEE, adjusted for multivariable) | 7 | 83680 | 606239 | 72304 | -1.75 | 8.0E-02 | 0 |
| mDC:%32+ | 271 | 566700 | 1021561 | 156415 | -1.75 | 8.0E-02 | 0 |
| Average weekly beer plus cider intake | 19 | 270300 | 910450 | 46256 | -1.75 | 8.0E-02 | 0 |
| Acceleration average | 8 | 147557 | 1199616 | 22268 | -1.75 | 8.1E-02 | 0 |
| von Willebrand Factor (at exam 5, GEE, adjusted for multivariable) | 4 | 67704 | 275860 | 52308 | -1.74 | 8.2E-02 | 0 |
| QT interval (cohort exam 11 offspring exam 1, GEE, adjusted for age nad RR) | 2 | 34503 | 43208 | 25797 | -1.73 | 8.3E-02 | 0 |
| Traumatic events - Someone to take to doctor when needed as a child | 2 | 35129 | 45148 | 25110 | -1.73 | 8.3E-02 | 0 |
| Low-density lipoprotein cholesterol (at exam 5, GEE, adjusted for age and sex) | 2 | 37573 | 38652 | 36493 | -1.72 | 8.5E-02 | 0 |
| Handedness (chirality/laterality) | 2 | 40610 | 48418 | 32802 | -1.72 | 8.6E-02 | 0 |
| HP - Haptoglobin | 2 | 44588 | 56868 | 32309 | -1.72 | 8.6E-02 | 0 |
| Reported occurrences of cancer | 2 | 37570 | 37957 | 37182 | -1.71 | 8.6E-02 | 0 |
| Doctor diagnosed asthma | 14 | 244103 | 755139 | 51321 | -1.70 | 8.9E-02 | 0 |
| Morbidity-free survival (at age 65, GEE, adjusted for birth-cohort) | 37 | 394573 | 904338 | 106636 | -1.70 | 9.0E-02 | 0 |
| Left thalamus proper | 2 | 49585 | 51112 | 48058 | -1.70 | 9.0E-02 | 0 |
| Spine Neck BMD (GEE, male, adjusted for multivariable) | 2 | 41796 | 44704 | 38889 | -1.70 | 9.0E-02 | 0 |
| BRCA1/2-negative breast cancer | 2 | 50715 | 51587 | 49842 | -1.68 | 9.2E-02 | 0 |
| Hip circumference (female) | 33 | 382215 | 1035158 | 79540 | -1.67 | 9.5E-02 | 0 |
| SDNN HRV (cohort exam 18 offsprint exam 3, GEE, adjusted for age and HR) | 2 | 55596 | 69385 | 41806 | -1.66 | 9.6E-02 | 0 |
| Ever depressed for a whole week | 5 | 56776 | 401733 | 42107 | -1.65 | 9.9E-02 | 0 |
| Schizophrenia/Bipolar disorder | 86 | 499765 | 1068881 | 81502 | -1.65 | 1.0E-01 | 0 |
| Usual weekday sleep duration (GEE, adjusted for age, sex and BMI) | 2 | 69364 | 69517 | 69212 | -1.64 | 1.0E-01 | 0 |
| Left precentral | 5 | 63278 | 867898 | 62100 | -1.64 | 1.0E-01 | 0 |
| Interpolated Age of participant when non-cancer illness first diagnosed | 6 | 144197 | 630851 | 89913 | -1.63 | 1.0E-01 | 0 |
| Interpolated Year when non-cancer illness first diagnosed | 6 | 144197 | 630851 | 89913 | -1.63 | 1.0E-01 | 0 |
| ::::X-11491 | 2 | 74898 | 87034 | 62762 | -1.61 | 1.1E-01 | 0 |

| | | | | | | | |
|---|---|---|---|---|---|---|---|
| Anterior limb of internal capsule mode of anisotropy | 4 | 113774 | 254376 | 55525 | -1.61 | 1.1E-01 | 0 |
| Suffer from 'nerves' (SUF-NERV) | 5 | 79348 | 586548 | 63709 | -1.61 | 1.1E-01 | 0 |
| Non-oily fish intake | 10 | 216765 | 424298 | 83193 | -1.61 | 1.1E-01 | 0 |
| Body Mass Index - Child | 17 | 292630 | 513870 | 76367 | -1.60 | 1.1E-01 | 0 |
| Mean Hippocampus | 2 | 81421 | 105141 | 57701 | -1.59 | 1.1E-01 | 0 |
| Peptide::gamma-glutamyl::gamma-glutamyltyrosine | 4 | 118692 | 357965 | 97545 | -1.59 | 1.1E-01 | 0 |
| Left lateral ventricle | 9 | 182377 | 474765 | 97114 | -1.58 | 1.2E-01 | 0 |
| Milk type used: Soya | 5 | 85606 | 1419258 | 53268 | -1.57 | 1.2E-01 | 0 |
| Cancer register - Type of cancer: ICD10: C44 Other and unspecified malignant neoplasm of skin | 28 | 376049 | 766590 | 99630 | -1.57 | 1.2E-01 | 0 |
| Amino acid::Valine, leucine and isoleucine metabolism::isovalerylcarnitine | 2 | 89642 | 110107 | 69178 | -1.56 | 1.2E-01 | 0 |
| Beef intake | 6 | 176647 | 724122 | 78859 | -1.56 | 1.2E-01 | 0 |
| Insulin sensitivity (GEE, adjusted for age and sex) | 5 | 80097 | 92573 | 72101 | -1.56 | 1.2E-01 | 0 |
| Triglycerides (at exam 5, GEE, adjusted for age and sex) | 2 | 102186 | 125350 | 79021 | -1.55 | 1.2E-01 | 0 |
| Electronic device use - Plays computer games | 27 | 378423 | 1017605 | 84049 | -1.54 | 1.2E-01 | 0 |
| Number of vehicles in household | 5 | 100707 | 138743 | 58540 | -1.53 | 1.3E-01 | 0 |
| Sagittal stratum mean diusivities | 3 | 23901 | 359703 | 23073 | -1.53 | 1.3E-01 | 0 |
| Thinness | 3 | 33968 | 518521 | 23297 | -1.52 | 1.3E-01 | 0 |
| Sagittal stratum radial diusivities | 3 | 23901 | 474499 | 23073 | -1.52 | 1.3E-01 | 0 |
| Interferon gamma-induced protein 10 (CXCL10) | 3 | 28732 | 811925 | 26743 | -1.52 | 1.3E-01 | 0 |
| TNF-related apoptosis inducing ligand | 3 | 24513 | 36174 | 22204 | -1.51 | 1.3E-01 | 0 |
| Interleukin 6 (at exam 7, GEE, adjusted for age and sex) | 2 | 114955 | 144786 | 85124 | -1.51 | 1.3E-01 | 0 |
| Pain type(s) experienced in last month: Neck or shoulder pain | 5 | 108061 | 787273 | 30242 | -1.50 | 1.3E-01 | 0 |
| Mouth/teeth dental problems: Loose teeth | 5 | 113557 | 782813 | 58698 | -1.50 | 1.3E-01 | 0 |
| MST1 - Hepatocyte growth factor-like protein | 3 | 44586 | 123463 | 38980 | -1.50 | 1.3E-01 | 0 |
| Ulcerative colitis | 294 | 587378 | 918819 | 199182 | -1.49 | 1.4E-01 | 0 |
| Cannabis use - Ever taken cannabis | 9 | 204782 | 1218098 | 50761 | -1.49 | 1.4E-01 | 0 |
| HOMA-IR (at exam 5, GEE, adjusted for age and sex) | 3 | 42572 | 198497 | 42306 | -1.49 | 1.4E-01 | 0 |
| Morbidity-free survival (at age 65, GEE, adjusted for multivariable) | 37 | 430495 | 904338 | 166969 | -1.48 | 1.4E-01 | 0 |
| Depression - Recent thoughts of suicide or self-harm | 4 | 156480 | 234614 | 123060 | -1.48 | 1.4E-01 | 0 |
| Right vessel | 2 | 133376 | 148423 | 118329 | -1.48 | 1.4E-01 | 0 |
| Usual side of head for mobile phone use | 2 | 129096 | 151873 | 106319 | -1.47 | 1.4E-01 | 0 |
| Lipid::Lysolipid::1-eicosatrienoylglycerophosphocholine* | 2 | 124936 | 151070 | 98801 | -1.47 | 1.4E-01 | 0 |
| Left precuneus | 3 | 46213 | 294738 | 46117 | -1.46 | 1.4E-01 | 0 |
| Length of time at current address | 5 | 123446 | 1197513 | 64740 | -1.46 | 1.4E-01 | 0 |
| Fluid intelligence score | 47 | 459465 | 1121114 | 81313 | -1.46 | 1.5E-01 | 0 |
| Plasma Apolipoprotein E level (at exam 5, GEE, adjusted for multivariable) | 3 | 51715 | 304511 | 39040 | -1.46 | 1.5E-01 | 0 |
| Total body BMD (30-45 years old) | 7 | 176519 | 813318 | 71063 | -1.46 | 1.5E-01 | 0 |
| Left lateral occipital | 3 | 57675 | 162368 | 46604 | -1.46 | 1.5E-01 | 0 |
| Neck Section Modulus (GEE, male, adjusted for multivariable) | 3 | 60104 | 98041 | 53237 | -1.45 | 1.5E-01 | 0 |
| Smoking cessation | 5 | 125509 | 875095 | 89821 | -1.45 | 1.5E-01 | 0 |
| TNC - Tenascin | 2 | 141702 | 145360 | 138044 | -1.45 | 1.5E-01 | 0 |
| Hands-free device/speakerphone use with mobile phone in last 3 month | 3 | 58721 | 336492 | 55004 | -1.45 | 1.5E-01 | 0 |
| High-density lipoprotein 2 cholesterol (at exam 4, GEE, adjusted for multivariable) | 7 | 179200 | 718004 | 109570 | -1.45 | 1.5E-01 | 0 |

| | | | | | | | |
|---|---|---|---|---|---|---|---|
| Cytomegalovirus IgG levels | 2 | 143819 | 203686 | 83952 | -1.44 | 1.5E-01 | 0 |
| Average across all tracts fractional anisotropy | 3 | 61894 | 460397 | 46179 | -1.44 | 1.5E-01 | 0 |
| Anterior limb of internal capsule fractional anisotropy | 2 | 143717 | 197707 | 89727 | -1.44 | 1.5E-01 | 0 |
| Depression - Recent feelings of tiredness or low enery | 3 | 66860 | 774248 | 55193 | -1.42 | 1.6E-01 | 0 |
| Major dietary changes in the last 5 years | 9 | 229329 | 889053 | 69644 | -1.42 | 1.6E-01 | 0 |
| Left cuneus | 3 | 74601 | 624878 | 66138 | -1.41 | 1.6E-01 | 0 |
| Nucleotide::Pyrimidine metabolism, uracil containing::uridine | 3 | 71641 | 618145 | 70840 | -1.41 | 1.6E-01 | 0 |
| Frequency of drinking alcohol | 6 | 221736 | 436783 | 67929 | -1.40 | 1.6E-01 | 0 |
| Mean Putamen | 3 | 75055 | 662063 | 63835 | -1.40 | 1.6E-01 | 0 |
| Plasma Apolipoprotein A-I level (GEE, multivariate adjusted) | 2 | 164747 | 227617 | 101877 | -1.40 | 1.6E-01 | 0 |
| Plasma Apolipoprotein A-I level (GEE, adjusted for age and sex) | 2 | 166038 | 228263 | 103813 | -1.40 | 1.6E-01 | 0 |
| Thyroid stimulation hormone (at exam 4, GEE, adjusted for age and sex) | 2 | 173253 | 208335 | 138172 | -1.38 | 1.7E-01 | 0 |
| Intermediate-density lipoprotein by NMR (at exam 4, GEE, adjusted for age and sex) | 11 | 266905 | 946700 | 66152 | -1.38 | 1.7E-01 | 0 |
| Reason for reducing amount of alcohol drunk: Health precaution | 2 | 179762 | 243505 | 116020 | -1.37 | 1.7E-01 | 0 |
| Amyotrophic lateral sclerosis (meta-analysis) | 3 | 87092 | 675340 | 51090 | -1.36 | 1.7E-01 | 0 |
| Mean FEV1 from exam 3 and 5 (GEE, adjusted for multiple covariates) | 3 | 80044 | 242560 | 75360 | -1.36 | 1.7E-01 | 0 |
| College completion | 3 | 95032 | 422132 | 83753 | -1.35 | 1.8E-01 | 0 |
| Legs-leg fat ratio (male) | 20 | 385794 | 1069323 | 78229 | -1.34 | 1.8E-01 | 0 |
| Pairs matching test - Number of incorrect matches in round | 6 | 239202 | 269988 | 95049 | -1.34 | 1.8E-01 | 0 |
| Traumatic events - Avoided activities or situations because of previous stressful experience in pa | 3 | 100716 | 240462 | 78602 | -1.33 | 1.8E-01 | 0 |
| Viscosity I (at exam 5, GEE, adjusted for multivariable) | 4 | 207244 | 576054 | 67891 | -1.33 | 1.8E-01 | 0 |
| Corticospinal tract fractional anisotropy | 2 | 186521 | 272234 | 100808 | -1.32 | 1.9E-01 | 0 |
| Platelet aggregation to Epinephrine (at exam 5, GEE, adjusted for multivariable) | 2 | 181712 | 252351 | 111073 | -1.32 | 1.9E-01 | 0 |
| Large high-density lipoprotein by NMR (at exam 4, GEE, adjusted for multivariable) | 4 | 210962 | 496726 | 102327 | -1.32 | 1.9E-01 | 0 |
| Lipid::Carnitine metabolism::X-13431--nonanoylcarnitine* | 2 | 178248 | 238599 | 117897 | -1.32 | 1.9E-01 | 0 |
| Sphingomyelins | 7 | 225348 | 1028358 | 41734 | -1.31 | 1.9E-01 | 0 |
| Amino acid::Valine, leucine and isoleucine metabolism::alpha-hydroxyisovalerate | 2 | 199588 | 257028 | 142149 | -1.31 | 1.9E-01 | 0 |
| Dehydroepiandrosterone sulfate (at exam 3, GEE, adjusted for age and sex) | 2 | 187750 | 258885 | 116615 | -1.31 | 1.9E-01 | 0 |
| Fornix (column and body of fornix) fractional anisotropy | 6 | 247492 | 1190918 | 106453 | -1.30 | 1.9E-01 | 0 |
| Trochanter BMD (GEE, adjusted for multivariable) | 2 | 196602 | 268736 | 124468 | -1.30 | 1.9E-01 | 0 |
| Posterior limb of internal capsule mode of anisotropy | 6 | 248493 | 506239 | 154241 | -1.29 | 2.0E-01 | 0 |
| Interferon-gamma | 1 | 13015 | 13015 | 13015 | -1.29 | 2.0E-01 | 0 |
| Plasma Apolipoprotein E level (at exam 5, GEE, adjusted for age and sex) | 4 | 230313 | 446009 | 45378 | -1.29 | 2.0E-01 | 0 |
| Brain natriuretic peptide (at exam 6, GEE, adjusted for multivariable) | 1 | 9422 | 9422 | 9422 | -1.28 | 2.0E-01 | 0 |
| Chronic kidney disease | 56 | 505864 | 1150924 | 78809 | -1.28 | 2.0E-01 | 0 |
| Tumor necrosis factor beta | 1 | 5901 | 5901 | 5901 | -1.27 | 2.0E-01 | 0 |
| Posterior corona radiata axial diusivities | 2 | 208609 | 296688 | 120530 | -1.27 | 2.0E-01 | 0 |
| Phosphorylated Tau at position 181 | 1 | 20758 | 20758 | 20758 | -1.27 | 2.0E-01 | 0 |
| Leucine | 4 | 226721 | 696908 | 55258 | -1.27 | 2.0E-01 | 0 |
| Health satisfaction | 1 | 18576 | 18576 | 18576 | -1.27 | 2.0E-01 | 0 |
| Superior longitudinal fasciculus axial diusivities | 4 | 227778 | 738823 | 63880 | -1.27 | 2.0E-01 | 0 |
| Extreme chronotype | 3 | 130692 | 203013 | 84443 | -1.27 | 2.0E-01 | 0 |
| Maximum carotid artery bulb IMT (at exam 6, GEE, adjusted for multivariable) | 1 | 12886 | 12886 | 12886 | -1.27 | 2.0E-01 | 0 |

| | | | | | | | |
|---|---|---|---|---|---|---|---|
| UCHL1 - Ubiquitin carboxyl-terminal hydrolase isozyme L1 | 1 | 14724 | 14724 | 14724 | -1.27 | 2.0E-01 | 0 |
| ::::X-12092 | 3 | 124129 | 477847 | 98449 | -1.27 | 2.0E-01 | 0 |
| Double eyelid | 2 | 219257 | 228872 | 209641 | -1.27 | 2.1E-01 | 0 |
| Bilirubin (at exam 2, GEE, adjusted for multivariable) | 2 | 216216 | 306358 | 126074 | -1.27 | 2.1E-01 | 0 |
| Right posterior cingulate | 1 | 21558 | 21558 | 21558 | -1.26 | 2.1E-01 | 0 |
| Amino acid::Urea cycle; arginine-, proline-, metabolism::citrulline | 3 | 135205 | 540796 | 96099 | -1.26 | 2.1E-01 | 0 |
| Parietal brain volume (GEE, adjusted for age and sex) | 1 | 24553 | 24553 | 24553 | -1.26 | 2.1E-01 | 0 |
| Peak insulin response (adjusted for BMI) | 3 | 134748 | 419900 | 78553 | -1.26 | 2.1E-01 | 0 |
| Medial collateral ligament injury | 1 | 26829 | 26829 | 26829 | -1.26 | 2.1E-01 | 0 |
| Survival past average life expectancy (GEE, adjusted for multivariable) | 1 | 29042 | 29042 | 29042 | -1.26 | 2.1E-01 | 0 |
| Peptide::Polypeptide::HWESASXX* | 1 | 26849 | 26849 | 26849 | -1.25 | 2.1E-01 | 0 |
| Duration of heavy DIY | 1 | 22587 | 22587 | 22587 | -1.25 | 2.1E-01 | 0 |
| Coffee type: Decaffeinated coffee (any type) | 1 | 32190 | 32190 | 32190 | -1.25 | 2.1E-01 | 0 |
| FRZB - Secreted frizzled-related protein 3 | 1 | 36235 | 36235 | 36235 | -1.25 | 2.1E-01 | 0 |
| THBS1 - Thrombospondin-1 | 1 | 25776 | 25776 | 25776 | -1.25 | 2.1E-01 | 0 |
| Shaft Width (GEE, adjusted for multivariable) | 1 | 26182 | 26182 | 26182 | -1.25 | 2.1E-01 | 0 |
| Xenobiotics::Drug::p-acetamidophenylglucuronide | 1 | 28473 | 28473 | 28473 | -1.25 | 2.1E-01 | 0 |
| Age started oral contraceptive pill (female) | 4 | 240136 | 512628 | 42467 | -1.25 | 2.1E-01 | 0 |
| LMNB1 - Lamin-B1 | 1 | 24460 | 24460 | 24460 | -1.25 | 2.1E-01 | 0 |
| Traumatic events - Felt distant from other people in past month | 1 | 26219 | 26219 | 26219 | -1.25 | 2.1E-01 | 0 |
| Parietal brain volume (GEE, adjusted for multivariable) | 1 | 24553 | 24553 | 24553 | -1.25 | 2.1E-01 | 0 |
| Immunosuppressants | 1 | 24454 | 24454 | 24454 | -1.25 | 2.1E-01 | 0 |
| Hepatocyte growth factor | 1 | 31244 | 31244 | 31244 | -1.24 | 2.1E-01 | 0 |
| Father's age at death | 4 | 234241 | 601883 | 45062 | -1.24 | 2.1E-01 | 0 |
| Rotator cuff injury | 1 | 34759 | 34759 | 34759 | -1.24 | 2.1E-01 | 0 |
| Anti-Mullerian hormone | 1 | 26754 | 26754 | 26754 | -1.24 | 2.1E-01 | 0 |
| Eyebrows | 1 | 36447 | 36447 | 36447 | -1.24 | 2.1E-01 | 0 |
| Diagnoses - secondary ICD10: Z96 Presence of other functional implants | 1 | 23028 | 23028 | 23028 | -1.24 | 2.1E-01 | 0 |
| CCL25 - C-C motif chemokine 25 | 1 | 27893 | 27893 | 27893 | -1.24 | 2.2E-01 | 0 |
| MPO - Myeloperoxidase | 1 | 31385 | 31385 | 31385 | -1.24 | 2.2E-01 | 0 |
| Alcohol - Alcohol drinker status: Previous vs Current | 1 | 25520 | 25520 | 25520 | -1.24 | 2.2E-01 | 0 |
| Shaft Section Modulus (GEE, adjussted for multivariable) | 1 | 23513 | 23513 | 23513 | -1.24 | 2.2E-01 | 0 |
| Frequency of failure to fulfil normal expectations fue to drinking alcohol in last year | 1 | 29260 | 29260 | 29260 | -1.24 | 2.2E-01 | 0 |
| Ankle-brachial index (at exam 7, GEE, adjusted for multivariable) | 1 | 29042 | 29042 | 29042 | -1.24 | 2.2E-01 | 0 |
| Tinnitus | 1 | 27175 | 27175 | 27175 | -1.24 | 2.2E-01 | 0 |
| Frequency of heavy DIY in last 4 weeks | 1 | 35995 | 35995 | 35995 | -1.24 | 2.2E-01 | 0 |
| Left handed | 1 | 40135 | 40135 | 40135 | -1.23 | 2.2E-01 | 0 |
| FUT5 - Alpha-(1,3)-fucosyltransferase 5 | 1 | 19351 | 19351 | 19351 | -1.23 | 2.2E-01 | 0 |
| Regulated on Activation, Normal T Cell Expressed and Secreted (CCL5) | 1 | 27087 | 27087 | 27087 | -1.23 | 2.2E-01 | 0 |
| Alcohol dependency (unrelated genotyped individuals) | 1 | 33942 | 33942 | 33942 | -1.23 | 2.2E-01 | 0 |
| Remnant lipoprotein cholesterol (at exam 4, GEE, adjusted for age and sex) | 1 | 48458 | 48458 | 48458 | -1.23 | 2.2E-01 | 0 |
| Depression - Did your sleep change? | 1 | 26515 | 26515 | 26515 | -1.23 | 2.2E-01 | 0 |
| Left fusiform | 4 | 234976 | 643218 | 18827 | -1.23 | 2.2E-01 | 0 |

| | | | | | | | |
|---|---|---|---|---|---|---|---|
| Type 2 Diabetes (Dominance deviation model) | 1 | 24626 | 24626 | 24626 | -1.23 | 2.2E-01 | 0 |
| Coronary heart disease (MI, CI or CHD death, GEE, adjusted for age and sex) | 2 | 221913 | 299518 | 144308 | -1.23 | 2.2E-01 | 0 |
| CXCL6 - C-X-C motif chemokine 6 | 1 | 36394 | 36394 | 36394 | -1.23 | 2.2E-01 | 0 |
| Fornix (column and body of fornix) mean diusivities | 4 | 247492 | 714670 | 145666 | -1.23 | 2.2E-01 | 0 |
| Interleukin-2 receptor, alpha subunit | 1 | 23983 | 23983 | 23983 | -1.23 | 2.2E-01 | 0 |
| Neuroticism (IRT) | 1 | 30622 | 30621 | 30621 | -1.22 | 2.2E-01 | 0 |
| Maternal smoking around birth | 14 | 364589 | 708594 | 50867 | -1.22 | 2.2E-01 | 0 |
| Fasting insulin interaction (adjusted for BMI) | 10 | 316486 | 995656 | 37916 | -1.22 | 2.2E-01 | 0 |
| NAAA - N-acylethanolamine-hydrolyzing acid amidase | 2 | 228643 | 320999 | 136287 | -1.22 | 2.2E-01 | 0 |
| Triglyceride / High-density lipoprotein cholesterol ratio (at exam 5, GEE, adjusted for age and sex | 3 | 148515 | 440528 | 102186 | -1.22 | 2.2E-01 | 0 |
| Number of older siblings | 1 | 48252 | 48252 | 48252 | -1.22 | 2.2E-01 | 0 |
| F5 - Coagulation Factor V | 1 | 42326 | 42326 | 42326 | -1.22 | 2.2E-01 | 0 |
| Alcohol dependency (unrelated individuals) | 1 | 33942 | 33942 | 33942 | -1.22 | 2.2E-01 | 0 |
| Total cholesterol (at exam 5, GEE, adjusted for age and sex) | 1 | 39731 | 39731 | 39731 | -1.22 | 2.2E-01 | 0 |
| Reason for glasses/contact lenses: For 'astigmatism' | 1 | 37587 | 37587 | 37587 | -1.22 | 2.2E-01 | 0 |
| Mean MMSE score for exam 5 and 7 (GEE, adjusted for age) | 5 | 213150 | 393503 | 89686 | -1.22 | 2.2E-01 | 0 |
| ADAMTS5 - A disintegrin and metalloproteinase with thrombospondin motifs 5 | 1 | 42435 | 42435 | 42435 | -1.22 | 2.2E-01 | 0 |
| TDGF1 - Teratocarcinoma-derived growth factor 1 | 1 | 46758 | 46758 | 46758 | -1.22 | 2.2E-01 | 0 |
| Splenium of corpus callosum mode of anisotropy | 1 | 35532 | 35532 | 35532 | -1.22 | 2.2E-01 | 0 |
| C2 - Complement C2 | 1 | 49177 | 49177 | 49177 | -1.22 | 2.2E-01 | 0 |
| Job involves shift work | 3 | 150650 | 995752 | 108329 | -1.21 | 2.2E-01 | 0 |
| High-density lipoprotein 3 cholesterol (at exam 5, GEE, adjusted for age and sex) | 1 | 41063 | 41063 | 41063 | -1.21 | 2.2E-01 | 0 |
| Frequency of light DIY in last 4 weeks | 1 | 41613 | 41613 | 41613 | -1.21 | 2.2E-01 | 0 |
| Illnesses of mother: Chronic bronchitis/emphysema | 1 | 49775 | 49775 | 49775 | -1.21 | 2.3E-01 | 0 |
| Follicle stimulating hormone (at exam 3, GEE, adjusted for multivariable) | 6 | 274744 | 876374 | 106641 | -1.21 | 2.3E-01 | 0 |
| Ratio of visceral-tosubcutaneous adipose tissue volume (adjusted for BMI) | 1 | 43488 | 43488 | 43488 | -1.21 | 2.3E-01 | 0 |
| Boston Naming Test score without cues (GEE, adjusted for multivariable) | 4 | 250470 | 460136 | 206938 | -1.21 | 2.3E-01 | 0 |
| Illnesses of father: Stroke | 1 | 40700 | 40700 | 40700 | -1.21 | 2.3E-01 | 0 |
| Duration walking for pleasure | 1 | 48891 | 48891 | 48891 | -1.21 | 2.3E-01 | 0 |
| Fornix (column and body of fornix) axial diusivities | 4 | 247492 | 396774 | 145666 | -1.21 | 2.3E-01 | 0 |
| TXNDC12 - Thioredoxin domain-containing protein 12 | 1 | 41374 | 41374 | 41374 | -1.21 | 2.3E-01 | 0 |
| CTSS - Cathepsin S | 1 | 47665 | 47665 | 47665 | -1.21 | 2.3E-01 | 0 |
| Meconium ileus in cystic fibrosis | 1 | 40478 | 40478 | 40478 | -1.21 | 2.3E-01 | 0 |
| HOMA-IR (at exam 5, GEE, adjusted for age, sex and BMI) | 2 | 232251 | 332114 | 132388 | -1.21 | 2.3E-01 | 0 |
| IL11RA - Interleukin-11 receptor subunit alpha | 1 | 47165 | 47165 | 47165 | -1.21 | 2.3E-01 | 0 |
| Diagnoses - main ICD10: K21 Gastro-esophageal reflux disease | 1 | 43810 | 43810 | 43810 | -1.20 | 2.3E-01 | 0 |
| CA13 - Carbonic anhydrase 13 | 1 | 52683 | 52683 | 52683 | -1.20 | 2.3E-01 | 0 |
| Amino acid::Lysine metabolism::pipecolate | 1 | 41635 | 41635 | 41635 | -1.20 | 2.3E-01 | 0 |
| MMP7 - Matrilysin | 1 | 41413 | 41413 | 41413 | -1.20 | 2.3E-01 | 0 |
| ::::X-11905 | 1 | 50626 | 50626 | 50626 | -1.20 | 2.3E-01 | 0 |
| Bilirubin (at exam 2, GEE, adjusted for age and sex) | 1 | 35932 | 35932 | 35932 | -1.20 | 2.3E-01 | 0 |
| ::::X-11529 | 1 | 50626 | 50626 | 50626 | -1.20 | 2.3E-01 | 0 |
| High-density lipoprotein cholesterol (at exam 5, GEE, adjusted for age and sex) | 1 | 41063 | 41063 | 41063 | -1.20 | 2.3E-01 | 0 |

| | | | | | | | |
|---|---|---|---|---|---|---|---|
| Lipid::Fatty acid, dicarboxylate::octadecanedioate | 1 | 49898 | 49898 | 49898 | -1.20 | 2.3E-01 | 0 |
| TFPI - Tissue factor pathway inhibitor | 1 | 49125 | 49125 | 49125 | -1.20 | 2.3E-01 | 0 |
| Traumatic events - Able to pay rent/morgage as an adult | 1 | 41906 | 41906 | 41906 | -1.20 | 2.3E-01 | 0 |
| Total cholesterol / High-density lipoprotein cholesterol ratio (at exam 4, GEE, adjusted for age an | 1 | 50557 | 50557 | 50557 | -1.20 | 2.3E-01 | 0 |
| MAP2K4 - Dual specificity mitogen-activated protein kinase kinase 4 | 2 | 233334 | 266697 | 199971 | -1.20 | 2.3E-01 | 0 |
| Fasting HbA1c (at exam 7, GEE, adjusted for age, sex and BMI) | 1 | 52810 | 52810 | 52810 | -1.19 | 2.3E-01 | 0 |
| BST1 - ADP-ribosyl cyclase/cyclic ADP-ribose hydrolase 2 | 1 | 56841 | 56841 | 56841 | -1.19 | 2.3E-01 | 0 |
| Chest pain or discomfort walking normally | 1 | 51049 | 51049 | 51049 | -1.19 | 2.3E-01 | 0 |
| MMP12 - Macrophage metalloelastase | 1 | 53852 | 53852 | 53852 | -1.19 | 2.3E-01 | 0 |
| Lean body mass (appendicular) | 1 | 50637 | 50637 | 50637 | -1.19 | 2.3E-01 | 0 |
| Smoking status: Previous vs Current | 1 | 56814 | 56814 | 56814 | -1.19 | 2.3E-01 | 0 |
| Right medial orbitofrontal | 1 | 55574 | 55574 | 55574 | -1.19 | 2.3E-01 | 0 |
| ANXA2 - Annexin A2 | 1 | 56800 | 56800 | 56800 | -1.19 | 2.4E-01 | 0 |
| Fibrinogen | 1 | 57474 | 57474 | 57474 | -1.18 | 2.4E-01 | 0 |
| Mean Amygdala | 1 | 50604 | 50604 | 50604 | -1.18 | 2.4E-01 | 0 |
| Macrophage inflammatory protein-1 (CCL4) | 2 | 249325 | 361553 | 137098 | -1.18 | 2.4E-01 | 0 |
| Granulocyte percentage of myeloid white cells (three-way meta) | 118 | 566890 | 1062923 | 61371 | -1.18 | 2.4E-01 | 0 |
| Superior corona radiata axial diusivities | 4 | 257590 | 583121 | 68972 | -1.18 | 2.4E-01 | 0 |
| Number of pregnancy terminations (female) | 9 | 296375 | 968948 | 83377 | -1.18 | 2.4E-01 | 0 |
| ::::X-12749 | 1 | 49556 | 49556 | 49556 | -1.18 | 2.4E-01 | 0 |
| Amino acid::Urea cycle; arginine-, proline-, metabolism::homocitrulline | 1 | 72770 | 72770 | 72770 | -1.18 | 2.4E-01 | 0 |
| Own or rent accommodation lived in: Own outright (by you or someone in your household) | 1 | 55775 | 55775 | 55775 | -1.18 | 2.4E-01 | 0 |
| CD200R1 - Cell surface glycoprotein CD200 receptor 1 | 1 | 62122 | 62122 | 62122 | -1.18 | 2.4E-01 | 0 |
| External capsule mode of anisotropy | 1 | 54619 | 54618 | 54618 | -1.18 | 2.4E-01 | 0 |
| Cardiovascular disease (MI, CI, CHD death or ABI, GEE, adjusted for age and sex) | 1 | 66703 | 66703 | 66703 | -1.18 | 2.4E-01 | 0 |
| SERPING1 - Plasma protease C1 inhibitor | 1 | 64831 | 64831 | 64831 | -1.18 | 2.4E-01 | 0 |
| Bipolar and major depression status | 1 | 66277 | 66277 | 66277 | -1.18 | 2.4E-01 | 0 |
| Amino acid::Glutamate metabolism::glutamine | 1 | 66319 | 66319 | 66319 | -1.18 | 2.4E-01 | 0 |
| Subcutaneous adipose tissue volume (female) | 1 | 58604 | 58604 | 58604 | -1.17 | 2.4E-01 | 0 |
| Number of cigarettes smoked per day | 1 | 53928 | 53928 | 53928 | -1.17 | 2.4E-01 | 0 |
| ::::X-13658 | 1 | 56255 | 56254 | 56254 | -1.17 | 2.4E-01 | 0 |
| CTSB - Cathepsin B | 1 | 64966 | 64966 | 64966 | -1.17 | 2.4E-01 | 0 |
| IL23R - Interleukin-23 receptor | 1 | 71257 | 71257 | 71257 | -1.17 | 2.4E-01 | 0 |
| ::::X-11858 | 1 | 53958 | 53958 | 53958 | -1.17 | 2.4E-01 | 0 |
| Cardiovascular disease (MI, CI, CHD death or ABI, GEE, adjusted for multivariable) | 1 | 66703 | 66703 | 66703 | -1.17 | 2.4E-01 | 0 |
| CFB - Complement factor B | 1 | 62138 | 62138 | 62138 | -1.17 | 2.4E-01 | 0 |
| Years since last breast cancer screening / mammogram (female) | 1 | 74025 | 74025 | 74025 | -1.17 | 2.4E-01 | 0 |
| IGF1 - Insulin-like growth factor I | 1 | 73830 | 73830 | 73830 | -1.17 | 2.4E-01 | 0 |
| Oily fish intake | 52 | 517029 | 1102269 | 70475 | -1.17 | 2.4E-01 | 0 |
| Frequency of depressed mood in last 2 weeks | 8 | 311489 | 801214 | 59210 | -1.17 | 2.4E-01 | 0 |
| Phltrium width | 1 | 69269 | 69269 | 69269 | -1.17 | 2.4E-01 | 0 |
| Total cholesterol (at exam 3, GEE, adjusted for age and sex) | 1 | 77581 | 77581 | 77581 | -1.17 | 2.4E-01 | 0 |
| C4A C4B - Complement C4b | 1 | 62587 | 62587 | 62587 | -1.17 | 2.4E-01 | 0 |

| | | | | | | | |
|---|---|---|---|---|---|---|---|
| IL6R - Interleukin-6 receptor subunit alpha | 1 | 63105 | 63104 | 63104 | -1.17 | 2.4E-01 | 0 |
| Factor VII (at exam 5, GEE, adjusted for multivariable) | 1 | 69430 | 69430 | 69430 | -1.17 | 2.4E-01 | 0 |
| Severe gingival inflammation | 1 | 62660 | 62660 | 62660 | -1.16 | 2.4E-01 | 0 |
| Cereal type: Biscuit cereal (e.g. Weetabix) | 1 | 63526 | 63526 | 63526 | -1.16 | 2.4E-01 | 0 |
| Amino acid::Alanine and aspartate metabolism::N-acetylalanine | 1 | 62631 | 62630 | 62630 | -1.16 | 2.4E-01 | 0 |
| Interleukin 6 (at exam 7, GEE, adjusted for multivariable) | 3 | 174616 | 459431 | 114955 | -1.16 | 2.5E-01 | 0 |
| Factor VII (at exam 5, GEE, adjusted for age and sex) | 1 | 69430 | 69430 | 69430 | -1.16 | 2.5E-01 | 0 |
| Diagnoses - secondary ICD10: R10 Abdominal and pelvic pain | 1 | 68013 | 68013 | 68013 | -1.16 | 2.5E-01 | 0 |
| Anterior limb of internal capsule mean diusivities | 4 | 270164 | 623454 | 64008 | -1.16 | 2.5E-01 | 0 |
| FTH1 FTL - Ferritin | 1 | 69148 | 69148 | 69148 | -1.16 | 2.5E-01 | 0 |
| Waist circumference (female) | 30 | 461045 | 957167 | 224040 | -1.16 | 2.5E-01 | 0 |
| Nonischemic cardiomyopathy | 1 | 76263 | 76263 | 76263 | -1.16 | 2.5E-01 | 0 |
| Age at death (GEE, adjusted for birth-cohort) | 13 | 357680 | 964500 | 127921 | -1.16 | 2.5E-01 | 0 |
| Residual from predicted FEV1 for latest exam (GEE, adjusted for multiple covariates) | 1 | 66461 | 66461 | 66461 | -1.15 | 2.5E-01 | 0 |
| ::::X-14473 | 1 | 82250 | 82250 | 82250 | -1.15 | 2.5E-01 | 0 |
| Mean internal carotid artery IMT (at exam 6, GEE, adjusted for age and sex) | 1 | 78194 | 78194 | 78194 | -1.15 | 2.5E-01 | 0 |
| ::::X-13741 | 1 | 72422 | 72422 | 72422 | -1.15 | 2.5E-01 | 0 |
| Mean high-density lipoprotein cholesterol from exam 1-7 (GEE, adjusted for multivariable) | 2 | 251804 | 300259 | 203348 | -1.15 | 2.5E-01 | 0 |
| Nucleotide::Purine metabolism, (hypo)xanthine, inosine containing::inosine | 1 | 71707 | 71707 | 71707 | -1.15 | 2.5E-01 | 0 |
| ENTPD5 - Ectonucleoside triphosphate diphosphohydrolase 5 | 1 | 76428 | 76428 | 76428 | -1.15 | 2.5E-01 | 0 |
| IGFBP3 - Insulin-like growth factor-binding protein 3 | 1 | 73830 | 73830 | 73830 | -1.15 | 2.5E-01 | 0 |
| Time spent driving | 2 | 256746 | 372968 | 140524 | -1.15 | 2.5E-01 | 0 |
| Triglyceride / High-density lipoprotein cholesterol ratio (at exam 3, GEE, adjusted for age and sex | 1 | 78118 | 78118 | 78118 | -1.14 | 2.5E-01 | 0 |
| Cofactors and vitamins::Pantothenate and CoA metabolism::pantothenate | 1 | 72590 | 72590 | 72590 | -1.14 | 2.5E-01 | 0 |
| Mean total cholesterol from exam 1-7 (GEE, adjusted for age and sex) | 1 | 75871 | 75871 | 75871 | -1.14 | 2.5E-01 | 0 |
| Alcohol dependency (random effect model for unrelated genotyped individuals) | 1 | 81986 | 81986 | 81986 | -1.14 | 2.5E-01 | 0 |
| Alcohol dependency (fixed effect model for unrelated genotyped individuals) | 1 | 81986 | 81986 | 81986 | -1.14 | 2.5E-01 | 0 |
| ::::X-12855 | 1 | 85556 | 85556 | 85556 | -1.14 | 2.5E-01 | 0 |
| MIA - Melanoma-derived growth regulatory protein | 1 | 71989 | 71989 | 71989 | -1.14 | 2.6E-01 | 0 |
| SIRT2 - NAD-dependent protein deacetylase sirtuin-2 | 1 | 83338 | 83338 | 83338 | -1.14 | 2.6E-01 | 0 |
| Vitamin K plasma phylloquinone (at exam 6 or 7, GEE, adjusted for multivariable) | 1 | 78622 | 78622 | 78622 | -1.13 | 2.6E-01 | 0 |
| NCR3 - Natural cytotoxicity triggering receptor 3 | 1 | 90180 | 90180 | 90180 | -1.13 | 2.6E-01 | 0 |
| Conscientiousness (NEO-FFI) | 1 | 75362 | 75362 | 75362 | -1.13 | 2.6E-01 | 0 |
| DKK4 - Dickkopf-related protein 4 | 1 | 83338 | 83338 | 83338 | -1.13 | 2.6E-01 | 0 |
| Own or rent accommodation lived in: Rent - from private landlord or letting agency | 1 | 87761 | 87761 | 87761 | -1.13 | 2.6E-01 | 0 |
| Body of corpus callosum mode of anisotropy | 2 | 261746 | 368255 | 155236 | -1.13 | 2.6E-01 | 0 |
| Right transverse temporal | 1 | 87790 | 87790 | 87790 | -1.13 | 2.6E-01 | 0 |
| Maximum internal carotid artery IMT (at exam 6, GEE, adjusted for age and sex) | 1 | 78194 | 78194 | 78194 | -1.13 | 2.6E-01 | 0 |
| THBS4 - Thrombospondin-4 | 1 | 98210 | 98210 | 98210 | -1.13 | 2.6E-01 | 0 |
| Frequency of consuming six or more units of alcohol | 6 | 306314 | 768224 | 34029 | -1.13 | 2.6E-01 | 0 |
| Education - Qualifications | 126 | 574844 | 1087806 | 70971 | -1.13 | 2.6E-01 | 0 |
| Diagnoses - main ICD10: M16 Osteoarthritis of hip | 10 | 343796 | 693370 | 106550 | -1.13 | 2.6E-01 | 0 |
| Lipid::Fatty acid, dicarboxylate::3-carboxy-4-methyl-5-propyl-2-furanpropanoate (CMPF) | 1 | 83759 | 83759 | 83759 | -1.12 | 2.6E-01 | 0 |

| | | | | | | | |
|---|---|---|---|---|---|---|---|
| Xenobiotics::Sugar, sugar substitute, starch::erythritol | 1 | 91344 | 91344 | 91344 | -1.12 | 2.6E-01 | 0 |
| KNG1 - Kininogen-1 | 1 | 76945 | 76945 | 76945 | -1.12 | 2.6E-01 | 0 |
| High-density lipoprotein 2 cholesterol (at exam 5, GEE, adjusted for age and sex) | 1 | 83680 | 83680 | 83680 | -1.12 | 2.6E-01 | 0 |
| Urinary albumin excertion in hypertensive enriched sample (at exam 6, GEE, adjusted for age an | 1 | 96326 | 96326 | 96326 | -1.11 | 2.7E-01 | 0 |
| Urinary albumin excertion in hypertensive enriched sample (at exam 6, GEE, adjusted for multiva | 1 | 96326 | 96326 | 96326 | -1.11 | 2.7E-01 | 0 |
| Mean common carotid artery IMT (at exam 6, GEE, adjusted for multivariable) | 1 | 92642 | 92642 | 92642 | -1.11 | 2.7E-01 | 0 |
| MICB - MHC class I polypeptide-related sequence B | 1 | 94062 | 94062 | 94062 | -1.11 | 2.7E-01 | 0 |
| External capsule mean diusivities | 2 | 270164 | 368162 | 172166 | -1.11 | 2.7E-01 | 0 |
| tPA (at exam 5, GEE, adjusted for multivariable) | 1 | 95528 | 95528 | 95528 | -1.11 | 2.7E-01 | 0 |
| Right cuneus | 5 | 257275 | 543158 | 151319 | -1.11 | 2.7E-01 | 0 |
| Insulin sensitivity (GEE, adjusted for age, sex and BMI) | 1 | 92573 | 92573 | 92573 | -1.11 | 2.7E-01 | 0 |
| Lipid::Bile acid metabolism::glycochenodeoxycholate | 1 | 97843 | 97843 | 97843 | -1.11 | 2.7E-01 | 0 |
| Illnesses of siblings: Severe depression | 1 | 109402 | 109402 | 109402 | -1.10 | 2.7E-01 | 0 |
| Splenium of corpus callosum axial diusivities | 2 | 276832 | 381652 | 172011 | -1.10 | 2.7E-01 | 0 |
| Gamma-glutamyl transferase (at exam 2, GEE, adjusted for age and sex) | 1 | 104295 | 104295 | 104295 | -1.10 | 2.7E-01 | 0 |
| Primary angle closure glaucoma | 8 | 336599 | 570258 | 53689 | -1.10 | 2.7E-01 | 0 |
| 1p deletion neuroblastoma | 3 | 200598 | 241084 | 121469 | -1.10 | 2.7E-01 | 0 |
| SIGLEC14 - Sialic acid-binding Ig-like lectin 14 | 1 | 100117 | 100117 | 100117 | -1.10 | 2.7E-01 | 0 |
| 11q deletion neuroblastoma | 3 | 200598 | 395628 | 170245 | -1.10 | 2.7E-01 | 0 |
| Ratio of bisLallylic bonds to double bonds in lipids | 2 | 279550 | 406535 | 152566 | -1.10 | 2.7E-01 | 0 |
| Neck cross-sectional moment of inertia (GEE, female, adjussted for age) | 1 | 108175 | 108174 | 108174 | -1.09 | 2.7E-01 | 0 |
| SERPINA3 - alpha-1-antichymotrypsin complex | 1 | 105475 | 105475 | 105475 | -1.09 | 2.7E-01 | 0 |
| Heart failure | 1 | 102578 | 102578 | 102578 | -1.09 | 2.8E-01 | 0 |
| Depression - Trouble falling or staying asleep. or sleeping too much | 1 | 96727 | 96727 | 96727 | -1.09 | 2.8E-01 | 0 |
| LGALS3 - Galectin-3 | 1 | 99542 | 99542 | 99542 | -1.09 | 2.8E-01 | 0 |
| SERPINA3 - Alpha-1-antichymotrypsin | 1 | 105475 | 105475 | 105475 | -1.08 | 2.8E-01 | 0 |
| Bread type: Wholemeal or wholegrain | 14 | 393296 | 1151058 | 77991 | -1.08 | 2.8E-01 | 0 |
| DKK3 - Dickkopf-related protein 3 | 1 | 105130 | 105130 | 105130 | -1.08 | 2.8E-01 | 0 |
| Posterior limb of internal capsule fractional anisotropy | 8 | 343513 | 806896 | 78111 | -1.08 | 2.8E-01 | 0 |
| Eye problems/disorders: Cataract | 2 | 277362 | 357695 | 197029 | -1.08 | 2.8E-01 | 0 |
| Trunk-trunk fat ratio (female) | 184 | 596538 | 1172396 | 86587 | -1.08 | 2.8E-01 | 0 |
| ACP1 - Low molecular weight phosphotyrosine protein phosphatase | 1 | 112461 | 112461 | 112461 | -1.07 | 2.8E-01 | 0 |
| Psoriasis | 5 | 269828 | 367203 | 157010 | -1.07 | 2.8E-01 | 0 |
| Reticulocyte count (three-way meta) | 109 | 570260 | 1100540 | 106267 | -1.07 | 2.8E-01 | 0 |
| CASP3 - Caspase-3 | 1 | 121828 | 121828 | 121828 | -1.07 | 2.9E-01 | 0 |
| GPNMB - Transmembrane glycoprotein NMB | 1 | 127137 | 127137 | 127137 | -1.07 | 2.9E-01 | 0 |
| CD123 on 11c+123+DC | 6 | 319690 | 510082 | 142118 | -1.06 | 2.9E-01 | 0 |
| Amino acid::Polyamine metabolism::X-03056--N-[3-(2-Oxopyrrolidin-1-yl)propyl]acetamide | 3 | 209582 | 343057 | 128382 | -1.06 | 2.9E-01 | 0 |
| Urinary albumin excertion of at least 30 in enriched hypertensive sample (at exam 6, GEE, adjust | 128 | 580163 | 962323 | 252430 | -1.06 | 2.9E-01 | 0 |
| Subjective well being | 1 | 117951 | 117951 | 117951 | -1.06 | 2.9E-01 | 0 |
| NOV - Protein NOV homolog | 1 | 129775 | 129775 | 129775 | -1.06 | 2.9E-01 | 0 |
| Disposition index (adjusted for BMI) | 1 | 134748 | 134748 | 134748 | -1.06 | 2.9E-01 | 0 |
| Gout | 2 | 288515 | 415423 | 161608 | -1.05 | 2.9E-01 | 0 |

| | | | | | | | |
|---|---|---|---|---|---|---|---|
| Shaft average buckling ratio (GEE, male, adjussted for multivariable) | 1 | 128658 | 128658 | 128658 | -1.05 | 2.9E-01 | 0 |
| Neck-Shaft Angle (GEE, adjusted for multivariable) | 1 | 130815 | 130815 | 130815 | -1.05 | 2.9E-01 | 0 |
| Left isthmus cingulate | 4 | 306495 | 721156 | 49208 | -1.05 | 3.0E-01 | 0 |
| Left vessel | 3 | 224226 | 605667 | 170768 | -1.05 | 3.0E-01 | 0 |
| ::::X-13069 | 1 | 123964 | 123964 | 123964 | -1.04 | 3.0E-01 | 0 |
| Diagnoses - secondary ICD10: E66 Overweight and obesity | 3 | 227601 | 386760 | 139639 | -1.04 | 3.0E-01 | 0 |
| Disposition index | 1 | 134748 | 134748 | 134748 | -1.04 | 3.0E-01 | 0 |
| Neck-Shaft Angle (GEE, adjusted for age and sex) | 1 | 130815 | 130815 | 130815 | -1.04 | 3.0E-01 | 0 |
| Alanine transaminase (at exam 2, GEE, adjusted for age and sex) | 1 | 133607 | 133607 | 133607 | -1.04 | 3.0E-01 | 0 |
| 3Lhydroxybutyrate | 1 | 127709 | 127709 | 127709 | -1.03 | 3.0E-01 | 0 |
| Diagnoses - secondary ICD10: Z86 Personal history of certain other diseases | 3 | 225979 | 1377815 | 131869 | -1.03 | 3.0E-01 | 0 |
| Triglyceride / High-density lipoprotein cholesterol ratio (at exam 4, GEE, adjusted for age and sex | 1 | 139078 | 139078 | 139078 | -1.03 | 3.1E-01 | 0 |
| AFM - Afamin | 1 | 146106 | 146106 | 146106 | -1.02 | 3.1E-01 | 0 |
| Diagnoses - secondary ICD10: F32 Major depressive disorder, single episode | 1 | 141002 | 141002 | 141002 | -1.02 | 3.1E-01 | 0 |
| Neck cross-sectional moment of inertia (GEE, male, adjussted for multivariable) | 1 | 135979 | 135979 | 135979 | -1.02 | 3.1E-01 | 0 |
| Right parahippocampal | 1 | 140254 | 140254 | 140254 | -1.02 | 3.1E-01 | 0 |
| Major depressive disorder (females) | 2 | 306760 | 420480 | 193040 | -1.01 | 3.1E-01 | 0 |
| Parkinson disease | 5 | 284488 | 1019892 | 115794 | -1.01 | 3.1E-01 | 0 |
| Coronary artery disease (SOFT definition including angina) | 19 | 437615 | 643464 | 204980 | -1.01 | 3.1E-01 | 0 |
| Pain type(s) experienced in last month: Hip pain | 1 | 144252 | 144252 | 144252 | -1.00 | 3.2E-01 | 0 |
| Energy::Oxidative phosphorylation::acetylphosphate | 2 | 305180 | 406679 | 203682 | -1.00 | 3.2E-01 | 0 |
| High-density lipoprotein particle size by NMR (at exam 4, GEE, adjusted for age and sex) | 1 | 148515 | 148515 | 148515 | -1.00 | 3.2E-01 | 0 |
| Extreme Body Mass Index | 6 | 339575 | 621187 | 72550 | -1.00 | 3.2E-01 | 0 |
| Mean thyroid stimulation hormone from exam 3 and 4 (GEE, adjusted for age and sex) | 3 | 243416 | 759696 | 173253 | -0.99 | 3.2E-01 | 0 |
| Glucocorticoids | 12 | 406321 | 922544 | 156640 | -0.99 | 3.2E-01 | 0 |
| Blond hair in Solomon Islanders | 1 | 150429 | 150429 | 150429 | -0.99 | 3.2E-01 | 0 |
| Bilateral oophorectomy (both ovaries removed) (female) | 6 | 351934 | 726293 | 99518 | -0.99 | 3.2E-01 | 0 |
| Primary sclerosing cholangitis | 1 | 160339 | 160339 | 160339 | -0.99 | 3.2E-01 | 0 |
| CGA FSHB - Follicle stimulating hormone | 1 | 156653 | 156653 | 156653 | -0.99 | 3.2E-01 | 0 |
| Ankle-brachial index (at exam 6, GEE, adjusted for age and sex) | 2 | 309009 | 398106 | 219912 | -0.99 | 3.2E-01 | 0 |
| Amino acid::Creatine metabolism::creatinine | 1 | 168503 | 168503 | 168503 | -0.99 | 3.2E-01 | 0 |
| Prospective memory test - Time to answer | 1 | 156966 | 156966 | 156966 | -0.98 | 3.3E-01 | 0 |
| Xenobiotics::Xanthine metabolism::7-methylxanthine | 1 | 169008 | 169008 | 169008 | -0.98 | 3.3E-01 | 0 |
| Percent predicted FEV1/FVC for latest exam (GEE, adjusted for multiple covariates) | 2 | 313860 | 444314 | 183406 | -0.98 | 3.3E-01 | 0 |
| Pericardial adipose tissue volume (adjusted for height and weight) | 3 | 249751 | 604247 | 153668 | -0.98 | 3.3E-01 | 0 |
| Anxiety - Recent worrying too much about different things | 1 | 165224 | 165224 | 165224 | -0.98 | 3.3E-01 | 0 |
| CD300C - CMRF35-like molecule 6 | 1 | 169106 | 169106 | 169106 | -0.98 | 3.3E-01 | 0 |
| PLA2G2A - Phospholipase A2, membrane associated | 1 | 162662 | 162662 | 162662 | -0.97 | 3.3E-01 | 0 |
| Residual from predicted FEV1/FVC for latest exam (GEE, adjusted for multiple covariates) | 2 | 313860 | 444314 | 183406 | -0.97 | 3.3E-01 | 0 |
| Estimated glomerular filtration rate based on cystain C | 21 | 461418 | 852025 | 75316 | -0.97 | 3.3E-01 | 0 |
| Coronary artery disease | 59 | 547253 | 942629 | 149970 | -0.97 | 3.3E-01 | 0 |
| Low-density lipoprotein cholesterol (at exam 7, GEE, adjusted for age and sex) | 3 | 258490 | 375309 | 198934 | -0.97 | 3.3E-01 | 0 |
| ::::X-13496 | 2 | 328038 | 463909 | 192168 | -0.97 | 3.3E-01 | 0 |

| | | | | | | | |
|---|---|---|---|---|---|---|---|
| LEPR - Leptin receptor | 1 | 171856 | 171856 | 171856 | -0.97 | 3.3E-01 | 0 |
| CPB2 - Carboxypeptidase B2 | 1 | 180436 | 180436 | 180436 | -0.96 | 3.4E-01 | 0 |
| Skin colour | 141 | 592280 | 1057190 | 80257 | -0.96 | 3.4E-01 | 0 |
| Small high-density lipoprotein by NMR (at exam 4, GEE, adjusted for age and sex) | 5 | 312530 | 581883 | 118377 | -0.96 | 3.4E-01 | 0 |
| Monocyte percentage of white cells (two-way meta) | 93 | 576443 | 1006610 | 51702 | -0.96 | 3.4E-01 | 0 |
| Low-density lipoprotein cholesterol (at exam 1, GEE, adjusted for multivariable) | 2 | 319872 | 454121 | 185623 | -0.95 | 3.4E-01 | 0 |
| Percent predicted FEV1 for latest exam (GEE, adjusted for multiple covariates) | 2 | 336659 | 470782 | 202535 | -0.95 | 3.4E-01 | 0 |
| Right lingual | 2 | 328520 | 445639 | 211401 | -0.95 | 3.4E-01 | 0 |
| Low-density lipoprotein cholesterol (at exam 3, GEE, adjusted for age and sex) | 2 | 332975 | 460673 | 205278 | -0.95 | 3.4E-01 | 0 |
| Miserableness (MIS) | 32 | 508028 | 1096294 | 83254 | -0.94 | 3.5E-01 | 0 |
| Non-butter spread type details: Soft (tub) margarine | 1 | 182155 | 182155 | 182155 | -0.94 | 3.5E-01 | 0 |
| NKearly:%337+158b+ | 1 | 178749 | 178749 | 178749 | -0.94 | 3.5E-01 | 0 |
| Urinary albumin excertion (at exam 6, GEE, adjusted for multivariable) | 1 | 186388 | 186388 | 186388 | -0.94 | 3.5E-01 | 0 |
| Asthma (fixed effect model) | 21 | 465193 | 1011813 | 136209 | -0.94 | 3.5E-01 | 0 |
| Social support - Leisure/social activities: Other group activity | 5 | 309555 | 984592 | 256188 | -0.93 | 3.5E-01 | 0 |
| Tobacco smoking | 4 | 338065 | 777539 | 40820 | -0.93 | 3.5E-01 | 0 |
| Types of physical activity in last 4 weeks: Other exercises (eg: swimming, cycling, keep fit, bowlir | 9 | 379430 | 638813 | 256820 | -0.92 | 3.6E-01 | 0 |
| Interleukin-13 | 1 | 200425 | 200425 | 200425 | -0.92 | 3.6E-01 | 0 |
| MAP2K2 - Dual specificity mitogen-activated protein kinase kinase 2 | 1 | 194036 | 194036 | 194036 | -0.92 | 3.6E-01 | 0 |
| Visual memory composite score (GEE, adjusted for multivariable) | 3 | 275695 | 650359 | 193286 | -0.92 | 3.6E-01 | 0 |
| Height SDS (male at age 12) | 1 | 203205 | 203205 | 203205 | -0.92 | 3.6E-01 | 0 |
| Milk type used: Skimmed | 5 | 323715 | 685363 | 67861 | -0.92 | 3.6E-01 | 0 |
| Ever had same-sex intercourse | 3 | 276390 | 337001 | 161982 | -0.92 | 3.6E-01 | 0 |
| Sodium | 11 | 407960 | 1181650 | 63995 | -0.92 | 3.6E-01 | 0 |
| Shaft average buckling ratio (GEE, female, adjussted for age) | 2 | 337619 | 467762 | 207476 | -0.92 | 3.6E-01 | 0 |
| Weight (male) | 9 | 382215 | 1187630 | 64317 | -0.91 | 3.6E-01 | 0 |
| Immunoglobulin A deficiency | 1 | 199865 | 199865 | 199865 | -0.91 | 3.6E-01 | 0 |
| MMSE score (at exam 5, GEE, adjusted for multivariable) | 3 | 274398 | 687744 | 163966 | -0.91 | 3.6E-01 | 0 |
| ::::X-11261 | 1 | 209582 | 209582 | 209582 | -0.91 | 3.6E-01 | 0 |
| Total body BMD (60 or older) | 17 | 449128 | 928078 | 87540 | -0.90 | 3.7E-01 | 0 |
| Platelets | 23 | 475608 | 950306 | 161699 | -0.90 | 3.7E-01 | 0 |
| AKR1A1 - Alcohol dehydrogenase [NADP(+)] | 1 | 202076 | 202076 | 202076 | -0.90 | 3.7E-01 | 0 |
| Lipid::Monoacylglycerol::1-linoleoylglycerol (1-monolinolein) | 1 | 201998 | 201998 | 201998 | -0.90 | 3.7E-01 | 0 |
| Albumin | 11 | 404728 | 1181053 | 85602 | -0.90 | 3.7E-01 | 0 |
| MET - Hepatocyte growth factor receptor | 2 | 348838 | 515803 | 181873 | -0.90 | 3.7E-01 | 0 |
| Mean MMSE score for exam 5 and 7 (GEE, adjusted for multivariable) | 1 | 213150 | 213150 | 213150 | -0.90 | 3.7E-01 | 0 |
| Phospholipids in medium LDL | 9 | 385130 | 818555 | 106456 | -0.90 | 3.7E-01 | 0 |
| Lipid::Lysolipid::2-stearoylglycerophosphocholine* | 1 | 213700 | 213700 | 213700 | -0.89 | 3.7E-01 | 0 |
| KDR - Vascular endothelial growth factor receptor 2 | 2 | 354096 | 518432 | 189760 | -0.89 | 3.7E-01 | 0 |
| Cisplatin-associated ototoxicity | 1 | 215406 | 215406 | 215406 | -0.89 | 3.7E-01 | 0 |
| Cystatin C (at exam 7, GEE, adjusted for age and sex) | 1 | 213150 | 213150 | 213150 | -0.89 | 3.8E-01 | 0 |
| CCL16 - C-C motif chemokine 16 | 1 | 213623 | 213623 | 213623 | -0.89 | 3.8E-01 | 0 |
| Amino acid::Valine, leucine and isoleucine metabolism::2-methylbutyroylcarnitine | 1 | 209582 | 209582 | 209582 | -0.88 | 3.8E-01 | 0 |

| | | | | | | | |
|---|---|---|---|---|---|---|---|
| RR interval (female, cohort exam 11 offspring exam 1, GEE, adjusted for age nad RR) | 2 | 344529 | 502637 | 186421 | -0.88 | 3.8E-01 | 0 |
| Dehydroepiandrosterone sulfate (at exam 3, GEE, adjusted for multivariable) | 5 | 330020 | 409593 | 45480 | -0.88 | 3.8E-01 | 0 |
| Lipid::Carnitine metabolism::laurylcarnitine | 1 | 216130 | 216130 | 216130 | -0.88 | 3.8E-01 | 0 |
| Right supramarginal | 3 | 300553 | 495426 | 162765 | -0.88 | 3.8E-01 | 0 |
| Total lipids in medium LDL | 12 | 435135 | 1050478 | 86744 | -0.87 | 3.8E-01 | 0 |
| Bisecting GlcNAc | 2 | 356250 | 447470 | 265030 | -0.87 | 3.8E-01 | 0 |
| Cystatin C (at exam 7, GEE, adjusted for multivariable) | 1 | 213150 | 213150 | 213150 | -0.87 | 3.8E-01 | 0 |
| Total cholesterol in IDL | 12 | 435135 | 1095218 | 95473 | -0.87 | 3.9E-01 | 0 |
| Left lateral orbitofrontal | 2 | 357608 | 493928 | 221288 | -0.86 | 3.9E-01 | 0 |
| Obesity class 3 | 2 | 360811 | 426552 | 295070 | -0.86 | 3.9E-01 | 0 |
| Lipid::Fatty acid, monohydroxy::2-hydroxypalmitate | 1 | 221208 | 221208 | 221208 | -0.86 | 3.9E-01 | 0 |
| Amino acid::Phenylalanine & tyrosine metabolism::X-11423--O-sulfo-L-tyrosine | 2 | 366162 | 524465 | 207859 | -0.86 | 3.9E-01 | 0 |
| ::::X-11334 | 1 | 213132 | 213132 | 213132 | -0.86 | 3.9E-01 | 0 |
| Mineral and other dietary supplements: Iron | 1 | 235206 | 235206 | 235206 | -0.86 | 3.9E-01 | 0 |
| Age at death (GEE, adjusted for multivariable) | 12 | 435141 | 746174 | 219740 | -0.86 | 3.9E-01 | 0 |
| Lipoprotein A (at exam 3, GEE, adjusted for age and sex) | 16 | 467165 | 918525 | 155511 | -0.86 | 3.9E-01 | 0 |
| Treatment/medication code: lisinopril | 3 | 298338 | 693795 | 174102 | -0.85 | 3.9E-01 | 0 |
| Celiac disease | 38 | 533051 | 1057677 | 126359 | -0.85 | 3.9E-01 | 0 |
| Vitamin D plasma 25(OH)-D (at exam 6 or 7, GEE, adjusted for multivariable) | 1 | 221840 | 221840 | 221840 | -0.85 | 3.9E-01 | 0 |
| Anterior corona radiata axial diusivities | 3 | 303995 | 416168 | 168223 | -0.85 | 3.9E-01 | 0 |
| Depression | 40 | 536490 | 958729 | 68205 | -0.85 | 4.0E-01 | 0 |
| SELP - P-Selectin | 2 | 363678 | 523223 | 204133 | -0.85 | 4.0E-01 | 0 |
| Periodontal complex trait 1 - Socransky trait | 5 | 348638 | 532088 | 57663 | -0.84 | 4.0E-01 | 0 |
| Dapsone hypersensitivity syndrome | 1 | 230508 | 230508 | 230508 | -0.84 | 4.0E-01 | 0 |
| Retrolenticular part of internal capsule radial diusivities | 6 | 388821 | 640041 | 97264 | -0.84 | 4.0E-01 | 0 |
| Neck Cross-Sectional Moment of Inertia (GEE, adjusted for age and sex) | 3 | 313900 | 675333 | 208294 | -0.84 | 4.0E-01 | 0 |
| Social support - Leisure/social activities: Religious group | 41 | 543158 | 1075395 | 84723 | -0.83 | 4.0E-01 | 0 |
| Fasting glucose main effect (adjusted for BMI) | 20 | 484658 | 1055847 | 117412 | -0.83 | 4.0E-01 | 0 |
| Triglycerides (at exam 3, GEE, adjusted for age and sex) | 1 | 237183 | 237183 | 237183 | -0.83 | 4.1E-01 | 0 |
| Time spent watching television (TV) | 94 | 588114 | 1210895 | 86453 | -0.83 | 4.1E-01 | 0 |
| HGFAC - Hepatocyte growth factor activator | 2 | 372572 | 539840 | 205304 | -0.83 | 4.1E-01 | 0 |
| Walking speed (at exam 7, GEE) | 2 | 373150 | 415107 | 331193 | -0.83 | 4.1E-01 | 0 |
| TLR4 LY96 - Toll-like receptor 4:Lymphocyte antigen 96 complex | 2 | 367728 | 525248 | 210208 | -0.82 | 4.1E-01 | 0 |
| SMPDL3A - Acid sphingomyelinase-like phosphodiesterase 3a | 1 | 254480 | 254480 | 254480 | -0.82 | 4.1E-01 | 0 |
| Tumor necrosis factor alpha (at exam 7, GEE, adjusted for age and sex) | 2 | 373314 | 541329 | 205298 | -0.82 | 4.1E-01 | 0 |
| Primary biliary cirrhosis | 17 | 471018 | 837185 | 72818 | -0.82 | 4.1E-01 | 0 |
| High-density lipoprotein cholesterol (at exam 6, GEE, adjusted for age and sex) | 1 | 261068 | 261068 | 261068 | -0.82 | 4.1E-01 | 0 |
| Frequency of other exercises in last 4 weeks | 1 | 247745 | 247745 | 247745 | -0.82 | 4.1E-01 | 0 |
| Walking speed (at offspring exam 7 cohort exam 27, GEE) | 2 | 373150 | 415107 | 331193 | -0.81 | 4.2E-01 | 0 |
| Sleep durataion (conditioning sex and BMI) | 8 | 415726 | 1113755 | 47191 | -0.81 | 4.2E-01 | 0 |
| CCL7 - C-C motif chemokine 7 | 1 | 253743 | 253743 | 253743 | -0.81 | 4.2E-01 | 0 |
| Total lipids in small HDL | 7 | 385130 | 870363 | 14263 | -0.81 | 4.2E-01 | 0 |
| IL22RA2 - Interleukin-22 receptor subunit alpha-2 | 1 | 256605 | 256605 | 256605 | -0.81 | 4.2E-01 | 0 |

| | | | | | | | |
|---|---|---|---|---|---|---|---|
| Lipid::Essential fatty acid::dihomo-linolenate (20:3n3 or n6) | 2 | 378892 | 532005 | 225779 | -0.81 | 4.2E-01 | 0 |
| CCL21 - C-C motif chemokine 21 | 1 | 261233 | 261232 | 261232 | -0.81 | 4.2E-01 | 0 |
| Albumin/globulin ratio | 36 | 538656 | 982466 | 159788 | -0.81 | 4.2E-01 | 0 |
| Bone Ultrasound Attenuation (GEE, adjusted for multivariable) | 1 | 258733 | 258732 | 258732 | -0.81 | 4.2E-01 | 0 |
| Diagnoses - secondary ICD10: I48 Atrial fibrillation and flutter | 18 | 482194 | 960957 | 85545 | -0.80 | 4.2E-01 | 0 |
| Sabuctaneous fat by CT (GEE, adjusted for age, sex, smoking and menopause) | 3 | 323475 | 725280 | 275570 | -0.79 | 4.3E-01 | 0 |
| CD27 - CD27 antigen | 1 | 254023 | 254023 | 254023 | -0.79 | 4.3E-01 | 0 |
| Similarities raw score (GEE, adjusted for multivariable) | 1 | 270908 | 270908 | 270908 | -0.79 | 4.3E-01 | 0 |
| Viscosity I (at exam 5, GEE, adjusted for age and sex) | 3 | 330808 | 821301 | 175665 | -0.79 | 4.3E-01 | 0 |
| Sabuctaneous fat by CT (GEE, adjusted for age and sex) | 3 | 323475 | 725280 | 275570 | -0.79 | 4.3E-01 | 0 |
| Educational attainment | 471 | 637770 | 1144603 | 112184 | -0.79 | 4.3E-01 | 0 |
| Mean high-density lipoprotein cholesterol from exam 1-7 (GEE, adjusted for age and sex) | 2 | 388291 | 512898 | 263684 | -0.79 | 4.3E-01 | 0 |
| Types of transport used (excluding work): Public transport | 4 | 391594 | 917194 | 79860 | -0.78 | 4.3E-01 | 0 |
| Adiponectin | 9 | 420263 | 508473 | 34787 | -0.78 | 4.3E-01 | 0 |
| Wide Range Achievement Test (GEE, adjusted for multivariable) | 1 | 262548 | 262548 | 262548 | -0.78 | 4.3E-01 | 0 |
| Ratio of visceral-tosubcutaneous adipose tissue volume (adjusted for BMI, female) | 1 | 269605 | 269605 | 269605 | -0.78 | 4.4E-01 | 0 |
| ANG - Angiogenin | 1 | 272390 | 272390 | 272390 | -0.78 | 4.4E-01 | 0 |
| Gray matter | 5 | 370350 | 921158 | 62138 | -0.77 | 4.4E-01 | 0 |
| High-density lipoprotein cholesterol (at exam 2, GEE, adjusted for age and sex) | 4 | 397353 | 671840 | 231969 | -0.77 | 4.4E-01 | 0 |
| NXPH1 - Neurexophilin-1 | 1 | 279333 | 279333 | 279333 | -0.77 | 4.4E-01 | 0 |
| FCN1 - Ficolin-1 | 1 | 278583 | 278583 | 278583 | -0.77 | 4.4E-01 | 0 |
| CCL1 - C-C motif chemokine 1 | 1 | 279333 | 279333 | 279333 | -0.76 | 4.5E-01 | 0 |
| Potassium | 7 | 412018 | 756058 | 51825 | -0.76 | 4.5E-01 | 0 |
| Femoral BMD (GEE, adjusted for multivariable) | 1 | 294198 | 294198 | 294198 | -0.76 | 4.5E-01 | 0 |
| FEV1/FCV | 6 | 415331 | 570311 | 315943 | -0.75 | 4.5E-01 | 0 |
| Reason for glasses/contact lenses: For long-sightedness, i.e. for distance and near, but particula | 4 | 401899 | 710946 | 134303 | -0.75 | 4.5E-01 | 0 |
| Basophil count | 23 | 512150 | 1190899 | 76009 | -0.75 | 4.5E-01 | 0 |
| Left caudal anterior cingulate | 4 | 402908 | 780317 | 91580 | -0.75 | 4.5E-01 | 0 |
| Troch Neck BMD (GEE, male, adjussted for multivariable) | 2 | 402444 | 584057 | 220831 | -0.75 | 4.5E-01 | 0 |
| Heart rate variability (pvRSA/HF) | 7 | 406100 | 930565 | 76558 | -0.75 | 4.5E-01 | 0 |
| Creatinine | 4 | 397978 | 715134 | 145730 | -0.75 | 4.6E-01 | 0 |
| Free cholesterol in large LDL | 10 | 453360 | 937892 | 173578 | -0.74 | 4.6E-01 | 0 |
| Fornix (column and body of fornix) radial diusivities | 4 | 403400 | 850859 | 243339 | -0.74 | 4.6E-01 | 0 |
| Depression - Recent changes in speed/amount of moving or speaking | 2 | 411778 | 589381 | 234174 | -0.74 | 4.6E-01 | 0 |
| Lipid::Medium chain fatty acid::caprate (10:0) | 2 | 412974 | 526115 | 299834 | -0.74 | 4.6E-01 | 0 |
| HFE2 - Hemojuvelin | 1 | 299470 | 299470 | 299470 | -0.74 | 4.6E-01 | 0 |
| Fasting HbA1c (at exam 7, GEE, adjusted for age and sex) | 1 | 294975 | 294975 | 294975 | -0.73 | 4.6E-01 | 0 |
| PNN50 HRV (cohort exam 18 offsprint exam 3, GEE, adjusted for age and HR) | 1 | 298460 | 298460 | 298460 | -0.73 | 4.7E-01 | 0 |
| Traumatic events - Witnessed sudden violent death | 1 | 295505 | 295505 | 295505 | -0.72 | 4.7E-01 | 0 |
| Body of corpus callosum radial diusivities | 1 | 303995 | 303995 | 303995 | -0.72 | 4.7E-01 | 0 |
| ESR1 - Estrogen receptor | 1 | 300060 | 300060 | 300060 | -0.72 | 4.7E-01 | 0 |
| Urinary albumin excertion of at least 30 in enriched hypertensive sample (at exam 6, GEE, adjust | 86 | 596151 | 921702 | 271061 | -0.72 | 4.7E-01 | 0 |
| Social support - Leisure/social activities: Adult education class | 2 | 419401 | 605997 | 232805 | -0.72 | 4.7E-01 | 0 |

| | | | | | | | |
|---|---|---|---|---|---|---|---|
| Mouth/teeth dental problems: Mouth ulcers | 27 | 528883 | 969554 | 173262 | -0.72 | 4.7E-01 | 0 |
| Glycine level | 34 | 546529 | 1088058 | 58887 | -0.72 | 4.7E-01 | 0 |
| Visuospatial memory and organization (GEE, adjusted for multivariable) | 1 | 296015 | 296015 | 296015 | -0.71 | 4.8E-01 | 0 |
| Amino acid::Valine, leucine and isoleucine metabolism::levulinate (4-oxovalerate) | 1 | 309273 | 309273 | 309273 | -0.71 | 4.8E-01 | 0 |
| Diagnoses - secondary ICD10: M19 Other and unspecified osteoarthritis | 1 | 313033 | 313033 | 313033 | -0.71 | 4.8E-01 | 0 |
| Concentration of small HDL particles | 5 | 395798 | 1344928 | 17101 | -0.71 | 4.8E-01 | 0 |
| Ulcerative Colitis | 27 | 531558 | 890296 | 164281 | -0.71 | 4.8E-01 | 0 |
| Father still alive | 3 | 361718 | 625385 | 200031 | -0.71 | 4.8E-01 | 0 |
| Water intake | 26 | 536603 | 1007956 | 81688 | -0.71 | 4.8E-01 | 0 |
| Xenobiotics::Drug::ibuprofen | 9 | 443588 | 805615 | 402078 | -0.70 | 4.8E-01 | 0 |
| Miserableness | 31 | 542315 | 958953 | 94404 | -0.70 | 4.8E-01 | 0 |
| Average across all tracts mean diusivities | 5 | 402258 | 466160 | 303995 | -0.69 | 4.9E-01 | 0 |
| CD4mem:%cFTH (1) | 1 | 321275 | 321275 | 321275 | -0.69 | 4.9E-01 | 0 |
| Medication for cholesterol, blood pressure, diabetes, or take exogenous hormones: Hormone rep | 2 | 427943 | 450684 | 405201 | -0.69 | 4.9E-01 | 0 |
| Glycine level (female) | 4 | 421852 | 962418 | 38242 | -0.69 | 4.9E-01 | 0 |
| Depression - Recent poor appetite or overeting | 1 | 323715 | 323715 | 323715 | -0.69 | 4.9E-01 | 0 |
| Illnesses of mother: Lung cancer | 2 | 420399 | 620102 | 220695 | -0.68 | 4.9E-01 | 0 |
| Myeloperoxidase (at exam 7, GEE, adjusted for multivariable) | 1 | 324093 | 324093 | 324093 | -0.68 | 4.9E-01 | 0 |
| Concentration of medium LDL particles | 13 | 485140 | 1055113 | 106456 | -0.68 | 5.0E-01 | 0 |
| Platelet aggregation to Epinephrine (at exam 5, GEE, adjusted for age and sex) | 1 | 322990 | 322990 | 322990 | -0.68 | 5.0E-01 | 0 |
| Myeloperoxidase (at exam 7, GEE, adjusted for age and sex) | 1 | 324093 | 324093 | 324093 | -0.68 | 5.0E-01 | 0 |
| Worrier / anxious feelings | 44 | 572380 | 946318 | 75698 | -0.68 | 5.0E-01 | 0 |
| Coffee type: Instant coffee | 3 | 386028 | 710076 | 275317 | -0.68 | 5.0E-01 | 0 |
| LILRB2 - Leukocyte immunoglobulin-like receptor subfamily B member 2 | 1 | 326925 | 326925 | 326925 | -0.67 | 5.0E-01 | 0 |
| ::::X-11315 | 2 | 425939 | 531712 | 320167 | -0.67 | 5.0E-01 | 0 |
| Lithium response in Bipolar I patients - Alda Scale of 4 to 5 | 1 | 335108 | 335108 | 335108 | -0.67 | 5.0E-01 | 0 |
| IL12RB2 - Interleukin-12 receptor subunit beta-2 | 1 | 334775 | 334775 | 334775 | -0.67 | 5.0E-01 | 0 |
| Cereal type: Oat cereal (e.g. Ready Brek, porridge) | 1 | 323715 | 323715 | 323715 | -0.67 | 5.0E-01 | 0 |
| Lithium response in Bipolar I patients - Alda Scale of 7 to 8 | 1 | 335108 | 335108 | 335108 | -0.67 | 5.1E-01 | 0 |
| Anxiety - Recent trouble relaxing | 2 | 441872 | 653636 | 230109 | -0.67 | 5.1E-01 | 0 |
| Lithium response in Bipolar I patients - Alda Scale of 5 to 6 | 1 | 335108 | 335108 | 335108 | -0.67 | 5.1E-01 | 0 |
| Lithium response in Bipolar I patients - Alda Scale of 6 to 7 | 1 | 335108 | 335108 | 335108 | -0.66 | 5.1E-01 | 0 |
| CCL14 - C-C motif chemokine 14 | 1 | 334625 | 334625 | 334625 | -0.66 | 5.1E-01 | 0 |
| CH2 groups in fatty acids | 3 | 385130 | 439593 | 202385 | -0.66 | 5.1E-01 | 0 |
| Insulin sensitivity index (combined influence of the genotype effect adjusted for BMI and the inter | 11 | 472168 | 593669 | 31838 | -0.66 | 5.1E-01 | 0 |
| Left pericalcarine | 5 | 414180 | 721700 | 84806 | -0.66 | 5.1E-01 | 0 |
| Intercellular adhesion molecule-1 (at exam 7, GEE, adjusted for age and sex) | 1 | 340870 | 340870 | 340870 | -0.66 | 5.1E-01 | 0 |
| KLKB1 - Plasma kallikrein | 2 | 444254 | 582827 | 305680 | -0.65 | 5.1E-01 | 0 |
| Left transverse temporal | 1 | 338685 | 338685 | 338685 | -0.65 | 5.1E-01 | 0 |
| CD4:CD8 lymphocyte ratio | 1 | 343630 | 343630 | 343630 | -0.65 | 5.1E-01 | 0 |
| KLK12 - Kallikrein-12 | 1 | 336030 | 336030 | 336030 | -0.65 | 5.1E-01 | 0 |
| Milk type used: Semi-skimmed | 2 | 438239 | 623706 | 252773 | -0.65 | 5.2E-01 | 0 |
| CCL23 - C-C motif chemokine 23 | 1 | 334625 | 334625 | 334625 | -0.64 | 5.2E-01 | 0 |

| | | | | | | | |
|---|---|---|---|---|---|---|---|
| Epithelial ovarian cancer (high -grade and low-grade serous) | 1 | 349565 | 349565 | 349565 | -0.64 | 5.2E-01 | 0 |
| Lipid::Lysolipid::1-arachidonoylglycerophosphoethanolamine* | 3 | 395798 | 858888 | 223212 | -0.64 | 5.2E-01 | 0 |
| White blood cell count | 30 | 556364 | 1138005 | 92236 | -0.64 | 5.2E-01 | 0 |
| P-selectin (at exam 7, GEE, adjusted for age and sex) | 2 | 439788 | 440254 | 439321 | -0.64 | 5.2E-01 | 0 |
| Amino acid::Valine, leucine and isoleucine metabolism::beta-hydroxyisovalerate | 1 | 350585 | 350585 | 350585 | -0.64 | 5.2E-01 | 0 |
| Fasting HbA1c (at exam 5, GEE, adjusted for age, sex and BMI) | 2 | 439373 | 610280 | 268466 | -0.64 | 5.2E-01 | 0 |
| Right paracentral | 1 | 345645 | 345645 | 345645 | -0.64 | 5.2E-01 | 0 |
| CD40 Ligand serum (at exam 7, GEE, adjusted for multivariable) | 2 | 447752 | 637900 | 257604 | -0.64 | 5.2E-01 | 0 |
| Age at natural menopause (GEE, adjusted for multivariable) | 2 | 443058 | 634202 | 251915 | -0.64 | 5.2E-01 | 0 |
| Lipid::Sterol, Steroid::epiandrosterone sulfate | 1 | 355103 | 355103 | 355103 | -0.64 | 5.2E-01 | 0 |
| Worrier / anxious feelings (WORRY) | 35 | 562025 | 903823 | 74896 | -0.64 | 5.2E-01 | 0 |
| Intercellular adhesion molecule-1 (at exam 7, GEE, adjusted for multivariable) | 1 | 340870 | 340870 | 340870 | -0.64 | 5.2E-01 | 0 |
| Diurnal inactivity duration | 2 | 447371 | 649368 | 245374 | -0.64 | 5.3E-01 | 0 |
| FCGR3B - Low affinity immunoglobulin gamma Fc region receptor III-B | 1 | 345368 | 345368 | 345368 | -0.63 | 5.3E-01 | 0 |
| Epithelial ovarian cancer (high-grade serous) | 1 | 349565 | 349565 | 349565 | -0.63 | 5.3E-01 | 0 |
| Monocyte percentage of white cells (three-way meta) | 123 | 615695 | 1089181 | 85329 | -0.63 | 5.3E-01 | 0 |
| Total cholesterol in LDL | 12 | 494950 | 950535 | 86744 | -0.63 | 5.3E-01 | 0 |
| Body Mass Index (male <= 50 yrs) | 6 | 456843 | 1041314 | 338085 | -0.63 | 5.3E-01 | 0 |
| Maximum carotid artery stenosis (at exam 6, GEE, adjusted for multivariable) | 1 | 356285 | 356285 | 356285 | -0.63 | 5.3E-01 | 0 |
| ::::X-02249 | 1 | 347683 | 347683 | 347683 | -0.63 | 5.3E-01 | 0 |
| Temporal brain volume (GEE, adjusted for multivariable) | 1 | 354463 | 354463 | 354463 | -0.63 | 5.3E-01 | 0 |
| BPI - Bactericidal permeability-increasing protein | 1 | 357798 | 357798 | 357798 | -0.62 | 5.3E-01 | 0 |
| Neutrophil count | 16 | 517698 | 843389 | 89762 | -0.62 | 5.3E-01 | 0 |
| Plasma Apolipoprotein C3 level (GEE, adjusted for age and sex) | 1 | 368395 | 368395 | 368395 | -0.62 | 5.4E-01 | 0 |
| IL17RA - Interleukin-17 receptor A | 1 | 350205 | 350205 | 350205 | -0.62 | 5.4E-01 | 0 |
| ::::X-13548 | 2 | 451773 | 501705 | 401840 | -0.62 | 5.4E-01 | 0 |
| ECE1 - Endothelin-converting enzyme 1 | 1 | 356988 | 356988 | 356988 | -0.61 | 5.4E-01 | 0 |
| Antihistamines for systemic use | 5 | 433560 | 1317225 | 407193 | -0.61 | 5.4E-01 | 0 |
| CD40 Ligand serum (at exam 7, GEE, adjusted for age and sex) | 2 | 451672 | 643780 | 259564 | -0.61 | 5.4E-01 | 0 |
| PIGR - Polymeric immunoglobulin receptor | 1 | 373535 | 373535 | 373535 | -0.60 | 5.5E-01 | 0 |
| Chest pain or discomfort | 2 | 460186 | 648639 | 271733 | -0.60 | 5.5E-01 | 0 |
| Thyroid stimulation hormone (at exam 4, GEE, adjusted for multivariable) | 2 | 457443 | 564457 | 350430 | -0.60 | 5.5E-01 | 0 |
| Lipid::Essential fatty acid::eicosapentaenoate (EPA; 20:5n3) | 1 | 369990 | 369990 | 369990 | -0.59 | 5.5E-01 | 0 |
| Acute insulin response (adjusted for BMI) | 2 | 456758 | 617763 | 295753 | -0.59 | 5.6E-01 | 0 |
| Fluid intelligence test - FI3 : word interpolation | 5 | 444250 | 1316713 | 183142 | -0.58 | 5.6E-01 | 0 |
| ::::X-11469 | 3 | 416563 | 544035 | 250161 | -0.58 | 5.6E-01 | 0 |
| Amino acid::Glycine, serine and threonine metabolism::serine | 2 | 461674 | 633210 | 290139 | -0.58 | 5.6E-01 | 0 |
| von Willebrand Factor (at exam 5, GEE, adjusted for age and sex) | 2 | 462091 | 664765 | 259416 | -0.58 | 5.6E-01 | 0 |
| Bone Ultrasound Speed (GEE, adjusted for multivariable) | 1 | 370965 | 370965 | 370965 | -0.58 | 5.6E-01 | 0 |
| Hair colour (natural, before greying): Red | 11 | 499715 | 898259 | 56085 | -0.58 | 5.6E-01 | 0 |
| Pancreatic cancer | 3 | 415423 | 739820 | 277350 | -0.57 | 5.7E-01 | 0 |
| Amino acid::Tryptophan metabolism::indoleacetate | 1 | 378023 | 378023 | 378023 | -0.57 | 5.7E-01 | 0 |
| CCL23 - Ck-beta-8-1 | 1 | 389920 | 389920 | 389920 | -0.57 | 5.7E-01 | 0 |

| | | | | | | | |
|---|---|---|---|---|---|---|---|
| Small high-density lipoprotein by NMR (at exam 4, GEE, adjusted for multivariable) | 5 | 450625 | 918710 | 130815 | -0.57 | 5.7E-01 | 0 |
| Usual weekday bedtime (GEE) | 2 | 474741 | 698103 | 251379 | -0.57 | 5.7E-01 | 0 |
| Medication for pain relief, constipation, heartburn: Laxatives (e.g. Dulcolax, Senokot) | 1 | 380250 | 380250 | 380250 | -0.57 | 5.7E-01 | 0 |
| Coronary heart disease (MI, CI or CHD death, GEE, adjusted for multivariable) | 1 | 377123 | 377123 | 377123 | -0.56 | 5.7E-01 | 0 |
| Major depressive disorder | 3 | 421838 | 627209 | 265592 | -0.56 | 5.7E-01 | 0 |
| THBS2 - Thrombospondin-2 | 1 | 386948 | 386948 | 386948 | -0.56 | 5.8E-01 | 0 |
| Carbohydrate::Glycolysis, gluconeogenesis, pyruvate metabolism::1,5-anhydroglucitol (1,5-AG) | 3 | 427225 | 1101838 | 404371 | -0.56 | 5.8E-01 | 0 |
| Usual weekday bedtime (GEE, adjusted for age, sex and BMI) | 2 | 474741 | 698103 | 251379 | -0.56 | 5.8E-01 | 0 |
| Current employment status: Unable to work because of sickness or disability | 1 | 390623 | 390623 | 390623 | -0.55 | 5.8E-01 | 0 |
| Left superior parietal | 6 | 483566 | 711248 | 166031 | -0.55 | 5.8E-01 | 0 |
| Energy::Krebs cycle::succinylcarnitine | 6 | 485817 | 948119 | 96404 | -0.55 | 5.8E-01 | 0 |
| Prostate cancer (GEE, adjusted for age) | 26 | 563086 | 1018012 | 155701 | -0.55 | 5.8E-01 | 0 |
| Right rostral middle frontal | 1 | 394190 | 394190 | 394190 | -0.55 | 5.8E-01 | 0 |
| Mother still alive | 4 | 468654 | 992926 | 45778 | -0.54 | 5.9E-01 | 0 |
| PI3 - Elafin | 1 | 400733 | 400733 | 400733 | -0.54 | 5.9E-01 | 0 |
| Fornix (column and body of fornix) mode of anisotropy | 2 | 487471 | 713441 | 261502 | -0.54 | 5.9E-01 | 0 |
| CLEC1B - C-type lectin domain family 1 member B | 1 | 406628 | 406628 | 406628 | -0.54 | 5.9E-01 | 0 |
| Treatment/medication code: omega-3/fish oil supplement | 1 | 400145 | 400145 | 400145 | -0.54 | 5.9E-01 | 0 |
| Maximum internal carotid artery IMT (at exam 6, GEE, adjusted for multivariable) | 1 | 396293 | 396293 | 396293 | -0.54 | 5.9E-01 | 0 |
| ::::X-11204 | 1 | 395798 | 395798 | 395798 | -0.54 | 5.9E-01 | 0 |
| Xenobiotics::Food component, Plant::N-(2-furoyl)glycine | 1 | 404515 | 404515 | 404515 | -0.54 | 5.9E-01 | 0 |
| Mean internal carotid artery IMT (at exam 6, GEE, adjusted for multivariable) | 1 | 396293 | 396293 | 396293 | -0.54 | 5.9E-01 | 0 |
| Long sleep | 5 | 454690 | 842328 | 380620 | -0.53 | 5.9E-01 | 0 |
| Age completed full time education | 6 | 484754 | 1301597 | 142163 | -0.53 | 5.9E-01 | 0 |
| Femoral Neck Length (GEE, adjusted for age and sex) | 2 | 489545 | 524355 | 454735 | -0.53 | 6.0E-01 | 0 |
| Hearing difficulty/problems | 25 | 560375 | 1064688 | 183421 | -0.53 | 6.0E-01 | 0 |
| Total cholesterol in medium LDL | 11 | 514640 | 831980 | 196658 | -0.53 | 6.0E-01 | 0 |
| Amino acid::Phenylalanine & tyrosine metabolism::phenylacetylglutamine | 1 | 410853 | 410852 | 410852 | -0.53 | 6.0E-01 | 0 |
| Amino acid::Butanoate metabolism::X-04499--3,4-dihydroxybutyrate | 1 | 403843 | 403843 | 403843 | -0.53 | 6.0E-01 | 0 |
| Remission after 12 weeks of antidepressant treatment in MDD | 1 | 406538 | 406538 | 406538 | -0.52 | 6.0E-01 | 0 |
| Inferior fronto-occipital fasciculus axial diusivities | 4 | 473609 | 582851 | 347378 | -0.52 | 6.0E-01 | 0 |
| Brain natriuretic peptide (at exam 6, GEE, adjusted for age and sex) | 3 | 442838 | 845003 | 226130 | -0.52 | 6.1E-01 | 0 |
| Heart rate | 10 | 514051 | 1113645 | 78809 | -0.51 | 6.1E-01 | 0 |
| CHIT1 - Chitotriosidase-1 | 1 | 414468 | 414468 | 414468 | -0.51 | 6.1E-01 | 0 |
| Osteoprotegerin (at exam 7, GEE, adjusted for multivariable | 2 | 490446 | 594783 | 386109 | -0.51 | 6.1E-01 | 0 |
| CST3 - Cystatin-C | 1 | 414238 | 414238 | 414238 | -0.51 | 6.1E-01 | 0 |
| CGA CGB - Human Chorionic Gonadotropin | 1 | 411445 | 411445 | 411445 | -0.51 | 6.1E-01 | 0 |
| Osteoarthritis | 29 | 568683 | 1044870 | 210350 | -0.51 | 6.1E-01 | 0 |
| Phospholipids in large LDL | 11 | 521590 | 1009733 | 240700 | -0.51 | 6.1E-01 | 0 |
| Lipid::Sterol/Steroid::X-11445--5-alpha-pregnan-3beta,20alpha-disulfate | 1 | 416563 | 416563 | 416563 | -0.50 | 6.2E-01 | 0 |
| Epithelial ovarian cancer (Clear cell) | 1 | 422213 | 422213 | 422213 | -0.49 | 6.2E-01 | 0 |
| Diagnoses - secondary ICD10: Z37 Outcome of delivery | 2 | 495450 | 726366 | 264534 | -0.49 | 6.2E-01 | 0 |
| Monocyte chemotactic protein-1 (CCL2) | 2 | 500645 | 736100 | 265190 | -0.49 | 6.2E-01 | 0 |

| | | | | | | | |
|---|---|---|---|---|---|---|---|
| Alanine transaminase (at exam 2, GEE, adjusted for multivariable) | 1 | 419925 | 419925 | 419925 | -0.49 | 6.2E-01 | 0 |
| Splenium of corpus callosum mean diusivities | 2 | 499750 | 597628 | 401873 | -0.49 | 6.2E-01 | 0 |
| High-density lipoprotein 3 cholesterol (at exam 4, GEE, adjusted for multivariable) | 3 | 451013 | 479461 | 245010 | -0.49 | 6.3E-01 | 0 |
| How are people in household related to participant: Son and/or daughter (include step-children) | 1 | 419693 | 419693 | 419693 | -0.49 | 6.3E-01 | 0 |
| ::::X-12524 | 1 | 424935 | 424935 | 424935 | -0.49 | 6.3E-01 | 0 |
| ::::X-13429 | 1 | 426800 | 426800 | 426800 | -0.49 | 6.3E-01 | 0 |
| Variation in diet | 10 | 520368 | 1001284 | 204209 | -0.48 | 6.3E-01 | 0 |
| Cingulum (cingulate gyrus) mode of anisotropy | 5 | 475135 | 875192 | 49476 | -0.48 | 6.3E-01 | 0 |
| Colorectal cancer | 9 | 507910 | 1064808 | 338440 | -0.48 | 6.3E-01 | 0 |
| Ever used hormone-replacement therapy (HRT) (female) | 5 | 475463 | 1105553 | 66235 | -0.48 | 6.3E-01 | 0 |
| Male-specific factors - Hair/balding pattern: Pattern 2 | 45 | 597060 | 1010020 | 217307 | -0.48 | 6.3E-01 | 0 |
| Cholesterol esters in medium LDL | 14 | 544955 | 1053568 | 173578 | -0.48 | 6.3E-01 | 0 |
| Obesity class 2 | 10 | 526273 | 640331 | 283569 | -0.48 | 6.3E-01 | 0 |
| Posterior thalamic radiation (include optic radiation) mean diusivities | 2 | 499750 | 597628 | 401873 | -0.48 | 6.3E-01 | 0 |
| ::::X-14626 | 1 | 426800 | 426800 | 426800 | -0.48 | 6.3E-01 | 0 |
| Total brain volume | 8 | 512325 | 1109881 | 224513 | -0.48 | 6.3E-01 | 0 |
| Fluid intelligence test - FI6 : conditional arithmetic | 1 | 428080 | 428080 | 428080 | -0.47 | 6.4E-01 | 0 |
| Numeric memory test - Maximum digits remembered correctly | 6 | 509016 | 1608511 | 62861 | -0.47 | 6.4E-01 | 0 |
| Lipid::Fatty acid, dicarboxylate::tetradecanedioate | 1 | 426800 | 426800 | 426800 | -0.47 | 6.4E-01 | 0 |
| Frequency of stair climbing in last 4 weeks | 6 | 504911 | 915942 | 413849 | -0.47 | 6.4E-01 | 0 |
| Number of children ever born (male) | 1 | 432875 | 432875 | 432875 | -0.47 | 6.4E-01 | 0 |
| Mean Arterial Pressure | 14 | 540890 | 934514 | 272220 | -0.47 | 6.4E-01 | 0 |
| Posterior thalamic radiation (include optic radiation) radial diusivities | 2 | 499750 | 597628 | 401873 | -0.47 | 6.4E-01 | 0 |
| Superior longitudinal fasciculus fractional anisotropy | 4 | 498528 | 751069 | 233972 | -0.47 | 6.4E-01 | 0 |
| Waist circumference by CT (GEE, adjusted for age and sex) | 2 | 505384 | 737206 | 273563 | -0.47 | 6.4E-01 | 0 |
| Usual walking pace | 40 | 598774 | 1174531 | 130097 | -0.47 | 6.4E-01 | 0 |
| Duration of moderate activity | 5 | 487838 | 856333 | 432443 | -0.47 | 6.4E-01 | 0 |
| Had menopause (female) | 28 | 580544 | 1140433 | 109921 | -0.47 | 6.4E-01 | 0 |
| Fasting proinsulin (adjusted for fasting insulin, age and sex) | 7 | 499810 | 935071 | 285636 | -0.47 | 6.4E-01 | 0 |
| Temporal brain volume (GEE, adjusted for age and sex) | 2 | 503445 | 577936 | 428954 | -0.46 | 6.4E-01 | 0 |
| Lipid::Essential fatty acid::docosapentaenoate (n3 DPA; 22:5n3) | 1 | 439805 | 439805 | 439805 | -0.46 | 6.4E-01 | 0 |
| Peptide::gamma-glutamyl::gamma-glutamylphenylalanine | 1 | 436478 | 436477 | 436477 | -0.46 | 6.5E-01 | 0 |
| CCL18 - C-C motif chemokine 18 | 1 | 451863 | 451863 | 451863 | -0.45 | 6.5E-01 | 0 |
| Waist circumference by CT (GEE, adjusted for age, sex, smoking and menopause) | 2 | 505384 | 737206 | 273563 | -0.45 | 6.5E-01 | 0 |
| PPIE - Peptidyl-prolyl cis-trans isomerase E | 1 | 445040 | 445040 | 445040 | -0.45 | 6.5E-01 | 0 |
| Temporal horn volume (GEE, adjusted for age and sex) | 1 | 437120 | 437120 | 437120 | -0.45 | 6.5E-01 | 0 |
| Lumbar Spine BMD (GEE, adjusted for multivariable) | 2 | 507674 | 743521 | 271828 | -0.45 | 6.5E-01 | 0 |
| NTN4 - Netrin-4 | 1 | 455813 | 455813 | 455813 | -0.45 | 6.6E-01 | 0 |
| Osteoarthritis of knee | 8 | 526686 | 1263233 | 396324 | -0.44 | 6.6E-01 | 0 |
| Diabetes (diagnosed by doctor) | 54 | 615524 | 1079431 | 158394 | -0.44 | 6.6E-01 | 0 |
| Positive affect (univariate) | 8 | 523256 | 1148566 | 94318 | -0.44 | 6.6E-01 | 0 |
| ::::X-11381 | 1 | 448098 | 448098 | 448098 | -0.44 | 6.6E-01 | 0 |
| Dysmenorrhea pain severity | 2 | 514651 | 746114 | 283188 | -0.44 | 6.6E-01 | 0 |

| | | | | | | | |
|---|---|---|---|---|---|---|---|
| Erectile dysfunction | 1 | 453883 | 453883 | 453883 | -0.43 | 6.6E-01 | 0 |
| CCL3 - C-C motif chemokine 3 | 1 | 451863 | 451863 | 451863 | -0.43 | 6.6E-01 | 0 |
| Total IgA levels | 1 | 456193 | 456193 | 456193 | -0.43 | 6.7E-01 | 0 |
| External capsule axial diusivities | 1 | 466160 | 466160 | 466160 | -0.43 | 6.7E-01 | 0 |
| CCL3L1 - C-C motif chemokine 3-like 1 | 1 | 451863 | 451863 | 451863 | -0.43 | 6.7E-01 | 0 |
| Ever had hysterectomy (womb removed) (female) | 2 | 524988 | 765437 | 284540 | -0.43 | 6.7E-01 | 0 |
| Maximum common carotid artery IMT (at exam 6, GEE, adjusted for age and sex) | 3 | 481390 | 683709 | 334883 | -0.42 | 6.7E-01 | 0 |
| Average across all tracts radial diusivities | 4 | 516526 | 761518 | 243521 | -0.42 | 6.7E-01 | 0 |
| Fatty acid length | 1 | 459733 | 459733 | 459733 | -0.42 | 6.8E-01 | 0 |
| ApoB | 14 | 554768 | 970919 | 173578 | -0.41 | 6.8E-01 | 0 |
| Urinary isoprostanes/creatinine (at exam 7, GEE, adjusted for age and sex) | 3 | 487203 | 581389 | 259864 | -0.41 | 6.8E-01 | 0 |
| Mineral and other dietary supplements: Zinc | 1 | 472285 | 472285 | 472285 | -0.40 | 6.9E-01 | 0 |
| Genu of corpus callosum mean diusivities | 5 | 504515 | 740255 | 303995 | -0.40 | 6.9E-01 | 0 |
| Eosinophil count | 13 | 550998 | 954342 | 96751 | -0.40 | 6.9E-01 | 0 |
| Age last used hormone-replacement therapy (HRT) (female) | 1 | 473425 | 473425 | 473425 | -0.40 | 6.9E-01 | 0 |
| IL18RAP - Interleukin-18 receptor accessory protein | 1 | 474333 | 474333 | 474333 | -0.40 | 6.9E-01 | 0 |
| Overweight | 13 | 555613 | 992923 | 323375 | -0.39 | 6.9E-01 | 0 |
| Body Mass Index (female <= 50 yrs) | 18 | 574599 | 1202961 | 192863 | -0.39 | 7.0E-01 | 0 |
| Lipid::Fatty acid, dicarboxylate::hexadecanedioate | 1 | 482085 | 482085 | 482085 | -0.39 | 7.0E-01 | 0 |
| Right lateral ventricle | 12 | 558718 | 912548 | 195172 | -0.39 | 7.0E-01 | 0 |
| Hippocampal volume (GEE, adjusted for multivariable with APOE) | 2 | 538554 | 599521 | 477587 | -0.38 | 7.0E-01 | 0 |
| Fasting glucose interaction (adjusted for BMI) | 6 | 533355 | 751131 | 516461 | -0.38 | 7.0E-01 | 0 |
| SPINT1 - Kunitz-type protease inhibitor 1 | 1 | 483008 | 483008 | 483008 | -0.38 | 7.1E-01 | 0 |
| MED1 - Mediator of RNA polymerase II transcription subunit 1 | 1 | 484898 | 484898 | 484898 | -0.38 | 7.1E-01 | 0 |
| Freckles | 4 | 530375 | 666042 | 333886 | -0.38 | 7.1E-01 | 0 |
| Reticulocyte fraction of red cells (three-way meta) | 114 | 637324 | 1092624 | 113725 | -0.38 | 7.1E-01 | 0 |
| Fasting glucose | 15 | 564913 | 974776 | 167229 | -0.38 | 7.1E-01 | 0 |
| ::::X-11538 | 1 | 482085 | 482085 | 482085 | -0.37 | 7.1E-01 | 0 |
| Ankle-brachial index (at exam 7, GEE, adjusted for age and sex) | 2 | 547524 | 551768 | 543279 | -0.37 | 7.1E-01 | 0 |
| Total cholesterol in large LDL | 12 | 563180 | 1090583 | 307823 | -0.37 | 7.1E-01 | 0 |
| Right precentral | 5 | 526250 | 875192 | 29254 | -0.36 | 7.2E-01 | 0 |
| MYCN amplification neuroblastoma | 2 | 547140 | 798676 | 295603 | -0.36 | 7.2E-01 | 0 |
| Interleukin-7 | 1 | 496443 | 496443 | 496443 | -0.36 | 7.2E-01 | 0 |
| Alcohol consumption (dichotomous) | 1 | 492088 | 492088 | 492088 | -0.35 | 7.2E-01 | 0 |
| How are people in household related to participant: Husband, wife or partner | 1 | 490698 | 490698 | 490698 | -0.35 | 7.2E-01 | 0 |
| Prothrombin time | 8 | 554565 | 1070068 | 373716 | -0.35 | 7.2E-01 | 0 |
| Left amygdala | 4 | 536671 | 727938 | 276053 | -0.35 | 7.3E-01 | 0 |
| Blood sugar | 15 | 569313 | 907016 | 90606 | -0.35 | 7.3E-01 | 0 |
| CAPN1 CAPNS1 - Calpain I | 1 | 495535 | 495535 | 495535 | -0.35 | 7.3E-01 | 0 |
| Waist-hip ratio (male > 50 yrs, adjusted for BMI) | 3 | 514313 | 794198 | 286605 | -0.35 | 7.3E-01 | 0 |
| LAG3 - Lymphocyte activation gene 3 protein | 1 | 495908 | 495908 | 495908 | -0.34 | 7.3E-01 | 0 |
| Immature fraction of reticulocytes (two-way meta) | 80 | 634590 | 1058338 | 111568 | -0.34 | 7.3E-01 | 0 |
| Chronotype (continuous) | 5 | 532973 | 965650 | 130692 | -0.34 | 7.3E-01 | 0 |

| | | | | | | | |
|---|---|---|---|---|---|---|---|
| CCL8 - C-C motif chemokine 8 | 2 | 558481 | 710851 | 406112 | -0.34 | 7.3E-01 | 0 |
| Genu of corpus callosum mode of anisotropy | 1 | 491515 | 491515 | 491515 | -0.34 | 7.4E-01 | 0 |
| Right superior parietal | 4 | 540659 | 764921 | 498629 | -0.34 | 7.4E-01 | 0 |
| Remnant lipoprotein triglycerides (at exam 4, GEE, adjusted for age and sex) | 2 | 560877 | 802005 | 319749 | -0.33 | 7.4E-01 | 0 |
| CD32 on 11c+123+DC | 1 | 508515 | 508515 | 508515 | -0.33 | 7.4E-01 | 0 |
| Reason for glasses/contact lenses: For short-sightedness, i.e. only or mainly for distance viewing | 38 | 614573 | 1209086 | 284724 | -0.33 | 7.4E-01 | 0 |
| Interleukin-18 | 2 | 551358 | 818120 | 284595 | -0.33 | 7.4E-01 | 0 |
| Cancer register - Histology of cancer tumour: Adenocarcinoma, NOS | 10 | 565555 | 852391 | 77790 | -0.33 | 7.4E-01 | 0 |
| High-density lipoprotein 2 cholesterol (at exam 4, GEE, adjusted for age and sex) | 2 | 565986 | 579462 | 552511 | -0.33 | 7.4E-01 | 0 |
| CFI - Complement factor I | 1 | 511358 | 511358 | 511358 | -0.33 | 7.5E-01 | 0 |
| Duration of light DIY | 2 | 560493 | 578058 | 542928 | -0.32 | 7.5E-01 | 0 |
| Atrial natriuretic peptide (at exam 6, GEE, adjusted for age and sex) | 1 | 511103 | 511102 | 511102 | -0.32 | 7.5E-01 | 0 |
| Amino acid::Glutathione metabolism::cysteine-glutathione disulfide | 1 | 506408 | 506408 | 506408 | -0.32 | 7.5E-01 | 0 |
| Cheese intake | 77 | 636550 | 1104145 | 87946 | -0.32 | 7.5E-01 | 0 |
| Friendships satisfaction | 3 | 524345 | 856514 | 474481 | -0.32 | 7.5E-01 | 0 |
| QT interval | 10 | 572545 | 862658 | 175184 | -0.31 | 7.6E-01 | 0 |
| Drinks per week | 47 | 624858 | 1257091 | 61640 | -0.31 | 7.6E-01 | 0 |
| Chloride | 14 | 582996 | 1098427 | 54292 | -0.31 | 7.6E-01 | 0 |
| White matter hyperintensities | 3 | 528005 | 810336 | 416841 | -0.31 | 7.6E-01 | 0 |
| Total body BMD (15 or younger) | 7 | 551500 | 1039185 | 244137 | -0.31 | 7.6E-01 | 0 |
| Eosinophil count (three-way meta) | 117 | 645863 | 1069568 | 84423 | -0.30 | 7.6E-01 | 0 |
| Cerebellar vermal lobules VI VII | 12 | 576470 | 843186 | 185615 | -0.30 | 7.6E-01 | 0 |
| Menstrual cycle length | 5 | 544048 | 1117228 | 78471 | -0.30 | 7.6E-01 | 0 |
| CDON - Cell adhesion molecule-related/down-regulated by oncogenes | 1 | 524285 | 524285 | 524285 | -0.29 | 7.7E-01 | 0 |
| CD4:%RTE | 1 | 524605 | 524605 | 524605 | -0.29 | 7.7E-01 | 0 |
| ::::X-11550 | 1 | 527550 | 527550 | 527550 | -0.29 | 7.7E-01 | 0 |
| Vitamin K percentage of undercarboxylated osteocalcin (at exam 6 or 7, GEE, adjusted for age a | 9 | 568983 | 576395 | 430827 | -0.29 | 7.7E-01 | 0 |
| Vitamin K plasma phylloquinone (at exam 6 or 7, GEE, adjusted for age and sex) | 2 | 579233 | 829539 | 328927 | -0.29 | 7.7E-01 | 0 |
| Anterior limb of internal capsule radial diusivities | 2 | 575501 | 835419 | 315582 | -0.29 | 7.7E-01 | 0 |
| Monocyte count (three-way meta) | 134 | 648628 | 1093106 | 97777 | -0.29 | 7.7E-01 | 0 |
| Age at first live birth (female) | 15 | 590413 | 1089984 | 109928 | -0.28 | 7.8E-01 | 0 |
| Asthma, hay fever or eczema | 58 | 633619 | 1197049 | 220662 | -0.28 | 7.8E-01 | 0 |
| Subcutaneous adipose tissue attenuation (male) | 1 | 544410 | 544410 | 544410 | -0.28 | 7.8E-01 | 0 |
| Cingulum (cingulate gyrus) axial diusivities | 4 | 557716 | 710658 | 368659 | -0.28 | 7.8E-01 | 0 |
| Inferior fronto-occipital fasciculus mean diusivities | 2 | 573296 | 633178 | 513414 | -0.28 | 7.8E-01 | 0 |
| ::::X-12456 | 2 | 576224 | 839023 | 313425 | -0.28 | 7.8E-01 | 0 |
| Total lipids in small LDL | 12 | 589284 | 1089553 | 296573 | -0.28 | 7.8E-01 | 0 |
| Posterior limb of internal capsule axial diusivities | 6 | 562368 | 1140271 | 372944 | -0.28 | 7.8E-01 | 0 |
| HOMA-IR (at exam 7, GEE, adjusted for age and sex) | 1 | 545980 | 545980 | 545980 | -0.27 | 7.8E-01 | 0 |
| Menstrual fever | 1 | 548240 | 548240 | 548240 | -0.27 | 7.9E-01 | 0 |
| Sleep duration (mean) | 7 | 568635 | 664266 | 43752 | -0.27 | 7.9E-01 | 0 |
| Vitiligo (late onset) | 5 | 559010 | 665375 | 86496 | -0.27 | 7.9E-01 | 0 |
| Total cholesterol in HDL | 11 | 586548 | 1042579 | 74752 | -0.27 | 7.9E-01 | 0 |

| | | | | | | | |
|---|---|---|---|---|---|---|---|
| Fasting insulin (at exam 7, GEE, adjusted for age and sex) | 1 | 545980 | 545980 | 545980 | -0.26 | 8.0E-01 | 0 |
| F11 - Coagulation Factor XI | 2 | 582585 | 651993 | 513178 | -0.25 | 8.0E-01 | 0 |
| Amino acid::Tryptophan metabolism::X-12100--hydroxytryptophan* | 3 | 552740 | 987160 | 508206 | -0.25 | 8.0E-01 | 0 |
| NID2 - Nidogen-2 | 1 | 556970 | 556970 | 556970 | -0.25 | 8.0E-01 | 0 |
| Peptide::Dipeptide::pro-hydroxy-pro | 1 | 551180 | 551180 | 551180 | -0.25 | 8.0E-01 | 0 |
| Types of physical activity in last 4 weeks: Light DIY (eg: pruning, watering the lawn) | 5 | 564225 | 842960 | 76649 | -0.25 | 8.0E-01 | 0 |
| Openness to Experience (NEO-FFI) | 1 | 550570 | 550570 | 550570 | -0.25 | 8.0E-01 | 0 |
| Double bonds in fatty acids | 3 | 548893 | 713123 | 521474 | -0.25 | 8.0E-01 | 0 |
| Relative age of first facial hair (male) | 78 | 641573 | 1085894 | 148743 | -0.25 | 8.0E-01 | 0 |
| F7 - Coagulation Factor VII | 2 | 591532 | 795001 | 388063 | -0.25 | 8.0E-01 | 0 |
| Low-density-lipoprotein cholesterol | 16 | 605781 | 972744 | 319617 | -0.25 | 8.0E-01 | 0 |
| Vitamin K percentage of undercarboxylated osteocalcin (at exam 6 or 7, GEE, adjusted for multiv | 6 | 579429 | 710328 | 550473 | -0.25 | 8.0E-01 | 0 |
| CD34 on Stem | 2 | 585721 | 747239 | 424203 | -0.25 | 8.0E-01 | 0 |
| PRSS3 - Trypsin-3 | 1 | 554135 | 554135 | 554135 | -0.25 | 8.0E-01 | 0 |
| Cingulum (cingulate gyrus) radial diusivities | 2 | 589824 | 732738 | 446909 | -0.25 | 8.1E-01 | 0 |
| Types of transport used (excluding work): Walk | 7 | 572478 | 1078124 | 475506 | -0.25 | 8.1E-01 | 0 |
| CCL17 - C-C motif chemokine 17 | 3 | 563553 | 799484 | 296644 | -0.24 | 8.1E-01 | 0 |
| Reaction time | 39 | 627113 | 991665 | 187568 | -0.24 | 8.1E-01 | 0 |
| MMSE score (at age 65, GEE, adjusted for birth-cohort) | 2 | 592761 | 814936 | 370587 | -0.24 | 8.1E-01 | 0 |
| Cholesterol esters in very large HDL | 7 | 576970 | 924116 | 300392 | -0.24 | 8.1E-01 | 0 |
| Reason for reducing amount of alcohol drunk: Other reason | 2 | 594386 | 865441 | 323332 | -0.23 | 8.1E-01 | 0 |
| Blood urea nitrogen | 160 | 655081 | 1009751 | 162559 | -0.23 | 8.2E-01 | 0 |
| Body Mass Index (female > 50 yrs) | 16 | 610561 | 831621 | 266421 | -0.23 | 8.2E-01 | 0 |
| Non-cancer illness code, self-reported: diabetes | 35 | 631385 | 1039403 | 127300 | -0.23 | 8.2E-01 | 0 |
| Suffer from 'nerves' | 9 | 586548 | 1331945 | 79348 | -0.23 | 8.2E-01 | 0 |
| High-density lipoprotein cholesterol (at exam 3, GEE, adjusted for age and sex) | 1 | 553705 | 553705 | 553705 | -0.23 | 8.2E-01 | 0 |
| Total cerebral brain volume (GEE, adjusted for age and sex) | 2 | 602395 | 802206 | 402583 | -0.22 | 8.2E-01 | 0 |
| Overall activity (conditioning sex and BMI) | 2 | 596295 | 876853 | 315738 | -0.22 | 8.2E-01 | 0 |
| Periodontal complex trait 4 | 1 | 570673 | 570673 | 570673 | -0.22 | 8.3E-01 | 0 |
| ITGA1 ITGB1 - Integrin alpha-I: beta-1 complex | 1 | 573303 | 573303 | 573303 | -0.22 | 8.3E-01 | 0 |
| Age spot | 2 | 604110 | 693733 | 514487 | -0.22 | 8.3E-01 | 0 |
| Body Mass Index (male > 50 yrs) | 18 | 614885 | 947845 | 132192 | -0.21 | 8.3E-01 | 0 |
| Mitral annular calcification | 1 | 579550 | 579550 | 579550 | -0.21 | 8.3E-01 | 0 |
| Triglycerides (at exam 2, GEE, adjusted for age and sex) | 1 | 570958 | 570958 | 570958 | -0.21 | 8.3E-01 | 0 |
| Verbal-numerical reasoning | 27 | 625013 | 1025048 | 106612 | -0.21 | 8.3E-01 | 0 |
| Concentration of small LDL particles | 8 | 589284 | 877694 | 382584 | -0.21 | 8.3E-01 | 0 |
| HIPK3 - Homeodomain-interacting protein kinase 3 | 1 | 562288 | 562288 | 562288 | -0.21 | 8.3E-01 | 0 |
| Visceral fat by CT (GEE, adjusted for age and sex) | 2 | 600713 | 867063 | 334363 | -0.21 | 8.3E-01 | 0 |
| Average across all tracts mode of anisotropy | 2 | 597093 | 793964 | 400222 | -0.21 | 8.4E-01 | 0 |
| Non-cancer illness code, self-reported: angina | 13 | 607173 | 911330 | 151638 | -0.21 | 8.4E-01 | 0 |
| Non-cancer illness code, self-reported: hayfever/allergic rhinitis | 13 | 604133 | 1105033 | 240679 | -0.21 | 8.4E-01 | 0 |
| Age at natural menopause (GEE) | 2 | 610135 | 672518 | 547753 | -0.20 | 8.4E-01 | 0 |
| Corrected insulin response (adjusted for insulin sensitivity index) | 2 | 608554 | 845457 | 371651 | -0.20 | 8.4E-01 | 0 |

| | | | | | | | |
|---|---|---|---|---|---|---|---|
| Visceral fat by CT (GEE, adjusted for age, sex, smoking, menopause) | 2 | 600713 | 867063 | 334363 | -0.20 | 8.4E-01 | 0 |
| Types of transport used (excluding work): Car/motor vehicle | 1 | 579423 | 579423 | 579423 | -0.20 | 8.5E-01 | 0 |
| Cholesterol esters in large LDL | 13 | 604770 | 1055113 | 374945 | -0.19 | 8.5E-01 | 0 |
| Aspartate aminotransferase | 20 | 619345 | 1047219 | 48244 | -0.19 | 8.5E-01 | 0 |
| Concentration of medium HDL particles | 3 | 586548 | 984386 | 328255 | -0.19 | 8.5E-01 | 0 |
| SEMA3E - Semaphorin-3E | 1 | 578428 | 578428 | 578428 | -0.19 | 8.5E-01 | 0 |
| Average across all tracts axial diusivities | 4 | 597609 | 837311 | 460609 | -0.19 | 8.5E-01 | 0 |
| Mean triglycerides from exam 1-7 (GEE, adjusted for multivariable) | 2 | 615177 | 804173 | 426180 | -0.19 | 8.5E-01 | 0 |
| Mean triglycerides from exam 1-7 (GEE, adjusted for age and sex) | 2 | 615177 | 804173 | 426180 | -0.18 | 8.5E-01 | 0 |
| Lipoprotein A (at exam 3, GEE, adjusted for multivariable) | 14 | 613269 | 1136285 | 107248 | -0.18 | 8.5E-01 | 0 |
| JAG1 - Protein jagged-1 | 2 | 619229 | 650998 | 587459 | -0.18 | 8.6E-01 | 0 |
| Overall health rating | 47 | 643415 | 1153283 | 125768 | -0.18 | 8.6E-01 | 0 |
| Focal Epilepsy, hippocampal sclerosis (HS) | 2 | 614200 | 817980 | 410421 | -0.18 | 8.6E-01 | 0 |
| Total cholesterol in small LDL | 9 | 604770 | 941313 | 385130 | -0.18 | 8.6E-01 | 0 |
| Number of self-reported cancers | 7 | 597025 | 744341 | 321497 | -0.18 | 8.6E-01 | 0 |
| Broad depression | 7 | 593940 | 769674 | 481033 | -0.17 | 8.6E-01 | 0 |
| Life span | 2 | 615808 | 891731 | 339884 | -0.17 | 8.6E-01 | 0 |
| Coffee intake | 22 | 626030 | 1211758 | 75279 | -0.17 | 8.6E-01 | 0 |
| ROR1 - Tyrosine-protein kinase transmembrane receptor ROR1 | 1 | 583968 | 583968 | 583968 | -0.17 | 8.6E-01 | 0 |
| PDCD1LG2 - Programmed cell death 1 ligand 2 | 1 | 588338 | 588338 | 588338 | -0.17 | 8.6E-01 | 0 |
| 28 year time-averaged Fasting plasma glucose (GEE, adjusted for age, sex and BMI) | 8 | 599569 | 935993 | 113052 | -0.17 | 8.6E-01 | 0 |
| Waist-hip ratio (male <= 50 yrs, adjusted for BMI) | 1 | 592133 | 592133 | 592133 | -0.17 | 8.7E-01 | 0 |
| Cofactors and vitamins::Hemoglobin and porphyrin metabolism::biliverdin | 1 | 600395 | 600395 | 600395 | -0.17 | 8.7E-01 | 0 |
| Low-density lipoprotein cholesterol (at exam 4, GEE, adjusted for age and sex) | 3 | 588370 | 945861 | 314979 | -0.17 | 8.7E-01 | 0 |
| Treatment/medication code: ventolin 100micrograms inhaler | 12 | 618721 | 860567 | 281457 | -0.16 | 8.7E-01 | 0 |
| Occipital brain volume (GEE, adjusted for multivariable) | 1 | 597905 | 597905 | 597905 | -0.16 | 8.7E-01 | 0 |
| Mean low-density lipoprotein cholesterol fromexam 1-7 (GEE, adjusted for age and sex) | 3 | 588370 | 836973 | 308513 | -0.16 | 8.7E-01 | 0 |
| Cofactors and vitamins::Hemoglobin and porphyrin metabolism::bilirubin (Z,Z) | 1 | 600395 | 600395 | 600395 | -0.16 | 8.7E-01 | 0 |
| Cofactors and vitamins::Hemoglobin and porphyrin metabolism::bilirubin (E,Z or Z,E)* | 1 | 600395 | 600395 | 600395 | -0.16 | 8.7E-01 | 0 |
| Activated partial thromboplastin time | 6 | 603606 | 912434 | 153682 | -0.16 | 8.7E-01 | 0 |
| Fasting plasma glucose (at exam 5, GEE, adjusted for age and sex) | 3 | 585880 | 744233 | 345087 | -0.16 | 8.7E-01 | 0 |
| Occipital brain volume (GEE, adjusted for age and sex) | 1 | 597905 | 597905 | 597905 | -0.16 | 8.8E-01 | 0 |
| Residual from predicted FEF25-75/FVC for latest exam (GEE, adjusted for multiple covariates) | 4 | 603766 | 803601 | 561251 | -0.15 | 8.8E-01 | 0 |
| Total cholesterol in very large HDL | 8 | 606231 | 1088621 | 528099 | -0.15 | 8.8E-01 | 0 |
| ApoA1 | 6 | 606613 | 1066224 | 196449 | -0.15 | 8.8E-01 | 0 |
| Total lipids in medium HDL | 3 | 586548 | 984386 | 328255 | -0.15 | 8.8E-01 | 0 |
| Drive faster than motorway speed limit | 21 | 634753 | 988613 | 213280 | -0.15 | 8.8E-01 | 0 |
| Years since last cervical smear test (female) | 6 | 603016 | 805232 | 178825 | -0.15 | 8.8E-01 | 0 |
| Cofactors and vitamins::Hemoglobin and porphyrin metabolism::X-11793--oxidized bilirubin* | 1 | 600395 | 600395 | 600395 | -0.15 | 8.8E-01 | 0 |
| Corticospinal tract mode of anisotropy | 1 | 604068 | 604068 | 604068 | -0.15 | 8.8E-01 | 0 |
| ::::X-11530 | 1 | 600395 | 600395 | 600395 | -0.15 | 8.8E-01 | 0 |
| SELL - L-Selectin | 1 | 608283 | 608283 | 608283 | -0.14 | 8.9E-01 | 0 |
| Cofactors and vitamins::Hemoglobin and porphyrin metabolism::bilirubin (E,E)* | 1 | 600395 | 600395 | 600395 | -0.14 | 8.9E-01 | 0 |

| | | | | | | | |
|---|---|---|---|---|---|---|---|
| ::::X-13215 | 1 | 603415 | 603415 | 603415 | -0.14 | 8.9E-01 | 0 |
| Phosphorus | 6 | 606580 | 927589 | 355484 | -0.14 | 8.9E-01 | 0 |
| Diagnoses - secondary ICD10: Z85 Personal history of malignant neoplasm | 1 | 609465 | 609465 | 609465 | -0.14 | 8.9E-01 | 0 |
| Lipid::Carnitine metabolism::cis-4-decenoyl carnitine | 2 | 625054 | 894417 | 355691 | -0.14 | 8.9E-01 | 0 |
| Concentration of chylomicrons and extremely large VLDL particles | 7 | 614910 | 976125 | 340115 | -0.14 | 8.9E-01 | 0 |
| D-dimer (at exam 6, GEE, adjusted for multivariable) | 2 | 631283 | 758655 | 503910 | -0.14 | 8.9E-01 | 0 |
| Left postcentral | 6 | 617603 | 665156 | 322343 | -0.13 | 9.0E-01 | 0 |
| Right thalamus proper | 4 | 613213 | 1213465 | 51112 | -0.13 | 9.0E-01 | 0 |
| Multiple Sclerosis | 18 | 636436 | 1019228 | 117572 | -0.13 | 9.0E-01 | 0 |
| IL15RA - Interleukin-15 receptor subunit alpha | 1 | 622523 | 622523 | 622523 | -0.12 | 9.0E-01 | 0 |
| Treatment/medication code: bendroflumethiazide | 16 | 632978 | 1072324 | 142796 | -0.12 | 9.0E-01 | 0 |
| Eosinophil percentage of granulocytes (two-way meta) | 97 | 660213 | 1131560 | 141814 | -0.12 | 9.0E-01 | 0 |
| B:%Mature | 1 | 617010 | 617010 | 617010 | -0.12 | 9.0E-01 | 0 |
| ::::X-11440 | 1 | 622075 | 622075 | 622075 | -0.12 | 9.0E-01 | 0 |
| Alcohol dependency (full discovery samples) | 1 | 620200 | 620200 | 620200 | -0.12 | 9.0E-01 | 0 |
| Neutrophil percentage of granulocytes (two-way meta) | 95 | 660213 | 1132070 | 178427 | -0.11 | 9.1E-01 | 0 |
| Diagnoses - main ICD10: M17 Osteoarthritis of knee | 3 | 614298 | 1249150 | 476524 | -0.11 | 9.1E-01 | 0 |
| Cigarettes per day (excluding nonsmokers) | 2 | 637711 | 918059 | 357363 | -0.11 | 9.1E-01 | 0 |
| Right inferior lateral ventricle | 3 | 610818 | 819869 | 320824 | -0.10 | 9.2E-01 | 0 |
| Cancer register - Histology of cancer tumour: Basal cell carcinoma, NOS | 24 | 644454 | 930931 | 98940 | -0.10 | 9.2E-01 | 0 |
| Lipid::Long chain fatty acid::adrenate (22:4n6) | 1 | 631788 | 631788 | 631788 | -0.10 | 9.2E-01 | 0 |
| Triglycerides in medium VLDL | 8 | 628013 | 1152693 | 85903 | -0.09 | 9.3E-01 | 0 |
| Glycine | 4 | 629152 | 1251847 | 41670 | -0.09 | 9.3E-01 | 0 |
| Asthma | 20 | 643551 | 1005998 | 292181 | -0.08 | 9.3E-01 | 0 |
| Sagittal stratum mode of anisotropy | 2 | 658305 | 969691 | 346919 | -0.08 | 9.3E-01 | 0 |
| Hypertriglyceridemia | 4 | 629379 | 711532 | 580843 | -0.08 | 9.3E-01 | 0 |
| Neck cross-sectional moment of inertia (GEE, male, adjussted for age) | 1 | 646793 | 646793 | 646793 | -0.08 | 9.4E-01 | 0 |
| Neutrophil percentage of white cells (two-way meta) | 80 | 663046 | 1133838 | 93413 | -0.08 | 9.4E-01 | 0 |
| NKeff:%314-158a+ | 330 | 669176 | 1027133 | 213042 | -0.08 | 9.4E-01 | 0 |
| Pain type(s) experienced in last month: Stomach or abdominal pain | 2 | 650809 | 888447 | 413171 | -0.07 | 9.4E-01 | 0 |
| Asthma (child-onset) | 67 | 661478 | 1088499 | 299055 | -0.07 | 9.4E-01 | 0 |
| Seen doctor (GP) for nerves, anxiety, tension or depression | 25 | 654258 | 1139650 | 95918 | -0.07 | 9.4E-01 | 0 |
| Wheeze or whistling in the chest in last year | 24 | 651598 | 1131414 | 293286 | -0.07 | 9.4E-01 | 0 |
| Morning/evening person (chronotype) | 112 | 667319 | 1162164 | 119263 | -0.07 | 9.4E-01 | 0 |
| Left insula | 4 | 627782 | 1311636 | 68113 | -0.07 | 9.4E-01 | 0 |
| Tea intake | 21 | 648410 | 1164618 | 306338 | -0.07 | 9.4E-01 | 0 |
| Total cholesterol / High-density lipoprotein cholesterol ratio (at exam 7, GEE, adjusted for age an | 1 | 637505 | 637505 | 637505 | -0.07 | 9.4E-01 | 0 |
| CD55 - Complement decay-accelerating factor | 1 | 648823 | 648823 | 648823 | -0.07 | 9.4E-01 | 0 |
| AIMP1 - Endothelial monocyte-activating polypeptide 2 | 1 | 649273 | 649273 | 649273 | -0.07 | 9.4E-01 | 0 |
| Focal Epilepsy | 1 | 649165 | 649165 | 649165 | -0.07 | 9.4E-01 | 0 |
| Diagnoses - secondary ICD10: E03 Other hypothyroidism | 40 | 657540 | 1108684 | 410239 | -0.07 | 9.5E-01 | 0 |
| Age-related Macular Degeneration | 31 | 656655 | 997913 | 263322 | -0.06 | 9.5E-01 | 0 |
| Age at first birth (female) | 5 | 629393 | 1276083 | 430185 | -0.06 | 9.5E-01 | 0 |

| | | | | | | | |
|---|---|---|---|---|---|---|---|
| Luteinizing hormone (at exam 3, GEE, adjusted for age and sex) | 3 | 632718 | 719479 | 625394 | -0.06 | 9.5E-01 | 0 |
| Hearing aid user | 3 | 626783 | 945589 | 324752 | -0.06 | 9.5E-01 | 0 |
| Nucleotide::Purine metabolism, adenine containing::N1-methyladenosine | 1 | 657820 | 657820 | 657820 | -0.06 | 9.6E-01 | 0 |
| Xenobiotics::Drug::2-methoxyacetaminophen sulfate* | 2 | 659686 | 938874 | 380497 | -0.06 | 9.6E-01 | 0 |
| Treatment/medication code: aspirin | 4 | 642720 | 938733 | 349516 | -0.06 | 9.6E-01 | 0 |
| Non-butter spread type details: Flora Pro-Active or Benecol | 6 | 636213 | 926591 | 402315 | -0.06 | 9.6E-01 | 0 |
| Maximum carotid artery bulb IMT (at exam 6, GEE, adjusted for age and sex) | 1 | 646203 | 646203 | 646203 | -0.05 | 9.6E-01 | 0 |
| ::::X-01911 | 1 | 646865 | 646865 | 646865 | -0.05 | 9.6E-01 | 0 |
| CAMK1 - Calcium/calmodulin-dependent protein kinase type 1 | 2 | 664188 | 932595 | 395781 | -0.05 | 9.6E-01 | 0 |
| Lipid::Medium chain fatty acid::caprylate (8:0) | 2 | 658515 | 894426 | 422604 | -0.05 | 9.6E-01 | 0 |
| Right entorhinal | 3 | 634860 | 931785 | 379027 | -0.05 | 9.6E-01 | 0 |
| Lipid::Sterol, Steroid::5alpha-androstan-3beta,17beta-diol disulfate | 3 | 640390 | 871106 | 345508 | -0.04 | 9.6E-01 | 0 |
| Retrolenticular part of internal capsule mode of anisotropy | 6 | 631640 | 953067 | 325622 | -0.04 | 9.7E-01 | 0 |
| Hippocampal volume (GEE, adjusted for multivariable) | 1 | 660488 | 660488 | 660488 | -0.04 | 9.7E-01 | 0 |
| Treatment/medication code: levothyroxine sodium | 47 | 665375 | 1100020 | 345475 | -0.04 | 9.7E-01 | 0 |
| Hippocampal volume (GEE, adjusted for age and sex) | 1 | 660488 | 660488 | 660488 | -0.04 | 9.7E-01 | 0 |
| Amino acid::Glutamate metabolism::pyroglutamine* | 4 | 639446 | 1027879 | 362622 | -0.04 | 9.7E-01 | 0 |
| AGER - Advanced glycosylation end product-specific receptor, soluble | 1 | 667833 | 667833 | 667833 | -0.03 | 9.7E-01 | 0 |
| Depression - Recent lack of interenst of pleasure in doing things | 1 | 662578 | 662578 | 662578 | -0.03 | 9.7E-01 | 0 |
| CST6 - Cystatin-M | 1 | 652085 | 652085 | 652085 | -0.03 | 9.7E-01 | 0 |
| Hemoglobin A1C | 9 | 645613 | 900685 | 505020 | -0.03 | 9.8E-01 | 0 |
| Left pars triangularis | 1 | 669333 | 669333 | 669333 | -0.03 | 9.8E-01 | 0 |
| Granulocyte percentage of myeloid white cells (two-way meta) | 97 | 667925 | 1115185 | 93711 | -0.03 | 9.8E-01 | 0 |
| Body of corpus callosum axial diusivities | 3 | 647943 | 1047060 | 407927 | -0.02 | 9.8E-01 | 0 |
| Tumor necrosis factor receptor II (at exam 7, GEE, adjusted for age and sex) | 1 | 673465 | 673465 | 673465 | -0.02 | 9.8E-01 | 0 |
| SNRPF - Small nuclear ribonucleoprotein F | 1 | 678693 | 678693 | 678693 | -0.02 | 9.9E-01 | 0 |
| Non-albumin protein | 39 | 667463 | 1067198 | 411438 | -0.01 | 9.9E-01 | 0 |
| Interleukin-12p70 | 3 | 657310 | 764339 | 383500 | -0.01 | 9.9E-01 | 0 |
| Shaft average buckling ratio (GEE, male, adjussted for age) | 3 | 651995 | 854344 | 390326 | -0.01 | 9.9E-01 | 0 |
| SERPINA1 - Alpha-1-antitrypsin | 1 | 675858 | 675858 | 675858 | -0.01 | 9.9E-01 | 0 |
| CTSA - Lysosomal protective protein | 1 | 679143 | 679143 | 679143 | -0.01 | 9.9E-01 | 0 |
| CD36 - Platelet glycoprotein 4 | 1 | 682768 | 682768 | 682768 | -0.01 | 9.9E-01 | 0 |
| Lipid::Essential fatty acid::X-12990--docosapentaenoic acid (n6-DPA) | 1 | 667463 | 667463 | 667463 | -0.01 | 9.9E-01 | 0 |
| Peptide::Fibrinogen cleavage peptide::ADpSGEGDFXAEGGGVR* | 1 | 682768 | 682768 | 682768 | -0.01 | 9.9E-01 | 0 |
| Total lipids in medium VLDL | 8 | 651848 | 860042 | 156479 | -0.01 | 1.0E+00 | 0 |
| Type of accommodation lived in: A house or bungalow | 2 | 673859 | 991085 | 356634 | 0.00 | 1.0E+00 | 0 |
| Gamma-glutamyl transferase | 32 | 665893 | 1172208 | 259429 | 0.00 | 1.0E+00 | 0 |
| Total cholesterol / High-density lipoprotein cholesterol ratio (at exam 1, GEE, adjusted for age an | 1 | 671785 | 671785 | 671785 | 0.00 | 1.0E+00 | 0 |
| Glucose levels 2 h after an oral glucose challenge | 1 | 683278 | 683278 | 683278 | 0.00 | 1.0E+00 | 0 |
| Lipid::Inositol metabolism::scyllo-inositol | 1 | 680895 | 680895 | 680895 | 0.00 | 1.0E+00 | 0 |
| Body Mass Index (male) | 363 | 672625 | 1140566 | 176507 | 0.00 | 1.0E+00 | 0 |
| APOE - Apolipoprotein E (isoform E2) | 1 | 678693 | 678693 | 678693 | 0.00 | 1.0E+00 | 0 |
| TIE1 - Tyrosine-protein kinase receptor Tie-1, soluble | 1 | 688525 | 688525 | 688525 | 0.00 | 1.0E+00 | 0 |

| | | | | | | | |
|---|---|---|---|---|---|---|---|
| IDS - Iduronate 2-sulfatase | 1 | 679143 | 679143 | 679143 | 0.00 | 1.0E+00 | 0 |
| Non-cancer illness code, self-reported: osteoarthritis | 3 | 654168 | 848608 | 343179 | 0.00 | 1.0E+00 | 0 |
| ICAM2 - Intercellular adhesion molecule 2 | 1 | 682768 | 682768 | 682768 | 0.01 | 1.0E+00 | 0 |
| CD200 - OX-2 membrane glycoprotein | 1 | 682768 | 682768 | 682768 | 0.01 | 1.0E+00 | 0 |
| ::::X-11441 | 1 | 680218 | 680218 | 680218 | 0.01 | 1.0E+00 | 0 |
| Salicylic acid and derivatives | 6 | 658126 | 995141 | 413272 | 0.01 | 1.0E+00 | 0 |
| Total-body less head BMD | 4 | 660009 | 985103 | 324646 | 0.01 | 9.9E-01 | 0 |
| CHST15 - Carbohydrate sulfotransferase 15 | 1 | 682768 | 682768 | 682768 | 0.01 | 9.9E-01 | 0 |
| SELE - E-Selectin | 1 | 682768 | 682768 | 682768 | 0.01 | 9.9E-01 | 0 |
| Basophil percentage of granulocytes (three-way meta) | 38 | 669313 | 1004079 | 126006 | 0.01 | 9.9E-01 | 0 |
| CTSZ - Cathepsin Z | 1 | 679143 | 679143 | 679143 | 0.01 | 9.9E-01 | 0 |
| ::::X-11442 | 1 | 680218 | 680218 | 680218 | 0.01 | 9.9E-01 | 0 |
| INSR - Insulin receptor | 1 | 682768 | 682768 | 682768 | 0.01 | 9.9E-01 | 0 |
| Lipid::Fatty acid metabolism (also BCAA metabolism)::butyrylcarnitine | 4 | 661675 | 1076433 | 267493 | 0.02 | 9.9E-01 | 0 |
| ::::X-12627 | 1 | 690760 | 690760 | 690760 | 0.02 | 9.9E-01 | 0 |
| Eosinophil percentage of granulocytes (three-way meta) | 114 | 673229 | 1086919 | 84834 | 0.02 | 9.9E-01 | 0 |
| Lipid::Long chain fatty acid::pentadecanoate (15:0) | 1 | 694215 | 694215 | 694215 | 0.02 | 9.9E-01 | 0 |
| ENG - Endoglin | 1 | 682768 | 682768 | 682768 | 0.02 | 9.8E-01 | 0 |
| Serum calcium (at exam 2, GEE, adjusted for age and sex) | 1 | 693178 | 693178 | 693178 | 0.02 | 9.8E-01 | 0 |
| Hand grip (at exam 7, GEE) | 2 | 691033 | 736609 | 645456 | 0.02 | 9.8E-01 | 0 |
| Had other major operations | 1 | 693428 | 693428 | 693428 | 0.02 | 9.8E-01 | 0 |
| Hand grip (at offspring exam 7 cohort exam 27, GEE) | 2 | 691033 | 736609 | 645456 | 0.02 | 9.8E-01 | 0 |
| Uncinate fasciculus axial diusivities | 1 | 693060 | 693060 | 693060 | 0.02 | 9.8E-01 | 0 |
| Citrate | 4 | 666720 | 896784 | 548718 | 0.02 | 9.8E-01 | 0 |
| Posterior thalamic radiation (include optic radiation) fractional anisotropy | 1 | 695505 | 695505 | 695505 | 0.02 | 9.8E-01 | 0 |
| Interleukin-10 | 2 | 694316 | 712819 | 675813 | 0.02 | 9.8E-01 | 0 |
| Inferior fronto-occipital fasciculus radial diusivities | 1 | 693060 | 693060 | 693060 | 0.03 | 9.8E-01 | 0 |
| ::::X-11799 | 1 | 691853 | 691853 | 691853 | 0.03 | 9.8E-01 | 0 |
| Intermediate-density lipoprotein by NMR (at exam 4, GEE, adjusted for multivariable) | 12 | 662720 | 987299 | 63628 | 0.03 | 9.8E-01 | 0 |
| Age of initiation of regular smoking | 5 | 662633 | 1188550 | 465385 | 0.03 | 9.7E-01 | 0 |
| Right postcentral | 3 | 669333 | 909503 | 351999 | 0.03 | 9.7E-01 | 0 |
| Daytime sleepiness / dozing | 1 | 701585 | 701585 | 701585 | 0.04 | 9.7E-01 | 0 |
| ::::X-02269 | 3 | 671508 | 1023313 | 547845 | 0.04 | 9.7E-01 | 0 |
| Lymphocyte percentage of white cells (three-way meta) | 96 | 678575 | 1135889 | 95754 | 0.04 | 9.7E-01 | 0 |
| Right pars triangularis | 5 | 669333 | 677188 | 59909 | 0.04 | 9.7E-01 | 0 |
| Uncinate fasciculus mean diusivities | 1 | 703043 | 703043 | 703043 | 0.04 | 9.6E-01 | 0 |
| Phospholipids in medium VLDL | 10 | 666598 | 1093759 | 253186 | 0.05 | 9.6E-01 | 0 |
| Cognitive performance | 153 | 676120 | 1082510 | 92619 | 0.05 | 9.6E-01 | 0 |
| Reason for glasses/contact lenses: For just reading/near work as you are getting older (called 'pr | 3 | 680180 | 840683 | 363239 | 0.05 | 9.6E-01 | 0 |
| Uncinate fasciculus radial diusivities | 1 | 703043 | 703043 | 703043 | 0.05 | 9.6E-01 | 0 |
| Subcutaneous adipose tissue volume | 1 | 712045 | 712045 | 712045 | 0.05 | 9.6E-01 | 0 |
| Basophil percentage of white cells (three-way meta) | 44 | 677086 | 1033089 | 124297 | 0.05 | 9.6E-01 | 0 |
| Geographic atrophy | 3 | 678693 | 758206 | 552133 | 0.05 | 9.6E-01 | 0 |

| | | | | | | | |
|---|---|---|---|---|---|---|---|
| MMP8 - Neutrophil collagenase | 3 | 677845 | 730991 | 567746 | 0.05 | 9.6E-01 | 0 |
| Left supramarginal | 5 | 669333 | 690300 | 66903 | 0.05 | 9.6E-01 | 0 |
| PLAUR - Urokinase plasminogen activator surface receptor | 1 | 710330 | 710330 | 710330 | 0.06 | 9.6E-01 | 0 |
| Basophil percentage of white cells (two-way meta) | 29 | 676158 | 1020317 | 126312 | 0.06 | 9.6E-01 | 0 |
| White blood cells | 7 | 673745 | 920073 | 480588 | 0.06 | 9.5E-01 | 0 |
| Morningness | 117 | 678165 | 1160463 | 120689 | 0.06 | 9.5E-01 | 0 |
| Weight (female) | 10 | 675399 | 1111026 | 315026 | 0.06 | 9.5E-01 | 0 |
| Thyroid-stimulating hormone (male) | 21 | 676045 | 919705 | 602028 | 0.07 | 9.5E-01 | 0 |
| Neutrophil count (three-way meta) | 95 | 678315 | 1169459 | 104621 | 0.07 | 9.5E-01 | 0 |
| FLT4 - Vascular endothelial growth factor receptor 3 | 3 | 688525 | 775538 | 685646 | 0.07 | 9.4E-01 | 0 |
| Lipid::Monoacylglycerol::1-stearoylglycerol (1-monostearin) | 1 | 712063 | 712063 | 712063 | 0.07 | 9.4E-01 | 0 |
| Interleukin-17 | 1 | 725510 | 725510 | 725510 | 0.07 | 9.4E-01 | 0 |
| Childhood obesity | 6 | 673271 | 1101319 | 319310 | 0.07 | 9.4E-01 | 0 |
| IL17RD - Interleukin-17 receptor D | 1 | 717630 | 717630 | 717630 | 0.07 | 9.4E-01 | 0 |
| Monocyte count | 27 | 680480 | 1095929 | 217991 | 0.07 | 9.4E-01 | 0 |
| Amino acid::Tryptophan metabolism::indolelactate | 1 | 723603 | 723603 | 723603 | 0.07 | 9.4E-01 | 0 |
| Lipid::Lysolipid::1-eicosadienoylglycerophosphocholine* | 1 | 717513 | 717513 | 717513 | 0.07 | 9.4E-01 | 0 |
| Right ventral DC | 6 | 679235 | 1226819 | 77010 | 0.08 | 9.4E-01 | 0 |
| CSF3 - Granulocyte colony-stimulating factor | 1 | 730453 | 730453 | 730453 | 0.08 | 9.4E-01 | 0 |
| Epworth Sleepiness Scale (GEE, adjusted for age, sex, BMI and usual sleep duration) | 3 | 695350 | 856394 | 672824 | 0.08 | 9.4E-01 | 0 |
| Glutamine | 6 | 678170 | 1395566 | 58720 | 0.08 | 9.4E-01 | 0 |
| High cholesterol lipoprotein | 17 | 675745 | 1322198 | 53854 | 0.08 | 9.4E-01 | 0 |
| Estimated glomerular filtration rate based on serum creatinine (non-diabetic) | 61 | 680143 | 1000742 | 147526 | 0.08 | 9.4E-01 | 0 |
| Monocyte count (two-way meta) | 113 | 680480 | 1150310 | 96751 | 0.08 | 9.3E-01 | 0 |
| Inferior fronto-occipital fasciculus mode of anisotropy | 2 | 710418 | 818784 | 602051 | 0.08 | 9.3E-01 | 0 |
| Zinc sulfate turbidity test | 2 | 713658 | 1014365 | 412950 | 0.08 | 9.3E-01 | 0 |
| Basophil count (two-way meta) | 41 | 680480 | 1065250 | 84667 | 0.08 | 9.3E-01 | 0 |
| Remnant lipoprotein cholesterol (at exam 4, GEE) | 2 | 714651 | 834193 | 595109 | 0.08 | 9.3E-01 | 0 |
| Frequency of friend/family visits | 10 | 677620 | 1352924 | 135529 | 0.09 | 9.3E-01 | 0 |
| Loneliness, isolation (LONE) | 4 | 690156 | 1055344 | 442737 | 0.09 | 9.3E-01 | 0 |
| Family relationship satisfaction | 1 | 735878 | 735878 | 735878 | 0.09 | 9.3E-01 | 0 |
| Left posterior cingulate | 1 | 727988 | 727988 | 727988 | 0.09 | 9.3E-01 | 0 |
| Epworth Sleepiness Scale (GEE, adjusted for age, sex and BMI) | 3 | 695350 | 856394 | 672824 | 0.09 | 9.3E-01 | 0 |
| Lipid::Sphingolipid::palmitoyl sphingomyelin | 1 | 737510 | 737510 | 737510 | 0.09 | 9.3E-01 | 0 |
| Diagnoses - main ICD10: C44 Other and unspecified malignant neoplasm of skin | 26 | 682456 | 967618 | 200524 | 0.09 | 9.3E-01 | 0 |
| Serum levels of soluble E-selectin with T1D | 1 | 730695 | 730695 | 730695 | 0.09 | 9.3E-01 | 0 |
| Right cerebellum white matter | 13 | 684965 | 1317335 | 399468 | 0.10 | 9.2E-01 | 0 |
| Amino acid::Glutathione metabolism::5-oxoproline | 1 | 737328 | 737328 | 737328 | 0.10 | 9.2E-01 | 0 |
| Biologic age by osseographic scoring system (GEE, adjusted for multivariable) | 1 | 731040 | 731040 | 731040 | 0.10 | 9.2E-01 | 0 |
| Total cholesterol (at exam 6, GEE, adjusted for age and sex) | 3 | 702050 | 989013 | 648325 | 0.10 | 9.2E-01 | 0 |
| IL5RA - Interleukin-5 receptor subunit alpha | 1 | 736593 | 736593 | 736593 | 0.10 | 9.2E-01 | 0 |
| Treatment/medication code: simvastatin | 20 | 683355 | 873274 | 215978 | 0.10 | 9.2E-01 | 0 |
| Advanced adenoma | 1 | 730130 | 730130 | 730130 | 0.10 | 9.2E-01 | 0 |

| | | | | | | | |
|---|---|---|---|---|---|---|---|
| Chronic kidney disease (at exam 7, GEE, adjusted for age and sex) | 33 | 686600 | 1054995 | 384010 | 0.10 | 9.2E-01 | 0 |
| Bust size | 2 | 721569 | 763804 | 679333 | 0.11 | 9.2E-01 | 0 |
| LCMT1 - Leucine carboxyl methyltransferase 1 | 2 | 727061 | 1052119 | 402003 | 0.11 | 9.1E-01 | 0 |
| MASP1 - Mannan-binding lectin serine protease 1 | 1 | 739498 | 739498 | 739498 | 0.11 | 9.1E-01 | 0 |
| Cofactors and vitamins::Ascorbate and aldarate metabolism::X-11593--O-methylascorbate* | 5 | 694273 | 810723 | 646865 | 0.11 | 9.1E-01 | 0 |
| Human-virus - gp41 C34 peptide, HIV | 1 | 740338 | 740338 | 740338 | 0.11 | 9.1E-01 | 0 |
| ::::X-11470 | 1 | 732723 | 732723 | 732723 | 0.11 | 9.1E-01 | 0 |
| Maximum carotid artery stenosis (at exam 6, GEE, adjusted for age and sex) | 2 | 724583 | 908731 | 540434 | 0.12 | 9.1E-01 | 0 |
| VEGFA - Vascular endothelial growth factor A, isoform 121 | 1 | 742125 | 742125 | 742125 | 0.12 | 9.1E-01 | 0 |
| LYZ - Lysozyme C | 1 | 745753 | 745753 | 745753 | 0.12 | 9.0E-01 | 0 |
| Leprosy | 2 | 730364 | 870164 | 590563 | 0.12 | 9.0E-01 | 0 |
| Types of transport used (excluding work): Cycle | 1 | 746725 | 746725 | 746725 | 0.12 | 9.0E-01 | 0 |
| QT interval (male, cohort exam 11 offspring exam 1, GEE, adjusted for age nad RR) | 3 | 711275 | 844996 | 549950 | 0.13 | 9.0E-01 | 0 |
| MMP1 - Interstitial collagenase | 1 | 752188 | 752188 | 752188 | 0.13 | 9.0E-01 | 0 |
| Spine Neck BMD (GEE, female, adjusted for multivariable) | 1 | 746478 | 746478 | 746478 | 0.13 | 9.0E-01 | 0 |
| Lipid::Sterol, Steroid::dehydroisoandrosterone sulfate (DHEA-S) | 2 | 729131 | 973597 | 484665 | 0.13 | 9.0E-01 | 0 |
| Lipid::Sterol, Steroid::4-androsten-3beta,17beta-diol disulfate 1* | 2 | 729131 | 973597 | 484665 | 0.13 | 9.0E-01 | 0 |
| Platelet aggregation to collagen (at exam 5, GEE, adjusted for age and sex) | 1 | 754025 | 754025 | 754025 | 0.13 | 9.0E-01 | 0 |
| Superior corona radiata fractional anisotropy | 4 | 697195 | 760934 | 523477 | 0.13 | 9.0E-01 | 0 |
| Moderate intesity | 1 | 760235 | 760235 | 760235 | 0.13 | 9.0E-01 | 0 |
| Lipid::Carnitine metabolism::palmitoylcarnitine | 1 | 758448 | 758448 | 758448 | 0.13 | 9.0E-01 | 0 |
| Tumor necrosis factor alpha (at exam 7, GEE, adjusted for multivariable) | 4 | 696709 | 742702 | 550973 | 0.13 | 9.0E-01 | 0 |
| Genu of corpus callosum axial diusivities | 3 | 707440 | 1132926 | 606061 | 0.13 | 8.9E-01 | 0 |
| Amino acid::Valine, leucine and isoleucine metabolism::isobutyrylcarnitine | 2 | 731771 | 992866 | 470677 | 0.14 | 8.9E-01 | 0 |
| XCL1 - Lymphotactin | 1 | 754380 | 754380 | 754380 | 0.14 | 8.9E-01 | 0 |
| Platelet aggregation to collagen (at exam 5, GEE, adjusted for multivariable) | 1 | 754025 | 754025 | 754025 | 0.14 | 8.9E-01 | 0 |
| Retrolenticular part of internal capsule fractional anisotropy | 4 | 699931 | 1028419 | 362049 | 0.14 | 8.9E-01 | 0 |
| Total cholesterol (at exam 4, GEE, adjusted for age and sex) | 2 | 737199 | 811613 | 662784 | 0.14 | 8.9E-01 | 0 |
| Fasting plasma glucose (at exam 7, GEE, adjusted for age, sex and BMI) | 5 | 707010 | 1081133 | 658788 | 0.14 | 8.9E-01 | 0 |
| Lipid::Carnitine metabolism::oleoylcarnitine | 1 | 758448 | 758448 | 758448 | 0.14 | 8.9E-01 | 0 |
| Thin vs extreme obese | 2 | 737646 | 830016 | 645277 | 0.14 | 8.9E-01 | 0 |
| CH2 groups to double bonds ratio | 4 | 705436 | 922774 | 496423 | 0.15 | 8.8E-01 | 0 |
| High-density-lipoprotein cholesterol | 29 | 694580 | 1052688 | 103741 | 0.15 | 8.8E-01 | 0 |
| Femoral Neck BMD (GEE, female, adjusted for multivariable) | 2 | 736355 | 866749 | 605961 | 0.16 | 8.8E-01 | 0 |
| Lymphocyte count | 10 | 698776 | 1347217 | 201352 | 0.16 | 8.8E-01 | 0 |
| Cholesterol esters in medium HDL | 6 | 702551 | 871428 | 199109 | 0.16 | 8.7E-01 | 0 |
| Symbol digit substitution test - Number of symbol digit matches attempted | 6 | 705734 | 1240339 | 455333 | 0.16 | 8.7E-01 | 0 |
| Antidepressants | 1 | 769455 | 769455 | 769455 | 0.16 | 8.7E-01 | 0 |
| SERPINA4 - Kallistatin | 1 | 775320 | 775320 | 775320 | 0.17 | 8.7E-01 | 0 |
| Posterior cortical atrophy | 3 | 719763 | 804408 | 387647 | 0.17 | 8.7E-01 | 0 |
| CCL15 - C-C motif chemokine 15 | 1 | 770930 | 770930 | 770930 | 0.17 | 8.6E-01 | 0 |
| Spread type: Butter/spreadable butter | 8 | 706016 | 1029411 | 349474 | 0.17 | 8.6E-01 | 0 |
| Superior longitudinal fasciculus mean diusivities | 3 | 729058 | 1076166 | 516526 | 0.17 | 8.6E-01 | 0 |

| | | | | | | | |
|---|---|---|---|---|---|---|---|
| Right amygdala | 6 | 709455 | 1023940 | 394565 | 0.18 | 8.6E-01 | 0 |
| Heart rate recovery at 20 secnds | 15 | 705170 | 1033409 | 198798 | 0.18 | 8.6E-01 | 0 |
| Right precuneus | 2 | 748906 | 1094522 | 403291 | 0.18 | 8.6E-01 | 0 |
| First PC of the four riskly behaviours | 64 | 694644 | 1229573 | 198975 | 0.18 | 8.6E-01 | 0 |
| Snoring | 34 | 698791 | 1242741 | 111614 | 0.18 | 8.6E-01 | 0 |
| Concentration of small VLDL particles | 12 | 702551 | 949047 | 421839 | 0.18 | 8.6E-01 | 0 |
| Medication for pain relief, constipation, heartburn: Aspirin | 4 | 719566 | 1061144 | 300735 | 0.18 | 8.6E-01 | 0 |
| Coffee cups per day | 1 | 780430 | 780430 | 780430 | 0.18 | 8.5E-01 | 0 |
| Impedance measures - Impedance of leg (left) | 447 | 682503 | 1155285 | 124853 | 0.19 | 8.5E-01 | 0 |
| Hemoglobin A1c | 21 | 705053 | 1112253 | 417280 | 0.20 | 8.4E-01 | 0 |
| Mean coronary artery calcification - Agatston score (at exam 7, GEE, adjusted for multivariable) | 1 | 781405 | 781405 | 781405 | 0.20 | 8.4E-01 | 0 |
| Amino acid::Phenylalanine & tyrosine metabolism::3-(4-hydroxyphenyl)lactate | 1 | 786553 | 786553 | 786553 | 0.20 | 8.4E-01 | 0 |
| Low-density lipoprotein cholesterol (at exam 2, GEE, adjusted for age and sex) | 1 | 781405 | 781405 | 781405 | 0.20 | 8.4E-01 | 0 |
| WIF1 - Wnt inhibitory factor 1 | 1 | 793583 | 793583 | 793583 | 0.20 | 8.4E-01 | 0 |
| ::::X-11412 | 1 | 797550 | 797550 | 797550 | 0.21 | 8.4E-01 | 0 |
| Mean thyroid stimulation hormone from exam 3 and 4 (GEE, adjusted for multivariable) | 2 | 759696 | 1017835 | 501556 | 0.21 | 8.3E-01 | 0 |
| ::::X-11792 | 2 | 755300 | 829280 | 681320 | 0.21 | 8.3E-01 | 0 |
| Right lateral orbitofrontal | 5 | 735818 | 1284313 | 695640 | 0.21 | 8.3E-01 | 0 |
| Blood clot, DVT, bronchitis, emphysema, asthma, rhinitis, eczema, allergy diagnosed by doctor: H | 111 | 690818 | 1120401 | 99927 | 0.21 | 8.3E-01 | 0 |
| Basophil percentage of granulocytes (two-way meta) | 28 | 703709 | 1128588 | 109064 | 0.22 | 8.3E-01 | 0 |
| Illnesses of mother: Breast cancer | 12 | 713113 | 1123981 | 72306 | 0.22 | 8.3E-01 | 0 |
| Cancer (diagnosed by doctor) | 5 | 729058 | 1116113 | 718645 | 0.22 | 8.2E-01 | 0 |
| Free thyroxine (FT4, male) | 7 | 719743 | 892615 | 514966 | 0.22 | 8.2E-01 | 0 |
| High-density lipoprotein cholesterol (at exam 4, GEE, adjusted for age and sex) | 2 | 768730 | 927589 | 609871 | 0.22 | 8.2E-01 | 0 |
| Right cerebellum exterior | 21 | 713013 | 1015942 | 37856 | 0.23 | 8.2E-01 | 0 |
| ::::X-18601 | 2 | 766418 | 937535 | 595300 | 0.23 | 8.2E-01 | 0 |
| Non-cancer illness code, self-reported: depression | 1 | 803498 | 803498 | 803498 | 0.23 | 8.1E-01 | 0 |
| HAVCR2 - Hepatitis A virus cellular receptor 2 | 1 | 802273 | 802273 | 802273 | 0.24 | 8.1E-01 | 0 |
| High-density lipoprotein 3 cholesterol (at exam 4, GEE, adjusted for age and sex) | 2 | 768730 | 927589 | 609871 | 0.24 | 8.1E-01 | 0 |
| Relative age voice broke (male) | 26 | 709688 | 1009222 | 141234 | 0.24 | 8.1E-01 | 0 |
| Current employment status: Doing unpaid or voluntary work | 3 | 760953 | 960884 | 418647 | 0.24 | 8.1E-01 | 0 |
| Amino acid::Creatine metabolism::creatine | 1 | 804745 | 804745 | 804745 | 0.24 | 8.1E-01 | 0 |
| Number of live births (female) | 7 | 734533 | 1061604 | 98554 | 0.24 | 8.1E-01 | 0 |
| Amino acid::Glycine, serine and threonine metabolism::glycine | 1 | 804745 | 804745 | 804745 | 0.24 | 8.1E-01 | 0 |
| Platelet distribution width (two-way meta) | 122 | 694173 | 1116600 | 144377 | 0.25 | 8.1E-01 | 0 |
| Forearm BMD | 3 | 760258 | 1174644 | 578291 | 0.25 | 8.1E-01 | 0 |
| Lipid::Monoacylglycerol::1-oleoylglycerol (1-monoolein) | 1 | 818555 | 818555 | 818555 | 0.25 | 8.0E-01 | 0 |
| CNDP1 - Beta-Ala-His dipeptidase | 1 | 811238 | 811238 | 811238 | 0.25 | 8.0E-01 | 0 |
| ::::X-11452 | 1 | 822285 | 822285 | 822285 | 0.25 | 8.0E-01 | 0 |
| Splenium of corpus callosum fractional anisotropy | 5 | 740255 | 970290 | 695505 | 0.26 | 8.0E-01 | 0 |
| Amino acid::Lysine metabolism::glutaroyl carnitine | 8 | 726981 | 1094302 | 443685 | 0.26 | 8.0E-01 | 0 |
| Eosinophil count (two-way meta) | 106 | 697781 | 1141567 | 130407 | 0.26 | 8.0E-01 | 0 |
| Ever had known person concerned about, or recommended reduction of alcohol consumption | 2 | 774169 | 838752 | 709586 | 0.26 | 7.9E-01 | 0 |

| | | | | | | | |
|---|---|---|---|---|---|---|---|
| Mineral and other dietary supplements: Calcium | 4 | 749432 | 1492680 | 40017 | 0.26 | 7.9E-01 | 0 |
| Albuminuria | 31 | 714893 | 1137285 | 249761 | 0.27 | 7.9E-01 | 0 |
| Total cholesterol / High-density lipoprotein cholesterol ratio (at exam 2, GEE, adjusted for age an | 3 | 766583 | 773994 | 513825 | 0.27 | 7.9E-01 | 0 |
| Total cholesterol (at exam 7, GEE, adjusted for age and sex) | 1 | 818283 | 818283 | 818283 | 0.27 | 7.9E-01 | 0 |
| CNTN4 - Contactin-4 | 1 | 823563 | 823563 | 823563 | 0.27 | 7.9E-01 | 0 |
| Age high blood pressure diagnosed | 9 | 729583 | 1372948 | 84496 | 0.27 | 7.9E-01 | 0 |
| Mean abdominal aortic calcification - Agatston score (at exam 7, GEE, adjusted for age and sex) | 2 | 778343 | 791410 | 765275 | 0.27 | 7.9E-01 | 0 |
| Ever unenthusiastic/disinterested for a whole week | 2 | 782587 | 1140742 | 424432 | 0.27 | 7.9E-01 | 0 |
| ::::X-11327 | 2 | 787613 | 1007905 | 567320 | 0.27 | 7.8E-01 | 0 |
| Lipid::Eicosanoid::X-12441--12-hydroxyeicosatetraenoate (12-HETE) | 1 | 831750 | 831750 | 831750 | 0.28 | 7.8E-01 | 0 |
| Inflammatory Bowel Disease | 132 | 697886 | 1126955 | 148760 | 0.29 | 7.8E-01 | 0 |
| ESD - S-formylglutathione hydrolase | 1 | 846518 | 846517 | 846517 | 0.29 | 7.7E-01 | 0 |
| Diagnoses - main ICD10: K57 Diverticular disease of intestine | 12 | 732865 | 1004836 | 394734 | 0.29 | 7.7E-01 | 0 |
| Frequency of tenseness / restlessness in last 2 weeks | 8 | 739929 | 1016108 | 310178 | 0.29 | 7.7E-01 | 0 |
| WFIKKN2 - WAP, Kazal, immunoglobulin, Kunitz and NTR domain-containing protein 2 | 1 | 840440 | 840440 | 840440 | 0.29 | 7.7E-01 | 0 |
| Cancer register - Behaviour of cancer tumour: Carcinoma in situ | 1 | 840790 | 840790 | 840790 | 0.29 | 7.7E-01 | 0 |
| Mean corpuscular hemoglobin (two-way meta) | 137 | 697765 | 1216680 | 220254 | 0.29 | 7.7E-01 | 0 |
| Disialylation | 1 | 835425 | 835425 | 835425 | 0.29 | 7.7E-01 | 0 |
| Obesity class 1 | 16 | 730111 | 1262095 | 383953 | 0.30 | 7.7E-01 | 0 |
| Heart rate variability (SDNN) | 6 | 751738 | 1298050 | 161926 | 0.30 | 7.7E-01 | 0 |
| Total bilirubin | 8 | 738733 | 1342429 | 91881 | 0.30 | 7.7E-01 | 0 |
| Cingulum (hippocampus) fractional anisotropy | 1 | 840440 | 840440 | 840440 | 0.30 | 7.6E-01 | 0 |
| Lipid::Fatty acid metabolism (also BCAA metabolism)::propionylcarnitine | 4 | 758270 | 1254596 | 388469 | 0.30 | 7.6E-01 | 0 |
| Uncinate fasciculus fractional anisotropy | 2 | 790013 | 994476 | 585549 | 0.30 | 7.6E-01 | 0 |
| Hyperthyroidism | 10 | 740721 | 1123445 | 424717 | 0.30 | 7.6E-01 | 0 |
| Sleep sedentary (conditioning sex and BMI) | 5 | 759793 | 1561180 | 701760 | 0.30 | 7.6E-01 | 0 |
| Drugs affecting bone structure and mineralization | 10 | 739629 | 1139094 | 433414 | 0.31 | 7.6E-01 | 0 |
| IL17RB - interleukin-17 receptor B | 1 | 843100 | 843100 | 843100 | 0.31 | 7.6E-01 | 0 |
| Amino acid::Phenylalanine & tyrosine metabolism::phenyllactate (PLA) | 1 | 841233 | 841233 | 841233 | 0.31 | 7.6E-01 | 0 |
| HbA1c | 54 | 714065 | 1209588 | 383161 | 0.31 | 7.6E-01 | 0 |
| Intermediate high-density lipoprotein by NMR (at exam 4, GEE, adjusted for age and sex) | 1 | 844268 | 844267 | 844267 | 0.31 | 7.6E-01 | 0 |
| Diagnoses - main ICD10: R10 Abdominal and pelvic pain | 1 | 845510 | 845510 | 845510 | 0.31 | 7.6E-01 | 0 |
| Arms-arm fat ratio (female) | 118 | 700878 | 1034723 | 102815 | 0.31 | 7.6E-01 | 0 |
| CST1 - Cystatin-SN | 2 | 803341 | 1158676 | 448007 | 0.31 | 7.5E-01 | 0 |
| Right hippocampus | 12 | 737163 | 905236 | 433041 | 0.32 | 7.5E-01 | 0 |
| BCL2A1 - Bcl-2-related protein A1 | 1 | 851908 | 851908 | 851908 | 0.32 | 7.5E-01 | 0 |
| Left inferior lateral ventricle | 4 | 766684 | 984178 | 446730 | 0.32 | 7.5E-01 | 0 |
| CST2 - Cystatin-SA | 2 | 803341 | 1158676 | 448007 | 0.33 | 7.4E-01 | 0 |
| Periodontal complex trait 3 - Aa trait | 2 | 798903 | 961373 | 636433 | 0.33 | 7.4E-01 | 0 |
| Creatine kinase | 30 | 723366 | 1011169 | 66381 | 0.33 | 7.4E-01 | 0 |
| Retrolenticular part of internal capsule mean diusivities | 2 | 801855 | 855030 | 748680 | 0.33 | 7.4E-01 | 0 |
| Mineral and other dietary supplements: Fish oil (including cod liver oil) | 6 | 753070 | 1034334 | 179905 | 0.33 | 7.4E-01 | 0 |
| Lipid::Sterol, Steroid::4-androsten-3beta,17beta-diol disulfate 2* | 2 | 802881 | 1179008 | 426753 | 0.33 | 7.4E-01 | 0 |

| | | | | | | | |
|---|---|---|---|---|---|---|---|
| Atrial fibrillation | 103 | 704448 | 1122404 | 342304 | 0.33 | 7.4E-01 | 0 |
| HRG - Histidine-rich glycoprotein | 1 | 859163 | 859163 | 859163 | 0.34 | 7.4E-01 | 0 |
| Lipid::Sterol, Steroid::cortisone | 1 | 870475 | 870475 | 870475 | 0.34 | 7.4E-01 | 0 |
| FETUB - Fetuin-B | 1 | 859163 | 859163 | 859163 | 0.34 | 7.4E-01 | 0 |
| Relative wall thickness | 1 | 853988 | 853988 | 853988 | 0.34 | 7.3E-01 | 0 |
| ETHE1 - Persulfide dioxygenase ETHE1, mitochondrial | 1 | 859163 | 859163 | 859163 | 0.34 | 7.3E-01 | 0 |
| Nap during day | 75 | 711078 | 1105469 | 130951 | 0.34 | 7.3E-01 | 0 |
| Free thyroxine (FT4, female) | 9 | 757165 | 821192 | 310785 | 0.34 | 7.3E-01 | 0 |
| ICAM5 - Intercellular adhesion molecule 5 | 1 | 863173 | 863173 | 863173 | 0.34 | 7.3E-01 | 0 |
| Heart rate variability (RMSSD) | 7 | 767923 | 1475635 | 326952 | 0.35 | 7.3E-01 | 0 |
| ICAM1 - Intercellular adhesion molecule 1 | 1 | 863173 | 863173 | 863173 | 0.35 | 7.3E-01 | 0 |
| Amino acid::Valine, leucine and isoleucine metabolism::hydroxyisovaleroyl carnitine | 2 | 809280 | 1142384 | 476177 | 0.35 | 7.3E-01 | 0 |
| Phospholipids in IDL | 12 | 746061 | 1154155 | 95473 | 0.35 | 7.3E-01 | 0 |
| AHSG - Alpha-2-HS-glycoprotein | 1 | 869150 | 869150 | 869150 | 0.35 | 7.3E-01 | 0 |
| Peptide::Dipeptide::X-14208--phenylalanylserine | 1 | 866553 | 866553 | 866553 | 0.35 | 7.3E-01 | 0 |
| Facial ageing | 55 | 718705 | 1052721 | 90395 | 0.35 | 7.2E-01 | 0 |
| Antihypertensives | 2 | 818203 | 848533 | 787873 | 0.36 | 7.2E-01 | 0 |
| N-glycosylation | 6 | 772176 | 1259329 | 502829 | 0.36 | 7.2E-01 | 0 |
| Visual scanning and motor speed (GEE, adjusted for multivariable) | 2 | 816872 | 1132989 | 500756 | 0.36 | 7.2E-01 | 0 |
| Inferior fronto-occipital fasciculus fractional anisotropy | 4 | 773926 | 961300 | 624258 | 0.36 | 7.2E-01 | 0 |
| Transport type for commuting to job workplace: Car/motor vehicle | 1 | 864035 | 864035 | 864035 | 0.37 | 7.1E-01 | 0 |
| Mean corpuscular hemoglobin | 65 | 714893 | 1101913 | 90714 | 0.37 | 7.1E-01 | 0 |
| Amino acid::Alanine and aspartate metabolism::asparagine | 3 | 809285 | 889586 | 803198 | 0.37 | 7.1E-01 | 0 |
| Fasting plasma glucose (at exam 7, GEE, adjusted for age and sex) | 12 | 751496 | 1085908 | 482089 | 0.37 | 7.1E-01 | 0 |
| Falls in the last year | 5 | 783183 | 893328 | 238323 | 0.37 | 7.1E-01 | 0 |
| High light scatter percentage of red cells (three-way meta) | 118 | 707310 | 1110043 | 119754 | 0.37 | 7.1E-01 | 0 |
| CPNE1 - Copine-1 | 1 | 876843 | 876842 | 876842 | 0.37 | 7.1E-01 | 0 |
| Alkaline phosphatase (at exam 2, GEE, adjusted for multivariable) | 2 | 816666 | 933526 | 699807 | 0.37 | 7.1E-01 | 0 |
| Vascular endothelial growth factor | 2 | 816755 | 1170288 | 463223 | 0.38 | 7.1E-01 | 0 |
| Total cholesterol in large VLDL | 11 | 765925 | 919409 | 116974 | 0.38 | 7.1E-01 | 0 |
| CD32 on mDC | 1 | 883748 | 883748 | 883748 | 0.38 | 7.1E-01 | 0 |
| Serum total cholesterol | 10 | 757501 | 1112504 | 143176 | 0.38 | 7.1E-01 | 0 |
| Male-specific factors - Hair/balding pattern: Pattern 4 | 145 | 703688 | 1127210 | 143498 | 0.38 | 7.0E-01 | 0 |
| Isoleucine | 2 | 824168 | 1228768 | 419568 | 0.38 | 7.0E-01 | 0 |
| Average weekly champagne plus white wine intake | 11 | 764545 | 1211046 | 454174 | 0.38 | 7.0E-01 | 0 |
| CAPG - Macrophage-capping protein | 1 | 891645 | 891645 | 891645 | 0.39 | 7.0E-01 | 0 |
| TIMP3 - Metalloproteinase inhibitor 3 | 1 | 888365 | 888365 | 888365 | 0.39 | 7.0E-01 | 0 |
| Illnesses of father: Alzheimer's disease/dementia | 1 | 889053 | 889053 | 889053 | 0.39 | 7.0E-01 | 0 |
| Splenium of corpus callosum radial diusivities | 8 | 768704 | 1355371 | 553884 | 0.39 | 7.0E-01 | 0 |
| ::::X-10510 | 3 | 816268 | 938858 | 810763 | 0.39 | 7.0E-01 | 0 |
| LGALS3BP - Galectin-3-binding protein | 1 | 879690 | 879690 | 879690 | 0.39 | 7.0E-01 | 0 |
| CD4:%Treg(39+73+) | 1 | 882988 | 882988 | 882988 | 0.39 | 7.0E-01 | 0 |
| Amino acid::Tryptophan metabolism::tryptophan betaine | 2 | 826493 | 979168 | 673818 | 0.39 | 6.9E-01 | 0 |

| | | | | | | | |
|---|---|---|---|---|---|---|---|
| Total protein | 24 | 743341 | 963183 | 392371 | 0.39 | 6.9E-01 | 0 |
| Left cerebellum exterior | 13 | 763213 | 1305810 | 51303 | 0.40 | 6.9E-01 | 0 |
| Weight change compared with 1 year ago | 1 | 889053 | 889053 | 889053 | 0.40 | 6.9E-01 | 0 |
| Drugs for peptic ulcer and gastro-oesophageal reflux disease (GORD) | 5 | 795498 | 1178155 | 66035 | 0.40 | 6.9E-01 | 0 |
| Cannabis use | 4 | 787915 | 1245809 | 353908 | 0.40 | 6.9E-01 | 0 |
| Mean low-density lipoprotein cholesterol fromexam 1-7 (GEE, adjusted for multivariable) | 2 | 836973 | 961274 | 712671 | 0.40 | 6.9E-01 | 0 |
| Neck Section Modulus (GEE, adjusted for multivariable) | 1 | 896600 | 896600 | 896600 | 0.40 | 6.9E-01 | 0 |
| Body of corpus callosum fractional anisotropy | 3 | 819578 | 883061 | 561786 | 0.40 | 6.9E-01 | 0 |
| ::::X-11483 | 1 | 889695 | 889695 | 889695 | 0.40 | 6.9E-01 | 0 |
| ::::X-14625 | 1 | 892930 | 892930 | 892930 | 0.40 | 6.9E-01 | 0 |
| PGC cross disorder | 4 | 799226 | 1225639 | 300538 | 0.41 | 6.9E-01 | 0 |
| 25-Hydroxyvitamin D level | 20 | 747458 | 986733 | 293376 | 0.41 | 6.8E-01 | 0 |
| Antisocial behavior (female) | 2 | 830614 | 1216454 | 444773 | 0.41 | 6.8E-01 | 0 |
| MMSE score (at exam 7, GEE, adjusted for age) | 12 | 762891 | 1060904 | 241497 | 0.41 | 6.8E-01 | 0 |
| Mean total cholesterol from exam 1-7 (GEE, adjusted for multivariable) | 2 | 836973 | 961274 | 712671 | 0.41 | 6.8E-01 | 0 |
| Diagnoses - secondary ICD10: E11 Type 2 diabetes mellitus | 39 | 732540 | 1086393 | 299993 | 0.42 | 6.8E-01 | 0 |
| Genetic generalized epilepsy | 1 | 903535 | 903535 | 903535 | 0.42 | 6.8E-01 | 0 |
| Osteoarthritis of hip | 28 | 739114 | 1045994 | 187140 | 0.42 | 6.8E-01 | 0 |
| Superior corona radiata mean diusivities | 2 | 835703 | 905801 | 765604 | 0.42 | 6.8E-01 | 0 |
| Phospholipids in very small VLDL | 14 | 761860 | 1037454 | 170611 | 0.42 | 6.8E-01 | 0 |
| Digalactosylation | 6 | 786509 | 1178976 | 574811 | 0.42 | 6.7E-01 | 0 |
| Peptide::Dipeptide::X-14189--leucylalanine | 2 | 843853 | 924395 | 763310 | 0.42 | 6.7E-01 | 0 |
| Anxiety disorder (factor score) | 1 | 903160 | 903160 | 903160 | 0.43 | 6.7E-01 | 0 |
| LDL diameter | 5 | 805883 | 955163 | 395798 | 0.43 | 6.7E-01 | 0 |
| CST5 - Cystatin-D | 1 | 897198 | 897198 | 897198 | 0.43 | 6.7E-01 | 0 |
| Treatment/medication code: ramipril | 3 | 826673 | 1222518 | 610704 | 0.43 | 6.7E-01 | 0 |
| Posterior limb of internal capsule mean diusivities | 3 | 838300 | 1088341 | 628503 | 0.43 | 6.7E-01 | 0 |
| Superior corona radiata radial diusivities | 4 | 800633 | 912428 | 524465 | 0.43 | 6.7E-01 | 0 |
| CA6 - Carbonic anhydrase 6 | 1 | 910498 | 910498 | 910498 | 0.43 | 6.7E-01 | 0 |
| Visceral sdipose tissue volume (adjusted for BMI) | 4 | 799906 | 1192394 | 394419 | 0.43 | 6.7E-01 | 0 |
| Vitiligo | 30 | 740455 | 1176916 | 116180 | 0.43 | 6.7E-01 | 0 |
| High-density lipoprotein cholesterol (at exam 1, GEE, adjusted for age and sex) | 1 | 902585 | 902585 | 902585 | 0.43 | 6.7E-01 | 0 |
| Peptide::gamma-glutamyl::gamma-glutamylglutamine | 2 | 849843 | 1241605 | 458081 | 0.43 | 6.6E-01 | 0 |
| Number of sexual partners | 72 | 721611 | 1162502 | 188661 | 0.44 | 6.6E-01 | 0 |
| MAPKAPK2 - MAP kinase-activated protein kinase 2 | 2 | 852759 | 1017141 | 688377 | 0.44 | 6.6E-01 | 0 |
| Estimated glomerular filtration rate based on serum creatinine | 105 | 714693 | 1015240 | 271088 | 0.44 | 6.6E-01 | 0 |
| Alcohol intake frequency | 70 | 724966 | 1125026 | 129270 | 0.44 | 6.6E-01 | 0 |
| MMSE score (at exam 5, GEE, adjusted for age) | 11 | 778853 | 1040979 | 695891 | 0.44 | 6.6E-01 | 0 |
| Triglycerides (at exam 1, GEE, adjusted for multivariable) | 1 | 913278 | 913278 | 913278 | 0.44 | 6.6E-01 | 0 |
| HDL diameter | 8 | 786835 | 1110696 | 567870 | 0.44 | 6.6E-01 | 0 |
| Vitiligo (early onset) | 1 | 922435 | 922435 | 922435 | 0.44 | 6.6E-01 | 0 |
| APOB - Apolipoprotein B | 1 | 916168 | 916167 | 916167 | 0.45 | 6.6E-01 | 0 |
| Fasting glucose main effect | 16 | 762186 | 1391916 | 191450 | 0.45 | 6.6E-01 | 0 |

| | | | | | | | |
|---|---|---|---|---|---|---|---|
| Concentration of very large HDL particles | 7 | 797550 | 1177365 | 584506 | 0.45 | 6.5E-01 | 0 |
| POSTN - Periostin | 1 | 911180 | 911180 | 911180 | 0.45 | 6.5E-01 | 0 |
| Illnesses of father: Prostate cancer | 3 | 842773 | 1055566 | 566590 | 0.45 | 6.5E-01 | 0 |
| Cancer (GEE, adjusted for multivariable) | 1 | 920393 | 920392 | 920392 | 0.45 | 6.5E-01 | 0 |
| Lipid::Lysolipid::1-palmitoylglycerophosphoethanolamine | 1 | 925430 | 925430 | 925430 | 0.45 | 6.5E-01 | 0 |
| Positive affect (MA GWAMA) | 82 | 720200 | 1165619 | 175515 | 0.45 | 6.5E-01 | 0 |
| Platelet derived growth factor BB | 3 | 836653 | 1103279 | 483783 | 0.45 | 6.5E-01 | 0 |
| Hypertension | 1 | 925610 | 925610 | 925610 | 0.46 | 6.5E-01 | 0 |
| Genu of corpus callosum radial diusivities | 4 | 807954 | 1008323 | 699594 | 0.46 | 6.5E-01 | 0 |
| Cancer (GEE, adjusted for age and sex) | 1 | 920393 | 920392 | 920392 | 0.46 | 6.5E-01 | 0 |
| Sum neutrophil eosinophil count (three-way meta) | 102 | 718523 | 1201344 | 155431 | 0.46 | 6.5E-01 | 0 |
| GFRA1 - GDNF family receptor alpha-1 | 2 | 855804 | 1020434 | 691173 | 0.46 | 6.4E-01 | 0 |
| Diagnoses - main ICD10: I25 Chronic ischemic heart disease | 25 | 755635 | 1079455 | 453870 | 0.46 | 6.4E-01 | 0 |
| Femoral Neck Length (GEE, adjusted for multivariable) | 3 | 848208 | 928820 | 634066 | 0.46 | 6.4E-01 | 0 |
| Cingulum (hippocampus) mean diusivities | 1 | 925098 | 925098 | 925098 | 0.46 | 6.4E-01 | 0 |
| Diagnoses - secondary ICD10: J45 Asthma | 21 | 760445 | 1105033 | 303590 | 0.46 | 6.4E-01 | 0 |
| Ischemic stroke | 1 | 928078 | 928078 | 928078 | 0.46 | 6.4E-01 | 0 |
| Lipid::Lysolipid::1-oleoylglycerophosphoethanolamine | 1 | 925430 | 925430 | 925430 | 0.46 | 6.4E-01 | 0 |
| PCSK7 - Proprotein convertase subtilisin/kexin type 7 | 1 | 922308 | 922308 | 922308 | 0.47 | 6.4E-01 | 0 |
| Beta nerve growth factor | 1 | 928360 | 928360 | 928360 | 0.47 | 6.4E-01 | 0 |
| Epworth Sleepiness Scale (GEE, adjusted for age, sex, BMI, usual sleep duration and additional | 2 | 856394 | 936916 | 775872 | 0.47 | 6.4E-01 | 0 |
| Concentration of large VLDL particles | 8 | 792240 | 1008001 | 391314 | 0.47 | 6.4E-01 | 0 |
| Phenylalanine | 2 | 854445 | 913310 | 795580 | 0.47 | 6.4E-01 | 0 |
| Sum basophil neutrophil count (three-way meta) | 100 | 718523 | 1184096 | 110826 | 0.47 | 6.4E-01 | 0 |
| Total cholesterol (at exam 2, GEE, adjusted for age and sex) | 4 | 816515 | 998514 | 616188 | 0.47 | 6.4E-01 | 0 |
| IL25 - Interleukin-25 | 1 | 923305 | 923305 | 923305 | 0.47 | 6.4E-01 | 0 |
| Lipid::Carnitine metabolism::octanoylcarnitine | 3 | 852333 | 1008056 | 695191 | 0.47 | 6.4E-01 | 0 |
| Vitamin D plasma 25(OH)-D (at exam 6 or 7, GEE, adjusted for age and sex) | 1 | 928385 | 928385 | 928385 | 0.47 | 6.4E-01 | 0 |
| Worry/vulnerability factors | 13 | 782163 | 1343513 | 551685 | 0.48 | 6.3E-01 | 0 |
| Left ventricular internal dimension in diastole | 3 | 853988 | 1185196 | 800129 | 0.48 | 6.3E-01 | 0 |
| Total-body less head BMD and total body lean mass (bivariate meta-analysis) | 9 | 796800 | 1004520 | 666668 | 0.48 | 6.3E-01 | 0 |
| Age at menopause | 42 | 739396 | 1113224 | 399028 | 0.48 | 6.3E-01 | 0 |
| Diagnoses - secondary ICD10: K21 Gastro-esophageal reflux disease | 1 | 940400 | 940400 | 940400 | 0.48 | 6.3E-01 | 0 |
| AMN - Protein amnionless | 1 | 938475 | 938475 | 938475 | 0.48 | 6.3E-01 | 0 |
| ::::X-03094 | 5 | 818555 | 1317188 | 83239 | 0.48 | 6.3E-01 | 0 |
| Neck Section Modulus (GEE, female, adjusted for multivariable) | 1 | 940688 | 940688 | 940688 | 0.48 | 6.3E-01 | 0 |
| Illnesses of siblings: Heart disease | 3 | 851270 | 1019795 | 775553 | 0.48 | 6.3E-01 | 0 |
| Crohn's Disease | 142 | 712213 | 1132919 | 347002 | 0.48 | 6.3E-01 | 0 |
| Cingulum (cingulate gyrus) fractional anisotropy | 4 | 818978 | 942934 | 579343 | 0.48 | 6.3E-01 | 0 |
| Superior longitudinal fasciculus radial diusivities | 4 | 818631 | 1036972 | 622792 | 0.48 | 6.3E-01 | 0 |
| Energy::Krebs cycle::malate | 1 | 938865 | 938865 | 938865 | 0.49 | 6.3E-01 | 0 |
| Worry subcluster | 59 | 731810 | 1311698 | 112775 | 0.49 | 6.3E-01 | 0 |
| Anxiety/tension factors | 13 | 782163 | 1343513 | 551685 | 0.49 | 6.2E-01 | 0 |

| | | | | | | | |
|---|---|---|---|---|---|---|---|
| MMSE score (at exam 7, GEE, adjusted for multivariable) | 2 | 858548 | 1016168 | 700928 | 0.49 | 6.2E-01 | 0 |
| MAPKAPK3 - MAP kinase-activated protein kinase 3 | 1 | 950015 | 950015 | 950015 | 0.49 | 6.2E-01 | 0 |
| ANXA1 - Annexin A1 | 1 | 928565 | 928565 | 928565 | 0.49 | 6.2E-01 | 0 |
| Free cholesterol in small VLDL | 11 | 797550 | 919060 | 423210 | 0.49 | 6.2E-01 | 0 |
| Lipid::Sterol, Steroid::cholesterol | 1 | 946783 | 946783 | 946783 | 0.49 | 6.2E-01 | 0 |
| Neck Section Modulus (GEE, female, adjusted for age) | 1 | 940688 | 940688 | 940688 | 0.49 | 6.2E-01 | 0 |
| Chronic kidney disease (at exam 7, GEE, adjusted for multivariable) | 12 | 789218 | 1085388 | 477651 | 0.50 | 6.2E-01 | 0 |
| Left ventricular internal dimension in systole | 3 | 853988 | 970369 | 800129 | 0.50 | 6.2E-01 | 0 |
| Low cholesterol lipoprotein | 14 | 775933 | 1169944 | 439964 | 0.50 | 6.2E-01 | 0 |
| Free cholesterol in medium VLDL | 7 | 818555 | 1096754 | 297742 | 0.50 | 6.2E-01 | 0 |
| Heart rate recovery at 10 secnds | 16 | 776683 | 1143033 | 108092 | 0.50 | 6.2E-01 | 0 |
| HOMA-B | 4 | 826464 | 1353527 | 275448 | 0.50 | 6.2E-01 | 0 |
| PROC - Vitamin K-dependent protein C | 1 | 950673 | 950673 | 950673 | 0.50 | 6.2E-01 | 0 |
| ::::X-12844 | 4 | 824953 | 1259806 | 359058 | 0.50 | 6.2E-01 | 0 |
| Heart rate recovery at 30 secnds | 21 | 767923 | 904060 | 115768 | 0.51 | 6.1E-01 | 0 |
| Phospholipids in large VLDL | 7 | 818555 | 919409 | 314614 | 0.51 | 6.1E-01 | 0 |
| Anterior corona radiata fractional anisotropy | 7 | 819578 | 848623 | 377580 | 0.51 | 6.1E-01 | 0 |
| Total lipids in chylomicrons and extremely large VLDL | 7 | 818555 | 979404 | 327468 | 0.51 | 6.1E-01 | 0 |
| Central corneal thickness | 29 | 759835 | 1053798 | 422213 | 0.51 | 6.1E-01 | 0 |
| Adrenergics, inhalants | 40 | 751529 | 1001276 | 318350 | 0.52 | 6.0E-01 | 0 |
| Pericardial adipose tissue volume (adjusted for height and weight, male) | 1 | 958743 | 958742 | 958742 | 0.52 | 6.0E-01 | 0 |
| Extreme obesity (childhood) | 2 | 876113 | 939594 | 812631 | 0.52 | 6.0E-01 | 0 |
| MIF - Macrophage migration inhibitory factor | 1 | 950015 | 950015 | 950015 | 0.52 | 6.0E-01 | 0 |
| Age when periods started (menarche) (female) | 184 | 712341 | 1122476 | 196464 | 0.52 | 6.0E-01 | 0 |
| Pericardial adipose tissue volume (male) | 1 | 958743 | 958742 | 958742 | 0.52 | 6.0E-01 | 0 |
| Asthma (adult-onset) | 25 | 767645 | 908280 | 347450 | 0.52 | 6.0E-01 | 0 |
| Worry too long after embarrassment | 22 | 767135 | 1021552 | 138923 | 0.52 | 6.0E-01 | 0 |
| IL1A - Interleukin-1 alpha | 1 | 950015 | 950015 | 950015 | 0.52 | 6.0E-01 | 0 |
| Total lipids in large LDL | 14 | 787651 | 1175429 | 377491 | 0.52 | 6.0E-01 | 0 |
| Total lipids in large VLDL | 7 | 818555 | 979404 | 478593 | 0.53 | 6.0E-01 | 0 |
| Peptide::Dipeptide::X-14304--leucylalanine | 2 | 874809 | 939873 | 809744 | 0.53 | 6.0E-01 | 0 |
| Extreme Waist-hip ratio (adjusted for BMI) | 2 | 880005 | 965983 | 794028 | 0.53 | 5.9E-01 | 0 |
| Cerebrospinal fluid levels of amyloid-beta 1-42 | 1 | 973443 | 973442 | 973442 | 0.53 | 5.9E-01 | 0 |
| Triglyceride | 27 | 765925 | 1023714 | 152967 | 0.53 | 5.9E-01 | 0 |
| SOD2 - Superoxide dismutase [Mn], mitochondrial | 1 | 962338 | 962338 | 962338 | 0.53 | 5.9E-01 | 0 |
| Probable major dpressive disorder | 1 | 966988 | 966988 | 966988 | 0.54 | 5.9E-01 | 0 |
| Phospholipids in small VLDL | 9 | 818555 | 949067 | 514640 | 0.54 | 5.9E-01 | 0 |
| ::::X-12094 | 1 | 967390 | 967390 | 967390 | 0.54 | 5.9E-01 | 0 |
| Infantile hypertrophic pyloric stenosis | 6 | 818603 | 1039886 | 334516 | 0.54 | 5.9E-01 | 0 |
| Phospholipids in very large VLDL | 7 | 836050 | 1068231 | 792240 | 0.54 | 5.9E-01 | 0 |
| Age at menopause (last menstrual period) (female) | 59 | 741363 | 1167133 | 120117 | 0.54 | 5.9E-01 | 0 |
| HCK - Tyrosine-protein kinase HCK | 1 | 969995 | 969995 | 969995 | 0.54 | 5.9E-01 | 0 |
| Cingulum (cingulate gyrus) mean diusivities | 3 | 875653 | 1126300 | 670906 | 0.54 | 5.9E-01 | 0 |

| | | | | | | | |
|---|---|---|---|---|---|---|---|
| Fluid intelligence test - FI7 : synonym | 1 | 961218 | 961217 | 961217 | 0.54 | 5.9E-01 | 0 |
| Sagittal diameter by CT (GEE, adjusted for age, sex, smoking and menopause) | 1 | 969028 | 969028 | 969028 | 0.54 | 5.9E-01 | 0 |
| SLAMF7 - SLAM family member 7 | 1 | 969403 | 969403 | 969403 | 0.54 | 5.9E-01 | 0 |
| Non-cancer illness code, self-reported: migraine | 13 | 800740 | 1061773 | 406448 | 0.55 | 5.8E-01 | 0 |
| Triglycerides in large VLDL | 6 | 824461 | 959667 | 532741 | 0.55 | 5.8E-01 | 0 |
| D-dimer (at exam 6, GEE, adjusted for age and sex) | 3 | 886028 | 988078 | 868826 | 0.55 | 5.8E-01 | 0 |
| GRAP2 - GRB2-related adapter protein 2 | 1 | 982440 | 982440 | 982440 | 0.55 | 5.8E-01 | 0 |
| Triglycerides in small HDL | 9 | 818555 | 1118528 | 765925 | 0.55 | 5.8E-01 | 0 |
| Concentration of large LDL particles | 14 | 787651 | 1175429 | 377491 | 0.55 | 5.8E-01 | 0 |
| Dysmenorrhea (quality of life impact) | 1 | 977578 | 977578 | 977578 | 0.55 | 5.8E-01 | 0 |
| Phospholipids in chylomicrons and extremely large VLDL | 6 | 827303 | 966318 | 800503 | 0.56 | 5.8E-01 | 0 |
| Nucleotide::NAD metabolism::X-12095--N1-methyl-3-pyridone-4-carboxamide | 1 | 967390 | 967390 | 967390 | 0.56 | 5.8E-01 | 0 |
| Diagnoses - main ICD10: R31 Hematuria | 4 | 841234 | 1251904 | 394397 | 0.56 | 5.8E-01 | 0 |
| Alkaline phosphatase | 24 | 772270 | 1046054 | 188651 | 0.56 | 5.8E-01 | 0 |
| Ever highly irritable/argumentative for 2 days | 3 | 886200 | 1003341 | 660144 | 0.56 | 5.7E-01 | 0 |
| Loneliness (MTAG) | 13 | 802985 | 1398988 | 70786 | 0.56 | 5.7E-01 | 0 |
| Plasma Lipoprotein A level | 9 | 824590 | 1055170 | 128412 | 0.56 | 5.7E-01 | 0 |
| White matter hyperintensity volume (GEE, adjusted for age and sex) | 3 | 890213 | 1139348 | 808465 | 0.56 | 5.7E-01 | 0 |
| Lipid::Long chain fatty acid::arachidonate (20:4n6) | 1 | 978718 | 978717 | 978717 | 0.57 | 5.7E-01 | 0 |
| PDGFRB - Platelet-derived growth factor receptor beta | 1 | 979123 | 979123 | 979123 | 0.57 | 5.7E-01 | 0 |
| TYK2 - Non-receptor tyrosine-protein kinase TYK2 | 1 | 982440 | 982440 | 982440 | 0.57 | 5.7E-01 | 0 |
| Myeloid white cell count (three-way meta) | 97 | 730250 | 1219753 | 129958 | 0.57 | 5.7E-01 | 0 |
| Total lipids in small VLDL | 11 | 818555 | 949054 | 430043 | 0.57 | 5.7E-01 | 0 |
| White matter hyperintensity volume (GEE, adjusted for multivariable) | 3 | 890213 | 1139348 | 808465 | 0.57 | 5.7E-01 | 0 |
| MSR1 - Macrophage scavenger receptor types I and II | 1 | 982440 | 982440 | 982440 | 0.58 | 5.7E-01 | 0 |
| Number of treatments/medications taken | 47 | 754590 | 1160480 | 264317 | 0.58 | 5.6E-01 | 0 |
| ABL2 - Abelson tyrosine-protein kinase 2 | 1 | 982440 | 982440 | 982440 | 0.58 | 5.6E-01 | 0 |
| 18:2, linoleic acid (LA) | 11 | 818555 | 1073441 | 216544 | 0.58 | 5.6E-01 | 0 |
| Triglycerides in very small VLDL | 11 | 818555 | 1041381 | 296602 | 0.58 | 5.6E-01 | 0 |
| Alcohol usually taken with meals | 13 | 809173 | 1017803 | 70284 | 0.59 | 5.6E-01 | 0 |
| Neck Section Modulus (GEE, male, adjusted for age) | 2 | 906885 | 1036931 | 776839 | 0.59 | 5.6E-01 | 0 |
| Antithrombotic agents | 9 | 833758 | 1082570 | 385130 | 0.59 | 5.6E-01 | 0 |
| VWF - von Willebrand factor | 1 | 986643 | 986642 | 986642 | 0.59 | 5.6E-01 | 0 |
| Low-density lipoprotein cholesterol (at exam 6, GEE, adjusted for age and sex) | 1 | 989188 | 989188 | 989188 | 0.59 | 5.6E-01 | 0 |
| Ever had prostate specific antigen (PSA) test | 3 | 895335 | 907265 | 760590 | 0.59 | 5.6E-01 | 0 |
| Total cholesterol in small VLDL | 11 | 818555 | 999000 | 444793 | 0.59 | 5.6E-01 | 0 |
| STK17B - Serine/threonine-protein kinase 17B | 1 | 982440 | 982440 | 982440 | 0.59 | 5.6E-01 | 0 |
| BCAR3 - Breast cancer anti-estrogen resistance protein 3 | 1 | 982440 | 982440 | 982440 | 0.59 | 5.5E-01 | 0 |
| IL12RB1 - Interleukin-12 receptor subunit beta-1 | 1 | 981160 | 981160 | 981160 | 0.59 | 5.5E-01 | 0 |
| PPA1 - Inorganic pyrophosphatase | 1 | 991410 | 991410 | 991410 | 0.59 | 5.5E-01 | 0 |
| OCIAD1 - OCIA domain-containing protein 1 | 1 | 982440 | 982440 | 982440 | 0.59 | 5.5E-01 | 0 |
| Stem cell factor | 2 | 896879 | 1077596 | 716162 | 0.59 | 5.5E-01 | 0 |
| Sleep durataion | 9 | 832098 | 982820 | 98504 | 0.59 | 5.5E-01 | 0 |

| | | | | | | | |
|---|---|---|---|---|---|---|---|
| Lipid::Medium chain fatty acid::10-undecenoate (11:1n1) | 1 | 992923 | 992923 | 992923 | 0.59 | 5.5E-01 | 0 |
| Peptide::Dipeptide::aspartylphenylalanine | 1 | 1004938 | 1004938 | 1004938 | 0.60 | 5.5E-01 | 0 |
| Small low-density lipoprotein by NMR (at exam 4, GEE, adjusted for multivariable) | 3 | 908985 | 1031613 | 859315 | 0.60 | 5.5E-01 | 0 |
| 22:6, docosahexaenoic acid (DHA) | 3 | 899838 | 1011653 | 696946 | 0.60 | 5.5E-01 | 0 |
| PR interval (cohort exam 11 offspring exam 1, GEE, adjusted for age nad RR) | 3 | 907803 | 1028435 | 680969 | 0.60 | 5.5E-01 | 0 |
| Left parahippocampal | 3 | 909943 | 1133163 | 502112 | 0.60 | 5.5E-01 | 0 |
| Fasting plasma glucose (at exam 5, GEE, adjusted for age, sex and BMI) | 3 | 902585 | 969401 | 503440 | 0.60 | 5.5E-01 | 0 |
| Treatment/medication code: ibuprofen | 3 | 912318 | 962444 | 467493 | 0.61 | 5.4E-01 | 0 |
| ::::X-14086 | 1 | 1004938 | 1004938 | 1004938 | 0.61 | 5.4E-01 | 0 |
| Lipid::Sterol, Steroid::androsterone sulfate | 2 | 914850 | 1052080 | 777620 | 0.61 | 5.4E-01 | 0 |
| Cerebellar vermal lobules I V | 14 | 807139 | 1032873 | 105391 | 0.61 | 5.4E-01 | 0 |
| Serum total triglycerides | 10 | 824461 | 934064 | 599834 | 0.62 | 5.4E-01 | 0 |
| Lipid::Lysolipid::1-arachidonoylglycerophosphocholine* | 3 | 918253 | 957725 | 903789 | 0.62 | 5.4E-01 | 0 |
| CSF | 7 | 854105 | 914416 | 200127 | 0.62 | 5.4E-01 | 0 |
| Cholesterol esters in medium VLDL | 13 | 818555 | 1048933 | 485140 | 0.62 | 5.4E-01 | 0 |
| Menstrual pain medicine use | 1 | 1000075 | 1000075 | 1000075 | 0.62 | 5.4E-01 | 0 |
| Lipid::Lysolipid::1-linoleoylglycerophosphoethanolamine* | 2 | 907378 | 916404 | 898351 | 0.62 | 5.4E-01 | 0 |
| Glucose | 4 | 863451 | 1014154 | 689373 | 0.62 | 5.4E-01 | 0 |
| Epworth Sleepiness Scale (GEE) | 1 | 1017438 | 1017438 | 1017438 | 0.62 | 5.3E-01 | 0 |
| Small low-density lipoprotein by NMR (at exam 4, GEE, adjusted for age and sex) | 4 | 860539 | 972134 | 783952 | 0.62 | 5.3E-01 | 0 |
| Number of days/week of vigorous physical activity 10+ minutes | 10 | 824791 | 1006328 | 634146 | 0.62 | 5.3E-01 | 0 |
| QRS interval | 8 | 833528 | 1174862 | 568435 | 0.62 | 5.3E-01 | 0 |
| Interleukin-16 | 2 | 916238 | 968164 | 864311 | 0.63 | 5.3E-01 | 0 |
| Morning person (binary) | 116 | 730095 | 1180721 | 106761 | 0.63 | 5.3E-01 | 0 |
| Total cerebral brain volume (GEE, adjusted for multivariable) | 1 | 1002018 | 1002017 | 1002017 | 0.63 | 5.3E-01 | 0 |
| Triglycerides in very large HDL | 10 | 823634 | 1163668 | 650053 | 0.63 | 5.3E-01 | 0 |
| Age started wearing glasses or contact lenses | 100 | 738014 | 1074956 | 148191 | 0.63 | 5.3E-01 | 0 |
| Seen a psychiatrist for nerves, anxiety, tension or depression | 3 | 920673 | 1054678 | 483596 | 0.63 | 5.3E-01 | 0 |
| Triglycerides in small VLDL | 11 | 830368 | 1137843 | 490970 | 0.63 | 5.3E-01 | 0 |
| Eosinophil percentage of white cells (two-way meta) | 97 | 736083 | 1223408 | 122584 | 0.63 | 5.3E-01 | 0 |
| ESAM - Endothelial cell-selective adhesion molecule | 1 | 1006610 | 1006610 | 1006610 | 0.63 | 5.3E-01 | 0 |
| GFRA2 - GDNF family receptor alpha-2 | 1 | 1023148 | 1023148 | 1023148 | 0.64 | 5.2E-01 | 0 |
| Inter-trochanteric buckling ratio (GEE, female, adjussted for age) | 1 | 1025960 | 1025960 | 1025960 | 0.64 | 5.2E-01 | 0 |
| Diagnoses - main ICD10: K40 Inguinal hernia | 14 | 820809 | 1125520 | 73491 | 0.64 | 5.2E-01 | 0 |
| Inter-trochanteric buckling ratio (GEE, male, adjussted for multivariable) | 1 | 1019215 | 1019215 | 1019215 | 0.64 | 5.2E-01 | 0 |
| Survival past average life expectancy (GEE, adjusted for birth-cohort) | 1 | 1020063 | 1020063 | 1020063 | 0.65 | 5.2E-01 | 0 |
| Insomnia | 14 | 814311 | 1289623 | 64540 | 0.65 | 5.2E-01 | 0 |
| 28 year time-averaged Fasting plasma glucose (GEE, adjusted for age and sex) | 9 | 851625 | 1214860 | 104295 | 0.65 | 5.2E-01 | 0 |
| Resting heart rate | 290 | 711600 | 1123018 | 159015 | 0.65 | 5.2E-01 | 0 |
| EMR2 - EGF-like module-containing mucin-like hormone receptor-like 2 | 1 | 1038403 | 1038403 | 1038403 | 0.65 | 5.2E-01 | 0 |
| FN1 - Fibronectin Fragment 4 | 1 | 1027838 | 1027838 | 1027838 | 0.65 | 5.1E-01 | 0 |
| Systemic Lupus Erythematosus | 61 | 756663 | 1285880 | 293270 | 0.65 | 5.1E-01 | 0 |
| Fractured/broken bones in last 5 years | 13 | 825893 | 1057395 | 434027 | 0.65 | 5.1E-01 | 0 |

| | | | | | | | |
|---|---|---|---|---|---|---|---|
| Thyroid-stimulating hormone (female) | 21 | 801673 | 1080473 | 576603 | 0.66 | 5.1E-01 | 0 |
| Neck Section Modulus (GEE, adjusted for age and sex) | 1 | 1036765 | 1036765 | 1036765 | 0.66 | 5.1E-01 | 0 |
| Nucleotide::Purine metabolism, (hypo)xanthine, inosine containing::hypoxanthine | 1 | 1036650 | 1036650 | 1036650 | 0.66 | 5.1E-01 | 0 |
| Posterior corona radiata fractional anisotropy | 2 | 922019 | 923558 | 920479 | 0.66 | 5.1E-01 | 0 |
| FN1 - Fibronectin Fragment 3 | 1 | 1027838 | 1027838 | 1027838 | 0.66 | 5.1E-01 | 0 |
| Coronary artery disease and high-density lipoprotein cholesterol (bivariate) | 144 | 725998 | 1125248 | 217187 | 0.66 | 5.1E-01 | 0 |
| Life satisfaction (MA GWAMA) | 32 | 779481 | 1198157 | 381741 | 0.66 | 5.1E-01 | 0 |
| Trunk-trunk fat ratio (male) | 15 | 822398 | 1220179 | 168480 | 0.66 | 5.1E-01 | 0 |
| FN1 - Fibronectin | 2 | 929173 | 978505 | 879840 | 0.66 | 5.1E-01 | 0 |
| Short sleep | 21 | 805503 | 1078588 | 91238 | 0.67 | 5.0E-01 | 0 |
| Resistin (GEE, adjusted for age, sex and BMI) | 2 | 927399 | 1096913 | 757884 | 0.67 | 5.0E-01 | 0 |
| Social support - Leisure/social activities: Pub or social club | 17 | 818465 | 1284278 | 506602 | 0.67 | 5.0E-01 | 0 |
| Coronary artery disease and low-density lipoprotein cholesterol (bivariate) | 133 | 730695 | 1139080 | 361015 | 0.67 | 5.0E-01 | 0 |
| Number of days/week of moderate physical activity 10+ minutes | 5 | 889053 | 1369603 | 558700 | 0.67 | 5.0E-01 | 0 |
| Shaft cross-sectional moment of inertia (GEE, female, adjussted for multivariable) | 2 | 932634 | 1174766 | 690502 | 0.67 | 5.0E-01 | 0 |
| Vitamin and mineral supplements: Multivitamins +/- minerals | 2 | 924765 | 1319770 | 529761 | 0.67 | 5.0E-01 | 0 |
| Glycoprotein acetyls, mainly a1Lacid glycoprotein | 4 | 883811 | 1120143 | 617628 | 0.67 | 5.0E-01 | 0 |
| NKeff:%337+ | 1 | 1037885 | 1037885 | 1037885 | 0.67 | 5.0E-01 | 0 |
| Mouth/teeth dental problems: Dentures | 30 | 784160 | 1066246 | 208581 | 0.67 | 5.0E-01 | 0 |
| GPC3 - Glypican-3 | 1 | 1028153 | 1028153 | 1028153 | 0.67 | 5.0E-01 | 0 |
| Granulocyte count (three-way meta) | 102 | 737218 | 1216351 | 155431 | 0.67 | 5.0E-01 | 0 |
| Urinary albumin-to-creatinine ratio | 2 | 929724 | 1058693 | 800754 | 0.67 | 5.0E-01 | 0 |
| Neck width (GEE, female, adjusted for multivariable) | 1 | 1025960 | 1025960 | 1025960 | 0.67 | 5.0E-01 | 0 |
| Neutrophil percentage of granulocytes (three-way meta) | 107 | 736083 | 1184711 | 138590 | 0.68 | 5.0E-01 | 0 |
| Coronary artery disease and total cholesterol (bivariate) | 135 | 730695 | 1120903 | 376863 | 0.68 | 5.0E-01 | 0 |
| Serum calcium (at exam 2, GEE, adjusted for age, sex and serum creatinine) | 2 | 933483 | 1053635 | 813330 | 0.68 | 5.0E-01 | 0 |
| Hot drink temperature | 45 | 767540 | 1294605 | 72826 | 0.68 | 5.0E-01 | 0 |
| Concentration of IDL particles | 12 | 831980 | 1050478 | 304479 | 0.68 | 5.0E-01 | 0 |
| Fasting insulin | 1 | 1031433 | 1031433 | 1031433 | 0.68 | 5.0E-01 | 0 |
| LRIG3 - Leucine-rich repeats and immunoglobulin-like domains protein 3 | 1 | 1039958 | 1039958 | 1039958 | 0.68 | 4.9E-01 | 0 |
| SPON1 - Spondin-1 | 1 | 1033753 | 1033753 | 1033753 | 0.68 | 4.9E-01 | 0 |
| OmegaL3 fatty acids | 5 | 899838 | 1045825 | 494055 | 0.69 | 4.9E-01 | 0 |
| ::::X-12450 | 1 | 1043453 | 1043453 | 1043453 | 0.69 | 4.9E-01 | 0 |
| KLRK1 - NKG2-D type II integral membrane protein | 1 | 1049338 | 1049338 | 1049338 | 0.69 | 4.9E-01 | 0 |
| ::::X-12644 | 1 | 1045825 | 1045825 | 1045825 | 0.69 | 4.9E-01 | 0 |
| Anxiety - Ever worried more than most people would in similar situation | 1 | 1052310 | 1052310 | 1052310 | 0.69 | 4.9E-01 | 0 |
| Thyroid-stimulating hormone | 39 | 778293 | 929274 | 498726 | 0.69 | 4.9E-01 | 0 |
| Age when last used oral contraceptive pill (female) | 1 | 1044205 | 1044205 | 1044205 | 0.70 | 4.9E-01 | 0 |
| Atrial natriuretic peptide (at exam 6, GEE, adjusted for multivariable) | 1 | 1053973 | 1053973 | 1053973 | 0.70 | 4.9E-01 | 0 |
| EPHB6 - Ephrin type-B receptor 6 | 1 | 1057895 | 1057895 | 1057895 | 0.70 | 4.9E-01 | 0 |
| Pericardial adipose tissue volume (adjusted for height and weight, female) | 1 | 1051960 | 1051960 | 1051960 | 0.70 | 4.9E-01 | 0 |
| Nucleotide::Purine metabolism, guanine containing::N2,N2-dimethylguanosine | 1 | 1060300 | 1060300 | 1060300 | 0.70 | 4.8E-01 | 0 |
| Histidine | 4 | 894400 | 1058084 | 640250 | 0.70 | 4.8E-01 | 0 |

| | | | | | | | |
|---|---|---|---|---|---|---|---|
| Atrial Fibrillation | 103 | 740255 | 1119101 | 348511 | 0.70 | 4.8E-01 | 0 |
| Lipid::Carnitine metabolism::stearoylcarnitine | 1 | 1051960 | 1051960 | 1051960 | 0.70 | 4.8E-01 | 0 |
| Fibrotic idiopathic interstitial pneumonias | 6 | 873483 | 1013531 | 497613 | 0.70 | 4.8E-01 | 0 |
| Immature fraction of reticulocytes (three-way meta) | 102 | 741580 | 1121044 | 240916 | 0.70 | 4.8E-01 | 0 |
| Hypothyroidism | 8 | 855323 | 1193860 | 369793 | 0.71 | 4.8E-01 | 0 |
| Posterior thalamic radiation (include optic radiation) axial diusivities | 2 | 938233 | 1255351 | 621114 | 0.71 | 4.8E-01 | 0 |
| Basophil count (three-way meta) | 53 | 766478 | 1112530 | 96751 | 0.71 | 4.8E-01 | 0 |
| Eosinophil percentage of white cells (three-way meta) | 119 | 739293 | 1151696 | 124665 | 0.71 | 4.8E-01 | 0 |
| PDGFB - Platelet-derived growth factor subunit B | 1 | 1051143 | 1051143 | 1051143 | 0.71 | 4.8E-01 | 0 |
| Depressive symptoms (univariate) | 53 | 767688 | 1141800 | 149863 | 0.71 | 4.8E-01 | 0 |
| MFGE8 - Lactadherin | 1 | 1055668 | 1055668 | 1055668 | 0.71 | 4.8E-01 | 0 |
| Periodontal complex trait 5 - Pg trait | 2 | 943671 | 982687 | 904656 | 0.71 | 4.8E-01 | 0 |
| Triglyceride / High-density lipoprotein cholesterol ratio (at exam 1, GEE, adjusted for multivariable | 1 | 1060918 | 1060918 | 1060918 | 0.71 | 4.8E-01 | 0 |
| Illnesses of siblings: Breast cancer | 3 | 951490 | 1005368 | 501975 | 0.71 | 4.8E-01 | 0 |
| MAPK3 - Mitogen-activated protein kinase 3 | 1 | 1054850 | 1054850 | 1054850 | 0.71 | 4.8E-01 | 0 |
| Alkaline phosphatase (at exam 2, GEE, adjusted for age and sex) | 1 | 1050385 | 1050385 | 1050385 | 0.71 | 4.8E-01 | 0 |
| Xenobiotics::Benzoate metabolism::4-vinylphenol sulfate | 1 | 1060360 | 1060360 | 1060360 | 0.71 | 4.8E-01 | 0 |
| Diagnoses - secondary ICD10: E78 Disorders of lipoprotein metabolism and other lipidemias | 21 | 818555 | 1079695 | 214394 | 0.71 | 4.8E-01 | 0 |
| Facial attractiveness (male-coder, female samples) | 1 | 1052485 | 1052485 | 1052485 | 0.71 | 4.8E-01 | 0 |
| Reaction time test - Mean time to correctly identify matches | 37 | 783943 | 1105768 | 73004 | 0.71 | 4.8E-01 | 0 |
| Total cholesterol in large HDL | 10 | 853804 | 1051374 | 596580 | 0.71 | 4.8E-01 | 0 |
| High-density lipoprotein cholesterol | 100 | 742574 | 1117746 | 230108 | 0.71 | 4.7E-01 | 0 |
| Heart rate recovery at 40 secnds | 18 | 818834 | 1056066 | 381888 | 0.72 | 4.7E-01 | 0 |
| Heart rate recovery at 50 secnds | 18 | 818834 | 1075021 | 381888 | 0.72 | 4.7E-01 | 0 |
| Diagnoses - secondary ICD10: K57 Diverticular disease of intestine | 10 | 849669 | 1025939 | 205592 | 0.72 | 4.7E-01 | 0 |
| FAP - Seprase | 1 | 1065780 | 1065780 | 1065780 | 0.72 | 4.7E-01 | 0 |
| Sitting height | 659 | 700768 | 1127885 | 108483 | 0.72 | 4.7E-01 | 0 |
| DYNLRB1 - Dynein light chain roadblock-type 1 | 1 | 1062243 | 1062243 | 1062243 | 0.72 | 4.7E-01 | 0 |
| CD209 - CD209 antigen | 1 | 1064218 | 1064218 | 1064218 | 0.72 | 4.7E-01 | 0 |
| METAP2 - Methionine aminopeptidase 2 | 1 | 1062243 | 1062243 | 1062243 | 0.73 | 4.7E-01 | 0 |
| Pulse rate (automated reading) | 202 | 723834 | 1153959 | 196015 | 0.73 | 4.7E-01 | 0 |
| Corrected insulin response | 3 | 960418 | 1036844 | 547583 | 0.73 | 4.6E-01 | 0 |
| CAT - Catalase | 2 | 953029 | 1052771 | 853287 | 0.73 | 4.6E-01 | 0 |
| ::::X-13859 | 2 | 956335 | 1410120 | 502550 | 0.74 | 4.6E-01 | 0 |
| MonoLunsaturated fatty acids | 6 | 883811 | 988469 | 250732 | 0.74 | 4.6E-01 | 0 |
| Medication for cholesterol, blood pressure or diabetes: Blood pressure medication | 50 | 770760 | 1296984 | 288661 | 0.74 | 4.6E-01 | 0 |
| Xenobiotics::Xanthine metabolism::3-methylxanthine | 2 | 953061 | 958687 | 947436 | 0.74 | 4.6E-01 | 0 |
| Vitamin and mineral supplements: Vitamin E | 1 | 1067763 | 1067763 | 1067763 | 0.74 | 4.6E-01 | 0 |
| ::::X-13435 | 2 | 948226 | 1030456 | 865997 | 0.74 | 4.6E-01 | 0 |
| Asthma (random effect model) | 18 | 825820 | 1147311 | 395958 | 0.74 | 4.6E-01 | 0 |
| Free thyroxine (FT4) | 18 | 822718 | 1196573 | 448066 | 0.75 | 4.6E-01 | 0 |
| Amino acid::Cysteine, methionine, SAM, taurine metabolism::X-11786--methylcysteine | 1 | 1075038 | 1075038 | 1075038 | 0.75 | 4.6E-01 | 0 |
| Age stopped smoking | 1 | 1076735 | 1076735 | 1076735 | 0.75 | 4.5E-01 | 0 |

| | | | | | | | |
|---|---|---|---|---|---|---|---|
| Medication for cholesterol, blood pressure, diabetes, or take exogenous hormones: Cholesterol lo | 27 | 809510 | 1178439 | 417343 | 0.75 | 4.5E-01 | 0 |
| ARSB - Arylsulfatase B | 1 | 1073073 | 1073073 | 1073073 | 0.75 | 4.5E-01 | 0 |
| GNLY - Granulysin | 1 | 1083200 | 1083200 | 1083200 | 0.75 | 4.5E-01 | 0 |
| FCER2 - Low affinity immunoglobulin epsilon Fc receptor | 1 | 1080683 | 1080683 | 1080683 | 0.75 | 4.5E-01 | 0 |
| Eotaxin (CCL11) | 3 | 971555 | 1741565 | 500645 | 0.75 | 4.5E-01 | 0 |
| ::::X-08988 | 3 | 972713 | 992768 | 888729 | 0.76 | 4.5E-01 | 0 |
| Mean low-density lipoprotein cholesterol fromexam 1-7 (FBAT, adjusted for multivariable) | 1 | 1085575 | 1085575 | 1085575 | 0.76 | 4.5E-01 | 0 |
| Peptide::Polypeptide::bradykinin, des-arg(9) | 3 | 972068 | 1051986 | 846734 | 0.76 | 4.5E-01 | 0 |
| Body fat percentage | 9 | 878838 | 1353228 | 382215 | 0.76 | 4.5E-01 | 0 |
| CMPK1 - UMP-CMP kinase | 1 | 1089855 | 1089855 | 1089855 | 0.76 | 4.5E-01 | 0 |
| Xenobiotics::Xanthine metabolism::caffeine | 1 | 1082660 | 1082660 | 1082660 | 0.76 | 4.5E-01 | 0 |
| Platelet distribution width (three-way meta) | 113 | 742700 | 1092335 | 354748 | 0.76 | 4.4E-01 | 0 |
| Vigorous physical activity | 5 | 930240 | 1133225 | 751333 | 0.77 | 4.4E-01 | 0 |
| Height SDS | 4 | 915130 | 1273486 | 454842 | 0.77 | 4.4E-01 | 0 |
| IL18R1 - Interleukin-18 receptor 1 | 1 | 1088285 | 1088285 | 1088285 | 0.77 | 4.4E-01 | 0 |
| Guilty feelings (GUILT) | 12 | 854145 | 965692 | 378950 | 0.77 | 4.4E-01 | 0 |
| Vascular/heart problems diagnosed by doctor: Angina | 14 | 842514 | 1116611 | 321236 | 0.77 | 4.4E-01 | 0 |
| PTHLH - Parathyroid hormone-related protein | 1 | 1088588 | 1088588 | 1088588 | 0.77 | 4.4E-01 | 0 |
| CAMK1D - Calcium/calmodulin-dependent protein kinase type 1D | 1 | 1089855 | 1089855 | 1089855 | 0.78 | 4.4E-01 | 0 |
| Lipid::Carnitine metabolism::carnitine | 16 | 838394 | 1061705 | 445802 | 0.78 | 4.4E-01 | 0 |
| Mental distress - Ever suffered mental distress preventing usual activities | 4 | 915493 | 1171415 | 523001 | 0.78 | 4.4E-01 | 0 |
| Left accumbens area | 6 | 894191 | 1283032 | 204119 | 0.78 | 4.4E-01 | 0 |
| Cholesterol esters in large HDL | 9 | 889053 | 946992 | 586548 | 0.78 | 4.4E-01 | 0 |
| IL5 - Interleukin-5 | 1 | 1089855 | 1089855 | 1089855 | 0.78 | 4.3E-01 | 0 |
| Mean low-density lipoprotein cholesterol fromexam 1-7 (FBAT, adjusted for age and sex) | 1 | 1085575 | 1085575 | 1085575 | 0.78 | 4.3E-01 | 0 |
| Ejection fraction | 2 | 970369 | 1028559 | 912178 | 0.78 | 4.3E-01 | 0 |
| Large high-density lipoprotein by NMR (at exam 4, GEE, adjusted for age and sex) | 1 | 1086448 | 1086448 | 1086448 | 0.79 | 4.3E-01 | 0 |
| Impedance measures - Impedance of leg (right) | 458 | 709886 | 1144254 | 164261 | 0.79 | 4.3E-01 | 0 |
| PRSS27 - Serine protease 27 | 1 | 1089855 | 1089855 | 1089855 | 0.79 | 4.3E-01 | 0 |
| Diagnoses - secondary ICD10: I25 Chronic ischemic heart disease | 15 | 851270 | 1309281 | 431029 | 0.79 | 4.3E-01 | 0 |
| WISP1 - WNT1-inducible-signaling pathway protein 1 | 1 | 1089855 | 1089855 | 1089855 | 0.79 | 4.3E-01 | 0 |
| Alcohol - Alcohol drinker status: Never | 6 | 898186 | 1250929 | 253162 | 0.79 | 4.3E-01 | 0 |
| Traumatic events - Felt loved as child | 3 | 976820 | 1178106 | 741361 | 0.79 | 4.3E-01 | 0 |
| Superior corona radiata mode of anisotropy | 3 | 982630 | 1103274 | 529679 | 0.79 | 4.3E-01 | 0 |
| SEMA3A - Semaphorin-3A | 2 | 974509 | 1032182 | 916836 | 0.79 | 4.3E-01 | 0 |
| Other serious medical condition/disability (diagnosed by doctor) | 3 | 991493 | 1408148 | 546305 | 0.79 | 4.3E-01 | 0 |
| Left rostral anterior cingulate | 4 | 922554 | 1338835 | 389791 | 0.79 | 4.3E-01 | 0 |
| Posterior corona radiata mode of anisotropy | 3 | 988860 | 1428891 | 754460 | 0.79 | 4.3E-01 | 0 |
| Red blood cell count | 41 | 787993 | 1045680 | 245007 | 0.79 | 4.3E-01 | 0 |
| Femoral Neck BMD | 36 | 796618 | 1241681 | 419366 | 0.79 | 4.3E-01 | 0 |
| ::::X-12798 | 3 | 986395 | 1216105 | 523425 | 0.80 | 4.3E-01 | 0 |
| Diagnoses - secondary ICD10: Z95 Presence of cardiac and vascular implants and grafts | 9 | 890533 | 1034870 | 678693 | 0.80 | 4.3E-01 | 0 |
| Lipid::Long chain fatty acid::myristoleate (14:1n5) | 1 | 1113115 | 1113115 | 1113115 | 0.80 | 4.3E-01 | 0 |

| | | | | | | | |
|---|---|---|---|---|---|---|---|
| VLDL diameter | 10 | 871420 | 1266549 | 782259 | 0.80 | 4.3E-01 | 0 |
| Lipid::Long chain fatty acid::margarate (17:0) | 1 | 1113115 | 1113115 | 1113115 | 0.80 | 4.3E-01 | 0 |
| Headaches for 3+ months | 4 | 924608 | 1075645 | 649757 | 0.80 | 4.2E-01 | 0 |
| Anti-inflammatory and antirheumatic products, non-steroids | 6 | 903869 | 1095183 | 247398 | 0.80 | 4.2E-01 | 0 |
| Pain type(s) experienced in last month: Back pain | 8 | 887976 | 998662 | 670678 | 0.80 | 4.2E-01 | 0 |
| Medication for pain relief, constipation, heartburn: Ibuprofen (e.g. Nurofen) | 7 | 912318 | 1326158 | 444606 | 0.80 | 4.2E-01 | 0 |
| Change in serum creatinine from exam 2 to 7 (GEE, adjusted for age and sex) | 1 | 1110343 | 1110343 | 1110343 | 0.81 | 4.2E-01 | 0 |
| Corticospinal tract radial diusivities | 5 | 948305 | 1178285 | 870740 | 0.81 | 4.2E-01 | 0 |
| Neck Width (GEE, adjusted for age and sex) | 1 | 1100235 | 1100235 | 1100235 | 0.81 | 4.2E-01 | 0 |
| Posterior corona radiata radial diusivities | 1 | 1113795 | 1113795 | 1113795 | 0.81 | 4.2E-01 | 0 |
| Lipid::Long chain fatty acid::palmitate (16:0) | 1 | 1113115 | 1113115 | 1113115 | 0.81 | 4.2E-01 | 0 |
| Birth length | 2 | 988131 | 1337664 | 638598 | 0.82 | 4.1E-01 | 0 |
| Lipid::Long chain fatty acid::palmitoleate (16:1n7) | 1 | 1113115 | 1113115 | 1113115 | 0.82 | 4.1E-01 | 0 |
| Lipid::Medium chain fatty acid::5-dodecenoate (12:1n7) | 1 | 1113115 | 1113115 | 1113115 | 0.82 | 4.1E-01 | 0 |
| Free cholesterol | 9 | 899838 | 1045825 | 374945 | 0.82 | 4.1E-01 | 0 |
| Mean corpuscular hemoglobin (three-way meta) | 145 | 738943 | 1129463 | 320653 | 0.82 | 4.1E-01 | 0 |
| Automobile speeding propensity | 31 | 813355 | 1119875 | 538151 | 0.82 | 4.1E-01 | 0 |
| Lipid::Long chain fatty acid::nonadecanoate (19:0) | 1 | 1113115 | 1113115 | 1113115 | 0.82 | 4.1E-01 | 0 |
| OmegaL7 and L9 and saturated fatty acids | 5 | 949068 | 1001603 | 818555 | 0.82 | 4.1E-01 | 0 |
| Lipid::Carnitine metabolism::3-dehydrocarnitine* | 2 | 980161 | 983278 | 977044 | 0.82 | 4.1E-01 | 0 |
| Triglycerides in very large VLDL | 6 | 910661 | 1100963 | 779083 | 0.82 | 4.1E-01 | 0 |
| Lipid::Lysolipid::1-palmitoleoylglycerophosphocholine* | 1 | 1113115 | 1113115 | 1113115 | 0.83 | 4.1E-01 | 0 |
| Systolic Blood Pressure (automated reading) | 209 | 730453 | 1122473 | 139634 | 0.83 | 4.1E-01 | 0 |
| APCS - Serum amyloid P-component | 1 | 1116670 | 1116670 | 1116670 | 0.83 | 4.1E-01 | 0 |
| Free cholesterol in large HDL | 11 | 889053 | 1130259 | 606613 | 0.83 | 4.1E-01 | 0 |
| PLXNC1 - Plexin-C1 | 1 | 1116948 | 1116948 | 1116948 | 0.83 | 4.1E-01 | 0 |
| Open-angle glaucoma (random-effect model) | 37 | 806553 | 1044898 | 168601 | 0.83 | 4.1E-01 | 0 |
| Lymphocyte count (three-way meta) | 114 | 749410 | 1111608 | 171860 | 0.83 | 4.0E-01 | 0 |
| Tyrosine | 3 | 998693 | 1148823 | 528410 | 0.83 | 4.0E-01 | 0 |
| ADAMTS13 - A disintegrin and metalloproteinase with thrombospondin motifs 13 | 1 | 1120203 | 1120203 | 1120203 | 0.83 | 4.0E-01 | 0 |
| Age started hormone-replacement therapy (HRT) (female) | 7 | 922255 | 1408741 | 442714 | 0.84 | 4.0E-01 | 0 |
| Offspring birthweight (maternal) | 7 | 924228 | 1220385 | 660623 | 0.84 | 4.0E-01 | 0 |
| KIR2DL4 - Killer cell immunoglobulin-like receptor 2DL4 | 1 | 1123560 | 1123560 | 1123560 | 0.84 | 4.0E-01 | 0 |
| PPP3CA PPP3R1 - Calcineurin | 1 | 1120980 | 1120980 | 1120980 | 0.84 | 4.0E-01 | 0 |
| Agents acting on the renin-angiotensin system | 145 | 741878 | 1136463 | 114163 | 0.84 | 4.0E-01 | 0 |
| KLK11 - Kallikrein-11 | 1 | 1135173 | 1135173 | 1135173 | 0.85 | 4.0E-01 | 0 |
| High light scatter reticulocyte count (three-way meta) | 120 | 748144 | 1137496 | 190531 | 0.85 | 4.0E-01 | 0 |
| Total cholesterol in medium VLDL | 10 | 883811 | 1099398 | 482911 | 0.85 | 4.0E-01 | 0 |
| Lipid::Lysolipid::1-stearoylglycerophosphoethanolamine | 1 | 1123655 | 1123655 | 1123655 | 0.85 | 4.0E-01 | 0 |
| SPARCL1 - SPARC-like protein 1 | 1 | 1135330 | 1135330 | 1135330 | 0.85 | 4.0E-01 | 0 |
| Triglycerides in chylomicrons and extremely large VLDL | 6 | 914148 | 1102706 | 303142 | 0.85 | 4.0E-01 | 0 |
| ::::X-08402 | 2 | 991993 | 1026720 | 957265 | 0.85 | 3.9E-01 | 0 |
| Hearing difficulty/problems with background noise | 25 | 830990 | 1001603 | 84269 | 0.85 | 3.9E-01 | 0 |

| | | | | | | | |
|---|---|---|---|---|---|---|---|
| Concentration of medium VLDL particles | 10 | 889718 | 1139916 | 625655 | 0.85 | 3.9E-01 | 0 |
| Guilty feelings | 13 | 881448 | 1264640 | 654675 | 0.86 | 3.9E-01 | 0 |
| Attention deficit hyperactivity disorder | 11 | 892855 | 1212943 | 497249 | 0.86 | 3.9E-01 | 0 |
| Fractional shortening | 3 | 1017888 | 1075320 | 935938 | 0.86 | 3.9E-01 | 0 |
| Amino acid::Lysine metabolism::lysine | 1 | 1137315 | 1137315 | 1137315 | 0.86 | 3.9E-01 | 0 |
| Fornix (cres) / Stria terminalis fractional anisotropy | 1 | 1138703 | 1138703 | 1138703 | 0.86 | 3.9E-01 | 0 |
| Comparative height size at age 10 | 457 | 713448 | 1114073 | 168115 | 0.86 | 3.9E-01 | 0 |
| Hip circumference (male) | 24 | 833950 | 1196401 | 458584 | 0.86 | 3.9E-01 | 0 |
| Amino acid::Tryptophan metabolism::tryptophan | 17 | 856618 | 1022270 | 525145 | 0.87 | 3.9E-01 | 0 |
| Illnesses of father: Heart disease | 20 | 846216 | 1119437 | 363832 | 0.88 | 3.8E-01 | 0 |
| Sagittal diameter by CT (GEE, adjusted for age and sex) | 1 | 1127085 | 1127085 | 1127085 | 0.88 | 3.8E-01 | 0 |
| Percent predicted FEF25-75 for latest exam (GEE, adjusted for multiple covariates) | 4 | 950311 | 1321710 | 604336 | 0.88 | 3.8E-01 | 0 |
| Mental health - Illness, injury, bereavement, stress in last 2 years: Financial difficulties | 3 | 1023008 | 1167565 | 854443 | 0.88 | 3.8E-01 | 0 |
| Beta blocking agents | 47 | 795763 | 1235108 | 491945 | 0.88 | 3.8E-01 | 0 |
| White matter hyperintensity volume (GEE, adjusted for multivariable with APOE) | 4 | 956653 | 1114440 | 849339 | 0.88 | 3.8E-01 | 0 |
| Lymphocyte percentage of white cells (two-way meta) | 83 | 768645 | 1144873 | 196233 | 0.88 | 3.8E-01 | 0 |
| Xenobiotics::Xanthine metabolism::1-methylxanthine | 1 | 1165490 | 1165490 | 1165490 | 0.88 | 3.8E-01 | 0 |
| Open-angle glaucoma (fixed-effect model) | 41 | 806553 | 1059860 | 174462 | 0.88 | 3.8E-01 | 0 |
| Lipid::Medium chain fatty acid::pelargonate (9:0) | 1 | 1151108 | 1151108 | 1151108 | 0.88 | 3.8E-01 | 0 |
| Neuroticism general factor | 46 | 796220 | 1230458 | 99213 | 0.88 | 3.8E-01 | 0 |
| CFHR5 - Complement factor H-related protein 5 | 1 | 1151555 | 1151555 | 1151555 | 0.88 | 3.8E-01 | 0 |
| Anterior limb of internal capsule axial diusivities | 2 | 1013764 | 1256476 | 771052 | 0.89 | 3.8E-01 | 0 |
| Treatment/medication code: metformin | 29 | 832305 | 1152513 | 428733 | 0.89 | 3.8E-01 | 0 |
| Left ventral DC | 6 | 933798 | 1184813 | 300938 | 0.89 | 3.7E-01 | 0 |
| Waist circumference (male, adjusted for BMI) | 37 | 813175 | 1190793 | 112308 | 0.89 | 3.7E-01 | 0 |
| Free cholesterol in large VLDL | 8 | 919409 | 1118562 | 805398 | 0.89 | 3.7E-01 | 0 |
| Mean arterial pressure | 21 | 848045 | 1122783 | 263795 | 0.89 | 3.7E-01 | 0 |
| Total lipids in very small VLDL | 12 | 883811 | 1050478 | 379486 | 0.89 | 3.7E-01 | 0 |
| Total cholesterol / High-density lipoprotein cholesterol ratio (at exam 6, GEE, adjusted for age an | 2 | 1013751 | 1234507 | 792996 | 0.89 | 3.7E-01 | 0 |
| Concentration of large HDL particles | 8 | 916110 | 1110696 | 760553 | 0.89 | 3.7E-01 | 0 |
| Acetoacetate | 2 | 1016169 | 1114976 | 917362 | 0.89 | 3.7E-01 | 0 |
| Lung canger (adjusted for age) | 3 | 1026643 | 1160283 | 980650 | 0.89 | 3.7E-01 | 0 |
| Mean corpuscular volume | 75 | 769498 | 1076421 | 198935 | 0.90 | 3.7E-01 | 0 |
| Lipid::Long chain fatty acid::X-12442--5,8-tetradecadienoate | 1 | 1158020 | 1158020 | 1158020 | 0.90 | 3.7E-01 | 0 |
| Daytime napping | 7 | 946520 | 1066675 | 399486 | 0.90 | 3.7E-01 | 0 |
| Amino acid::Phenylalanine & tyrosine metabolism::phenylalanine | 1 | 1148495 | 1148495 | 1148495 | 0.90 | 3.7E-01 | 0 |
| Concentration of very small VLDL particles | 12 | 883811 | 1070123 | 381858 | 0.90 | 3.7E-01 | 0 |
| Concentration of very large VLDL particles | 7 | 949068 | 1071718 | 669135 | 0.90 | 3.7E-01 | 0 |
| Amyotrophic lateral sclerosis | 6 | 935901 | 1369743 | 171442 | 0.90 | 3.7E-01 | 0 |
| NRP1 - Neuropilin-1 | 1 | 1147413 | 1147413 | 1147413 | 0.90 | 3.7E-01 | 0 |
| Amino acid::Glycine, serine and threonine metabolism::N-acetylglycine | 2 | 1018876 | 1125942 | 911811 | 0.90 | 3.7E-01 | 0 |
| Total body BMD | 61 | 781903 | 1203333 | 339343 | 0.90 | 3.7E-01 | 0 |
| Anterior corona radiata mean diusivities | 5 | 975900 | 1147350 | 466160 | 0.91 | 3.6E-01 | 0 |

| | | | | | | | |
|---|---|---|---|---|---|---|---|
| RGMA - Repulsive guidance molecule A | 1 | 1152513 | 1152513 | 1152513 | 0.91 | 3.6E-01 | 0 |
| Total lipids in very large VLDL | 7 | 949068 | 1071718 | 486283 | 0.91 | 3.6E-01 | 0 |
| GRN - Granulins | 1 | 1152513 | 1152513 | 1152513 | 0.91 | 3.6E-01 | 0 |
| Total lipids in large HDL | 8 | 916110 | 1110696 | 760553 | 0.91 | 3.6E-01 | 0 |
| Noisy workplace | 1 | 1164618 | 1164618 | 1164618 | 0.91 | 3.6E-01 | 0 |
| Left lingual | 5 | 983585 | 1497278 | 94282 | 0.91 | 3.6E-01 | 0 |
| Open-angle glaucoma | 6 | 935514 | 1277202 | 646358 | 0.91 | 3.6E-01 | 0 |
| Lipid::Lysolipid::1-arachidonoylglycerophosphoinositol* | 4 | 968436 | 1194688 | 708983 | 0.91 | 3.6E-01 | 0 |
| PPID - Peptidyl-prolyl cis-trans isomerase D | 1 | 1163780 | 1163780 | 1163780 | 0.92 | 3.6E-01 | 0 |
| Waist-hip ratio (male, adjusted for BMI) | 10 | 905414 | 1397184 | 692481 | 0.92 | 3.6E-01 | 0 |
| Lamb/mutton intake | 18 | 867938 | 1034478 | 69147 | 0.92 | 3.6E-01 | 0 |
| Comparative body size at age 10 | 172 | 740809 | 1201341 | 137596 | 0.92 | 3.6E-01 | 0 |
| Total cholesterol | 121 | 755865 | 1162523 | 387173 | 0.92 | 3.6E-01 | 0 |
| Percent predicted FEF25-75/FVC for latest exam (GEE, adjusted for multiple covariates) | 4 | 969626 | 1350683 | 562959 | 0.92 | 3.6E-01 | 0 |
| Amino acid::Valine, leucine and isoleucine metabolism::leucine | 11 | 913785 | 1076463 | 599483 | 0.92 | 3.6E-01 | 0 |
| Worry too long after embarrassment (WORR-EMB) | 18 | 861511 | 1050194 | 352567 | 0.92 | 3.6E-01 | 0 |
| Mean corpuscular volume (two-way meta) | 147 | 748590 | 1158769 | 158722 | 0.92 | 3.6E-01 | 0 |
| Number of self-reported non-cancer illnesses | 36 | 817938 | 1414458 | 363062 | 0.92 | 3.6E-01 | 0 |
| Male pattern baldness (BOLT LMM non-infinitesimal mixed model) | 347 | 723280 | 1105631 | 179495 | 0.92 | 3.6E-01 | 0 |
| IMPDH2 - Inosine-5'-monophosphate dehydrogenase 2 | 1 | 1162328 | 1162328 | 1162328 | 0.92 | 3.6E-01 | 0 |
| Amino acid::Glycine, serine and threonine metabolism::betaine | 3 | 1036218 | 1223268 | 920481 | 0.92 | 3.6E-01 | 0 |
| Diagnoses - main ICD10: K80 Cholelithiasis | 15 | 881008 | 1127658 | 80472 | 0.92 | 3.6E-01 | 0 |
| GPC5 - Glypican-5 | 1 | 1184868 | 1184868 | 1184868 | 0.93 | 3.5E-01 | 0 |
| CNTN2 - Contactin-2 | 1 | 1175578 | 1175578 | 1175578 | 0.93 | 3.5E-01 | 0 |
| GOT1 - Aspartate aminotransferase, cytoplasmic | 1 | 1154430 | 1154430 | 1154430 | 0.93 | 3.5E-01 | 0 |
| Shaft Section Modulus (GEE, male, adjussted for multivariable) | 1 | 1167850 | 1167850 | 1167850 | 0.93 | 3.5E-01 | 0 |
| Aspartate aminotransferase (at exam 2, GEE, adjusted for multivariable) | 2 | 1036616 | 1174206 | 899027 | 0.94 | 3.5E-01 | 0 |
| Shaft Width (GEE, male, adjussted for multivariable) | 1 | 1167850 | 1167850 | 1167850 | 0.94 | 3.5E-01 | 0 |
| Impedance measures - Trunk fat mass | 428 | 717713 | 1141230 | 118706 | 0.94 | 3.5E-01 | 0 |
| Moderate to vigorous physical activity levels | 19 | 873448 | 1319581 | 61573 | 0.94 | 3.5E-01 | 0 |
| Diverticular disease | 61 | 789088 | 1138955 | 162788 | 0.94 | 3.5E-01 | 0 |
| Shaft cross-sectional moment of inertia (GEE, male, adjussted for multivariable) | 1 | 1167850 | 1167850 | 1167850 | 0.95 | 3.4E-01 | 0 |
| PEF | 266 | 730150 | 1167076 | 165896 | 0.95 | 3.4E-01 | 0 |
| Lymphocyte count (two-way meta) | 112 | 760833 | 1144230 | 205409 | 0.95 | 3.4E-01 | 0 |
| Serum creatinine | 51 | 804745 | 1107733 | 201162 | 0.95 | 3.4E-01 | 0 |
| Happiness and subjective well-being - General happiness | 1 | 1181195 | 1181195 | 1181195 | 0.95 | 3.4E-01 | 0 |
| Left superior temporal | 1 | 1191618 | 1191618 | 1191618 | 0.95 | 3.4E-01 | 0 |
| Lipid::Long chain fatty acid::stearidonate (18:4n3) | 1 | 1181688 | 1181688 | 1181688 | 0.95 | 3.4E-01 | 0 |
| Cancer register - Type of cancer: ICD10: C50 Malignant neoplasm of breast | 9 | 936240 | 1069450 | 464265 | 0.95 | 3.4E-01 | 0 |
| Never eat eggs, dairy, wheat, sugar: I eat all of the above | 11 | 924228 | 1128154 | 624635 | 0.95 | 3.4E-01 | 0 |
| Diagnoses - secondary ICD10: Z92 Personal history of medical treatment | 3 | 1053010 | 1791330 | 1049563 | 0.96 | 3.4E-01 | 0 |
| Age hay fever, rhinitis or eczema diagnosed | 24 | 849218 | 1202956 | 498121 | 0.96 | 3.4E-01 | 0 |
| Phospholipids in large HDL | 9 | 943168 | 1141990 | 818555 | 0.96 | 3.4E-01 | 0 |

| | | | | | | | |
|---|---|---|---|---|---|---|---|
| Amino acid::Alanine and aspartate metabolism::alanine | 2 | 1036141 | 1334754 | 737528 | 0.96 | 3.4E-01 | 0 |
| Systolic Blood Pressure | 29 | 848045 | 1103473 | 223135 | 0.96 | 3.4E-01 | 0 |
| Triglyceride / High-density lipoprotein cholesterol ratio (at exam 2, GEE, adjusted for age and sex | 1 | 1183588 | 1183588 | 1183588 | 0.96 | 3.4E-01 | 0 |
| Microalbuminuria | 1 | 1187663 | 1187663 | 1187663 | 0.96 | 3.4E-01 | 0 |
| Glioma | 12 | 899935 | 1304187 | 617321 | 0.96 | 3.3E-01 | 0 |
| Other polyunsaturated fatty acids than 18:2 | 6 | 957164 | 1027002 | 636089 | 0.97 | 3.3E-01 | 0 |
| Mean erythrocyte cell volume | 4 | 978601 | 1075023 | 887002 | 0.97 | 3.3E-01 | 0 |
| Follicle stimulating hormone (at exam 3, GEE, adjusted for age and sex) | 3 | 1058510 | 1095961 | 563762 | 0.97 | 3.3E-01 | 0 |
| Social support - Leisure/social activities: Sports club or gym | 9 | 948488 | 1036333 | 104020 | 0.97 | 3.3E-01 | 0 |
| Left basal forebrain | 3 | 1065723 | 1487795 | 555382 | 0.97 | 3.3E-01 | 0 |
| Medication for cholesterol, blood pressure or diabetes: Cholesterol lowering medication | 33 | 833758 | 1047450 | 229329 | 0.97 | 3.3E-01 | 0 |
| EDAR - Tumor necrosis factor receptor superfamily member EDAR | 2 | 1044644 | 1280314 | 808973 | 0.98 | 3.3E-01 | 0 |
| Stem cell growth factor beta | 2 | 1044773 | 1117041 | 972504 | 0.98 | 3.3E-01 | 0 |
| Non-cancer illness code, self-reported: hypothyroidism/myxoedema | 75 | 783140 | 1139713 | 327709 | 0.98 | 3.3E-01 | 0 |
| Total lipids in very large HDL | 9 | 946993 | 1100265 | 797550 | 0.98 | 3.3E-01 | 0 |
| Lactate dehydrogenase | 5 | 1004188 | 1217050 | 617268 | 0.98 | 3.3E-01 | 0 |
| Parkinson Disease | 6 | 956171 | 1299603 | 637970 | 0.99 | 3.2E-01 | 0 |
| CCL4L1 - C-C motif chemokine 4-like | 1 | 1196450 | 1196450 | 1196450 | 0.99 | 3.2E-01 | 0 |
| Anxiety disorder (case/control) | 1 | 1201733 | 1201733 | 1201733 | 0.99 | 3.2E-01 | 0 |
| Illnesses of father: Bowel cancer | 3 | 1064808 | 1233548 | 842486 | 0.99 | 3.2E-01 | 0 |
| Pain type(s) experienced in last month: Knee pain | 8 | 942316 | 1020380 | 732324 | 0.99 | 3.2E-01 | 0 |
| 4th ventricle | 13 | 918940 | 1192940 | 158949 | 0.99 | 3.2E-01 | 0 |
| Nervous feelings (NERV-FEEL) | 35 | 835355 | 1175465 | 235547 | 0.99 | 3.2E-01 | 0 |
| Diagnoses - secondary ICD10: Z82 Family history of certain disabilities and chronic diseases (lea | 6 | 964495 | 1123575 | 380211 | 0.99 | 3.2E-01 | 0 |
| Illnesses of mother: High blood pressure | 17 | 892065 | 1073905 | 341380 | 1.00 | 3.2E-01 | 0 |
| TGFBI - Transforming growth factor-beta-induced protein ig-h3 | 1 | 1206840 | 1206840 | 1206840 | 1.00 | 3.2E-01 | 0 |
| Impedance measures - Basal metabolic rate | 583 | 713940 | 1157539 | 106956 | 1.00 | 3.2E-01 | 0 |
| Right isthmus cingulate | 3 | 1073430 | 1189656 | 992874 | 1.00 | 3.2E-01 | 0 |
| CFP - Properdin | 1 | 1217590 | 1217590 | 1217590 | 1.01 | 3.1E-01 | 0 |
| CAST - Calpastatin | 1 | 1201003 | 1201003 | 1201003 | 1.01 | 3.1E-01 | 0 |
| Phospholipids in very large HDL | 10 | 927768 | 1087011 | 598784 | 1.01 | 3.1E-01 | 0 |
| Rapid alternating stimulus test for letters/numbers | 1 | 1213078 | 1213078 | 1213078 | 1.01 | 3.1E-01 | 0 |
| MBL2 - Mannose-binding protein C | 2 | 1068624 | 1236331 | 900917 | 1.01 | 3.1E-01 | 0 |
| 3rd ventricle | 13 | 921338 | 1065428 | 398773 | 1.01 | 3.1E-01 | 0 |
| Hip circumference (female, adjusted for BMI) | 54 | 807940 | 1223229 | 189748 | 1.01 | 3.1E-01 | 0 |
| AGRP - Agouti-related protein | 1 | 1225853 | 1225853 | 1225853 | 1.02 | 3.1E-01 | 0 |
| Opioids | 2 | 1048161 | 1179488 | 916834 | 1.02 | 3.1E-01 | 0 |
| IL1RAP - Interleukin-1 Receptor accessory protein | 1 | 1231095 | 1231095 | 1231095 | 1.02 | 3.1E-01 | 0 |
| Rapid automised naming and rapid alternating stimulus test performance | 1 | 1213078 | 1213078 | 1213078 | 1.02 | 3.1E-01 | 0 |
| Illnesses of mother: Diabetes | 19 | 889053 | 1137050 | 314279 | 1.02 | 3.1E-01 | 0 |
| Insomnia (female) | 8 | 948676 | 1381783 | 76377 | 1.02 | 3.1E-01 | 0 |
| Number of sleep episodes | 19 | 889053 | 973685 | 247575 | 1.02 | 3.1E-01 | 0 |
| Calcium | 13 | 927923 | 1173788 | 602963 | 1.02 | 3.1E-01 | 0 |

| | | | | | | | |
|---|---|---|---|---|---|---|---|
| Right inferior parietal | 6 | 973244 | 1042853 | 896782 | 1.02 | 3.1E-01 | 0 |
| FCN2 - Ficolin-2 | 1 | 1217490 | 1217490 | 1217490 | 1.03 | 3.0E-01 | 0 |
| Rate of decline FEF25-75 (GEE, adjusted for multiple covariates) | 3 | 1081133 | 1268198 | 809241 | 1.03 | 3.0E-01 | 0 |
| Antiglaucoma preparations and miotics | 11 | 940400 | 1241610 | 149024 | 1.03 | 3.0E-01 | 0 |
| Never eat eggs, dairy, wheat, sugar: Sugar or foods/drinks containing sugar | 7 | 991630 | 1132256 | 486269 | 1.03 | 3.0E-01 | 0 |
| Height (female) | 52 | 807424 | 1156217 | 151947 | 1.03 | 3.0E-01 | 0 |
| Frequency of walking for pleasure in last 4 weeks | 5 | 1025960 | 1286093 | 972390 | 1.03 | 3.0E-01 | 0 |
| Amino acid::Valine, leucine and isoleucine metabolism::valine | 1 | 1226363 | 1226363 | 1226363 | 1.03 | 3.0E-01 | 0 |
| Low-density lipoprotein cholesterol (at exam 1, GEE, adjusted for age and sex) | 1 | 1214963 | 1214963 | 1214963 | 1.03 | 3.0E-01 | 0 |
| Right handed | 1 | 1233555 | 1233555 | 1233555 | 1.03 | 3.0E-01 | 0 |
| Symbol digit substitution test - Number of symbol digit matches made correctly | 4 | 1003465 | 1162194 | 775033 | 1.03 | 3.0E-01 | 0 |
| POMC - Beta-endorphin | 1 | 1234340 | 1234340 | 1234340 | 1.03 | 3.0E-01 | 0 |
| Cryptorchidism (group 2) | 5 | 1031778 | 1061213 | 78554 | 1.04 | 3.0E-01 | 0 |
| Irritability (IRR) | 35 | 844035 | 1222873 | 187028 | 1.04 | 3.0E-01 | 0 |
| Hemoglobin | 11 | 946020 | 1321773 | 266267 | 1.04 | 3.0E-01 | 0 |
| Diagnoses - secondary ICD10: I20 Angina pectoris | 12 | 921491 | 1185414 | 543718 | 1.04 | 3.0E-01 | 0 |
| Phosphatidylcholine and other cholines | 4 | 1008179 | 1067793 | 963848 | 1.04 | 3.0E-01 | 0 |
| Neck width (GEE, female, adjusted for age) | 2 | 1063098 | 1081666 | 1044529 | 1.04 | 3.0E-01 | 0 |
| Neuroblastoma | 5 | 1027838 | 1050213 | 787600 | 1.04 | 3.0E-01 | 0 |
| Monocyte chemoattractant protein 1 (at exam 7, GEE, adjusted for age and sex) | 1 | 1242235 | 1242235 | 1242235 | 1.05 | 2.9E-01 | 0 |
| Osteoarthritis of hit or knee | 22 | 874041 | 1222210 | 167860 | 1.05 | 2.9E-01 | 0 |
| Number of days/week walked 10+ minutes | 10 | 945119 | 1173691 | 89270 | 1.05 | 2.9E-01 | 0 |
| Baldness | 59 | 803585 | 1165716 | 433620 | 1.05 | 2.9E-01 | 0 |
| Total fatty acids | 9 | 970533 | 1097950 | 889053 | 1.05 | 2.9E-01 | 0 |
| Esterified cholesterol | 9 | 970533 | 1048933 | 385130 | 1.06 | 2.9E-01 | 0 |
| Monocyte chemoattractant protein 1 (at exam 7, FBAT, adjusted for multivariable) | 1 | 1242235 | 1242235 | 1242235 | 1.06 | 2.9E-01 | 0 |
| Amino acid::Valine, leucine and isoleucine metabolism::3-methyl-2-oxobutyrate | 1 | 1226363 | 1226363 | 1226363 | 1.06 | 2.9E-01 | 0 |
| Diagnoses - main ICD10: R07 Pain in throat and chest | 1 | 1246235 | 1246235 | 1246235 | 1.07 | 2.9E-01 | 0 |
| Mean FEV1/FVC from exam 3 and 5 (GEE, adjusted for multiple covariates) | 2 | 1078585 | 1172401 | 984769 | 1.07 | 2.9E-01 | 0 |
| Current tobacco smoking | 11 | 957335 | 1211318 | 260396 | 1.07 | 2.8E-01 | 0 |
| ::::X-12728 | 1 | 1244233 | 1244233 | 1244233 | 1.07 | 2.8E-01 | 0 |
| OmegaL6 fatty acids | 7 | 1001603 | 1073441 | 677831 | 1.07 | 2.8E-01 | 0 |
| Anterior corona radiata radial diusivities | 9 | 975900 | 1268678 | 466160 | 1.07 | 2.8E-01 | 0 |
| ::::X-09789 | 2 | 1078950 | 1149318 | 1008583 | 1.08 | 2.8E-01 | 0 |
| Ratio of visceral-tosubcutaneous adipose tissue volume | 3 | 1109360 | 1275428 | 607840 | 1.08 | 2.8E-01 | 0 |
| Types of physical activity in last 4 weeks: Heavy DIY (eg: weeding, lawn mowing, carpentry, diggi | 6 | 983833 | 1250692 | 846796 | 1.08 | 2.8E-01 | 0 |
| Gastric cancer | 4 | 1026389 | 1326258 | 716506 | 1.08 | 2.8E-01 | 0 |
| CCL28 - C-C motif chemokine 28 | 1 | 1241580 | 1241580 | 1241580 | 1.08 | 2.8E-01 | 0 |
| Carbohydrate::Glycolysis, gluconeogenesis, pyruvate metabolism::1,6-anhydroglucose | 1 | 1260335 | 1260335 | 1260335 | 1.08 | 2.8E-01 | 0 |
| Cutaneous T-cell attracting (CCL27) | 3 | 1103595 | 1213264 | 564904 | 1.08 | 2.8E-01 | 0 |
| Lipid::Carnitine metabolism::acetylcarnitine | 1 | 1255280 | 1255280 | 1255280 | 1.08 | 2.8E-01 | 0 |
| Hair colour (natural, before greying): Black | 148 | 762259 | 1158253 | 99273 | 1.08 | 2.8E-01 | 0 |
| Ratio of bisLallylic bonds to total fatty acids in lipids | 2 | 1083444 | 1358406 | 808482 | 1.09 | 2.8E-01 | 0 |

| | | | | | | | |
|---|---|---|---|---|---|---|---|
| Superior longitudinal fasciculus mode of anisotropy | 1 | 1255143 | 1255143 | 1255143 | 1.09 | 2.8E-01 | 0 |
| High-density lipoprotein - Birth cohorts | 4 | 1025219 | 1240917 | 912017 | 1.09 | 2.7E-01 | 0 |
| Illnesses of father: Chronic bronchitis/emphysema | 4 | 1026484 | 1485056 | 431966 | 1.09 | 2.7E-01 | 0 |
| Illnesses of father: High blood pressure | 9 | 985248 | 1091518 | 479195 | 1.09 | 2.7E-01 | 0 |
| Lipid::Carnitine metabolism::2-tetradecenoyl carnitine | 1 | 1255280 | 1255280 | 1255280 | 1.09 | 2.7E-01 | 0 |
| Amino acid::Guanidino and acetamido metabolism::4-acetamidobutanoate | 2 | 1078410 | 1149521 | 1007299 | 1.10 | 2.7E-01 | 0 |
| Doctor diagnosed hayfever or allergic rhinitis | 21 | 896303 | 1174765 | 476268 | 1.10 | 2.7E-01 | 0 |
| Right inferior temporal | 2 | 1086555 | 1407824 | 765286 | 1.10 | 2.7E-01 | 0 |
| Hematocrit | 16 | 913133 | 1317203 | 374489 | 1.10 | 2.7E-01 | 0 |
| ::::X-10395 | 1 | 1267410 | 1267410 | 1267410 | 1.10 | 2.7E-01 | 0 |
| Mean platelet volume (two-way meta) | 147 | 763848 | 1170245 | 323021 | 1.11 | 2.7E-01 | 0 |
| ::::X-10429 | 1 | 1267410 | 1267410 | 1267410 | 1.11 | 2.7E-01 | 0 |
| Lipid::Carnitine metabolism::decanoylcarnitine | 3 | 1112685 | 1138233 | 825368 | 1.11 | 2.7E-01 | 0 |
| RARRES2 - Retinoic acid receptor responder protein 2 | 1 | 1266463 | 1266463 | 1266463 | 1.11 | 2.7E-01 | 0 |
| Amount of alcohol drunk on a typical drinking day | 6 | 999435 | 1344179 | 841460 | 1.12 | 2.6E-01 | 0 |
| ERAP1 - Endoplasmic reticulum aminopeptidase 1 | 1 | 1264990 | 1264990 | 1264990 | 1.12 | 2.6E-01 | 0 |
| ASAH2 - Neutral ceramidase | 1 | 1275198 | 1275198 | 1275198 | 1.12 | 2.6E-01 | 0 |
| Amino acid::Cysteine, methionine, SAM, taurine metabolism::methionine | 1 | 1270505 | 1270505 | 1270505 | 1.12 | 2.6E-01 | 0 |
| C-reactive protein - Birth cohorts | 2 | 1100296 | 1377998 | 822594 | 1.12 | 2.6E-01 | 0 |
| IGFBP7 - Insulin-like growth factor-binding protein 7 | 1 | 1272163 | 1272163 | 1272163 | 1.12 | 2.6E-01 | 0 |
| Total phosphoglycerides | 6 | 1008179 | 1111728 | 950479 | 1.12 | 2.6E-01 | 0 |
| Parkinson's disease age at onset | 1 | 1275658 | 1275658 | 1275658 | 1.13 | 2.6E-01 | 0 |
| Epstein Barr Virus antigen EBNA seropositivity | 1 | 1266218 | 1266218 | 1266218 | 1.13 | 2.6E-01 | 0 |
| Left cerebellum white matter | 11 | 977470 | 1141903 | 675613 | 1.13 | 2.6E-01 | 0 |
| Resistin (GEE, adjusted for age and sex) | 3 | 1133978 | 1200203 | 861174 | 1.13 | 2.6E-01 | 0 |
| High light scatter percentage of red cells (two-way meta) | 103 | 783420 | 1135919 | 194959 | 1.14 | 2.6E-01 | 0 |
| IMPDH1 - Inosine-5'-monophosphate dehydrogenase 1 | 3 | 1129118 | 1327356 | 1095680 | 1.14 | 2.6E-01 | 0 |
| CD109 - CD109 antigen | 1 | 1288055 | 1288055 | 1288055 | 1.14 | 2.6E-01 | 0 |
| Nervous feelings | 30 | 868378 | 1199436 | 373306 | 1.14 | 2.6E-01 | 0 |
| Treatment/medication code: atenolol | 7 | 1027155 | 1152028 | 631976 | 1.14 | 2.5E-01 | 0 |
| Neutrophil count (two-way meta) | 68 | 806130 | 1222529 | 97643 | 1.14 | 2.5E-01 | 0 |
| Illnesses of siblings: Diabetes | 7 | 1027155 | 1168869 | 957019 | 1.14 | 2.5E-01 | 0 |
| Alanine aminotransferase | 21 | 905515 | 1113980 | 392805 | 1.14 | 2.5E-01 | 0 |
| Reticulocyte count (two-way meta) | 110 | 778138 | 1085710 | 132467 | 1.14 | 2.5E-01 | 0 |
| Neovascular disease | 10 | 966043 | 1142473 | 580831 | 1.15 | 2.5E-01 | 0 |
| Type 2 Diabetes (adjusted for BMI) | 153 | 763233 | 1077243 | 353550 | 1.15 | 2.5E-01 | 0 |
| Waist circumference (male) | 23 | 898940 | 1300483 | 499163 | 1.15 | 2.5E-01 | 0 |
| Schizophrenia | 193 | 755908 | 1223613 | 128092 | 1.16 | 2.5E-01 | 0 |
| Cigarettes per day | 19 | 922358 | 1098394 | 322805 | 1.16 | 2.5E-01 | 0 |
| Nucleotide::Purine metabolism, urate metabolism::urate | 1 | 1294678 | 1294678 | 1294678 | 1.16 | 2.5E-01 | 0 |
| Lipid::Long chain fatty acid::10-nonadecenoate (19:1n9) | 1 | 1301655 | 1301655 | 1301655 | 1.17 | 2.4E-01 | 0 |
| Antimigraine preparations | 12 | 956521 | 1305876 | 419438 | 1.17 | 2.4E-01 | 0 |
| Aspartate aminotransferase (at exam 2, GEE, adjusted for age and sex) | 1 | 1311795 | 1311795 | 1311795 | 1.17 | 2.4E-01 | 0 |

| | | | | | | | |
|---|---|---|---|---|---|---|---|
| Cereal intake | 27 | 889053 | 1246224 | 83385 | 1.17 | 2.4E-01 | 0 |
| Growth regulated oncogene (CXCL1) | 2 | 1120801 | 1195424 | 1046178 | 1.17 | 2.4E-01 | 0 |
| ::::X-12704 | 1 | 1285133 | 1285133 | 1285133 | 1.17 | 2.4E-01 | 0 |
| Illnesses of mother: Heart disease | 2 | 1122575 | 1258228 | 986923 | 1.18 | 2.4E-01 | 0 |
| Male-specific factors - Hair/balding pattern: Pattern 1 | 166 | 764678 | 1093732 | 345728 | 1.18 | 2.4E-01 | 0 |
| DAPK2 - Death-associated protein kinase 2 | 1 | 1305810 | 1305810 | 1305810 | 1.19 | 2.4E-01 | 0 |
| Lipid::Long chain fatty acid::oleate (18:1n9) | 1 | 1301655 | 1301655 | 1301655 | 1.19 | 2.4E-01 | 0 |
| Neuroticism (univariate) | 72 | 807849 | 1108384 | 503082 | 1.19 | 2.3E-01 | 0 |
| Fed-up feelings | 27 | 889275 | 1346724 | 93923 | 1.19 | 2.3E-01 | 0 |
| Paternal history of Alzheimer's disease | 2 | 1126449 | 1245147 | 1007751 | 1.19 | 2.3E-01 | 0 |
| CXCL5 - C-X-C motif chemokine 5 | 1 | 1298125 | 1298125 | 1298125 | 1.19 | 2.3E-01 | 0 |
| Mean corpuscular volume (three-way meta) | 162 | 765151 | 1153711 | 202809 | 1.19 | 2.3E-01 | 0 |
| Mental distress - Ever sought or received professional help for mental distress | 2 | 1129539 | 1134594 | 1124483 | 1.19 | 2.3E-01 | 0 |
| Low-density lipoprotein - Birth cohorts | 3 | 1152513 | 1276488 | 940048 | 1.19 | 2.3E-01 | 0 |
| Excessive hairiness | 3 | 1145635 | 1332681 | 912813 | 1.19 | 2.3E-01 | 0 |
| Total lipids in IDL | 13 | 970533 | 1055113 | 385130 | 1.20 | 2.3E-01 | 0 |
| Irritability | 34 | 869086 | 1230843 | 195670 | 1.20 | 2.3E-01 | 0 |
| SPINT2 - Kunitz-type protease inhibitor 2 | 1 | 1313543 | 1313543 | 1313543 | 1.21 | 2.3E-01 | 0 |
| Right fusiform | 1 | 1317115 | 1317115 | 1317115 | 1.21 | 2.3E-01 | 0 |
| HMG CoA reductase inhibitors | 58 | 826156 | 1142679 | 194463 | 1.21 | 2.3E-01 | 0 |
| Strenuous sports or other exercises | 12 | 966571 | 1341801 | 557027 | 1.21 | 2.3E-01 | 0 |
| High light scatter reticulocyte count (two-way meta) | 98 | 791799 | 1131342 | 261510 | 1.21 | 2.2E-01 | 0 |
| Left entorhinal | 4 | 1068055 | 1685108 | 392550 | 1.22 | 2.2E-01 | 0 |
| Free cholesterol in IDL | 11 | 996508 | 1132234 | 84490 | 1.22 | 2.2E-01 | 0 |
| Autism spectrum disorder | 3 | 1164070 | 1190443 | 970190 | 1.22 | 2.2E-01 | 0 |
| Coronary artery disease in diabetes | 1 | 1335308 | 1335308 | 1335308 | 1.23 | 2.2E-01 | 0 |
| C7 - Complement component C7 | 2 | 1140420 | 1208021 | 1072819 | 1.23 | 2.2E-01 | 0 |
| IL6ST - Interleukin-6 receptor subunit beta | 1 | 1329585 | 1329585 | 1329585 | 1.23 | 2.2E-01 | 0 |
| Daytime dozing / sleeping (narcolepsy) | 26 | 897286 | 1134078 | 242213 | 1.23 | 2.2E-01 | 0 |
| Ever smoked regulary | 100 | 793695 | 1193218 | 160938 | 1.23 | 2.2E-01 | 0 |
| Sleep sedentary | 5 | 1102738 | 1605853 | 759793 | 1.23 | 2.2E-01 | 0 |
| Triglycerides in IDL | 12 | 976043 | 1106157 | 705657 | 1.23 | 2.2E-01 | 0 |
| Trail making test - Duration to complete alphanumeric path (trail #2) | 8 | 1011706 | 1550558 | 847574 | 1.23 | 2.2E-01 | 0 |
| Heel bone mineral density | 308 | 741698 | 1114096 | 149482 | 1.23 | 2.2E-01 | 0 |
| CST7 - Cystatin-F | 1 | 1331835 | 1331835 | 1331835 | 1.24 | 2.2E-01 | 0 |
| Cingulum (hippocampus) radial diusivities | 1 | 1347053 | 1347053 | 1347053 | 1.24 | 2.2E-01 | 0 |
| Peptide::Dipeptide::X-14205--alpha-glutamyltyrosine | 2 | 1143096 | 1212176 | 1074017 | 1.24 | 2.2E-01 | 0 |
| Xenobiotics::Xanthine metabolism::1-methylurate | 1 | 1346083 | 1346083 | 1346083 | 1.24 | 2.1E-01 | 0 |
| Job involves heavy manual or physical work | 8 | 1018508 | 1274003 | 561957 | 1.24 | 2.1E-01 | 0 |
| Amino acid::Butanoate metabolism::2-aminobutyrate | 2 | 1149435 | 1207090 | 1091780 | 1.25 | 2.1E-01 | 0 |
| CXCL11 - C-X-C motif chemokine 11 | 1 | 1336048 | 1336048 | 1336048 | 1.25 | 2.1E-01 | 0 |
| Taking other prescription medications | 10 | 994278 | 1603742 | 489303 | 1.26 | 2.1E-01 | 0 |
| Standing height | 876 | 715844 | 1114119 | 124907 | 1.26 | 2.1E-01 | 0 |

| | | | | | | | |
|---|---|---|---|---|---|---|---|
| Free cholesterol in very large HDL | 10 | 997120 | 1266549 | 669396 | 1.26 | 2.1E-01 | 0 |
| Skin fluorescence | 1 | 1346083 | 1346083 | 1346083 | 1.27 | 2.1E-01 | 0 |
| Amino acid::Urea cycle; arginine-, proline-, metabolism::proline | 2 | 1151220 | 1207369 | 1095071 | 1.27 | 2.0E-01 | 0 |
| Bread intake | 16 | 951480 | 1685187 | 120441 | 1.27 | 2.0E-01 | 0 |
| Right accumbens area | 6 | 1055123 | 1240547 | 763986 | 1.27 | 2.0E-01 | 0 |
| Non-cancer illness code, self-reported: asthma | 51 | 848655 | 1136361 | 228042 | 1.27 | 2.0E-01 | 0 |
| Male-specific factors - Hair/balding pattern: Pattern 3 | 40 | 864370 | 1164823 | 210260 | 1.27 | 2.0E-01 | 0 |
| Insulin sensitivity index (adjusted for age, sex. Bmi) | 1 | 1353228 | 1353228 | 1353228 | 1.28 | 2.0E-01 | 0 |
| CXCL1 - Growth-regulated alpha protein | 1 | 1355818 | 1355818 | 1355818 | 1.28 | 2.0E-01 | 0 |
| Diagnoses - main ICD10: H26 Other cataract | 6 | 1048938 | 1397449 | 342022 | 1.28 | 2.0E-01 | 0 |
| NAGK - N-acetyl-D-glucosamine kinase | 1 | 1346420 | 1346420 | 1346420 | 1.28 | 2.0E-01 | 0 |
| Coffee type: Ground coffee (include espresso, filter etc) | 17 | 960338 | 1291558 | 407690 | 1.29 | 2.0E-01 | 0 |
| Peptide::Dipeptide::X-12244--N-acetylcarnosine | 3 | 1191685 | 1234166 | 1125289 | 1.29 | 2.0E-01 | 0 |
| Bipolar disorder | 12 | 984339 | 1212962 | 495684 | 1.30 | 2.0E-01 | 0 |
| Impedance measures - Leg fat-free mass (left) | 515 | 728210 | 1145316 | 113642 | 1.30 | 1.9E-01 | 0 |
| Plateletcrit (two-way meta) | 129 | 786968 | 1097068 | 338498 | 1.30 | 1.9E-01 | 0 |
| White matter | 10 | 1017278 | 1266022 | 618400 | 1.30 | 1.9E-01 | 0 |
| DPT - Dermatopontin | 1 | 1368958 | 1368958 | 1368958 | 1.30 | 1.9E-01 | 0 |
| Energy::Krebs cycle::citrate | 4 | 1108011 | 1261512 | 879573 | 1.30 | 1.9E-01 | 0 |
| PDK1 - [Pyruvate dehydrogenase (acetyl-transferring)] kinase isozyme 1, mitochondrial | 1 | 1372680 | 1372680 | 1372680 | 1.30 | 1.9E-01 | 0 |
| ::::X-12556 | 1 | 1361353 | 1361353 | 1361353 | 1.31 | 1.9E-01 | 0 |
| Duration of walks | 5 | 1121600 | 1341578 | 757313 | 1.31 | 1.9E-01 | 0 |
| Superior fronto-occipital fasciculus axial diusivities | 1 | 1381228 | 1381228 | 1381228 | 1.31 | 1.9E-01 | 0 |
| Ever smoked | 41 | 869535 | 1255143 | 96500 | 1.31 | 1.9E-01 | 0 |
| NKeff:%314-R7- | 1 | 1371008 | 1371008 | 1371008 | 1.32 | 1.9E-01 | 0 |
| Neuroticism score | 72 | 823763 | 1246051 | 393650 | 1.32 | 1.9E-01 | 0 |
| Blood clot, DVT, bronchitis, emphysema, asthma, rhinitis, eczema, allergy diagnosed by doctor: A | 66 | 830396 | 1120137 | 217690 | 1.32 | 1.9E-01 | 0 |
| Leptin (adjusted for BMI) | 2 | 1159548 | 1169260 | 1149835 | 1.32 | 1.9E-01 | 0 |
| Anilides | 7 | 1088348 | 1271423 | 553504 | 1.32 | 1.9E-01 | 0 |
| Mania - Ever had period extreme irritability | 1 | 1360448 | 1360448 | 1360448 | 1.33 | 1.8E-01 | 0 |
| Sagittal stratum fractional anisotropy | 6 | 1070869 | 1154893 | 876299 | 1.33 | 1.8E-01 | 0 |
| Xenobiotics::Drug::metoprolol acid metabolite* | 1 | 1381228 | 1381228 | 1381228 | 1.33 | 1.8E-01 | 0 |
| Types of physical activity in last 4 weeks: Strenuous sports | 4 | 1107429 | 1337814 | 688202 | 1.33 | 1.8E-01 | 0 |
| Impedance measures - Trunk fat percentage | 352 | 744063 | 1170616 | 102215 | 1.33 | 1.8E-01 | 0 |
| IDUA - alpha-L-iduronidase | 1 | 1390215 | 1390215 | 1390215 | 1.33 | 1.8E-01 | 0 |
| NKearly:%335+314- | 1 | 1371008 | 1371008 | 1371008 | 1.33 | 1.8E-01 | 0 |
| TEK - Angiopoietin-1 receptor, soluble | 1 | 1386483 | 1386483 | 1386483 | 1.33 | 1.8E-01 | 0 |
| Left caudate | 8 | 1049194 | 1116236 | 901826 | 1.34 | 1.8E-01 | 0 |
| Illnesses of mother: Severe depression | 1 | 1401083 | 1401083 | 1401083 | 1.34 | 1.8E-01 | 0 |
| Sum eosinophil basophil count (three-way meta) | 125 | 791080 | 1089978 | 159373 | 1.34 | 1.8E-01 | 0 |
| Diuretics | 72 | 827005 | 1110203 | 341408 | 1.35 | 1.8E-01 | 0 |
| Non-cancer illness code, self-reported: high cholesterol | 48 | 860018 | 1209234 | 392902 | 1.35 | 1.8E-01 | 0 |
| Amino acid::Phenylalanine & tyrosine metabolism::tyrosine | 3 | 1217793 | 1366889 | 864193 | 1.35 | 1.8E-01 | 0 |

| | | | | | | | |
|---|---|---|---|---|---|---|---|
| Femoral Neck BMD (males) | 1 | 1384985 | 1384985 | 1384985 | 1.35 | 1.8E-01 | 0 |
| Average weekly red wine intake | 5 | 1131905 | 1158533 | 831242 | 1.35 | 1.8E-01 | 0 |
| Illnesses of mother: Alzheimer's disease/dementia | 2 | 1183306 | 1273576 | 1093037 | 1.35 | 1.8E-01 | 0 |
| Maternal history of Alzheimer's disease | 2 | 1183306 | 1273576 | 1093037 | 1.36 | 1.8E-01 | 0 |
| Length of mobile phone use | 21 | 948435 | 1247098 | 119511 | 1.36 | 1.8E-01 | 0 |
| Right pericalcarine | 3 | 1220193 | 1407921 | 1160199 | 1.36 | 1.7E-01 | 0 |
| Heart rate increase | 15 | 997005 | 1280579 | 671900 | 1.36 | 1.7E-01 | 0 |
| Posterior limb of internal capsule radial diusivities | 8 | 1050655 | 1283801 | 532716 | 1.36 | 1.7E-01 | 0 |
| RET - Proto-oncogene tyrosine-protein kinase receptor Ret | 1 | 1393185 | 1393185 | 1393185 | 1.36 | 1.7E-01 | 0 |
| Vasodilators used in cardiac diseases | 1 | 1393880 | 1393880 | 1393880 | 1.36 | 1.7E-01 | 0 |
| C-reactive protein | 14 | 988990 | 1190721 | 318864 | 1.37 | 1.7E-01 | 0 |
| Lean body mass (whole body) | 1 | 1408560 | 1408560 | 1408560 | 1.37 | 1.7E-01 | 0 |
| Mood swings | 36 | 888069 | 1397594 | 231215 | 1.37 | 1.7E-01 | 0 |
| Estimated BMD | 194 | 768539 | 1158503 | 209888 | 1.37 | 1.7E-01 | 0 |
| Impedance measures - Body fat percentage | 389 | 741980 | 1148463 | 108235 | 1.37 | 1.7E-01 | 0 |
| Corticospinal tract mean diusivities | 2 | 1196910 | 1206223 | 1187598 | 1.37 | 1.7E-01 | 0 |
| Fractures | 16 | 976246 | 1276098 | 548429 | 1.38 | 1.7E-01 | 0 |
| Lumbar Spine BMD (females) | 20 | 953276 | 1213724 | 385099 | 1.38 | 1.7E-01 | 0 |
| Neuroticism sum score | 92 | 812364 | 1268068 | 187236 | 1.38 | 1.7E-01 | 0 |
| Uncinate fasciculus mode of anisotropy | 1 | 1409495 | 1409495 | 1409495 | 1.38 | 1.7E-01 | 0 |
| Loneliness | 12 | 1010251 | 1348374 | 156278 | 1.39 | 1.7E-01 | 0 |
| Diastolic Blood Pressure | 24 | 935636 | 1244447 | 407697 | 1.39 | 1.6E-01 | 0 |
| Illnesses of father: Diabetes | 19 | 975063 | 1239001 | 472826 | 1.39 | 1.6E-01 | 0 |
| ::::X-13671 | 1 | 1432335 | 1432335 | 1432335 | 1.40 | 1.6E-01 | 0 |
| Posterior corona radiata mean diusivities | 1 | 1423275 | 1423275 | 1423275 | 1.40 | 1.6E-01 | 0 |
| Lipid::Long chain fatty acid::stearate (18:0) | 2 | 1207385 | 1254520 | 1160250 | 1.40 | 1.6E-01 | 0 |
| MMSE score (at age 65, GEE, adjusted for multivariable) | 3 | 1238728 | 1355005 | 915233 | 1.40 | 1.6E-01 | 0 |
| Treatment/medication code: amlodipine | 15 | 1005775 | 1356610 | 769329 | 1.40 | 1.6E-01 | 0 |
| Amino acid::Valine, leucine and isoleucine metabolism::2-hydroxyisobutyrate | 3 | 1233350 | 1380466 | 1152186 | 1.40 | 1.6E-01 | 0 |
| Risk taking | 16 | 982553 | 1272574 | 294044 | 1.40 | 1.6E-01 | 0 |
| Hip circumference (adjusted for BMI) | 102 | 810285 | 1126301 | 248288 | 1.41 | 1.6E-01 | 0 |
| Childhood Absence Epilepsy | 2 | 1201125 | 1328860 | 1073390 | 1.41 | 1.6E-01 | 0 |
| Weekly usage of mobile phone in last 3 months | 8 | 1059716 | 1351618 | 1000390 | 1.41 | 1.6E-01 | 0 |
| Mood swings (MOOD) | 40 | 888069 | 1370185 | 132709 | 1.41 | 1.6E-01 | 0 |
| TNFSF8 - Tumor necrosis factor ligand superfamily member 8 | 1 | 1440178 | 1440178 | 1440178 | 1.42 | 1.6E-01 | 0 |
| Glucose - Birth cohorts | 1 | 1422368 | 1422368 | 1422368 | 1.42 | 1.6E-01 | 0 |
| Long-standing illness, disability or infirmity | 12 | 1021243 | 1503525 | 508155 | 1.42 | 1.6E-01 | 0 |
| ECM1 - Extracellular matrix protein 1 | 1 | 1438543 | 1438543 | 1438543 | 1.42 | 1.6E-01 | 0 |
| NID1 - Nidogen-1 | 1 | 1427540 | 1427540 | 1427540 | 1.42 | 1.5E-01 | 0 |
| Pericardial adipose tissue volume | 2 | 1218301 | 1348081 | 1088522 | 1.42 | 1.5E-01 | 0 |
| Non-cancer illness code, self-reported: hypertension | 151 | 788825 | 1139530 | 161005 | 1.43 | 1.5E-01 | 0 |
| Impedance measures - Leg predicted mass (left) | 513 | 735633 | 1170118 | 114349 | 1.43 | 1.5E-01 | 0 |
| Waist-hip ratio (female, adjusted for BMI) | 42 | 884941 | 1215864 | 470843 | 1.43 | 1.5E-01 | 0 |

| | | | | | | | |
|---|---|---|---|---|---|---|---|
| Amino acid::Urea cycle; arginine-, proline-, metabolism::dimethylarginine (SDMA + ADMA) | 1 | 1443173 | 1443173 | 1443173 | 1.44 | 1.5E-01 | 0 |
| Vascular/heart problems diagnosed by doctor: High blood pressure | 189 | 776138 | 1124403 | 174418 | 1.44 | 1.5E-01 | 0 |
| Lumbar Spine BMD | 38 | 894429 | 1223609 | 403358 | 1.44 | 1.5E-01 | 0 |
| Uric acid | 21 | 963458 | 1223408 | 70411 | 1.44 | 1.5E-01 | 0 |
| PR interval | 7 | 1119863 | 1212410 | 889086 | 1.44 | 1.5E-01 | 0 |
| Milk type used: Full cream | 4 | 1148098 | 1407964 | 742005 | 1.44 | 1.5E-01 | 0 |
| Loneliness, isolation | 4 | 1141716 | 1363014 | 813379 | 1.45 | 1.5E-01 | 0 |
| IL7R - Interleukin-7 receptor subunit alpha | 1 | 1439100 | 1439100 | 1439100 | 1.45 | 1.5E-01 | 0 |
| Low-density lipoprotein cholesterol | 77 | 833758 | 1278630 | 471527 | 1.45 | 1.5E-01 | 0 |
| Ease of getting up in the morning | 53 | 866553 | 1234990 | 187736 | 1.45 | 1.5E-01 | 0 |
| NK:%Eff | 1 | 1454618 | 1454618 | 1454618 | 1.45 | 1.5E-01 | 0 |
| Depressive affect subcluster | 58 | 855294 | 1307893 | 220314 | 1.45 | 1.5E-01 | 0 |
| IGF2R - Cation-independent mannose-6-phosphate receptor | 1 | 1455465 | 1455465 | 1455465 | 1.47 | 1.4E-01 | 0 |
| Right insula | 5 | 1176590 | 1426355 | 854958 | 1.47 | 1.4E-01 | 0 |
| Amyotrophic lateral sclerosis (logistic) | 3 | 1263588 | 1334358 | 675340 | 1.47 | 1.4E-01 | 0 |
| Estimated bone mineral density from heel ultrasounds | 517 | 736918 | 1128618 | 201271 | 1.47 | 1.4E-01 | 0 |
| Vitamin and mineral supplements: Vitamin D | 1 | 1466193 | 1466193 | 1466193 | 1.47 | 1.4E-01 | 0 |
| FUT3 - Galactoside 3(4)-L-fucosyltransferase | 1 | 1464083 | 1464083 | 1464083 | 1.47 | 1.4E-01 | 0 |
| Valine | 3 | 1264745 | 1382799 | 1153795 | 1.48 | 1.4E-01 | 0 |
| Diagnoses - secondary ICD10: I10 Essential (primary) hypertension | 82 | 831810 | 1139037 | 359159 | 1.48 | 1.4E-01 | 0 |
| CD163 - Scavenger receptor cysteine-rich type 1 protein M130 | 1 | 1464448 | 1464448 | 1464448 | 1.49 | 1.4E-01 | 0 |
| Tense / 'highly strung' (TENSE) | 20 | 976814 | 1202053 | 345118 | 1.49 | 1.4E-01 | 0 |
| Adenocarcinoma (adjusted for age) | 4 | 1160283 | 1346019 | 942873 | 1.49 | 1.4E-01 | 0 |
| High-density lipoprotein 3 cholesterol (at exam 5, GEE, adjusted for multivariable) | 1 | 1455263 | 1455263 | 1455263 | 1.49 | 1.4E-01 | 0 |
| CD8:%39+ | 1 | 1480198 | 1480198 | 1480198 | 1.50 | 1.3E-01 | 0 |
| Reticulocyte fraction of red cells (two-way meta) | 108 | 816821 | 1102706 | 264560 | 1.50 | 1.3E-01 | 0 |
| Plateletcrit (three-way meta) | 163 | 786968 | 1148721 | 331799 | 1.51 | 1.3E-01 | 0 |
| Ankylosing spondylitis | 16 | 1009141 | 1215242 | 412496 | 1.51 | 1.3E-01 | 0 |
| Alcohol consumption | 1 | 1484388 | 1484388 | 1484388 | 1.51 | 1.3E-01 | 0 |
| Neck/shoulder pain for 3+ months | 1 | 1477533 | 1477533 | 1477533 | 1.51 | 1.3E-01 | 0 |
| Mean corpuscular hemoglobin concentration | 35 | 924508 | 1191553 | 170161 | 1.52 | 1.3E-01 | 0 |
| Frequency of unenthusiasm / disinterest in last 2 weeks | 4 | 1168816 | 1387924 | 801984 | 1.52 | 1.3E-01 | 0 |
| Chronotype | 168 | 788309 | 1169536 | 120038 | 1.52 | 1.3E-01 | 0 |
| High frequency power HRV (cohort exam 18 offsprint exam 3, GEE, adjusted for age and HR) | 1 | 1491978 | 1491978 | 1491978 | 1.52 | 1.3E-01 | 0 |
| Bread type: White | 13 | 1051725 | 1229118 | 114657 | 1.52 | 1.3E-01 | 0 |
| Right caudate | 6 | 1129000 | 1171759 | 1021284 | 1.53 | 1.3E-01 | 0 |
| Lipid::Inositol metabolism::myo-inositol | 2 | 1247600 | 1248508 | 1246693 | 1.53 | 1.3E-01 | 0 |
| Illnesses of siblings: High blood pressure | 15 | 1035558 | 1242953 | 388766 | 1.53 | 1.3E-01 | 0 |
| SAA1 - Serum amyloid A-1 protein | 1 | 1485705 | 1485705 | 1485705 | 1.54 | 1.2E-01 | 0 |
| CD8:%SCM | 1 | 1489373 | 1489373 | 1489373 | 1.54 | 1.2E-01 | 0 |
| Male pattern baldness (BOLT LMM infinitesimal mixed model) | 326 | 757014 | 1093209 | 242427 | 1.54 | 1.2E-01 | 0 |
| Residual from predicted FEF25-75 for latest exam (GEE, adjusted for multiple covariates) | 3 | 1295718 | 1347703 | 950311 | 1.54 | 1.2E-01 | 0 |
| IL19 - Interleukin-19 | 2 | 1248780 | 1609211 | 888349 | 1.55 | 1.2E-01 | 0 |

| | | | | | | | |
|---|---|---|---|---|---|---|---|
| Lipid::Medium chain fatty acid::heptanoate (7:0) | 2 | 1267661 | 1356518 | 1178804 | 1.56 | 1.2E-01 | 0 |
| Impedance measures - Leg predicted mass (right) | 525 | 740235 | 1158053 | 111811 | 1.56 | 1.2E-01 | 0 |
| ::::X-12850 | 2 | 1267595 | 1350481 | 1184709 | 1.56 | 1.2E-01 | 0 |
| Breastfed as a baby | 1 | 1504390 | 1504390 | 1504390 | 1.56 | 1.2E-01 | 0 |
| CCL5 - C-C motif chemokine 5 | 1 | 1496075 | 1496075 | 1496075 | 1.57 | 1.2E-01 | 0 |
| Neuroticism | 152 | 798790 | 1187466 | 226861 | 1.57 | 1.2E-01 | 0 |
| Waist-hip ratio (female <= 50 yrs, adjusted for BMI) | 9 | 1131123 | 1348585 | 792000 | 1.57 | 1.2E-01 | 0 |
| Non-cancer illness code, self-reported: eczema/dermatitis | 12 | 1063461 | 1186424 | 731673 | 1.57 | 1.2E-01 | 0 |
| ::::X-12696 | 2 | 1265263 | 1684281 | 846244 | 1.57 | 1.2E-01 | 0 |
| Right putamen | 10 | 1084045 | 1411990 | 235232 | 1.58 | 1.1E-01 | 0 |
| Amyotrophic lateral sclerosis (linear mixed model) | 5 | 1224425 | 1263588 | 1174275 | 1.58 | 1.1E-01 | 0 |
| Ratio of visceral-tosubcutaneous adipose tissue volume (female) | 1 | 1505643 | 1505643 | 1505643 | 1.58 | 1.1E-01 | 0 |
| Leg pain on walking | 1 | 1525045 | 1525045 | 1525045 | 1.58 | 1.1E-01 | 0 |
| Impedance measures - Arm fat-free mass (left) | 553 | 740420 | 1170118 | 104035 | 1.58 | 1.1E-01 | 0 |
| AGT - Angiotensinogen | 1 | 1516068 | 1516068 | 1516068 | 1.59 | 1.1E-01 | 0 |
| Shaft Section Modulus (GEE, female, adjussted for multivariable) | 1 | 1527753 | 1527753 | 1527753 | 1.59 | 1.1E-01 | 0 |
| Drinks per day | 28 | 952556 | 1272131 | 87686 | 1.59 | 1.1E-01 | 0 |
| Lipid::Medium chain fatty acid::laurate (12:0) | 2 | 1276010 | 1437755 | 1114265 | 1.60 | 1.1E-01 | 0 |
| General risk tolerance | 24 | 978341 | 1113830 | 548938 | 1.60 | 1.1E-01 | 0 |
| Body Mass Index (Dominance deviation model) | 1 | 1527753 | 1527753 | 1527753 | 1.60 | 1.1E-01 | 0 |
| Myeloid white cell count (two-way meta) | 82 | 845603 | 1210711 | 119599 | 1.61 | 1.1E-01 | 0 |
| Lipid::Bile acid metabolism::deoxycholate | 1 | 1529150 | 1529150 | 1529150 | 1.61 | 1.1E-01 | 0 |
| Fluid intelligence test - Number of fluid intelligence questions attempted within time limit | 9 | 1137883 | 1319015 | 160094 | 1.61 | 1.1E-01 | 0 |
| Hip circumference (male, adjusted for BMI) | 38 | 922968 | 1252582 | 141925 | 1.62 | 1.1E-01 | 0 |
| Lipid::Carnitine metabolism::hexanoylcarnitine | 4 | 1200301 | 1352965 | 830523 | 1.62 | 1.1E-01 | 0 |
| Diagnoses - secondary ICD10: Z72 Problems related to lifestyle | 1 | 1529998 | 1529998 | 1529998 | 1.62 | 1.1E-01 | 0 |
| RELT - Tumor necrosis factor receptor superfamily member 19L | 1 | 1537180 | 1537180 | 1537180 | 1.62 | 1.1E-01 | 0 |
| Biliary atresia | 3 | 1330175 | 1602635 | 1157248 | 1.62 | 1.0E-01 | 0 |
| TNFAIP6 - Tumor necrosis factor-inducible gene 6 protein | 1 | 1543070 | 1543070 | 1543070 | 1.63 | 1.0E-01 | 0 |
| Alcohol intake versus 10 years previously | 10 | 1101795 | 1591109 | 794321 | 1.63 | 1.0E-01 | 0 |
| Alzheimer's disease | 11 | 1109820 | 1196119 | 576701 | 1.63 | 1.0E-01 | 0 |
| Galactosylation | 8 | 1131278 | 1199662 | 845010 | 1.64 | 1.0E-01 | 0 |
| Arms-arm fat ratio (male) | 34 | 942438 | 1231051 | 222527 | 1.64 | 1.0E-01 | 0 |
| Frequency of tiredness / lethargy in last 2 weeks | 21 | 1011308 | 1222053 | 398010 | 1.64 | 1.0E-01 | 0 |
| CNTN5 - Contactin-5 | 1 | 1536328 | 1536328 | 1536328 | 1.64 | 1.0E-01 | 0 |
| Adiponectin level | 11 | 1111730 | 1281746 | 589694 | 1.64 | 1.0E-01 | 0 |
| PLAU - Urokinase-type plasminogen activator | 1 | 1547740 | 1547740 | 1547740 | 1.65 | 1.0E-01 | 0 |
| Treatment/medication code: glucosamine product | 1 | 1558458 | 1558458 | 1558458 | 1.65 | 1.0E-01 | 0 |
| Generalised epilepsy | 10 | 1098946 | 1325679 | 772659 | 1.65 | 9.9E-02 | 0 |
| Transport type for commuting to job workplace: Cycle | 1 | 1559753 | 1559753 | 1559753 | 1.65 | 9.8E-02 | 0 |
| Left inferior parietal | 4 | 1214493 | 1386809 | 799920 | 1.66 | 9.7E-02 | 0 |
| Sialylation | 3 | 1343885 | 1737911 | 1089655 | 1.66 | 9.7E-02 | 0 |
| Total cholesterol in medium HDL | 5 | 1249520 | 1391218 | 889053 | 1.66 | 9.7E-02 | 0 |

| | | | | | | | |
|---|---|---|---|---|---|---|---|
| Red blood cell count (three-way meta) | 132 | 812140 | 1141739 | 281164 | 1.67 | 9.5E-02 | 0 |
| Parkinson disease of sibling pairs (tier 1) | 17 | 1047335 | 1358525 | 557648 | 1.67 | 9.5E-02 | 0 |
| Thyroid preparations | 99 | 839208 | 1173291 | 436561 | 1.68 | 9.3E-02 | 0 |
| Sum basophil neutrophil count (two-way meta) | 71 | 865400 | 1231061 | 101524 | 1.68 | 9.2E-02 | 0 |
| Lumbar Spine BMD (males) | 3 | 1356720 | 1525356 | 936653 | 1.69 | 9.1E-02 | 0 |
| Past tobacco smoking | 65 | 876990 | 1208848 | 87778 | 1.69 | 9.1E-02 | 0 |
| Time spend outdoors in summer | 47 | 914723 | 1164664 | 103434 | 1.69 | 9.1E-02 | 0 |
| Peptide::gamma-glutamyl::gamma-glutamylvaline | 1 | 1565210 | 1565210 | 1565210 | 1.69 | 9.0E-02 | 0 |
| Impedance measures - Arm predicted mass (right) | 526 | 745993 | 1162627 | 108436 | 1.69 | 9.0E-02 | 0 |
| Osteoarthritis (self reported) | 1 | 1573255 | 1573255 | 1573255 | 1.70 | 8.8E-02 | 0 |
| Fraction of accelerations | 2 | 1315089 | 1408074 | 1222103 | 1.71 | 8.8E-02 | 0 |
| Sum neutrophil eosinophil count (two-way meta) | 77 | 865400 | 1216848 | 169561 | 1.71 | 8.7E-02 | 0 |
| Calcium channel blockers | 82 | 858356 | 1221251 | 317506 | 1.72 | 8.6E-02 | 0 |
| Treatment/medication code: paracetamol | 4 | 1243300 | 1426584 | 1081704 | 1.72 | 8.6E-02 | 0 |
| Late pubertal growth | 1 | 1588765 | 1588765 | 1588765 | 1.72 | 8.6E-02 | 0 |
| CD4:%Treg(39+73-) | 1 | 1592028 | 1592028 | 1592028 | 1.72 | 8.6E-02 | 0 |
| Well-being spectrum | 129 | 820555 | 1240205 | 126968 | 1.72 | 8.6E-02 | 0 |
| Serum Urate | 23 | 1015240 | 1329810 | 453078 | 1.72 | 8.5E-02 | 0 |
| Superior fronto-occipital fasciculus radial diusivities | 2 | 1332534 | 1489222 | 1175846 | 1.73 | 8.4E-02 | 0 |
| CD4:%Treg(39+) | 1 | 1592028 | 1592028 | 1592028 | 1.73 | 8.4E-02 | 0 |
| Light smokers, at least 100 smokes in lifetime | 2 | 1328129 | 1415968 | 1240289 | 1.73 | 8.3E-02 | 0 |
| Diastolic Blood Pressure (automated reading) | 194 | 796325 | 1152338 | 139206 | 1.73 | 8.3E-02 | 0 |
| Waist-hip ratio (female > 50 yrs, adjusted for BMI) | 13 | 1109360 | 1305485 | 743133 | 1.73 | 8.3E-02 | 0 |
| CD39 on CD4 T | 1 | 1611068 | 1611068 | 1611068 | 1.74 | 8.2E-02 | 0 |
| Drugs used in diabetes | 45 | 922743 | 1007783 | 353550 | 1.74 | 8.1E-02 | 0 |
| Tanner scale | 1 | 1613123 | 1613123 | 1613123 | 1.74 | 8.1E-02 | 0 |
| Right pallidum | 14 | 1080873 | 1448788 | 982972 | 1.75 | 8.0E-02 | 0 |
| Fasting insulin main effect | 3 | 1382225 | 1433306 | 1214708 | 1.75 | 7.9E-02 | 0 |
| Major depressive disorder (ICD-coded) | 1 | 1604963 | 1604963 | 1604963 | 1.76 | 7.9E-02 | 0 |
| Impedance measures - Leg fat percentage (right) | 378 | 761860 | 1166589 | 211315 | 1.76 | 7.8E-02 | 0 |
| Triglycerides cholesterol | 61 | 895060 | 1138703 | 417360 | 1.76 | 7.8E-02 | 0 |
| Monogalactosylation | 2 | 1335314 | 1733626 | 937002 | 1.77 | 7.8E-02 | 0 |
| FVC | 398 | 761659 | 1180471 | 212957 | 1.77 | 7.7E-02 | 0 |
| Number of cigarettes previously smoked daily | 2 | 1347998 | 1502146 | 1193849 | 1.77 | 7.6E-02 | 0 |
| Triglycerides - Birth cohorts | 1 | 1633368 | 1633368 | 1633368 | 1.77 | 7.6E-02 | 0 |
| Fucosylation | 2 | 1335314 | 1733626 | 937002 | 1.78 | 7.6E-02 | 0 |
| Impedance measures - Leg fat-free mass (right) | 517 | 750835 | 1173203 | 114014 | 1.78 | 7.6E-02 | 0 |
| Rheumatoid Arthritis | 66 | 886414 | 1181998 | 439329 | 1.78 | 7.6E-02 | 0 |
| Sensitivity / hurt feelings | 30 | 984788 | 1155241 | 102087 | 1.78 | 7.5E-02 | 0 |
| White blood cell count (three-way meta) | 121 | 832258 | 1216848 | 241180 | 1.78 | 7.4E-02 | 0 |
| ALPL - Alkaline phosphatase, tissue-nonspecific isozyme | 1 | 1633185 | 1633185 | 1633185 | 1.79 | 7.4E-02 | 0 |
| Diagnoses - secondary ICD10: K44 Diaphragmatic hernia | 3 | 1398910 | 1434606 | 1106865 | 1.79 | 7.4E-02 | 0 |
| Amino acid::Tryptophan metabolism::kynurenine | 2 | 1332175 | 1733535 | 930815 | 1.79 | 7.4E-02 | 0 |

| | | | | | | | |
|---|---|---|---|---|---|---|---|
| Schizophrenia vs Bipolar disorder | 75 | 873278 | 1273734 | 166938 | 1.79 | 7.3E-02 | 0 |
| Tense / 'highly strung' | 21 | 1039700 | 1240205 | 773518 | 1.79 | 7.3E-02 | 0 |
| Impedance measures - Trunk fat-free mass | 640 | 743453 | 1189853 | 138389 | 1.79 | 7.3E-02 | 0 |
| Types of physical activity in last 4 weeks: Walking for pleasure (not as a means of transport) | 3 | 1400850 | 1478315 | 752347 | 1.79 | 7.3E-02 | 0 |
| Impedance measures - Trunk predicted mass | 640 | 743453 | 1192461 | 145750 | 1.80 | 7.3E-02 | 0 |
| LY9 - T-lymphocyte surface antigen Ly-9 | 1 | 1614420 | 1614420 | 1614420 | 1.80 | 7.2E-02 | 0 |
| Impedance measures - Body Mass Index (BMI) | 401 | 763488 | 1183325 | 244295 | 1.80 | 7.2E-02 | 0 |
| Pyruvate | 1 | 1633368 | 1633368 | 1633368 | 1.80 | 7.1E-02 | 0 |
| Low-density lipoprotein particle size by NMR (at exam 4, GEE, adjusted for multivariable) | 2 | 1353031 | 1661883 | 1044179 | 1.81 | 7.0E-02 | 0 |
| Medication for pain relief, constipation, heartburn: Paracetamol | 9 | 1193768 | 1440695 | 1061773 | 1.81 | 7.0E-02 | 0 |
| Carbohydrate::Fructose, mannose, galactose, starch, and sucrose metabolism::mannose | 1 | 1633368 | 1633368 | 1633368 | 1.81 | 7.0E-02 | 0 |
| Depressive symptoms | 11 | 1162908 | 1220804 | 457311 | 1.81 | 7.0E-02 | 0 |
| Hematocrit (two-way meta) | 91 | 859663 | 1147904 | 245206 | 1.82 | 7.0E-02 | 0 |
| Impedance measures - Arm fat-free mass (right) | 527 | 751563 | 1188536 | 110709 | 1.84 | 6.6E-02 | 0 |
| Mean corpuscular hemoglobin concentration (two-way meta) | 43 | 949068 | 1230773 | 543046 | 1.84 | 6.6E-02 | 0 |
| Pubertal growth | 1 | 1642628 | 1642628 | 1642628 | 1.85 | 6.5E-02 | 0 |
| PDE5A - cGMP-specific 3',5'-cyclic phosphodiesterase | 1 | 1653040 | 1653040 | 1653040 | 1.85 | 6.5E-02 | 0 |
| Mother's age | 2 | 1362870 | 1561574 | 1164166 | 1.86 | 6.3E-02 | 0 |
| Total cholesterol / High-density lipoprotein cholesterol ratio (at exam 1, GEE, adjusted for multiva | 1 | 1655515 | 1655515 | 1655515 | 1.86 | 6.3E-02 | 0 |
| Amino acid::Valine, leucine and isoleucine metabolism::4-methyl-2-oxopentanoate | 2 | 1364501 | 1433571 | 1295432 | 1.86 | 6.3E-02 | 0 |
| IL1RL2 - Interleukin-1 receptor-like 2 | 1 | 1661675 | 1661675 | 1661675 | 1.86 | 6.2E-02 | 0 |
| Type 2 Diabetes | 417 | 763233 | 1115525 | 357085 | 1.87 | 6.1E-02 | 0 |
| Amino acid::Amino fatty acid::X-12510--2-aminooctanoic acid | 2 | 1378240 | 1449574 | 1306906 | 1.88 | 6.1E-02 | 0 |
| FEV1/FVC ratio | 478 | 757640 | 1155103 | 162547 | 1.88 | 6.0E-02 | 0 |
| CD4:%Treg(73+) | 1 | 1680665 | 1680665 | 1680665 | 1.88 | 6.0E-02 | 0 |
| CD4:%Treg(39-73+) | 1 | 1680665 | 1680665 | 1680665 | 1.88 | 6.0E-02 | 0 |
| Treatment/medication code: atorvastatin | 16 | 1097148 | 1162892 | 741303 | 1.88 | 6.0E-02 | 0 |
| Time spent outdoors in winter | 11 | 1181195 | 1235116 | 555412 | 1.88 | 5.9E-02 | 0 |
| Age at first birth | 8 | 1209933 | 1261298 | 869150 | 1.90 | 5.7E-02 | 0 |
| Granulocyte count (two-way meta) | 76 | 889219 | 1232085 | 248232 | 1.91 | 5.7E-02 | 0 |
| Overall activity | 2 | 1395661 | 1403164 | 1388158 | 1.92 | 5.5E-02 | 0 |
| Left putamen | 9 | 1231710 | 1265200 | 1042845 | 1.92 | 5.5E-02 | 0 |
| Platelet count (two-way meta) | 141 | 832258 | 1144778 | 151916 | 1.92 | 5.5E-02 | 0 |
| ::::X-12063 | 4 | 1313896 | 1412121 | 926203 | 1.93 | 5.4E-02 | 0 |
| Proxy and clinically diagnosed Alzheimer's disease | 33 | 998360 | 1277675 | 127536 | 1.93 | 5.4E-02 | 0 |
| Hair colour (natural, before greying): Blonde | 124 | 844196 | 1116346 | 253876 | 1.94 | 5.2E-02 | 0 |
| Sensitivity / hurt feelings (HURT) | 26 | 1034370 | 1155241 | 311621 | 1.95 | 5.2E-02 | 0 |
| Monocyte chemoattractant protein 1 (at exam 7, GEE, adjusted for multivariable) | 2 | 1401405 | 1480990 | 1321820 | 1.95 | 5.1E-02 | 0 |
| Impedance measures - Arm predicted mass (left) | 548 | 754924 | 1172258 | 129881 | 1.95 | 5.1E-02 | 0 |
| EPHA1 - Ephrin type-A receptor 1 | 1 | 1697610 | 1697610 | 1697610 | 1.95 | 5.1E-02 | 0 |
| Waist-hip ratio (adjusted for BMI) | 60 | 916700 | 1301019 | 420873 | 1.95 | 5.1E-02 | 0 |
| Platelet count (three-way meta) | 171 | 821920 | 1174625 | 212250 | 1.95 | 5.1E-02 | 0 |
| Frequent insomnia symptoms | 40 | 970969 | 1204728 | 282681 | 1.96 | 4.9E-02 | 0 |

| | | | | | | | |
|---|---|---|---|---|---|---|---|
| Height (male) | 38 | 981094 | 1174986 | 303788 | 1.97 | 4.9E-02 | 0 |
| Impedance measures - Whole body fat-free mass | 631 | 751563 | 1188323 | 123781 | 1.97 | 4.9E-02 | 0 |
| Sleeplessness / insomnia | 28 | 1022165 | 1090000 | 490857 | 1.97 | 4.9E-02 | 0 |
| Juvenile Myoclonic Epilepsy (JME) | 1 | 1725310 | 1725310 | 1725310 | 1.98 | 4.8E-02 | 0 |
| Amino acid::Valine, leucine and isoleucine metabolism::3-methyl-2-oxovalerate | 2 | 1429865 | 1531616 | 1328114 | 1.99 | 4.7E-02 | 0 |
| Able to confide | 8 | 1230328 | 1433348 | 842774 | 1.99 | 4.7E-02 | 0 |
| Fornix (cres) / Stria terminalis radial diusivities | 2 | 1420088 | 1560780 | 1279395 | 2.00 | 4.6E-02 | 0 |
| Amyotrophic lateral sclerosis (LMM) | 4 | 1334358 | 1863639 | 969464 | 2.00 | 4.6E-02 | 0 |
| Coronary artery disease and triglyceride (bivariate) | 125 | 850163 | 1055668 | 388948 | 2.00 | 4.6E-02 | 0 |
| Genu of corpus callosum fractional anisotropy | 5 | 1377788 | 1381305 | 577610 | 2.00 | 4.5E-02 | 0 |
| Neuroticism (MA GWAMA) | 138 | 840680 | 1297067 | 433145 | 2.00 | 4.5E-02 | 0 |
| Left pallidum | 12 | 1174571 | 1424523 | 1031357 | 2.00 | 4.5E-02 | 0 |
| Impedance measures - Impedance of whole body | 495 | 762915 | 1174960 | 133896 | 2.01 | 4.5E-02 | 0 |
| Mean platelet volume (three-way meta) | 156 | 829454 | 1151882 | 276298 | 2.01 | 4.4E-02 | 0 |
| Cerebellar vermal lobules VIII X | 25 | 1063038 | 1375498 | 596680 | 2.02 | 4.4E-02 | 0 |
| Platelet count | 75 | 897073 | 1147104 | 415343 | 2.02 | 4.3E-02 | 0 |
| Extreme Height | 49 | 955428 | 1164553 | 345608 | 2.02 | 4.3E-02 | 0 |
| Left inferior temporal | 2 | 1438299 | 1493834 | 1382763 | 2.03 | 4.2E-02 | 0 |
| Age at menarche | 254 | 798183 | 1153408 | 326588 | 2.04 | 4.1E-02 | 0 |
| Superior fronto-occipital fasciculus mean diusivities | 3 | 1499188 | 1602503 | 1468346 | 2.05 | 4.1E-02 | 0 |
| Age first had sexual intercourse | 90 | 884554 | 1258953 | 444909 | 2.05 | 4.0E-02 | 0 |
| Impedance measures - Whole body fat mass | 416 | 772084 | 1141230 | 141360 | 2.05 | 4.0E-02 | 0 |
| Epilepsy | 4 | 1357533 | 1549868 | 1062998 | 2.06 | 3.9E-02 | 0 |
| FGF2 - Fibroblast growth factor 2 | 1 | 1783258 | 1783258 | 1783258 | 2.06 | 3.9E-02 | 0 |
| Sum eosinophil basophil count (two-way meta) | 104 | 874868 | 1138026 | 238267 | 2.07 | 3.9E-02 | 0 |
| Large low-density lipoprotein by NMR (at exam 4, GEE, adjusted for age and sex) | 1 | 1766975 | 1766975 | 1766975 | 2.07 | 3.8E-02 | 0 |
| Right middle temporal | 1 | 1772910 | 1772910 | 1772910 | 2.08 | 3.8E-02 | 0 |
| Impedance measures - Weight | 534 | 762673 | 1167138 | 139750 | 2.08 | 3.8E-02 | 0 |
| Total body BMD (45-60 years old) | 16 | 1143158 | 1458020 | 525667 | 2.09 | 3.7E-02 | 0 |
| Carbohydrate::Aminosugars metabolism::erythronate* | 3 | 1516068 | 1629749 | 1335585 | 2.09 | 3.6E-02 | 0 |
| Phospholipids in medium HDL | 4 | 1363576 | 1384329 | 1155333 | 2.11 | 3.5E-02 | 0 |
| Medication for cholesterol, blood pressure, diabetes, or take exogenous hormones: Blood pressu | 76 | 912393 | 1185896 | 201241 | 2.12 | 3.4E-02 | 0 |
| Hemoglobin concentration (two-way meta) | 85 | 896078 | 1134575 | 465000 | 2.12 | 3.4E-02 | 0 |
| Weight | 549 | 763615 | 1183063 | 164584 | 2.14 | 3.2E-02 | 0 |
| Free cholesterol in medium HDL | 4 | 1386433 | 1415448 | 1183306 | 2.16 | 3.1E-02 | 0 |
| Red cell distribution width (three-way meta) | 119 | 870308 | 1173549 | 307930 | 2.17 | 3.0E-02 | 0 |
| Job involves mainly walking or standing | 10 | 1245224 | 1470354 | 973370 | 2.18 | 2.9E-02 | 0 |
| Amino acid::Histidine metabolism::histidine | 1 | 1843255 | 1843255 | 1843255 | 2.19 | 2.8E-02 | 0 |
| Mean corpuscular hemoglobin concentration (three-way meta) | 61 | 949068 | 1246753 | 557063 | 2.19 | 2.8E-02 | 0 |
| Fed-up feelings (FED_UP) | 28 | 1072941 | 1317129 | 510283 | 2.21 | 2.7E-02 | 0 |
| Type 1 Diabetes | 78 | 916961 | 1193800 | 379350 | 2.21 | 2.7E-02 | 0 |
| Impedance measures - Arm fat percentage (right) | 357 | 788650 | 1198243 | 156515 | 2.21 | 2.7E-02 | 0 |
| Impedance measures - Arm fat mass (right) | 317 | 795785 | 1190990 | 319590 | 2.22 | 2.6E-02 | 0 |

| | | | | | | | |
|---|---|---|---|---|---|---|---|
| Poultry intake | 12 | 1226236 | 1249541 | 393457 | 2.23 | 2.6E-02 | 0 |
| Pain type(s) experienced in last month: Headache | 31 | 1061773 | 1216263 | 200157 | 2.25 | 2.4E-02 | 0 |
| Large low-density lipoprotein by NMR (at exam 4, GEE, adjusted for multivariable) | 2 | 1511128 | 1639051 | 1383204 | 2.26 | 2.4E-02 | 0 |
| White blood cell count (two-way meta) | 104 | 889219 | 1272826 | 322686 | 2.26 | 2.4E-02 | 0 |
| Impedance measures - Impedance of arm (left) | 426 | 783736 | 1189558 | 155748 | 2.28 | 2.3E-02 | 0 |
| Family history of Alzheimer's disease | 6 | 1365889 | 1602587 | 1348568 | 2.28 | 2.3E-02 | 0 |
| Diagnoses - main ICD10: M23 Internal derangement of knee | 1 | 1884003 | 1884003 | 1884003 | 2.29 | 2.2E-02 | 0 |
| Impedance measures - Impedance of arm (right) | 420 | 784519 | 1167779 | 151590 | 2.30 | 2.1E-02 | 0 |
| Waist-hip ratio (female) | 293 | 806353 | 1157218 | 379398 | 2.31 | 2.1E-02 | 0 |
| Impedance measures - Arm fat percentage (left) | 364 | 793465 | 1177763 | 240071 | 2.32 | 2.1E-02 | 0 |
| Impedance measures - Arm fat mass (left) | 300 | 803375 | 1180508 | 374714 | 2.32 | 2.1E-02 | 0 |
| Estimated glomerular filtration rate | 459 | 779760 | 1148385 | 262656 | 2.32 | 2.1E-02 | 0 |
| Birth weight of first child (female) | 40 | 1026731 | 1303378 | 134069 | 2.34 | 1.9E-02 | 0 |
| Eczema | 10 | 1296890 | 1401259 | 492823 | 2.35 | 1.9E-02 | 0 |
| Waist-hip ratio (male) | 133 | 873133 | 1210258 | 318705 | 2.35 | 1.9E-02 | 0 |
| IL16 - Interleukin-16 | 1 | 1904160 | 1904160 | 1904160 | 2.35 | 1.9E-02 | 0 |
| Smoking status: Never | 65 | 963830 | 1307260 | 123532 | 2.37 | 1.8E-02 | 0 |
| Lactate | 2 | 1566465 | 1599916 | 1533014 | 2.38 | 1.7E-02 | 0 |
| Waist-hip ratio (adjusted for BMI, female) | 317 | 806353 | 1140565 | 265245 | 2.42 | 1.5E-02 | 0 |
| Red cell distribution width (two-way meta) | 105 | 906293 | 1212675 | 276643 | 2.43 | 1.5E-02 | 0 |
| IL1B - Interleukin-1 beta | 1 | 1969643 | 1969643 | 1969643 | 2.43 | 1.5E-02 | 0 |
| BMP10 - Bone morphogenetic protein 10 | 1 | 1969643 | 1969643 | 1969643 | 2.44 | 1.5E-02 | 0 |
| Waist circumference (female, adjusted for BMI) | 36 | 1069156 | 1223924 | 760473 | 2.44 | 1.5E-02 | 0 |
| Alanine | 4 | 1476583 | 1625281 | 1015272 | 2.45 | 1.4E-02 | 0 |
| COLEC11 - Collectin-11 | 1 | 1969643 | 1969643 | 1969643 | 2.46 | 1.4E-02 | 0 |
| Impedance measures - Leg fat mass (left) | 343 | 805280 | 1170864 | 156766 | 2.46 | 1.4E-02 | 0 |
| Fasting insulin main effect (adjusted for BMI) | 9 | 1391218 | 1518550 | 936215 | 2.47 | 1.4E-02 | 0 |
| Depressive symptoms (MA GWAMA) | 118 | 894703 | 1299175 | 446739 | 2.47 | 1.3E-02 | 0 |
| Hair (Straight/curly) | 1 | 1981195 | 1981195 | 1981195 | 2.48 | 1.3E-02 | 0 |
| Birth weight | 162 | 868386 | 1151339 | 307451 | 2.48 | 1.3E-02 | 0 |
| Waist circumference | 372 | 801440 | 1191453 | 297668 | 2.48 | 1.3E-02 | 0 |
| NACA - Nascent polypeptide-associated complex subunit alpha | 1 | 1969643 | 1969643 | 1969643 | 2.48 | 1.3E-02 | 0 |
| Low-density lipoprotein particle size by NMR (at exam 4, GEE, adjusted for age and sex) | 1 | 1970735 | 1970735 | 1970735 | 2.49 | 1.3E-02 | 0 |
| Femoral Neck BMD (females) | 10 | 1349605 | 1404806 | 992534 | 2.53 | 1.1E-02 | 0 |
| Intelligence | 197 | 852238 | 1183653 | 97955 | 2.56 | 1.0E-02 | 0 |
| Why stopped smoking: Health precaution | 2 | 1648120 | 2260548 | 1035693 | 2.60 | 9.3E-03 | 0 |
| Ever smoker | 161 | 873400 | 1234615 | 159386 | 2.60 | 9.2E-03 | 0 |
| PRSS2 - Trypsin-2 | 1 | 2039865 | 2039865 | 2039865 | 2.62 | 8.7E-03 | 0 |
| Impedance measures - Whole body water mass | 635 | 778098 | 1193205 | 134399 | 2.66 | 7.8E-03 | 0 |
| Hemoglobin concentration (three-way meta) | 95 | 941758 | 1314673 | 365079 | 2.66 | 7.8E-03 | 0 |
| Impedance measures - Leg fat percentage (left) | 389 | 805280 | 1193700 | 183401 | 2.66 | 7.8E-03 | 0 |
| Hematocrit (three-way meta) | 83 | 963458 | 1265828 | 339836 | 2.67 | 7.5E-03 | 0 |
| Excessive sweating | 2 | 1686329 | 1737813 | 1634844 | 2.71 | 6.8E-03 | 0 |

| | | | | | | | |
|---|---|---|---|---|---|---|---|
| Monosialylation | 1 | 2131938 | 2131938 | 2131938 | 2.76 | 5.8E-03 | 0 |
| Getting up in morning | 52 | 1038246 | 1241453 | 185757 | 2.77 | 5.6E-03 | 0 |
| Impedance measures - Leg fat mass (right) | 341 | 823015 | 1190990 | 232272 | 2.81 | 4.9E-03 | 0 |
| Sleep duration | 104 | 946296 | 1238599 | 343410 | 2.86 | 4.3E-03 | 0 |
| Red blood cell count (two-way meta) | 121 | 925328 | 1219650 | 489015 | 2.86 | 4.3E-03 | 0 |
| Hip circumference | 436 | 810475 | 1208070 | 195594 | 2.88 | 4.0E-03 | 0 |
| High blood pressure | 235 | 858543 | 1126979 | 237043 | 2.89 | 3.8E-03 | 0 |
| Salt added to food | 72 | 1011131 | 1252551 | 476143 | 2.91 | 3.7E-03 | 0 |
| Body Mass Index (female) | 453 | 813918 | 1172715 | 364585 | 3.04 | 2.4E-03 | 0 |
| FEV1 | 406 | 826861 | 1198660 | 317436 | 3.15 | 1.6E-03 | 0 |
| Waist-hip ratio (adjusted for BMI, male) | 128 | 954580 | 1197550 | 273060 | 3.24 | 1.2E-03 | 0 |
| Waist-hip ratio | 465 | 825688 | 1157315 | 344535 | 3.31 | 9.2E-04 | 0 |
| Waist circumference (adjusted for BMI) | 89 | 1036333 | 1250248 | 550950 | 3.49 | 4.9E-04 | 0 |
| Age at last live birth (female) | 1 | 2660000 | 2660000 | 2660000 | 3.74 | 1.8E-04 | 0 |
| Body Mass Index | 1671 | 794265 | 1180003 | 210393 | 4.96 | 6.9E-07 | 1 |
| Height | 1693 | 811305 | 1161105 | 308065 | 5.75 | 9.1E-09 | 1 |
