## Supplementary Table 2 for "Mapping the genetic evolutionary timeline of human neural and cognitive traits"

| Domain | N SNPs | Median age | Age 75% upper bound | Age 25% lower bound | Median MAF | Z-score | P value | Bonferroni |
|---|---|---|---|---|---|---|---|---|
| Psychiatric | 3577 | 475833 | 1078170 | 48958 | 0.242 | -11.84 | 2.4E-32 | 1 |
| Activities | 1435 | 644450 | 1130073 | 81190 | 0.295 | -4.57 | 5.0E-06 | 1 |
| Environment | 862 | 626965 | 1123312 | 79835 | 0.282 | -3.90 | 9.7E-05 | 1 |
| Social Interactions | 160 | 474527 | 1046098 | 55032 | 0.254 | -2.88 | 4.0E-03 | 0 |
| Reproduction | 1064 | 611368 | 1094320 | 63490 | 0.260 | -2.44 | 1.5E-02 | 0 |
| Ophthalmological | 382 | 355860 | 945678 | 35806 | 0.097 | -2.24 | 2.5E-02 | 0 |
| Cognitive | 530 | 682674 | 1102371 | 90468 | 0.308 | -1.87 | 6.2E-02 | 0 |
| Ear, Nose, Throat | 52 | 642079 | 1078489 | 96134 | 0.343 | -1.49 | 1.4E-01 | 0 |
| Body Structures | 90 | 699615 | 1100887 | 166689 | 0.319 | -0.88 | 3.8E-01 | 0 |
| Respiratory | 1746 | 761659 | 1155551 | 207961 | 0.300 | -0.81 | 4.2E-01 | 0 |
| Endocrine | 858 | 759645 | 1091699 | 368652 | 0.291 | -0.10 | 9.2E-01 | 0 |
| Connective Tissue | 94 | 816138 | 1178558 | 318661 | 0.318 | 0.09 | 9.3E-01 | 0 |
| Hematological | 57 | 595340 | 1052398 | 214394 | 0.189 | 0.13 | 9.0E-01 | 0 |
| Skeletal | 4386 | 764959 | 1143706 | 212789 | 0.285 | 0.19 | 8.5E-01 | 0 |
| Infection | 6 | 752978 | 1142349 | 462069 | 0.211 | 0.36 | 7.2E-01 | 0 |
| Cardiovascular | 1992 | 759193 | 1137015 | 229188 | 0.283 | 0.39 | 7.0E-01 | 0 |
| Environmental | 599 | 803743 | 1150583 | 266960 | 0.298 | 0.80 | 4.2E-01 | 0 |
| Muscular | 3 | 782185 | 1153615 | 691033 | 0.019 | 0.85 | 4.0E-01 | 0 |
| Nutritional | 623 | 771528 | 1174960 | 85350 | 0.289 | 0.98 | 3.3E-01 | 0 |
| Neurological | 815 | 800850 | 1174015 | 173721 | 0.304 | 1.23 | 2.2E-01 | 0 |
| Gastrointestinal | 598 | 667234 | 1072639 | 202736 | 0.216 | 1.29 | 2.0E-01 | 0 |
| Dermatological | 1594 | 703298 | 1103587 | 157359 | 0.242 | 1.48 | 1.4E-01 | 0 |
| Mortality | 139 | 652970 | 1049563 | 131524 | 0.143 | 1.55 | 1.2E-01 | 0 |
| Immunological | 4076 | 752814 | 1135338 | 220139 | 0.256 | 2.91 | 3.6E-03 | 0 |

| | | | | | | | | |
|---|---|---|---|---|---|---|---|---|
| Metabolic | 10131 | 785573 | 1171886 | 213102 | 0.290 | 4.01 | 6.0E-05 | 1 |
| Neoplasms | 373 | 591690 | 1000520 | 125718 | 0.044 | 4.46 | 8.1E-06 | 1 |

| Chapter | N SNPs | Median age | Age 75% upper bound | Age 25% lower bound | Median MAF | Z-score | P value | Bonferroni |
|---|---|---|---|---|---|---|---|---|
| Mental and Behavioural Disorders | 1893 | 83250 | 869535 | 21940 | 0.047 | -17.55 | 5.7E-69 | 1 |
| Major Life Areas | 817 | 617780 | 1117690 | 77641 | 0.281 | -3.97 | 7.3E-05 | 1 |
| Self-Care | 1965 | 685023 | 1145540 | 82951 | 0.291 | -3.01 | 2.6E-03 | 0 |
| Domestic Life | 6 | 94093 | 928996 | 64279 | 0.375 | -2.93 | 3.4E-03 | 0 |
| Interpersonal Interactions and Relationships | 160 | 474527 | 1046098 | 55032 | 0.254 | -2.85 | 4.3E-03 | 0 |
| Genitourinary and Reproductive Functions | 953 | 601310 | 1094123 | 61485 | 0.260 | -2.54 | 1.1E-02 | 0 |
| The Eye, Ear and Related Structures | 315 | 254530 | 845168 | 29329 | 0.030 | -2.46 | 1.4E-02 | 0 |
| Injuries to the Wrist or Hand | 12 | 89668 | 637349 | 43850 | 0.224 | -2.41 | 1.6E-02 | 0 |
| Voice and Speech Functions | 1 | 69269 | 69269 | 69269 | 0.290 | -1.34 | 1.8E-01 | 0 |
| Mental Functions | 2238 | 782174 | 1181555 | 122777 | 0.313 | -1.22 | 2.2E-01 | 0 |
| Certain Conditions Originating in the Perinatal Period | 1 | 40478 | 40478 | 40478 | 0.111 | -1.15 | 2.5E-01 | 0 |
| Communication | 59 | 698518 | 1110025 | 114528 | 0.317 | -0.98 | 3.3E-01 | 0 |
| Diseases of the Eye and Adnexa | 72 | 680315 | 1062517 | 140268 | 0.327 | -0.89 | 3.7E-01 | 0 |
| Structure Involved in Voice and Speech | 60 | 697368 | 1088796 | 149532 | 0.325 | -0.84 | 4.0E-01 | 0 |
| Symptoms, Signs and Abnormal Clinical and Laboratory Findings, Not Elsewhere Classified | 28 | 651598 | 1192498 | 293286 | 0.249 | -0.76 | 4.5E-01 | 0 |
| Persons Encountering Health Services in Circumstances related to Reproduction | 1 | 33618 | 33618 | 33618 | 0.018 | -0.74 | 4.6E-01 | 0 |
| Disease of the Respiratory System | 188 | 700429 | 1144659 | 141717 | 0.267 | -0.73 | 4.7E-01 | 0 |
| Structures of the Nervous System | 554 | 728523 | 1136119 | 163595 | 0.299 | -0.70 | 4.9E-01 | 0 |
| Diseases of the Respiratory System | 228 | 764024 | 1113379 | 232326 | 0.305 | -0.63 | 5.3E-01 | 0 |
| Diseases of the Musculoskeletal System and Connective Tissue | 231 | 746580 | 1173024 | 309821 | 0.304 | -0.63 | 5.3E-01 | 0 |
| Mobility | 15 | 572478 | 791724 | 344053 | 0.238 | -0.56 | 5.7E-01 | 0 |
| Endocrine, Nutritional and Metabolic Diseases | 898 | 755668 | 1110452 | 316624 | 0.294 | -0.55 | 5.8E-01 | 0 |
| Factors Influencing Health Status and Contact with Health Services | 12 | 656330 | 1023916 | 166671 | 0.218 | -0.50 | 6.2E-01 | 0 |
| Benign Neoplasms | 11 | 622368 | 978608 | 94680 | 0.310 | -0.41 | 6.8E-01 | 0 |
| Diseases of the Skin and Subcutaneous Tissue | 59 | 750523 | 1221463 | 144052 | 0.316 | -0.29 | 7.7E-01 | 0 |
| Diseases of the Circulatory System | 829 | 755865 | 1131780 | 301905 | 0.281 | -0.08 | 9.3E-01 | 0 |
| Structures Related to Genitourinary and Reproductive Functions | 2 | 721569 | 763804 | 679333 | 0.243 | -0.03 | 9.8E-01 | 0 |
| Injury, Poisoning and Certain Other Consequences of External Causes | 32 | 804538 | 1073765 | 208062 | 0.316 | 0.15 | 8.8E-01 | 0 |
| Functions of the Skin and Related Structures | 717 | 775515 | 1109543 | 300378 | 0.292 | 0.41 | 6.8E-01 | 0 |
| Certain conditions originating in the perinatal period | 7 | 924228 | 1220385 | 660623 | 0.291 | 0.43 | 6.7E-01 | 0 |
| Pregnancy, Childbirth and the Puerperium | 6 | 325911 | 879928 | 30598 | 0.001 | 0.44 | 6.6E-01 | 0 |
| Structures Related to Movement | 4240 | 768539 | 1143659 | 210562 | 0.285 | 0.47 | 6.4E-01 | 0 |
| Sensory Functions and Pain | 116 | 848390 | 1136323 | 168780 | 0.317 | 0.77 | 4.4E-01 | 0 |
| Products and Technology | 632 | 803074 | 1163988 | 245322 | 0.294 | 0.85 | 3.9E-01 | 0 |
| Neuromusculoskeletal and Movement-Related Functions | 4 | 691033 | 967900 | 542651 | 0.015 | 0.97 | 3.3E-01 | 0 |
| Persons with Potential Health Hazards Related to Family and Personal History and Certain Conditions Influencing Health Status | 18 | 990826 | 1104152 | 626772 | 0.236 | 1.33 | 1.8E-01 | 0 |
| Diseases of the Digestive System | 597 | 669370 | 1074650 | 206778 | 0.218 | 1.36 | 1.7E-01 | 0 |
| Mortality | 121 | 616138 | 1020063 | 105694 | 0.083 | 1.39 | 1.6E-01 | 0 |
| Persons Encountering Health Status and Contact with Health Services | 1 | 1529998 | 1529998 | 1529998 | 0.222 | 1.58 | 1.1E-01 | 0 |
| Skin and Related Structures | 830 | 634305 | 1090996 | 97258 | 0.184 | 1.78 | 7.4E-02 | 0 |
| Diseases of the Genitourinary System | 89 | 661478 | 1090430 | 207140 | 0.046 | 1.99 | 4.6E-02 | 0 |
| Diseases of the Nervous System | 206 | 933313 | 1244547 | 286080 | 0.307 | 2.58 | 1.0E-02 | 0 |
| Functions of the Cardiovascular, Haematological, Immunological and Respiratory Systems | 6700 | 758878 | 1142685 | 218702 | 0.268 | 2.69 | 7.2E-03 | 0 |
| Functions of the Digestive, Metabolic and Endocrine Systems | 10147 | 785400 | 1170475 | 217856 | 0.290 | 3.78 | 1.6E-04 | 1 |
| Malignant Neoplasms | 363 | 590658 | 995774 | 132831 | 0.042 | 4.51 | 6.4E-06 | 1 |

| Subchapter | N SNPs | Median age | Age 75% upper bound | Age 25% lower bound | Median MAF | Z-score | P value | Bonferroni |
|---|---|---|---|---|---|---|---|---|
| Mental and Behavioural Disorders Due to Use of Alcohol | 481 | 39549 | 611800 | 17611 | 0.010 | -6.90 | 5.2E-12 | 1 |
| Sexual Functions | 349 | 76815 | 853113 | 17446 | 0.016 | -6.86 | 7.1E-12 | 1 |
| Education | 578 | 637646 | 1131909 | 103298 | 0.306 | -4.27 | 2.0E-05 | 1 |
| Looking After One's Health | 1359 | 646078 | 1131673 | 80516 | 0.294 | -4.25 | 2.2E-05 | 1 |
| Depressive Episode | 658 | 24471 | 78286 | 14889 | 0.004 | -3.76 | 1.7E-04 | 1 |
| Mental Functions, Unspecified | 4 | 48181 | 350559 | 36350 | 0.342 | -2.54 | 1.1E-02 | 0 |
| Family Relationships | 68 | 176333 | 821454 | 25694 | 0.120 | -2.54 | 1.1E-02 | 0 |
| Structure of Eyeball | 315 | 254530 | 845168 | 29329 | 0.030 | -2.49 | 1.3E-02 | 0 |
| Carpal Tunnel Syndrome | 12 | 89668 | 637349 | 43850 | 0.224 | -2.49 | 1.3E-02 | 0 |
| Unspecified | 5 | 123446 | 1197513 | 64740 | 0.349 | -2.28 | 2.3E-02 | 0 |
| Work and Employment | 90 | 22719 | 618689 | 12761 | 0.006 | -2.16 | 3.1E-02 | 0 |
| Higher-Level Cognitive Functions | 434 | 741750 | 1128273 | 96597 | 0.324 | -2.05 | 4.0E-02 | 0 |
| Acquisition of Necessities | 1 | 55775 | 55775 | 55775 | 0.491 | -1.79 | 7.4E-02 | 0 |
| Obesity | 38 | 602585 | 1028234 | 283569 | 0.339 | -1.76 | 7.8E-02 | 0 |
| Problems Related to Upbringing | 14 | 364589 | 708594 | 50867 | 0.227 | -1.71 | 8.7E-02 | 0 |
| Coxarthrosis [Anthrosis of Hip] | 10 | 343796 | 693370 | 106550 | 0.259 | -1.70 | 8.9E-02 | 0 |
| Informal Social Relationships | 87 | 667720 | 1213893 | 117543 | 0.307 | -1.70 | 9.0E-02 | 0 |
| Schizophrenia/Bipolar Affective Disorder | 152 | 699801 | 1168273 | 96495 | 0.324 | -1.61 | 1.1E-01 | 0 |
| Parent-Child Relationships | 6 | 31772 | 84455 | 28499 | 0.117 | -1.57 | 1.2E-01 | 0 |
| Hearing Functions | 52 | 642079 | 1078489 | 96134 | 0.343 | -1.50 | 1.3E-01 | 0 |
| Intestinal Malabsorption | 38 | 533051 | 1057677 | 126359 | 0.223 | -1.48 | 1.4E-01 | 0 |
| Fibrosis and Cirrhosis of Liver | 17 | 471018 | 837185 | 72818 | 0.285 | -1.47 | 1.4E-01 | 0 |
| Acquired Absence of Organs, Not Elsewhere Classified | 6 | 351934 | 726293 | 99518 | 0.207 | -1.42 | 1.6E-01 | 0 |
| Glaucoma | 65 | 633208 | 1048700 | 143303 | 0.332 | -1.40 | 1.6E-01 | 0 |
| Disorders of Gallbladder, Biliary Tract and Pancreas | 1 | 160339 | 160339 | 160339 | 0.321 | -1.38 | 1.7E-01 | 0 |
| Structure of Nose | 1 | 69269 | 69269 | 69269 | 0.290 | -1.34 | 1.8E-01 | 0 |
| Cardiomiopathy | 1 | 76263 | 76263 | 76263 | 0.223 | -1.33 | 1.8E-01 | 0 |
| Conversation and Use of Communication Devices and Techniques | 58 | 675238 | 1114448 | 112036 | 0.320 | -1.23 | 2.2E-01 | 0 |
| Other Arthrosis | 84 | 712899 | 1059573 | 205613 | 0.359 | -1.21 | 2.3E-01 | 0 |
| Digestive System Disorders of Fetus and Newborn | 1 | 40478 | 40478 | 40478 | 0.111 | -1.13 | 2.6E-01 | 0 |
| Gout | 2 | 288515 | 415423 | 161608 | 0.166 | -1.12 | 2.6E-01 | 0 |
| Psychomotor Functions | 39 | 627113 | 991665 | 187568 | 0.307 | -1.07 | 2.9E-01 | 0 |
| Preterm Labour and Delivery | 2 | 21325 | 24569 | 18081 | 0.005 | -1.07 | 2.9E-01 | 0 |
| Other Hypothyroidism | 46 | 657540 | 1097048 | 387348 | 0.323 | -1.06 | 2.9E-01 | 0 |
| Maligrant Neoplasms of Pancreas | 3 | 415423 | 739820 | 277350 | 0.310 | -1.05 | 2.9E-01 | 0 |
| Disorders of Lens | 2 | 277362 | 357695 | 197029 | 0.142 | -0.96 | 3.4E-01 | 0 |
| Memory Functions | 27 | 112065 | 783248 | 26478 | 0.009 | -0.95 | 3.4E-01 | 0 |
| Disorders of Puberty, Not Elsewhere Classified | 95 | 680763 | 1100234 | 148938 | 0.295 | -0.89 | 3.7E-01 | 0 |
| Schizophrenia | 193 | 755908 | 1218233 | 128092 | 0.327 | -0.89 | 3.7E-01 | 0 |
| Abnormalities of Breathing | 24 | 651598 | 1131414 | 293286 | 0.251 | -0.87 | 3.9E-01 | 0 |
| Structure of Mouth | 60 | 697368 | 1088796 | 149532 | 0.325 | -0.82 | 4.1E-01 | 0 |
| Malignant Neoplasms of Skin | 47 | 564945 | 902261 | 132566 | 0.244 | -0.81 | 4.2E-01 | 0 |
| Asthma | 203 | 760445 | 1110224 | 224877 | 0.310 | -0.79 | 4.3E-01 | 0 |
| Pain in Chest | 4 | 460186 | 939378 | 75222 | 0.232 | -0.76 | 4.5E-01 | 0 |
| Dislocation, Sprain and Strain of Joints and Ligaments of Knee | 1 | 26829 | 26829 | 26829 | 0.046 | -0.75 | 4.5E-01 | 0 |
| Outcome of Delivery | 1 | 33618 | 33618 | 33618 | 0.018 | -0.74 | 4.6E-01 | 0 |

| | | | | | | | | |
|---|---|---|---|---|---|---|---|---|
| Injury of Muscle and Tendon at Shoulder and Upper Arm Level | 1 | 34759 | 34759 | 34759 | 0.047 | -0.73 | 4.7E-01 | 0 |
| Menstruation Functions | 542 | 765793 | 1156609 | 248200 | 0.308 | -0.72 | 4.7E-01 | 0 |
| Structure of Brain | 554 | 728523 | 1136119 | 163595 | 0.299 | -0.72 | 4.7E-01 | 0 |
| Bipolar Affective Disorder/Depressive Episode | 1 | 66277 | 66277 | 66277 | 0.065 | -0.66 | 5.1E-01 | 0 |
| Vasomotor and Allergic Rhinitis | 202 | 700429 | 1146708 | 141077 | 0.267 | -0.65 | 5.2E-01 | 0 |
| Maligrant Neoplasms of Colon | 10 | 616168 | 981138 | 374662 | 0.297 | -0.60 | 5.5E-01 | 0 |
| Failure of Genital Response | 1 | 453883 | 453883 | 453883 | 0.246 | -0.57 | 5.7E-01 | 0 |
| Sleep Functions | 698 | 769318 | 1192279 | 125031 | 0.294 | -0.51 | 6.1E-01 | 0 |
| Personal History of Malignant Neoplasm | 1 | 609465 | 609465 | 609465 | 0.392 | -0.51 | 6.1E-01 | 0 |
| Exercise Tolerance Functions | 1350 | 765109 | 1163311 | 206824 | 0.302 | -0.50 | 6.2E-01 | 0 |
| Malignant Neoplasms of Ovary | 2 | 385889 | 404051 | 367727 | 0.120 | -0.49 | 6.3E-01 | 0 |
| Gonarthrosis [Arthrosis of Knee] | 3 | 614298 | 1249150 | 476524 | 0.285 | -0.48 | 6.3E-01 | 0 |
| Walking and Moving | 9 | 552143 | 836723 | 398870 | 0.262 | -0.48 | 6.3E-01 | 0 |
| Diabetes Mellitus | 71 | 698965 | 1085338 | 127300 | 0.280 | -0.48 | 6.3E-01 | 0 |
| Disorders of eyelid, lacrimal system and orbit | 2 | 219257 | 228872 | 209641 | 0.054 | -0.44 | 6.6E-01 | 0 |
| Benign Neoplasm of Colon, Rectum, Anus and Anal Canal | 11 | 622368 | 978608 | 94680 | 0.310 | -0.41 | 6.8E-01 | 0 |
| Chronic Ischaemic Heart Disease | 425 | 740605 | 1120763 | 322275 | 0.276 | -0.39 | 6.9E-01 | 0 |
| Personal History of Certain Other Diseases | 3 | 225979 | 1377815 | 131869 | 0.023 | -0.38 | 7.0E-01 | 0 |
| Vitiligo | 34 | 731529 | 1131053 | 99137 | 0.315 | -0.32 | 7.5E-01 | 0 |
| Other disorders of pigmentation | 5 | 433813 | 626938 | 424865 | 0.143 | -0.29 | 7.7E-01 | 0 |
| Presence of Other Device | 3 | 626783 | 945589 | 324752 | 0.466 | -0.28 | 7.8E-01 | 0 |
| Moving Around Using Transportation | 6 | 637646 | 734011 | 210343 | 0.218 | -0.24 | 8.1E-01 | 0 |
| Temperament and Personality Functions | 1059 | 813625 | 1198890 | 150027 | 0.324 | -0.22 | 8.3E-01 | 0 |
| Maligrant Neoplasms of Brain | 22 | 661039 | 1044619 | 281435 | 0.247 | -0.21 | 8.4E-01 | 0 |
| Structures related to movement, unspecified | 926 | 745888 | 1141189 | 177289 | 0.275 | -0.20 | 8.4E-01 | 0 |
| Type 2 Diabetes Mellitus | 525 | 763615 | 1102403 | 373483 | 0.291 | -0.16 | 8.7E-01 | 0 |
| Inguinal Hernia | 14 | 820809 | 1125520 | 73491 | 0.355 | -0.16 | 8.7E-01 | 0 |
| Thyroiditis | 75 | 783140 | 1139713 | 327709 | 0.332 | -0.13 | 9.0E-01 | 0 |
| Breast and Nipple | 2 | 721569 | 763804 | 679333 | 0.243 | -0.04 | 9.7E-01 | 0 |
| Migraine | 13 | 800740 | 1061773 | 406448 | 0.316 | -0.02 | 9.8E-01 | 0 |
| Heart Functions | 525 | 733008 | 1154463 | 216580 | 0.274 | -0.01 | 9.9E-01 | 0 |
| Gastro-Oesophageal Reflux Disease | 1 | 940400 | 940400 | 940400 | 0.448 | 0.07 | 9.5E-01 | 0 |
| Structure of Head and Neck Region | 59 | 853965 | 1291146 | 404181 | 0.334 | 0.08 | 9.4E-01 | 0 |
| Superficial Injuries Involving Multiple Body Regions | 30 | 832467 | 1106505 | 287249 | 0.329 | 0.09 | 9.3E-01 | 0 |
| Atrial Fibrillation and Flutter | 186 | 759971 | 1150789 | 337274 | 0.275 | 0.16 | 8.7E-01 | 0 |
| Thyrotoxicosis | 10 | 740721 | 1123445 | 424717 | 0.273 | 0.19 | 8.5E-01 | 0 |
| Hyperkinetic Disorders | 11 | 892855 | 1212943 | 497249 | 0.426 | 0.22 | 8.3E-01 | 0 |
| Sensations Associated with Genital and Reproductive Functions | 4 | 762909 | 983202 | 424111 | 0.314 | 0.25 | 8.0E-01 | 0 |
| Assets | 33 | 773968 | 1187630 | 93906 | 0.272 | 0.30 | 7.6E-01 | 0 |
| Endocrine Gland Functions | 114 | 721111 | 928598 | 419004 | 0.264 | 0.31 | 7.6E-01 | 0 |
| Malignant Neoplasms | 47 | 718645 | 1024981 | 280109 | 0.244 | 0.40 | 6.9E-01 | 0 |
| Discomfort associated with menopause | 8 | 744573 | 1367268 | 434703 | 0.269 | 0.43 | 6.7E-01 | 0 |
| Disorders related to length of gestation and fetal growth | 7 | 924228 | 1220385 | 660623 | 0.291 | 0.43 | 6.7E-01 | 0 |
| Functions of Hair | 717 | 775515 | 1109543 | 300378 | 0.292 | 0.44 | 6.6E-01 | 0 |
| Height | 3231 | 777000 | 1145015 | 223525 | 0.288 | 0.44 | 6.6E-01 | 0 |
| Presence of Cardiac and Vascular Implants and Grafts | 8 | 918658 | 1054393 | 808126 | 0.236 | 0.53 | 5.9E-01 | 0 |
| Procreation Functions | 58 | 712216 | 1147821 | 73834 | 0.250 | 0.56 | 5.8E-01 | 0 |

| | | | | | | | | |
|---|---|---|---|---|---|---|---|---|
| Other Congenital Malformations of the Dygestive System | 6 | 818603 | 1039886 | 334516 | 0.215 | 0.56 | 5.7E-01 | 0 |
| Persons with Potential Health Hazards Related to Socioeconomic and Psychosocial Circumstances | 3 | 1023008 | 1167565 | 854443 | 0.228 | 0.58 | 5.6E-01 | 0 |
| Haematological System Functions | 26 | 452849 | 863486 | 86642 | 0.009 | 0.59 | 5.5E-01 | 0 |
| Essential (Primary) Hypertension | 210 | 796706 | 1132637 | 155420 | 0.296 | 0.61 | 5.4E-01 | 0 |
| Structure of Hair | 443 | 634015 | 1088624 | 115796 | 0.204 | 0.62 | 5.4E-01 | 0 |
| Family History of Certain Disabilities and Chronic Diseases Leading to Disablement | 5 | 1077720 | 1138860 | 851270 | 0.406 | 0.64 | 5.2E-01 | 0 |
| Angina Pectoris | 26 | 881300 | 1172282 | 321236 | 0.292 | 0.64 | 5.2E-01 | 0 |
| Other degenerative diseases of the nervous system | 3 | 719763 | 804408 | 387647 | 0.106 | 0.65 | 5.2E-01 | 0 |
| Mental and Behavioural Disorders Due to Use of Tobacco | 431 | 835703 | 1218840 | 110210 | 0.320 | 0.67 | 5.0E-01 | 0 |
| Water, Mineral and Electrolyte Balance Functions | 1387 | 767985 | 1127289 | 292274 | 0.289 | 0.69 | 4.9E-01 | 0 |
| Hypertrichosis | 3 | 1145635 | 1332681 | 912813 | 0.459 | 0.70 | 4.8E-01 | 0 |
| Unspecified Haematuria | 4 | 841234 | 1251904 | 394397 | 0.188 | 0.72 | 4.7E-01 | 0 |
| Disorders of Lipoprotein Metabolism and Other lipidaemias | 21 | 818555 | 1079695 | 214394 | 0.222 | 0.75 | 4.5E-01 | 0 |
| Products or Substances for Personal Consumption | 599 | 803743 | 1150583 | 266960 | 0.298 | 0.81 | 4.2E-01 | 0 |
| Cholelithiasis | 15 | 881008 | 1127658 | 80472 | 0.249 | 0.81 | 4.2E-01 | 0 |
| Other Rheumatoid Arthritis | 65 | 870620 | 1184573 | 431055 | 0.288 | 0.81 | 4.2E-01 | 0 |
| Systemic Lupus Erythematosus | 59 | 746580 | 1215221 | 276415 | 0.226 | 0.85 | 4.0E-01 | 0 |
| Pervasive Developmental Disorders | 3 | 1164070 | 1190443 | 970190 | 0.346 | 0.87 | 3.8E-01 | 0 |
| Blood Pressure Functions | 753 | 793230 | 1143225 | 187424 | 0.296 | 0.88 | 3.8E-01 | 0 |
| Bipolar Affective Disorder | 12 | 984339 | 1212962 | 495684 | 0.357 | 0.90 | 3.7E-01 | 0 |
| Food | 623 | 771528 | 1174960 | 85350 | 0.289 | 0.97 | 3.3E-01 | 0 |
| Structure of Trunk | 37 | 955958 | 1314745 | 427510 | 0.338 | 0.98 | 3.2E-01 | 0 |
| Functions of the Cardiovascular System | 6 | 1052319 | 1121252 | 894963 | 0.275 | 1.01 | 3.1E-01 | 0 |
| Hematological System Functions | 32 | 917845 | 1122026 | 347349 | 0.278 | 1.10 | 2.7E-01 | 0 |
| Pain in Abdomen and Pelvic | 1 | 845510 | 845510 | 845510 | 0.062 | 1.13 | 2.6E-01 | 0 |
| Diverticular Disease of Intestine | 68 | 901831 | 1167521 | 451644 | 0.260 | 1.13 | 2.6E-01 | 0 |
| Mental and Behavioural Disorders Due to Use of Cannabinoid | 7 | 904135 | 1237786 | 168857 | 0.126 | 1.18 | 2.4E-01 | 0 |
| Epilepsy | 21 | 1021760 | 1437625 | 722640 | 0.321 | 1.19 | 2.3E-01 | 0 |
| Type 1 Diabetes Mellitus | 75 | 911180 | 1213378 | 360986 | 0.283 | 1.21 | 2.3E-01 | 0 |
| Other Cataract | 6 | 1048938 | 1397449 | 342022 | 0.259 | 1.25 | 2.1E-01 | 0 |
| Control of Voluntary Movement Functions | 1 | 1525045 | 1525045 | 1525045 | 0.492 | 1.28 | 2.0E-01 | 0 |
| All-Cause Mortality | 121 | 616138 | 1020063 | 105694 | 0.083 | 1.40 | 1.6E-01 | 0 |
| Sleep Disorders | 74 | 970969 | 1183357 | 116859 | 0.335 | 1.41 | 1.6E-01 | 0 |
| Atopic Dermatitis | 21 | 1079008 | 1342805 | 531513 | 0.382 | 1.47 | 1.4E-01 | 0 |
| Conduct Disorder | 2 | 830614 | 1216454 | 444773 | 0.064 | 1.51 | 1.3E-01 | 0 |
| Structures Related to the Genitourinary and Reproductive Systems, Other Specified | 1 | 1613123 | 1613123 | 1613123 | 0.336 | 1.52 | 1.3E-01 | 0 |
| Problems Related to Lifestyle | 1 | 1529998 | 1529998 | 1529998 | 0.222 | 1.56 | 1.2E-01 | 0 |
| Sensation of Pain | 59 | 948753 | 1173568 | 241821 | 0.316 | 1.57 | 1.2E-01 | 0 |
| Personal History of Medical Treatment | 3 | 1053010 | 1791330 | 1049563 | 0.099 | 1.59 | 1.1E-01 | 0 |
| Noninfective entertitis and Colitis | 424 | 635919 | 1052809 | 221098 | 0.186 | 1.60 | 1.1E-01 | 0 |
| Parkinson Disease | 28 | 1028789 | 1293166 | 481049 | 0.264 | 1.64 | 1.0E-01 | 0 |
| Alzheimer disease | 38 | 958383 | 1224558 | 200994 | 0.277 | 1.68 | 9.4E-02 | 0 |
| Diaphragmatic Hernia | 3 | 1398910 | 1434606 | 1106865 | 0.314 | 1.72 | 8.6E-02 | 0 |
| Weight Maintenance Functions | 7748 | 807771 | 1186445 | 211168 | 0.301 | 1.86 | 6.2E-02 | 0 |
| Internal Derangement of Knee | 1 | 1884003 | 1884003 | 1884003 | 0.411 | 1.97 | 4.9E-02 | 0 |
| Renal Failure | 89 | 661478 | 1090430 | 207140 | 0.046 | 2.02 | 4.3E-02 | 0 |
| Structure of Areas of Skin | 397 | 635863 | 1091155 | 82497 | 0.142 | 2.21 | 2.7E-02 | 0 |

| | | | | | | | | |
|---|---|---|---|---|---|---|---|---|
| Potential Health Hazards Related to Socioeconomic and Psychosocial Circumstances | 138 | 953724 | 1197395 | 298738 | 0.292 | 2.51 | 1.2E-02 | 0 |
| Spinal Muscular Strophy and Related Syndrome | 10 | 1199350 | 1369743 | 217373 | 0.204 | 2.58 | 9.8E-03 | 0 |
| Immunological System Functions | 4076 | 752814 | 1135338 | 220139 | 0.256 | 2.89 | 3.9E-03 | 0 |
| General Metabolic Functions | 1043 | 667463 | 1076378 | 178804 | 0.140 | 4.54 | 5.8E-06 | 1 |
| Malignant Neoplasms of Breast | 215 | 487202 | 977286 | 101615 | 0.026 | 8.23 | 1.9E-16 | 1 |

| Trait | N SNPs | Median age | Age 75% upper bound | Age 25% lower bound | Median MAF | Z-score | P value | Bonferroni |
|---|---|---|---|---|---|---|---|---|
| Use of sun/uv protection | 28 | 185084 | 995476 | 56749 | 0.323 | -4.28 | 1.9E-05 | 1 |
| Educational attainment | 465 | 637523 | 1143858 | 111495 | 0.306 | -3.87 | 1.1E-04 | 0 |
| Fluid intelligence score | 47 | 459465 | 1121114 | 81313 | 0.386 | -3.76 | 1.7E-04 | 0 |
| Fluid intelligence test - Fluid intelligence score | 13 | 82009 | 877208 | 57133 | 0.377 | -3.73 | 1.9E-04 | 0 |
| Schizophrenia/Bipolar disorder | 86 | 499765 | 1068881 | 81502 | 0.328 | -3.62 | 2.9E-04 | 0 |
| Depression - Lifetime number of depressed periods | 352 | 19939 | 35225 | 13316 | 0.004 | -3.27 | 1.1E-03 | 0 |
| Legs-leg fat ratio (female) | 143 | 506280 | 1133235 | 61213 | 0.284 | -3.27 | 1.1E-03 | 0 |
| Hip circumference (female) | 33 | 382215 | 1035158 | 79540 | 0.354 | -3.09 | 2.0E-03 | 0 |
| Average weekly beer plus cider intake | 18 | 297008 | 990736 | 50671 | 0.383 | -2.81 | 4.9E-03 | 0 |
| Electronic device use - Plays computer games | 26 | 400844 | 1033210 | 90037 | 0.334 | -2.75 | 6.0E-03 | 0 |
| Impedance measures - Impedance of leg (left) | 447 | 682503 | 1155285 | 124853 | 0.301 | -2.66 | 7.9E-03 | 0 |
| Body Mass Index (male) | 362 | 675208 | 1140772 | 189329 | 0.317 | -2.66 | 7.9E-03 | 0 |
| Left hippocampus | 8 | 76249 | 633694 | 42060 | 0.348 | -2.60 | 9.5E-03 | 0 |
| Average weekly intake of other alcoholic drinks | 252 | 19507 | 33543 | 13699 | 0.005 | -2.59 | 9.6E-03 | 0 |
| Self-rated health | 6 | 71608 | 174189 | 56422 | 0.373 | -2.58 | 9.9E-03 | 0 |
| Cup area | 51 | 61882 | 743535 | 19603 | 0.070 | -2.52 | 1.2E-02 | 0 |
| Carpal tunnel syndrome | 12 | 89668 | 637349 | 43850 | 0.224 | -2.46 | 1.4E-02 | 0 |
| Depression | 40 | 536490 | 958729 | 68205 | 0.317 | -2.46 | 1.4E-02 | 0 |
| Neutrophil percentage of white cells (three-way meta) | 92 | 480893 | 1133946 | 81624 | 0.248 | -2.43 | 1.5E-02 | 0 |
| Social support - Leisure/social activities: Religious group | 41 | 543158 | 1075395 | 84723 | 0.320 | -2.37 | 1.8E-02 | 0 |
| Glucocorticoids | 12 | 406321 | 922544 | 156640 | 0.358 | -2.37 | 1.8E-02 | 0 |
| Asthma (fixed effect model) | 21 | 465193 | 1011813 | 136209 | 0.354 | -2.35 | 1.9E-02 | 0 |
| Length of time at current address | 5 | 123446 | 1197513 | 64740 | 0.349 | -2.35 | 1.9E-02 | 0 |
| Total body BMD (30-45 years old) | 7 | 176519 | 813318 | 71063 | 0.282 | -2.35 | 1.9E-02 | 0 |
| Beef intake | 6 | 176647 | 724122 | 78859 | 0.400 | -2.33 | 2.0E-02 | 0 |
| Father's age at death | 3 | 45592 | 592226 | 44533 | 0.331 | -2.32 | 2.0E-02 | 0 |
| Mean Putamen | 3 | 75055 | 662063 | 63835 | 0.419 | -2.31 | 2.1E-02 | 0 |
| Reticulocyte count (three-way meta) | 106 | 559426 | 1071839 | 100936 | 0.285 | -2.28 | 2.3E-02 | 0 |
| Number of full brothers | 28 | 71124 | 1129505 | 27961 | 0.150 | -2.27 | 2.3E-02 | 0 |
| Right vessel | 2 | 133376 | 148423 | 118329 | 0.400 | -2.24 | 2.5E-02 | 0 |
| Number of depression episodes | 233 | 27446 | 80091 | 16053 | 0.002 | -2.22 | 2.7E-02 | 0 |
| Time spent watching television (TV) | 90 | 610313 | 1210895 | 91628 | 0.349 | -2.21 | 2.7E-02 | 0 |
| Lifetime number of sexual partners | 191 | 18224 | 34894 | 11733 | 0.004 | -2.19 | 2.8E-02 | 0 |
| Waist circumference (female) | 30 | 461045 | 957167 | 224040 | 0.278 | -2.16 | 3.1E-02 | 0 |
| Handedness (chirality/laterality) | 2 | 40610 | 48418 | 32802 | 0.268 | -2.14 | 3.2E-02 | 0 |
| Intracranial Volume | 6 | 86245 | 470171 | 44418 | 0.223 | -2.13 | 3.3E-02 | 0 |
| Worrier / anxious feelings | 44 | 572380 | 946318 | 75698 | 0.326 | -2.13 | 3.3E-02 | 0 |
| Peptide::gamma-glutamyl::gamma-glutamyltyrosine | 4 | 118692 | 357965 | 97545 | 0.333 | -2.12 | 3.4E-02 | 0 |
| Impedance measures - Impedance of leg (right) | 458 | 709886 | 1144254 | 164261 | 0.307 | -2.12 | 3.4E-02 | 0 |
| Brain stem | 16 | 249259 | 719940 | 68144 | 0.276 | -2.11 | 3.4E-02 | 0 |
| Interpolated Age of participant when non-cancer illness first diagnosed | 6 | 144197 | 630851 | 89913 | 0.258 | -2.11 | 3.5E-02 | 0 |
| Left lateral ventricle | 9 | 182377 | 474765 | 97114 | 0.389 | -2.11 | 3.5E-02 | 0 |
| Interpolated Year when non-cancer illness first diagnosed | 6 | 144197 | 630851 | 89913 | 0.258 | -2.10 | 3.6E-02 | 0 |
| Nucleotide::Pyrimidine metabolism, uracil containing::uridine | 3 | 71641 | 618145 | 70840 | 0.338 | -2.08 | 3.7E-02 | 0 |
| Doctor diagnosed asthma | 14 | 244103 | 755139 | 51321 | 0.227 | -2.04 | 4.1E-02 | 0 |
| Cereal type: Muesli | 8 | 66258 | 307335 | 40122 | 0.253 | -2.03 | 4.2E-02 | 0 |
| Amino acid::Valine, leucine and isoleucine metabolism::alpha-hydroxyisovalerate | 2 | 199588 | 257028 | 142149 | 0.412 | -2.01 | 4.4E-02 | 0 |
| Cognitive performance | 153 | 676120 | 1082510 | 92619 | 0.321 | -2.01 | 4.4E-02 | 0 |
| Frequency of drinking alcohol | 6 | 221736 | 436783 | 67929 | 0.378 | -2.00 | 4.6E-02 | 0 |
| Major dietary changes in the last 5 years | 9 | 229329 | 889053 | 69644 | 0.395 | -1.99 | 4.7E-02 | 0 |
| Bread type: Wholemeal or wholegrain | 14 | 393296 | 1151058 | 77991 | 0.342 | -1.98 | 4.8E-02 | 0 |
| Number of vehicles in household | 5 | 100707 | 138743 | 58540 | 0.249 | -1.95 | 5.1E-02 | 0 |
| Legs-leg fat ratio (male) | 20 | 385794 | 1069323 | 78229 | 0.308 | -1.95 | 5.1E-02 | 0 |
| Posterior limb of internal capsule mode of anisotropy | 6 | 248493 | 506239 | 154241 | 0.326 | -1.94 | 5.2E-02 | 0 |
| L5 timing | 4 | 46067 | 359654 | 40801 | 0.284 | -1.94 | 5.2E-02 | 0 |

| | | | | | | | | |
|---|---|---|---|---|---|---|---|---|
| Left precentral | 5 | 63278 | 867898 | 62100 | 0.241 | -1.93 | 5.3E-02 | 0 |
| Left vessel | 3 | 224226 | 605667 | 170768 | 0.408 | -1.91 | 5.6E-02 | 0 |
| Illnesses of father: Lung cancer | 4 | 51982 | 283413 | 43957 | 0.218 | -1.89 | 5.8E-02 | 0 |
| Worrier / anxious feelings (WORRY) | 35 | 562025 | 903823 | 74896 | 0.292 | -1.89 | 5.9E-02 | 0 |
| Left cuneus | 3 | 74601 | 624878 | 66138 | 0.345 | -1.88 | 6.1E-02 | 0 |
| Smoking cessation | 5 | 125509 | 875095 | 89821 | 0.280 | -1.87 | 6.1E-02 | 0 |
| Trunk-trunk fat ratio (female) | 180 | 602353 | 1172396 | 86587 | 0.267 | -1.85 | 6.4E-02 | 0 |
| Body Mass Index - Child | 17 | 292630 | 513870 | 76367 | 0.205 | -1.84 | 6.5E-02 | 0 |
| Fornix (column and body of fornix) fractional anisotropy | 6 | 247492 | 1190918 | 106453 | 0.363 | -1.82 | 6.9E-02 | 0 |
| Reported occurrences of cancer | 2 | 37570 | 37957 | 37182 | 0.158 | -1.81 | 7.1E-02 | 0 |
| Reason for glasses/contact lenses: For short-sightedness, i.e. only or mainly for distance viewing such as driving, cinema etc (called 'myopia') | 38 | 614573 | 1209086 | 284724 | 0.347 | -1.80 | 7.2E-02 | 0 |
| Depression - Recent feelings of tiredness or low enery | 3 | 66860 | 774248 | 55193 | 0.345 | -1.80 | 7.2E-02 | 0 |
| Lipid::Lysolipid::1-eicosatrienoylglycerophosphocholine* | 2 | 124936 | 151070 | 98801 | 0.297 | -1.80 | 7.2E-02 | 0 |
| Amino acid::Urea cycle; arginine-, proline-, metabolism::citrulline | 3 | 135205 | 540796 | 96099 | 0.337 | -1.79 | 7.3E-02 | 0 |
| Male-specific factors - Hair/balding pattern: Pattern 2 | 45 | 597060 | 1010020 | 217307 | 0.316 | -1.76 | 7.9E-02 | 0 |
| Weight (male) | 9 | 382215 | 1187630 | 64317 | 0.391 | -1.76 | 7.9E-02 | 0 |
| Ratio of visceral-tosubcutaneous adipose tissue volume (adjusted for BMI) | 1 | 43488 | 43488 | 43488 | 0.440 | -1.75 | 7.9E-02 | 0 |
| Number of full sisters | 33 | 200859 | 761838 | 19366 | 0.118 | -1.75 | 8.0E-02 | 0 |
| First PC of the four riskly behaviours | 64 | 694644 | 1229573 | 198975 | 0.354 | -1.74 | 8.3E-02 | 0 |
| Suffer from 'nerves' (SUF-NERV) | 5 | 79348 | 586548 | 63709 | 0.262 | -1.73 | 8.4E-02 | 0 |
| Own or rent accommodation lived in: Own outright (by you or someone in your household) | 1 | 55775 | 55775 | 55775 | 0.491 | -1.73 | 8.4E-02 | 0 |
| Reason for reducing amount of alcohol drunk: Health precaution | 2 | 179762 | 243505 | 116020 | 0.326 | -1.73 | 8.4E-02 | 0 |
| Diagnoses - main ICD10: M16 Osteoarthritis of hip | 10 | 343796 | 693370 | 106550 | 0.259 | -1.72 | 8.5E-02 | 0 |
| Maternal smoking around birth | 14 | 364589 | 708594 | 50867 | 0.227 | -1.71 | 8.7E-02 | 0 |
| College completion | 3 | 95032 | 422132 | 83753 | 0.204 | -1.71 | 8.8E-02 | 0 |
| Glycine level (male) | 12 | 102899 | 563625 | 27396 | 0.122 | -1.70 | 8.8E-02 | 0 |
| Usual side of head for mobile phone use | 2 | 129096 | 151873 | 106319 | 0.240 | -1.70 | 8.9E-02 | 0 |
| Left lateral orbitofrontal | 1 | 84968 | 84968 | 84968 | 0.453 | -1.68 | 9.3E-02 | 0 |
| Conscientiousness (NEO-FFI) | 1 | 75362 | 75362 | 75362 | 0.416 | -1.67 | 9.4E-02 | 0 |
| Types of physical activity in last 4 weeks: Other exercises (eg: swimming, cycling, keep fit, bowling) | 9 | 379430 | 638813 | 256820 | 0.280 | -1.67 | 9.5E-02 | 0 |
| Lipid::Fatty acid, dicarboxylate::3-carboxy-4-methyl-5-propyl-2-furanpropanoate (CMPF) | 1 | 83759 | 83759 | 83759 | 0.422 | -1.67 | 9.6E-02 | 0 |
| Ulcerative Colitis | 26 | 541460 | 892418 | 190238 | 0.321 | -1.65 | 9.8E-02 | 0 |
| Acceleration average | 8 | 147557 | 1199616 | 22268 | 0.171 | -1.65 | 9.8E-02 | 0 |
| Amino acid::Valine, leucine and isoleucine metabolism::isovalerylcarnitine | 2 | 89642 | 110107 | 69178 | 0.279 | -1.65 | 9.9E-02 | 0 |
| Posterior corona radiata axial diusivities | 2 | 208609 | 296688 | 120530 | 0.277 | -1.65 | 9.9E-02 | 0 |
| Diagnoses - secondary ICD10: E66 Overweight and obesity | 3 | 227601 | 386760 | 139639 | 0.375 | -1.64 | 1.0E-01 | 0 |
| Traumatic events - Avoided activities or situations because of previous stressful experience in past month | 2 | 218348 | 299278 | 137419 | 0.277 | -1.64 | 1.0E-01 | 0 |
| Reason for glasses/contact lenses: For 'astigmatism' | 1 | 37587 | 37587 | 37587 | 0.360 | -1.62 | 1.0E-01 | 0 |
| Duration of moderate activity | 2 | 266121 | 349282 | 182960 | 0.329 | -1.62 | 1.1E-01 | 0 |
| Non-oily fish intake | 10 | 216765 | 424298 | 83193 | 0.190 | -1.62 | 1.1E-01 | 0 |
| Illnesses of mother: Chronic bronchitis/emphysema | 1 | 49775 | 49775 | 49775 | 0.326 | -1.61 | 1.1E-01 | 0 |
| Left handed | 1 | 40135 | 40135 | 40135 | 0.373 | -1.61 | 1.1E-01 | 0 |
| Obesity class 3 | 2 | 360811 | 426552 | 295070 | 0.437 | -1.60 | 1.1E-01 | 0 |
| Reticulocyte fraction of red cells (three-way meta) | 111 | 614160 | 1092275 | 108049 | 0.284 | -1.60 | 1.1E-01 | 0 |
| Childhood sunburn occasions | 107 | 19893 | 41180 | 12236 | 0.003 | -1.59 | 1.1E-01 | 0 |
| Overall health rating | 47 | 643415 | 1153283 | 125768 | 0.347 | -1.59 | 1.1E-01 | 0 |
| Body of corpus callosum mode of anisotropy | 2 | 261746 | 368255 | 155236 | 0.328 | -1.59 | 1.1E-01 | 0 |
| Verbal-numerical reasoning | 27 | 625013 | 1025048 | 106612 | 0.342 | -1.59 | 1.1E-01 | 0 |
| Impedance measures - Trunk fat mass | 421 | 732603 | 1144175 | 126329 | 0.304 | -1.59 | 1.1E-01 | 0 |
| Oily fish intake | 50 | 581130 | 1111271 | 65904 | 0.280 | -1.59 | 1.1E-01 | 0 |
| Duration walking for pleasure | 1 | 48891 | 48891 | 48891 | 0.314 | -1.59 | 1.1E-01 | 0 |
| Number of cigarettes smoked per day | 1 | 53928 | 53928 | 53928 | 0.377 | -1.58 | 1.1E-01 | 0 |
| Number of children fathered (male) | 6 | 31772 | 84455 | 28499 | 0.117 | -1.58 | 1.1E-01 | 0 |
| Right lingual | 2 | 328520 | 445639 | 211401 | 0.399 | -1.58 | 1.1E-01 | 0 |
| Sleep efficiency | 4 | 34179 | 44624 | 29291 | 0.127 | -1.57 | 1.2E-01 | 0 |
| Osteoarthritis | 29 | 568683 | 1044870 | 210350 | 0.349 | -1.57 | 1.2E-01 | 0 |

| | | | | | | | | |
|---|---|---|---|---|---|---|---|---|
| Intraocular pressure | 36 | 37645 | 531498 | 21006 | 0.019 | -1.56 | 1.2E-01 | 0 |
| Drinks per week | 47 | 624858 | 1257091 | 61640 | 0.328 | -1.56 | 1.2E-01 | 0 |
| Cofactors and vitamins::Pantothenate and CoA metabolism::pantothenate | 1 | 72590 | 72590 | 72590 | 0.306 | -1.54 | 1.2E-01 | 0 |
| ::::X-12092 | 3 | 124129 | 477847 | 98449 | 0.258 | -1.52 | 1.3E-01 | 0 |
| Impedance measures - Leg fat-free mass (left) | 513 | 728310 | 1145570 | 114349 | 0.308 | -1.52 | 1.3E-01 | 0 |
| BRCA1/2-negative breast cancer | 2 | 50715 | 51587 | 49842 | 0.153 | -1.52 | 1.3E-01 | 0 |
| Cancer register - Type of cancer: ICD10: C44 Other and unspecified malignant neoplasm of skin | 28 | 376049 | 766590 | 99630 | 0.194 | -1.52 | 1.3E-01 | 0 |
| ::::X-12855 | 1 | 85556 | 85556 | 85556 | 0.339 | -1.51 | 1.3E-01 | 0 |
| Amino acid::Creatine metabolism::creatinine | 1 | 168503 | 168503 | 168503 | 0.493 | -1.51 | 1.3E-01 | 0 |
| ::::X-11491 | 2 | 74898 | 87034 | 62762 | 0.145 | -1.50 | 1.3E-01 | 0 |
| Anterior limb of internal capsule fractional anisotropy | 2 | 143717 | 197707 | 89727 | 0.172 | -1.49 | 1.4E-01 | 0 |
| Celiac disease | 38 | 533051 | 1057677 | 126359 | 0.223 | -1.49 | 1.4E-01 | 0 |
| Miserableness (MIS) | 32 | 508027 | 1096294 | 83254 | 0.295 | -1.49 | 1.4E-01 | 0 |
| Mouth/teeth dental problems: Mouth ulcers | 27 | 528883 | 969554 | 173262 | 0.242 | -1.48 | 1.4E-01 | 0 |
| Treatment/medication code: lisinopril | 3 | 298338 | 693795 | 174102 | 0.386 | -1.47 | 1.4E-01 | 0 |
| Job involves shift work | 3 | 150650 | 995752 | 108329 | 0.418 | -1.47 | 1.4E-01 | 0 |
| Hearing difficulty/problems | 25 | 560375 | 1064688 | 183421 | 0.349 | -1.46 | 1.4E-01 | 0 |
| Left isthmus cingulate | 4 | 306495 | 721156 | 49208 | 0.280 | -1.44 | 1.5E-01 | 0 |
| Right parahippocampal | 1 | 140254 | 140254 | 140254 | 0.361 | -1.44 | 1.5E-01 | 0 |
| Primary biliary cirrhosis | 17 | 471018 | 837185 | 72818 | 0.285 | -1.44 | 1.5E-01 | 0 |
| Osteoarthritis of knee | 8 | 526686 | 1263233 | 396324 | 0.376 | -1.44 | 1.5E-01 | 0 |
| Insomnia (male) | 4 | 33202 | 35982 | 30368 | 0.100 | -1.43 | 1.5E-01 | 0 |
| Pain type(s) experienced in last month: Neck or shoulder pain | 5 | 108061 | 787273 | 30242 | 0.330 | -1.43 | 1.5E-01 | 0 |
| ::::X-11334 | 1 | 213132 | 213132 | 213132 | 0.458 | -1.43 | 1.5E-01 | 0 |
| Bilateral oophorectomy (both ovaries removed) (female) | 6 | 351934 | 726293 | 99518 | 0.207 | -1.42 | 1.5E-01 | 0 |
| Extreme chronotype | 3 | 130692 | 203013 | 84443 | 0.192 | -1.42 | 1.6E-01 | 0 |
| Illnesses of father: Stroke | 1 | 40700 | 40700 | 40700 | 0.218 | -1.41 | 1.6E-01 | 0 |
| Amino acid::Lysine metabolism::pipecolate | 1 | 41635 | 41635 | 41635 | 0.254 | -1.40 | 1.6E-01 | 0 |
| Fasting glucose main effect (adjusted for BMI) | 20 | 484658 | 1055847 | 117412 | 0.286 | -1.40 | 1.6E-01 | 0 |
| Average across all tracts fractional anisotropy | 3 | 61894 | 460397 | 46179 | 0.240 | -1.40 | 1.6E-01 | 0 |
| Lean body mass (appendicular) | 1 | 50637 | 50637 | 50637 | 0.275 | -1.39 | 1.6E-01 | 0 |
| Immature fraction of reticulocytes (two-way meta) | 79 | 614160 | 1059983 | 108206 | 0.320 | -1.39 | 1.6E-01 | 0 |
| Estimated glomerular filtration rate based on serum creatinine (non-diabetic) | 61 | 680143 | 1000742 | 147526 | 0.309 | -1.39 | 1.6E-01 | 0 |
| Frequency of light DIY in last 4 weeks | 1 | 41613 | 41613 | 41613 | 0.216 | -1.39 | 1.7E-01 | 0 |
| Time spent driving | 2 | 256746 | 372968 | 140524 | 0.244 | -1.39 | 1.7E-01 | 0 |
| Treatment/medication code: levothyroxine sodium | 47 | 665375 | 1100020 | 345475 | 0.338 | -1.38 | 1.7E-01 | 0 |
| Usual walking pace | 40 | 598774 | 1174531 | 130097 | 0.316 | -1.37 | 1.7E-01 | 0 |
| Hands-free device/speakerphone use with mobile phone in last 3 month | 1 | 51287 | 51287 | 51287 | 0.241 | -1.37 | 1.7E-01 | 0 |
| Primary sclerosing cholangitis | 1 | 160339 | 160339 | 160339 | 0.321 | -1.37 | 1.7E-01 | 0 |
| Frequency of other exercises in last 4 weeks | 1 | 247745 | 247745 | 247745 | 0.484 | -1.36 | 1.7E-01 | 0 |
| Fornix (column and body of fornix) mean diusivities | 4 | 247492 | 714670 | 145666 | 0.284 | -1.36 | 1.7E-01 | 0 |
| Blood urea nitrogen | 156 | 655081 | 1009751 | 162559 | 0.289 | -1.36 | 1.7E-01 | 0 |
| Milk type used: Skimmed | 5 | 323715 | 685363 | 67861 | 0.286 | -1.36 | 1.7E-01 | 0 |
| Fluid intelligence test - FI3 : word interpolation | 5 | 444250 | 1316713 | 183142 | 0.310 | -1.36 | 1.7E-01 | 0 |
| Body Mass Index (female <= 50 yrs) | 18 | 574599 | 1202961 | 192863 | 0.326 | -1.36 | 1.8E-01 | 0 |
| Fornix (column and body of fornix) axial diusivities | 4 | 247492 | 396774 | 145666 | 0.308 | -1.35 | 1.8E-01 | 0 |
| Extreme Body Mass Index | 6 | 339575 | 621187 | 72550 | 0.239 | -1.35 | 1.8E-01 | 0 |
| Leucine | 4 | 226721 | 696908 | 55258 | 0.252 | -1.35 | 1.8E-01 | 0 |
| Energy::Krebs cycle::succinylcarnitine | 6 | 485817 | 948119 | 96404 | 0.392 | -1.33 | 1.8E-01 | 0 |
| Arms-arm fat ratio (female) | 115 | 696565 | 1014908 | 103687 | 0.317 | -1.33 | 1.8E-01 | 0 |
| Phltrium width | 1 | 69269 | 69269 | 69269 | 0.290 | -1.33 | 1.8E-01 | 0 |
| Impedance measures - Leg predicted mass (right) | 522 | 740891 | 1158870 | 114098 | 0.311 | -1.33 | 1.8E-01 | 0 |
| Number of sexual partners | 72 | 721611 | 1162502 | 188661 | 0.355 | -1.33 | 1.8E-01 | 0 |
| Lipid::Medium chain fatty acid::caprate (10:0) | 2 | 412974 | 526115 | 299834 | 0.403 | -1.33 | 1.9E-01 | 0 |
| Asthma, hay fever or eczema | 57 | 605760 | 1192875 | 215168 | 0.270 | -1.32 | 1.9E-01 | 0 |
| Water intake | 24 | 624239 | 1013437 | 81388 | 0.335 | -1.31 | 1.9E-01 | 0 |

| | | | | | | | | |
|---|---|---|---|---|---|---|---|---|
| Nonischemic cardiomyopathy | 1 | 76263 | 76263 | 76263 | 0.223 | -1.31 | 1.9E-01 | 0 |
| Immunoglobulin A deficiency | 1 | 199865 | 199865 | 199865 | 0.397 | -1.31 | 1.9E-01 | 0 |
| Miserableness | 31 | 542315 | 958953 | 94404 | 0.308 | -1.31 | 1.9E-01 | 0 |
| Right transverse temporal | 1 | 87790 | 87790 | 87790 | 0.277 | -1.30 | 1.9E-01 | 0 |
| Xenobiotics::Sugar, sugar substitute, starch::erythritol | 1 | 91344 | 91344 | 91344 | 0.233 | -1.29 | 2.0E-01 | 0 |
| Posterior limb of internal capsule fractional anisotropy | 8 | 343513 | 806896 | 78111 | 0.322 | -1.29 | 2.0E-01 | 0 |
| Dried fruit intake | 7 | 79302 | 845108 | 28059 | 0.150 | -1.29 | 2.0E-01 | 0 |
| Inflammatory Bowel Disease | 129 | 677545 | 1122875 | 149469 | 0.324 | -1.28 | 2.0E-01 | 0 |
| Depression - Trouble falling or staying asleep. or sleeping too much | 1 | 96727 | 96727 | 96727 | 0.274 | -1.28 | 2.0E-01 | 0 |
| Corticospinal tract fractional anisotropy | 2 | 186521 | 272234 | 100808 | 0.178 | -1.27 | 2.0E-01 | 0 |
| Posterior limb of internal capsule axial diusivities | 6 | 562368 | 1140271 | 372944 | 0.447 | -1.27 | 2.0E-01 | 0 |
| Parkinson disease | 5 | 284488 | 1019892 | 115794 | 0.215 | -1.27 | 2.0E-01 | 0 |
| Chronic kidney disease | 54 | 505864 | 1183621 | 85285 | 0.289 | -1.27 | 2.0E-01 | 0 |
| Coronary artery disease | 59 | 547253 | 942629 | 149970 | 0.223 | -1.27 | 2.0E-01 | 0 |
| Lipid::Carnitine metabolism::X-13431--nonanoylcarnitine* | 2 | 178248 | 238599 | 117897 | 0.234 | -1.26 | 2.1E-01 | 0 |
| Subjective well being | 1 | 117951 | 117951 | 117951 | 0.249 | -1.26 | 2.1E-01 | 0 |
| Superior longitudinal fasciculus axial diusivities | 4 | 227778 | 738823 | 63880 | 0.189 | -1.26 | 2.1E-01 | 0 |
| Impedance measures - Leg predicted mass (left) | 510 | 740891 | 1172431 | 120680 | 0.307 | -1.25 | 2.1E-01 | 0 |
| Distance between home and job workplace | 56 | 17572 | 26267 | 11515 | 0.005 | -1.25 | 2.1E-01 | 0 |
| Lipid::Carnitine metabolism::laurylcarnitine | 1 | 216130 | 216130 | 216130 | 0.314 | -1.24 | 2.2E-01 | 0 |
| Amino acid::Valine, leucine and isoleucine metabolism::levulinate (4-oxovalerate) | 1 | 309273 | 309273 | 309273 | 0.448 | -1.23 | 2.2E-01 | 0 |
| Granulocyte percentage of myeloid white cells (three-way meta) | 114 | 607731 | 1072918 | 75980 | 0.264 | -1.22 | 2.2E-01 | 0 |
| Mean Hippocampus | 2 | 81421 | 105141 | 57701 | 0.130 | -1.22 | 2.2E-01 | 0 |
| Tobacco smoking | 4 | 338065 | 777539 | 40820 | 0.252 | -1.21 | 2.3E-01 | 0 |
| Sitting height | 652 | 704985 | 1130723 | 114463 | 0.277 | -1.21 | 2.3E-01 | 0 |
| Right posterior cingulate | 1 | 21558 | 21558 | 21558 | 0.122 | -1.21 | 2.3E-01 | 0 |
| Disposition index | 1 | 134748 | 134748 | 134748 | 0.275 | -1.20 | 2.3E-01 | 0 |
| Disposition index (adjusted for BMI) | 1 | 134748 | 134748 | 134748 | 0.275 | -1.20 | 2.3E-01 | 0 |
| Energy::Oxidative phosphorylation::acetylphosphate | 2 | 305180 | 406679 | 203682 | 0.270 | -1.20 | 2.3E-01 | 0 |
| Cereal type: Oat cereal (e.g. Ready Brek, porridge) | 1 | 323715 | 323715 | 323715 | 0.403 | -1.20 | 2.3E-01 | 0 |
| Frequency of stair climbing in last 4 weeks | 6 | 504911 | 915942 | 413849 | 0.376 | -1.20 | 2.3E-01 | 0 |
| Impedance measures - Body fat percentage | 387 | 742268 | 1157380 | 111967 | 0.302 | -1.20 | 2.3E-01 | 0 |
| Immunosuppressants | 1 | 24454 | 24454 | 24454 | 0.102 | -1.19 | 2.3E-01 | 0 |
| Neuroticism (IRT) | 1 | 30621 | 30621 | 30621 | 0.195 | -1.19 | 2.3E-01 | 0 |
| Amino acid::Glycine, serine and threonine metabolism::serine | 2 | 461674 | 633210 | 290139 | 0.359 | -1.18 | 2.4E-01 | 0 |
| Frequency of solarium/sunlamp use | 44 | 17000 | 27919 | 12505 | 0.003 | -1.18 | 2.4E-01 | 0 |
| Asthma (child-onset) | 64 | 660470 | 1093462 | 299600 | 0.279 | -1.18 | 2.4E-01 | 0 |
| Depression - Recent poor appetite or overeting | 1 | 323715 | 323715 | 323715 | 0.403 | -1.18 | 2.4E-01 | 0 |
| Positive affect (MA GWAMA) | 82 | 720200 | 1165619 | 175515 | 0.324 | -1.18 | 2.4E-01 | 0 |
| Left fusiform | 4 | 234976 | 643218 | 18827 | 0.216 | -1.17 | 2.4E-01 | 0 |
| Coronary artery disease (SOFT definition including angina) | 19 | 437615 | 643464 | 204980 | 0.179 | -1.17 | 2.4E-01 | 0 |
| Age started oral contraceptive pill (female) | 4 | 240136 | 512628 | 42467 | 0.160 | -1.16 | 2.5E-01 | 0 |
| Milk type used: Semi-skimmed | 2 | 438239 | 623706 | 252773 | 0.385 | -1.16 | 2.5E-01 | 0 |
| Total body BMD (60 or older) | 17 | 449127 | 928078 | 87540 | 0.282 | -1.16 | 2.5E-01 | 0 |
| Meconium ileus in cystic fibrosis | 1 | 40478 | 40478 | 40478 | 0.111 | -1.15 | 2.5E-01 | 0 |
| Left transverse temporal | 1 | 338685 | 338685 | 338685 | 0.446 | -1.15 | 2.5E-01 | 0 |
| Diagnoses - secondary ICD10: E03 Other hypothyroidism | 40 | 657540 | 1108684 | 410239 | 0.325 | -1.15 | 2.5E-01 | 0 |
| ::::X-11529 | 1 | 50626 | 50626 | 50626 | 0.143 | -1.15 | 2.5E-01 | 0 |
| Numeric memory test - Maximum digits remembered correctly | 6 | 509016 | 1608511 | 62861 | 0.340 | -1.15 | 2.5E-01 | 0 |
| Number of older siblings | 1 | 48252 | 48252 | 48252 | 0.168 | -1.14 | 2.5E-01 | 0 |
| Lipid::Fatty acid, dicarboxylate::octadecanedioate | 1 | 49898 | 49898 | 49898 | 0.194 | -1.13 | 2.6E-01 | 0 |
| Xenobiotics::Xanthine metabolism::7-methylxanthine | 1 | 169008 | 169008 | 169008 | 0.294 | -1.13 | 2.6E-01 | 0 |
| Chest pain or discomfort walking normally | 1 | 51049 | 51049 | 51049 | 0.136 | -1.12 | 2.6E-01 | 0 |
| Left thalamus proper | 1 | 46532 | 46532 | 46532 | 0.102 | -1.12 | 2.6E-01 | 0 |
| Right cuneus | 5 | 257275 | 543158 | 151319 | 0.248 | -1.12 | 2.6E-01 | 0 |
| Social support - Leisure/social activities: Other group activity | 5 | 309555 | 984592 | 256188 | 0.223 | -1.12 | 2.6E-01 | 0 |

| | | | | | | | | |
|---|---|---|---|---|---|---|---|---|
| Peak insulin response (adjusted for BMI) | 3 | 134748 | 419900 | 78553 | 0.260 | -1.12 | 2.6E-01 | 0 |
| ::::X-13496 | 2 | 328038 | 463909 | 192168 | 0.281 | -1.12 | 2.6E-01 | 0 |
| Fibrinogen | 1 | 57474 | 57474 | 57474 | 0.184 | -1.12 | 2.6E-01 | 0 |
| Amyotrophic lateral sclerosis (meta-analysis) | 3 | 87092 | 675340 | 51090 | 0.176 | -1.12 | 2.6E-01 | 0 |
| Smoking status: Previous vs Current | 1 | 56814 | 56814 | 56814 | 0.106 | -1.12 | 2.6E-01 | 0 |
| Amino acid::Alanine and aspartate metabolism::N-acetylalanine | 1 | 62630 | 62630 | 62630 | 0.139 | -1.12 | 2.6E-01 | 0 |
| Gout | 2 | 288515 | 415423 | 161608 | 0.166 | -1.11 | 2.7E-01 | 0 |
| Impedance measures - Basal metabolic rate | 574 | 730324 | 1160342 | 110691 | 0.303 | -1.11 | 2.7E-01 | 0 |
| Impedance measures - Leg fat-free mass (right) | 514 | 752146 | 1176354 | 120680 | 0.313 | -1.11 | 2.7E-01 | 0 |
| Sleep durataion (conditioning sex and BMI) | 8 | 415726 | 1113755 | 47191 | 0.237 | -1.11 | 2.7E-01 | 0 |
| Cereal type: Biscuit cereal (e.g. Weetabix) | 1 | 63526 | 63526 | 63526 | 0.118 | -1.11 | 2.7E-01 | 0 |
| Amino acid::Glutamate metabolism::glutamine | 1 | 66319 | 66319 | 66319 | 0.181 | -1.11 | 2.7E-01 | 0 |
| Superior corona radiata axial diusivities | 4 | 257590 | 583121 | 68972 | 0.218 | -1.10 | 2.7E-01 | 0 |
| Age when periods started (menarche) (female) | 184 | 712341 | 1122476 | 196464 | 0.303 | -1.10 | 2.7E-01 | 0 |
| FVC | 398 | 761659 | 1180471 | 212957 | 0.315 | -1.10 | 2.7E-01 | 0 |
| Waist-hip ratio (male > 50 yrs, adjusted for BMI) | 3 | 514313 | 794198 | 286605 | 0.471 | -1.10 | 2.7E-01 | 0 |
| Ever taken oral contraceptive pill (female) | 2 | 15392 | 16507 | 14276 | 0.007 | -1.10 | 2.7E-01 | 0 |
| Diabetes (diagnosed by doctor) | 54 | 615524 | 1079431 | 158394 | 0.278 | -1.10 | 2.7E-01 | 0 |
| Morningness | 117 | 678165 | 1160463 | 120689 | 0.283 | -1.10 | 2.7E-01 | 0 |
| Anterior corona radiata axial diusivities | 3 | 303995 | 416168 | 168223 | 0.265 | -1.09 | 2.8E-01 | 0 |
| Creatinine | 4 | 397978 | 715134 | 145730 | 0.267 | -1.09 | 2.8E-01 | 0 |
| Monocyte count (three-way meta) | 130 | 648628 | 1104121 | 101781 | 0.273 | -1.09 | 2.8E-01 | 0 |
| Overweight | 13 | 555613 | 992923 | 323375 | 0.225 | -1.08 | 2.8E-01 | 0 |
| Exposure to tobacco smoke outside home | 3 | 28787 | 265987 | 23680 | 0.001 | -1.08 | 2.8E-01 | 0 |
| Reaction time | 39 | 627113 | 991665 | 187568 | 0.307 | -1.08 | 2.8E-01 | 0 |
| ::::X-13741 | 1 | 72422 | 72422 | 72422 | 0.115 | -1.08 | 2.8E-01 | 0 |
| Height SDS (male at age 12) | 1 | 203205 | 203205 | 203205 | 0.298 | -1.08 | 2.8E-01 | 0 |
| Number of stillbirths (female) | 2 | 21325 | 24569 | 18081 | 0.005 | -1.08 | 2.8E-01 | 0 |
| Diurnal inactivity duration | 2 | 447371 | 649368 | 245374 | 0.372 | -1.08 | 2.8E-01 | 0 |
| Morning/evening person (chronotype) | 112 | 667319 | 1162164 | 119263 | 0.262 | -1.08 | 2.8E-01 | 0 |
| Variation in diet | 10 | 520368 | 1001284 | 204209 | 0.368 | -1.08 | 2.8E-01 | 0 |
| Crohn's Disease | 139 | 712183 | 1132675 | 347151 | 0.324 | -1.08 | 2.8E-01 | 0 |
| Reason for glasses/contact lenses: For long-sightedness, i.e. for distance and near, but particularly for near tasks like reading (called 'hypermetropia') | 4 | 401899 | 710946 | 134303 | 0.363 | -1.07 | 2.8E-01 | 0 |
| Obesity class 2 | 10 | 526273 | 640331 | 283569 | 0.304 | -1.07 | 2.8E-01 | 0 |
| Fasting proinsulin (adjusted for fasting insulin, age and sex) | 7 | 499810 | 935071 | 285636 | 0.291 | -1.07 | 2.8E-01 | 0 |
| Average weekly spirits intake | 2 | 17094 | 19245 | 14943 | 0.005 | -1.07 | 2.8E-01 | 0 |
| Systolic Blood Pressure (automated reading) | 208 | 742521 | 1123735 | 139634 | 0.319 | -1.07 | 2.9E-01 | 0 |
| Amino acid::Polyamine metabolism::X-03056--N-[3-(2-Oxopyrrolidin-1-yl)propyl]acetamide | 3 | 209582 | 343057 | 128382 | 0.192 | -1.06 | 2.9E-01 | 0 |
| Major depressive disorder | 3 | 421838 | 627209 | 265592 | 0.466 | -1.06 | 2.9E-01 | 0 |
| Body Mass Index (female > 50 yrs) | 16 | 610561 | 831621 | 266421 | 0.264 | -1.06 | 2.9E-01 | 0 |
| ::::X-14473 | 1 | 82250 | 82250 | 82250 | 0.151 | -1.06 | 2.9E-01 | 0 |
| Splenium of corpus callosum axial diusivities | 2 | 276832 | 381652 | 172011 | 0.250 | -1.06 | 2.9E-01 | 0 |
| Colorectal cancer | 9 | 507910 | 1064808 | 338440 | 0.329 | -1.06 | 2.9E-01 | 0 |
| Sagittal stratum mean diusivities | 3 | 23901 | 359703 | 23073 | 0.055 | -1.06 | 2.9E-01 | 0 |
| Gray matter | 5 | 370350 | 921158 | 62138 | 0.360 | -1.06 | 2.9E-01 | 0 |
| Comparative height size at age 10 | 457 | 713448 | 1114073 | 168115 | 0.278 | -1.05 | 2.9E-01 | 0 |
| Pancreatic cancer | 3 | 415423 | 739820 | 277350 | 0.310 | -1.05 | 2.9E-01 | 0 |
| Ever depressed for a whole week | 5 | 56776 | 401733 | 42107 | 0.070 | -1.05 | 2.9E-01 | 0 |
| Sagittal stratum radial diusivities | 3 | 23901 | 474499 | 23073 | 0.055 | -1.05 | 2.9E-01 | 0 |
| Body Mass Index (male <= 50 yrs) | 6 | 456843 | 1041314 | 338085 | 0.223 | -1.05 | 2.9E-01 | 0 |
| Had menopause (female) | 28 | 580544 | 1140433 | 109921 | 0.344 | -1.04 | 3.0E-01 | 0 |
| Treatment/medication code: ventolin 100micrograms inhaler | 12 | 618721 | 860567 | 281457 | 0.322 | -1.04 | 3.0E-01 | 0 |
| Right lateral ventricle | 12 | 558718 | 912548 | 195172 | 0.311 | -1.04 | 3.0E-01 | 0 |
| Other sociodemographic factors - Private healthcare | 2 | 32169 | 42186 | 22151 | 0.008 | -1.04 | 3.0E-01 | 0 |
| Lipid::Fatty acid, monohydroxy::2-hydroxypalmitate | 1 | 221208 | 221208 | 221208 | 0.276 | -1.03 | 3.0E-01 | 0 |
| Heart failure | 1 | 102578 | 102578 | 102578 | 0.159 | -1.03 | 3.0E-01 | 0 |

| | | | | | | | | |
|---|---|---|---|---|---|---|---|---|
| Cerebellar vermal lobules VI VII | 12 | 576470 | 843186 | 185615 | 0.303 | -1.02 | 3.1E-01 | 0 |
| Pericardial adipose tissue volume (adjusted for height and weight) | 3 | 249751 | 604247 | 153668 | 0.203 | -1.01 | 3.1E-01 | 0 |
| Seen doctor (GP) for nerves, anxiety, tension or depression | 25 | 654258 | 1139650 | 95918 | 0.370 | -1.00 | 3.2E-01 | 0 |
| Monocyte percentage of white cells (two-way meta) | 90 | 579946 | 1012198 | 52222 | 0.248 | -1.00 | 3.2E-01 | 0 |
| Lipid::Essential fatty acid::dihomo-linolenate (20:3n3 or n6) | 2 | 378892 | 532005 | 225779 | 0.249 | -1.00 | 3.2E-01 | 0 |
| Mother still alive | 4 | 468654 | 992926 | 45778 | 0.269 | -1.00 | 3.2E-01 | 0 |
| Average across all tracts mean diusivities | 5 | 402258 | 466160 | 303995 | 0.265 | -0.99 | 3.2E-01 | 0 |
| Relative age of first facial hair (male) | 78 | 641573 | 1085894 | 148743 | 0.278 | -0.99 | 3.2E-01 | 0 |
| Friendships satisfaction | 3 | 524345 | 856514 | 474481 | 0.384 | -0.99 | 3.2E-01 | 0 |
| 3Lhydroxybutyrate | 1 | 127709 | 127709 | 127709 | 0.157 | -0.98 | 3.3E-01 | 0 |
| Frequency of travelling from home to job workplace | 11 | 13217 | 101009 | 9575 | 0.002 | -0.98 | 3.3E-01 | 0 |
| Trail making test - Duration to complete numeric path (trail #1) | 2 | 54729 | 66145 | 43313 | 0.001 | -0.98 | 3.3E-01 | 0 |
| Vertical cup-disc ratio | 145 | 130496 | 687760 | 25328 | 0.006 | -0.98 | 3.3E-01 | 0 |
| Ratio of bisLallylic bonds to double bonds in lipids | 2 | 279550 | 406535 | 152566 | 0.206 | -0.97 | 3.3E-01 | 0 |
| Thinness | 3 | 33968 | 518521 | 23297 | 0.010 | -0.97 | 3.3E-01 | 0 |
| Acute insulin response (adjusted for BMI) | 2 | 456758 | 617763 | 295753 | 0.302 | -0.96 | 3.4E-01 | 0 |
| Impedance measures - Body Mass Index (BMI) | 401 | 763488 | 1183325 | 244295 | 0.318 | -0.96 | 3.4E-01 | 0 |
| Eye problems/disorders: Cataract | 2 | 277362 | 357695 | 197029 | 0.142 | -0.96 | 3.4E-01 | 0 |
| Lipid::Essential fatty acid::eicosapentaenoate (EPA; 20:5n3) | 1 | 369990 | 369990 | 369990 | 0.311 | -0.93 | 3.5E-01 | 0 |
| Citrate | 4 | 666720 | 896784 | 548718 | 0.468 | -0.93 | 3.5E-01 | 0 |
| Genu of corpus callosum mean diusivities | 5 | 504515 | 740255 | 303995 | 0.306 | -0.93 | 3.5E-01 | 0 |
| Ratio of visceral-tosubcutaneous adipose tissue volume (adjusted for BMI, female) | 1 | 269605 | 269605 | 269605 | 0.283 | -0.93 | 3.5E-01 | 0 |
| Prospective memory test - Time to answer | 1 | 156966 | 156966 | 156966 | 0.132 | -0.93 | 3.5E-01 | 0 |
| Types of transport used (excluding work): Public transport | 4 | 391594 | 917194 | 79860 | 0.201 | -0.92 | 3.6E-01 | 0 |
| Monocyte percentage of white cells (three-way meta) | 119 | 652628 | 1089650 | 89219 | 0.278 | -0.92 | 3.6E-01 | 0 |
| Schizophrenia | 190 | 757061 | 1219503 | 130813 | 0.327 | -0.92 | 3.6E-01 | 0 |
| Longest period of depression | 8 | 25889 | 107296 | 12834 | 0.002 | -0.91 | 3.6E-01 | 0 |
| Heart rate | 10 | 514051 | 1113645 | 78809 | 0.332 | -0.91 | 3.6E-01 | 0 |
| Heart rate variability (pvRSA/HF) | 7 | 406100 | 930565 | 76558 | 0.349 | -0.90 | 3.7E-01 | 0 |
| Sphingomyelins | 7 | 225348 | 1028358 | 41734 | 0.119 | -0.89 | 3.7E-01 | 0 |
| Right rostral middle frontal | 1 | 394190 | 394190 | 394190 | 0.330 | -0.89 | 3.7E-01 | 0 |
| Left pericalcarine | 5 | 414180 | 721700 | 84806 | 0.216 | -0.89 | 3.7E-01 | 0 |
| Left precuneus | 3 | 46213 | 294738 | 46117 | 0.090 | -0.89 | 3.7E-01 | 0 |
| Number in household | 4 | 41094 | 141118 | 22657 | 0.002 | -0.89 | 3.8E-01 | 0 |
| Mineral and other dietary supplements: Zinc | 1 | 472285 | 472285 | 472285 | 0.498 | -0.88 | 3.8E-01 | 0 |
| Social support - Leisure/social activities: Adult education class | 2 | 419401 | 605997 | 232805 | 0.226 | -0.88 | 3.8E-01 | 0 |
| Wheeze or whistling in the chest in last year | 24 | 651598 | 1131414 | 293286 | 0.251 | -0.88 | 3.8E-01 | 0 |
| Non-butter spread type details: Soft (tub) margarine | 1 | 182155 | 182155 | 182155 | 0.145 | -0.88 | 3.8E-01 | 0 |
| Cingulum (cingulate gyrus) mode of anisotropy | 5 | 475135 | 875192 | 49476 | 0.241 | -0.87 | 3.9E-01 | 0 |
| PEF | 265 | 730555 | 1171643 | 165279 | 0.293 | -0.86 | 3.9E-01 | 0 |
| Body of corpus callosum radial diusivities | 1 | 303995 | 303995 | 303995 | 0.265 | -0.86 | 3.9E-01 | 0 |
| Mean corpuscular hemoglobin (two-way meta) | 130 | 697610 | 1211348 | 178649 | 0.274 | -0.86 | 3.9E-01 | 0 |
| Types of transport used (excluding work): Walk | 7 | 572478 | 1078124 | 475506 | 0.303 | -0.86 | 3.9E-01 | 0 |
| Age at menopause | 42 | 739396 | 1113224 | 399028 | 0.343 | -0.86 | 3.9E-01 | 0 |
| Chest pain or discomfort | 2 | 460186 | 648639 | 271733 | 0.311 | -0.85 | 3.9E-01 | 0 |
| Left lateral occipital | 3 | 57675 | 162368 | 46604 | 0.093 | -0.85 | 3.9E-01 | 0 |
| Education - Qualifications | 110 | 713543 | 1110859 | 92875 | 0.318 | -0.85 | 4.0E-01 | 0 |
| Lipid::Essential fatty acid::docosapentaenoate (n3 DPA; 22:5n3) | 1 | 439805 | 439805 | 439805 | 0.320 | -0.85 | 4.0E-01 | 0 |
| ::::X-11469 | 3 | 416563 | 544035 | 250161 | 0.282 | -0.85 | 4.0E-01 | 0 |
| Posterior thalamic radiation (include optic radiation) radial diusivities | 2 | 499750 | 597628 | 401873 | 0.285 | -0.85 | 4.0E-01 | 0 |
| Genu of corpus callosum mode of anisotropy | 1 | 491515 | 491515 | 491515 | 0.419 | -0.84 | 4.0E-01 | 0 |
| Lipid::Lysolipid::1-arachidonoylglycerophosphoethanolamine* | 3 | 395798 | 858888 | 223212 | 0.299 | -0.84 | 4.0E-01 | 0 |
| Fluid intelligence test - FI6 : conditional arithmetic | 1 | 428080 | 428080 | 428080 | 0.325 | -0.84 | 4.0E-01 | 0 |
| Peptide::gamma-glutamyl::gamma-glutamylphenylalanine | 1 | 436477 | 436477 | 436477 | 0.306 | -0.84 | 4.0E-01 | 0 |
| Amino acid::Valine, leucine and isoleucine metabolism::2-methylbutyroylcarnitine | 1 | 209582 | 209582 | 209582 | 0.192 | -0.83 | 4.1E-01 | 0 |
| Prospective memory test - Number of attempts | 13 | 27942 | 768348 | 22606 | 0.002 | -0.83 | 4.1E-01 | 0 |

| | | | | | | | | |
|---|---|---|---|---|---|---|---|---|
| Total body BMD (15 or younger) | 7 | 551500 | 1039185 | 244137 | 0.276 | -0.83 | 4.1E-01 | 0 |
| Posterior thalamic radiation (include optic radiation) mean diusivities | 2 | 499750 | 597628 | 401873 | 0.285 | -0.83 | 4.1E-01 | 0 |
| Eosinophil percentage of granulocytes (two-way meta) | 94 | 675448 | 1120079 | 116859 | 0.259 | -0.82 | 4.1E-01 | 0 |
| Splenium of corpus callosum mean diusivities | 2 | 499750 | 597628 | 401873 | 0.285 | -0.82 | 4.1E-01 | 0 |
| Frequency of needing morning drink of alcohol after heavy drinking session in last year | 16 | 40179 | 56124 | 16995 | 0.001 | -0.82 | 4.1E-01 | 0 |
| Dysmenorrhea pain severity | 2 | 514651 | 746114 | 283188 | 0.314 | -0.82 | 4.1E-01 | 0 |
| Fornix (column and body of fornix) radial diusivities | 4 | 403400 | 850859 | 243339 | 0.262 | -0.82 | 4.1E-01 | 0 |
| MYCN amplification neuroblastoma | 2 | 547140 | 798676 | 295603 | 0.343 | -0.82 | 4.1E-01 | 0 |
| Left amygdala | 4 | 536671 | 727938 | 276053 | 0.323 | -0.81 | 4.2E-01 | 0 |
| Fasting glucose | 15 | 564913 | 974776 | 167229 | 0.280 | -0.81 | 4.2E-01 | 0 |
| Eosinophil count (three-way meta) | 114 | 638685 | 1069538 | 81190 | 0.274 | -0.80 | 4.2E-01 | 0 |
| ::::X-11261 | 1 | 209582 | 209582 | 209582 | 0.192 | -0.80 | 4.2E-01 | 0 |
| Frequency of consuming six or more units of alcohol | 6 | 306314 | 768224 | 34029 | 0.155 | -0.80 | 4.2E-01 | 0 |
| Average weekly fortified wine intake | 10 | 44812 | 190484 | 14535 | 0.002 | -0.80 | 4.2E-01 | 0 |
| Lipid::Lysolipid::2-stearoylglycerophosphocholine* | 1 | 213700 | 213700 | 213700 | 0.120 | -0.80 | 4.3E-01 | 0 |
| ::::X-02249 | 1 | 347683 | 347683 | 347683 | 0.281 | -0.79 | 4.3E-01 | 0 |
| Age completed full time education | 6 | 484754 | 1301597 | 142163 | 0.236 | -0.79 | 4.3E-01 | 0 |
| Chronotype (continuous) | 5 | 532973 | 965650 | 130692 | 0.355 | -0.78 | 4.4E-01 | 0 |
| Age started wearing glasses or contact lenses | 100 | 738014 | 1074956 | 148191 | 0.305 | -0.78 | 4.4E-01 | 0 |
| Interferon-gamma | 1 | 13015 | 13015 | 13015 | 0.010 | -0.78 | 4.4E-01 | 0 |
| Hot drink temperature | 45 | 767540 | 1294605 | 72826 | 0.371 | -0.78 | 4.4E-01 | 0 |
| Menstrual cycle length | 5 | 544048 | 1117228 | 78471 | 0.323 | -0.78 | 4.4E-01 | 0 |
| Nap during day | 75 | 711078 | 1105469 | 130951 | 0.303 | -0.78 | 4.4E-01 | 0 |
| Health satisfaction | 1 | 18576 | 18576 | 18576 | 0.013 | -0.77 | 4.4E-01 | 0 |
| Amino acid::Tryptophan metabolism::X-12100--hydroxytryptophan* | 3 | 552740 | 987160 | 508206 | 0.375 | -0.77 | 4.4E-01 | 0 |
| Duration of heavy DIY | 1 | 22587 | 22587 | 22587 | 0.035 | -0.77 | 4.4E-01 | 0 |
| Illnesses of mother: Lung cancer | 1 | 20992 | 20992 | 20992 | 0.008 | -0.76 | 4.4E-01 | 0 |
| Impedance measures - Weight | 522 | 771358 | 1173641 | 169746 | 0.317 | -0.76 | 4.5E-01 | 0 |
| Life span | 2 | 615808 | 891731 | 339884 | 0.405 | -0.76 | 4.5E-01 | 0 |
| TNF-related apoptosis inducing ligand | 3 | 24513 | 36174 | 22204 | 0.007 | -0.76 | 4.5E-01 | 0 |
| Antihistamines for systemic use | 5 | 433560 | 1317225 | 407193 | 0.257 | -0.76 | 4.5E-01 | 0 |
| Traumatic events - Felt distant from other people in past month | 1 | 26219 | 26219 | 26219 | 0.013 | -0.76 | 4.5E-01 | 0 |
| Alcohol - Alcohol drinker status: Previous vs Current | 1 | 25520 | 25520 | 25520 | 0.032 | -0.76 | 4.5E-01 | 0 |
| Primary angle closure glaucoma | 8 | 336599 | 570258 | 53689 | 0.170 | -0.76 | 4.5E-01 | 0 |
| Depression - Did your sleep change? | 1 | 26515 | 26515 | 26515 | 0.006 | -0.76 | 4.5E-01 | 0 |
| Medial collateral ligament injury | 1 | 26829 | 26829 | 26829 | 0.046 | -0.75 | 4.5E-01 | 0 |
| Cholesterol esters in large VLDL | 10 | 319575 | 805398 | 59010 | 0.124 | -0.75 | 4.5E-01 | 0 |
| Tinnitus | 1 | 27175 | 27175 | 27175 | 0.032 | -0.75 | 4.5E-01 | 0 |
| Type 2 Diabetes (Dominance deviation model) | 1 | 24626 | 24626 | 24626 | 0.028 | -0.75 | 4.5E-01 | 0 |
| Peptide::Polypeptide::HWESASXX* | 1 | 26849 | 26849 | 26849 | 0.089 | -0.75 | 4.5E-01 | 0 |
| Carbohydrate::Glycolysis, gluconeogenesis, pyruvate metabolism::1,5-anhydroglucitol (1,5-AG) | 3 | 427225 | 1101838 | 404371 | 0.267 | -0.75 | 4.5E-01 | 0 |
| Relative age voice broke (male) | 26 | 709688 | 1009222 | 141234 | 0.321 | -0.74 | 4.6E-01 | 0 |
| Basophil count (two-way meta) | 41 | 680480 | 1065250 | 84667 | 0.342 | -0.74 | 4.6E-01 | 0 |
| Peptide::Dipeptide::pro-hydroxy-pro | 1 | 551180 | 551180 | 551180 | 0.489 | -0.74 | 4.6E-01 | 0 |
| Frequency of failure to fulfil normal expectations fue to drinking alcohol in last year | 1 | 29260 | 29260 | 29260 | 0.019 | -0.74 | 4.6E-01 | 0 |
| Hepatocyte growth factor | 1 | 31244 | 31244 | 31244 | 0.066 | -0.74 | 4.6E-01 | 0 |
| Splenium of corpus callosum mode of anisotropy | 1 | 35532 | 35532 | 35532 | 0.093 | -0.73 | 4.6E-01 | 0 |
| Right superior parietal | 4 | 540659 | 764921 | 498629 | 0.429 | -0.73 | 4.6E-01 | 0 |
| Diagnoses - secondary ICD10: Z37 Outcome of delivery | 1 | 33618 | 33618 | 33618 | 0.018 | -0.73 | 4.6E-01 | 0 |
| Coffee type: Decaffeinated coffee (any type) | 1 | 32190 | 32190 | 32190 | 0.074 | -0.73 | 4.7E-01 | 0 |
| Rotator cuff injury | 1 | 34759 | 34759 | 34759 | 0.047 | -0.73 | 4.7E-01 | 0 |
| Number of unsuccessful stop-smoking attempts | 13 | 43876 | 423055 | 24088 | 0.004 | -0.72 | 4.7E-01 | 0 |
| Frequency of heavy DIY in last 4 weeks | 1 | 35995 | 35995 | 35995 | 0.005 | -0.72 | 4.7E-01 | 0 |
| Impedance measures - Whole body fat mass | 412 | 781795 | 1143120 | 142431 | 0.317 | -0.72 | 4.7E-01 | 0 |
| Reason for reducing amount of alcohol drunk: Other reason | 2 | 594386 | 865441 | 323332 | 0.374 | -0.71 | 4.8E-01 | 0 |
| Total cholesterol in IDL | 12 | 435135 | 1095218 | 95473 | 0.220 | -0.71 | 4.8E-01 | 0 |

| | | | | | | | | |
|---|---|---|---|---|---|---|---|---|
| Macrophage inflammatory protein-1 (CCL4) | 2 | 249325 | 361553 | 137098 | 0.086 | -0.71 | 4.8E-01 | 0 |
| Interleukin-12p70 | 3 | 657310 | 764339 | 383500 | 0.425 | -0.71 | 4.8E-01 | 0 |
| Pairs matching test - Number of incorrect matches in round | 2 | 135922 | 197432 | 74411 | 0.001 | -0.70 | 4.8E-01 | 0 |
| ::::X-12749 | 1 | 49556 | 49556 | 49556 | 0.081 | -0.70 | 4.8E-01 | 0 |
| Alcohol intake frequency | 70 | 724966 | 1125026 | 129270 | 0.338 | -0.70 | 4.8E-01 | 0 |
| Hair colour (natural, before greying): Red | 11 | 499715 | 898259 | 56085 | 0.212 | -0.70 | 4.8E-01 | 0 |
| Mean Amygdala | 1 | 50604 | 50604 | 50604 | 0.012 | -0.70 | 4.8E-01 | 0 |
| Comparative body size at age 10 | 172 | 740809 | 1201341 | 137596 | 0.304 | -0.70 | 4.8E-01 | 0 |
| Depressive symptoms (univariate) | 53 | 767688 | 1141800 | 149863 | 0.347 | -0.69 | 4.9E-01 | 0 |
| ::::X-11204 | 1 | 395798 | 395798 | 395798 | 0.299 | -0.69 | 4.9E-01 | 0 |
| Neutrophil percentage of granulocytes (two-way meta) | 92 | 678844 | 1125098 | 161238 | 0.262 | -0.69 | 4.9E-01 | 0 |
| Male-specific factors - Hair/balding pattern: Pattern 4 | 145 | 703688 | 1127210 | 143498 | 0.262 | -0.69 | 4.9E-01 | 0 |
| Neutrophil count (three-way meta) | 93 | 678315 | 1180640 | 104136 | 0.274 | -0.69 | 4.9E-01 | 0 |
| Age at first birth (female) | 5 | 629393 | 1276083 | 430185 | 0.386 | -0.69 | 4.9E-01 | 0 |
| Right medial orbitofrontal | 1 | 55574 | 55574 | 55574 | 0.065 | -0.68 | 5.0E-01 | 0 |
| Retrolenticular part of internal capsule radial diusivities | 6 | 388821 | 640041 | 97264 | 0.253 | -0.68 | 5.0E-01 | 0 |
| FEV1/FCV | 6 | 415331 | 570311 | 315943 | 0.200 | -0.68 | 5.0E-01 | 0 |
| Father still alive | 3 | 361718 | 625385 | 200031 | 0.150 | -0.68 | 5.0E-01 | 0 |
| Exposure to tobacco smoke at home | 1 | 62071 | 62071 | 62071 | 0.014 | -0.67 | 5.0E-01 | 0 |
| Average across all tracts radial diusivities | 4 | 516526 | 761518 | 243521 | 0.253 | -0.67 | 5.0E-01 | 0 |
| Subcutaneous adipose tissue volume (female) | 1 | 58604 | 58604 | 58604 | 0.042 | -0.67 | 5.0E-01 | 0 |
| Myeloid white cell count (three-way meta) | 96 | 704283 | 1220718 | 123745 | 0.275 | -0.67 | 5.0E-01 | 0 |
| Bipolar and major depression status | 1 | 66277 | 66277 | 66277 | 0.065 | -0.66 | 5.1E-01 | 0 |
| Amino acid::Phenylalanine & tyrosine metabolism::phenylacetylglutamine | 1 | 410852 | 410852 | 410852 | 0.232 | -0.66 | 5.1E-01 | 0 |
| Double bonds in fatty acids | 3 | 548893 | 713123 | 521474 | 0.347 | -0.66 | 5.1E-01 | 0 |
| Glycine level | 33 | 579855 | 1108530 | 64591 | 0.275 | -0.65 | 5.1E-01 | 0 |
| ::::X-13548 | 2 | 451773 | 501705 | 401840 | 0.195 | -0.65 | 5.2E-01 | 0 |
| Traumatic events - Witnessed sudden violent death | 1 | 295505 | 295505 | 295505 | 0.131 | -0.65 | 5.2E-01 | 0 |
| Pain type(s) experienced in last month: Stomach or abdominal pain | 2 | 650809 | 888447 | 413171 | 0.378 | -0.64 | 5.2E-01 | 0 |
| Inferior fronto-occipital fasciculus axial diusivities | 4 | 473609 | 582851 | 347378 | 0.270 | -0.64 | 5.2E-01 | 0 |
| Thyroid-stimulating hormone (male) | 21 | 676045 | 919705 | 602028 | 0.270 | -0.64 | 5.2E-01 | 0 |
| ::::X-11550 | 1 | 527550 | 527550 | 527550 | 0.324 | -0.64 | 5.2E-01 | 0 |
| Lipid::Medium chain fatty acid::caprylate (8:0) | 2 | 658515 | 894426 | 422604 | 0.391 | -0.64 | 5.2E-01 | 0 |
| Phospholipids in medium LDL | 9 | 385130 | 818555 | 106456 | 0.129 | -0.64 | 5.2E-01 | 0 |
| Pulse pressure | 20 | 394819 | 988123 | 96934 | 0.152 | -0.64 | 5.2E-01 | 0 |
| Anterior limb of internal capsule mode of anisotropy | 3 | 63578 | 113774 | 47473 | 0.044 | -0.64 | 5.2E-01 | 0 |
| Body Mass Index (male > 50 yrs) | 18 | 614885 | 947845 | 132192 | 0.277 | -0.63 | 5.3E-01 | 0 |
| Cancer register - Histology of cancer tumour: Adenocarcinoma, NOS | 10 | 565555 | 852391 | 77790 | 0.235 | -0.63 | 5.3E-01 | 0 |
| Waist-hip ratio (male <= 50 yrs, adjusted for BMI) | 1 | 592133 | 592133 | 592133 | 0.475 | -0.63 | 5.3E-01 | 0 |
| Alcohol dependency (fixed effect model for unrelated genotyped individuals) | 1 | 81986 | 81986 | 81986 | 0.045 | -0.63 | 5.3E-01 | 0 |
| Number of children ever born (male) | 1 | 432875 | 432875 | 432875 | 0.253 | -0.63 | 5.3E-01 | 0 |
| Potassium | 6 | 232177 | 424758 | 51569 | 0.085 | -0.62 | 5.3E-01 | 0 |
| Alcohol dependency (random effect model for unrelated genotyped individuals) | 1 | 81986 | 81986 | 81986 | 0.045 | -0.62 | 5.3E-01 | 0 |
| Impedance measures - Trunk fat percentage | 348 | 747969 | 1182900 | 103698 | 0.295 | -0.62 | 5.3E-01 | 0 |
| Weight (female) | 10 | 675399 | 1111026 | 315026 | 0.276 | -0.62 | 5.3E-01 | 0 |
| Prothrombin time | 8 | 554565 | 1070068 | 373716 | 0.271 | -0.62 | 5.3E-01 | 0 |
| Lipid::Carnitine metabolism::cis-4-decenoyl carnitine | 2 | 625054 | 894417 | 355691 | 0.305 | -0.62 | 5.3E-01 | 0 |
| Positive affect (univariate) | 8 | 523256 | 1148566 | 94318 | 0.228 | -0.62 | 5.4E-01 | 0 |
| ::::X-13215 | 1 | 603415 | 603415 | 603415 | 0.455 | -0.61 | 5.4E-01 | 0 |
| Interleukin-10 | 2 | 694316 | 712819 | 675813 | 0.450 | -0.61 | 5.4E-01 | 0 |
| Total brain volume | 8 | 512325 | 1109881 | 224513 | 0.221 | -0.61 | 5.4E-01 | 0 |
| Childhood obesity | 6 | 673271 | 1101319 | 319310 | 0.394 | -0.61 | 5.4E-01 | 0 |
| Non-cancer illness code, self-reported: diabetes | 35 | 631385 | 1039403 | 127300 | 0.275 | -0.61 | 5.4E-01 | 0 |
| Corticospinal tract mode of anisotropy | 1 | 604068 | 604068 | 604068 | 0.448 | -0.60 | 5.5E-01 | 0 |
| Diagnoses - secondary ICD10: M19 Other and unspecified osteoarthritis | 1 | 313033 | 313033 | 313033 | 0.188 | -0.60 | 5.5E-01 | 0 |
| Left superior parietal | 6 | 483566 | 711248 | 166031 | 0.184 | -0.60 | 5.5E-01 | 0 |

| | | | | | | | | |
|---|---|---|---|---|---|---|---|---|
| Granulocyte percentage of myeloid white cells (two-way meta) | 95 | 667925 | 1116856 | 93867 | 0.263 | -0.59 | 5.5E-01 | 0 |
| Blood clot, DVT, bronchitis, emphysema, asthma, rhinitis, eczema, allergy diagnosed by doctor: Hayfever, allergic rhinitis or eczema | 111 | 690818 | 1120401 | 99927 | 0.272 | -0.59 | 5.5E-01 | 0 |
| Morning person (binary) | 116 | 730095 | 1180721 | 106761 | 0.293 | -0.58 | 5.6E-01 | 0 |
| Lymphocyte percentage of white cells (three-way meta) | 95 | 670950 | 1136449 | 94757 | 0.258 | -0.58 | 5.6E-01 | 0 |
| Lipid::Bile acid metabolism::glycochenodeoxycholate | 1 | 97843 | 97843 | 97843 | 0.075 | -0.58 | 5.6E-01 | 0 |
| Ever smoked regulary | 100 | 793695 | 1193218 | 160938 | 0.334 | -0.58 | 5.6E-01 | 0 |
| Right cerebellum white matter | 13 | 684965 | 1317335 | 399468 | 0.350 | -0.58 | 5.6E-01 | 0 |
| Diagnoses - secondary ICD10: I48 Atrial fibrillation and flutter | 18 | 482194 | 960957 | 85545 | 0.182 | -0.58 | 5.6E-01 | 0 |
| Pulse rate (automated reading) | 202 | 723834 | 1153959 | 196015 | 0.286 | -0.58 | 5.6E-01 | 0 |
| Spread type: Butter/spreadable butter | 8 | 706016 | 1029411 | 349474 | 0.441 | -0.58 | 5.7E-01 | 0 |
| Anxiety - Recent trouble relaxing | 2 | 441872 | 653636 | 230109 | 0.214 | -0.57 | 5.7E-01 | 0 |
| Drive faster than motorway speed limit | 21 | 634753 | 988613 | 213280 | 0.278 | -0.57 | 5.7E-01 | 0 |
| Tea intake | 21 | 648410 | 1164618 | 306338 | 0.286 | -0.56 | 5.7E-01 | 0 |
| Zinc sulfate turbidity test | 2 | 713658 | 1014365 | 412950 | 0.459 | -0.56 | 5.7E-01 | 0 |
| Number of treatments/medications taken | 47 | 754590 | 1160480 | 264317 | 0.348 | -0.56 | 5.7E-01 | 0 |
| Monocyte count (two-way meta) | 112 | 683720 | 1150709 | 99829 | 0.289 | -0.56 | 5.7E-01 | 0 |
| Amino acid::Glutamate metabolism::pyroglutamine* | 4 | 639446 | 1027879 | 362622 | 0.318 | -0.56 | 5.8E-01 | 0 |
| Fatty acid length | 1 | 459733 | 459733 | 459733 | 0.206 | -0.56 | 5.8E-01 | 0 |
| Erectile dysfunction | 1 | 453883 | 453883 | 453883 | 0.246 | -0.56 | 5.8E-01 | 0 |
| Impedance measures - Leg fat percentage (right) | 378 | 761860 | 1166589 | 211315 | 0.305 | -0.56 | 5.8E-01 | 0 |
| Amino acid::Tryptophan metabolism::tryptophan | 17 | 856617 | 1022270 | 525145 | 0.420 | -0.56 | 5.8E-01 | 0 |
| Illnesses of siblings: Severe depression | 1 | 109402 | 109402 | 109402 | 0.052 | -0.55 | 5.8E-01 | 0 |
| Amino acid::Phenylalanine & tyrosine metabolism::X-11423--O-sulfo-L-tyrosine | 2 | 366162 | 524465 | 207859 | 0.142 | -0.55 | 5.8E-01 | 0 |
| Anterior limb of internal capsule radial diusivities | 2 | 575501 | 835419 | 315582 | 0.301 | -0.55 | 5.9E-01 | 0 |
| Hair colour (natural, before greying): Light brown | 65 | 598353 | 929055 | 65472 | 0.266 | -0.54 | 5.9E-01 | 0 |
| Right paracentral | 1 | 345645 | 345645 | 345645 | 0.150 | -0.54 | 5.9E-01 | 0 |
| ::::X-11315 | 2 | 425939 | 531712 | 320167 | 0.124 | -0.53 | 5.9E-01 | 0 |
| Amino acid::Valine, leucine and isoleucine metabolism::beta-hydroxyisovalerate | 1 | 350585 | 350585 | 350585 | 0.132 | -0.53 | 6.0E-01 | 0 |
| ::::X-01911 | 1 | 646865 | 646865 | 646865 | 0.481 | -0.53 | 6.0E-01 | 0 |
| Non-cancer illness code, self-reported: angina | 13 | 607173 | 911330 | 151638 | 0.262 | -0.53 | 6.0E-01 | 0 |
| Prospective memory test - Duration screen displayed | 5 | 89317 | 211086 | 20485 | 0.009 | -0.53 | 6.0E-01 | 0 |
| ::::X-13069 | 1 | 123964 | 123964 | 123964 | 0.025 | -0.52 | 6.0E-01 | 0 |
| How are people in household related to participant: Husband, wife or partner | 1 | 490698 | 490698 | 490698 | 0.293 | -0.52 | 6.1E-01 | 0 |
| Cofactors and vitamins::Hemoglobin and porphyrin metabolism::X-11793--oxidized bilirubin* | 1 | 600395 | 600395 | 600395 | 0.320 | -0.52 | 6.1E-01 | 0 |
| Coronary artery disease and high-density lipoprotein cholesterol (bivariate) | 144 | 725998 | 1125248 | 217187 | 0.283 | -0.51 | 6.1E-01 | 0 |
| Resting heart rate | 290 | 711600 | 1123018 | 159015 | 0.275 | -0.51 | 6.1E-01 | 0 |
| Fresh fruit intake | 36 | 752959 | 1152049 | 111678 | 0.345 | -0.51 | 6.1E-01 | 0 |
| Eosinophil count (two-way meta) | 105 | 694678 | 1135363 | 122305 | 0.286 | -0.51 | 6.1E-01 | 0 |
| Sum neutrophil eosinophil count (three-way meta) | 100 | 718523 | 1199696 | 146348 | 0.282 | -0.51 | 6.1E-01 | 0 |
| Non-cancer illness code, self-reported: hayfever/allergic rhinitis | 13 | 604133 | 1105033 | 240679 | 0.253 | -0.50 | 6.2E-01 | 0 |
| Cofactors and vitamins::Hemoglobin and porphyrin metabolism::bilirubin (E,Z or Z,E)* | 1 | 600395 | 600395 | 600395 | 0.320 | -0.50 | 6.2E-01 | 0 |
| Obesity class 1 | 16 | 730111 | 1262095 | 383953 | 0.339 | -0.50 | 6.2E-01 | 0 |
| Estimated glomerular filtration rate based on cystain C | 13 | 552465 | 852025 | 113585 | 0.222 | -0.49 | 6.2E-01 | 0 |
| Depression - Recent lack of interenst of pleasure in doing things | 1 | 662578 | 662578 | 662578 | 0.452 | -0.49 | 6.2E-01 | 0 |
| Cofactors and vitamins::Hemoglobin and porphyrin metabolism::bilirubin (Z,Z) | 1 | 600395 | 600395 | 600395 | 0.320 | -0.49 | 6.2E-01 | 0 |
| Vitiligo (late onset) | 5 | 559010 | 665375 | 86496 | 0.397 | -0.49 | 6.3E-01 | 0 |
| Cofactors and vitamins::Hemoglobin and porphyrin metabolism::bilirubin (E,E)* | 1 | 600395 | 600395 | 600395 | 0.320 | -0.49 | 6.3E-01 | 0 |
| ::::X-11530 | 1 | 600395 | 600395 | 600395 | 0.320 | -0.49 | 6.3E-01 | 0 |
| Diagnoses - main ICD10: M17 Osteoarthritis of knee | 3 | 614298 | 1249150 | 476524 | 0.285 | -0.49 | 6.3E-01 | 0 |
| Cofactors and vitamins::Hemoglobin and porphyrin metabolism::biliverdin | 1 | 600395 | 600395 | 600395 | 0.320 | -0.48 | 6.3E-01 | 0 |
| Free thyroxine (FT4, male) | 7 | 719743 | 892615 | 514966 | 0.328 | -0.48 | 6.3E-01 | 0 |
| Amino acid::Tryptophan metabolism::indoleacetate | 1 | 378023 | 378023 | 378023 | 0.116 | -0.48 | 6.3E-01 | 0 |
| Inferior fronto-occipital fasciculus mode of anisotropy | 2 | 710418 | 818784 | 602051 | 0.369 | -0.48 | 6.3E-01 | 0 |
| Pain type(s) experienced in last month: Hip pain | 1 | 144252 | 144252 | 144252 | 0.088 | -0.48 | 6.3E-01 | 0 |
| Mouth/teeth dental problems: Loose teeth | 4 | 420755 | 887251 | 46011 | 0.168 | -0.48 | 6.3E-01 | 0 |
| Diagnoses - secondary ICD10: Z85 Personal history of malignant neoplasm | 1 | 609465 | 609465 | 609465 | 0.392 | -0.47 | 6.4E-01 | 0 |

| | | | | | | | | |
|---|---|---|---|---|---|---|---|---|
| Sum eosinophil basophil count (three-way meta) | 123 | 683713 | 1087115 | 151872 | 0.274 | -0.47 | 6.4E-01 | 0 |
| Alcohol - Alcohol drinker status: Never | 1 | 149328 | 149328 | 149328 | 0.022 | -0.47 | 6.4E-01 | 0 |
| Impedance measures - Impedance of whole body | 494 | 763985 | 1175106 | 134889 | 0.304 | -0.46 | 6.4E-01 | 0 |
| Weight | 538 | 783496 | 1190523 | 187480 | 0.316 | -0.46 | 6.4E-01 | 0 |
| Number of self-reported cancers | 6 | 594358 | 696049 | 186400 | 0.243 | -0.46 | 6.5E-01 | 0 |
| Time spent using computer | 13 | 617728 | 1010805 | 71445 | 0.342 | -0.46 | 6.5E-01 | 0 |
| Glycine level (female) | 4 | 421852 | 962418 | 38242 | 0.108 | -0.46 | 6.5E-01 | 0 |
| 11q deletion neuroblastoma | 3 | 200598 | 395628 | 170245 | 0.077 | -0.45 | 6.5E-01 | 0 |
| ::::X-12627 | 1 | 690760 | 690760 | 690760 | 0.420 | -0.45 | 6.5E-01 | 0 |
| Impedance measures - Arm fat percentage (left) | 363 | 795785 | 1178351 | 250636 | 0.319 | -0.45 | 6.5E-01 | 0 |
| Double eyelid | 2 | 219257 | 228872 | 209641 | 0.054 | -0.45 | 6.6E-01 | 0 |
| Average across all tracts mode of anisotropy | 2 | 597093 | 793964 | 400222 | 0.242 | -0.45 | 6.6E-01 | 0 |
| Impedance measures - Leg fat mass (left) | 342 | 805615 | 1171188 | 163614 | 0.332 | -0.44 | 6.6E-01 | 0 |
| White blood cell count | 27 | 669445 | 1148834 | 231501 | 0.298 | -0.44 | 6.6E-01 | 0 |
| Albumin/globulin ratio | 30 | 598881 | 1226794 | 355130 | 0.249 | -0.44 | 6.6E-01 | 0 |
| Eosinophil percentage of granulocytes (three-way meta) | 111 | 681065 | 1078959 | 83013 | 0.264 | -0.44 | 6.6E-01 | 0 |
| Average across all tracts axial diusivities | 4 | 597609 | 837311 | 460609 | 0.275 | -0.43 | 6.6E-01 | 0 |
| Lipid::Carnitine metabolism::carnitine | 16 | 838394 | 1061705 | 445802 | 0.382 | -0.43 | 6.6E-01 | 0 |
| Treatment/medication code: bendroflumethiazide | 16 | 632978 | 1072324 | 142796 | 0.331 | -0.43 | 6.7E-01 | 0 |
| White matter hyperintensities | 3 | 528005 | 810336 | 416841 | 0.188 | -0.43 | 6.7E-01 | 0 |
| Lipid::Long chain fatty acid::adrenate (22:4n6) | 1 | 631788 | 631788 | 631788 | 0.347 | -0.43 | 6.7E-01 | 0 |
| Suffer from 'nerves' | 9 | 586548 | 1331945 | 79348 | 0.273 | -0.42 | 6.7E-01 | 0 |
| Amino acid::Butanoate metabolism::X-04499--3,4-dihydroxybutyrate | 1 | 403843 | 403843 | 403843 | 0.183 | -0.42 | 6.8E-01 | 0 |
| Impedance measures - Trunk predicted mass | 632 | 750450 | 1194322 | 172464 | 0.286 | -0.42 | 6.8E-01 | 0 |
| Diagnoses - main ICD10: D12 Benign neoplasm of colon, rectum, anus and anal canal | 11 | 622368 | 978608 | 94680 | 0.310 | -0.42 | 6.8E-01 | 0 |
| Facial ageing | 55 | 718705 | 1052721 | 90395 | 0.278 | -0.41 | 6.8E-01 | 0 |
| Broad depression | 7 | 593940 | 769674 | 481033 | 0.226 | -0.41 | 6.8E-01 | 0 |
| Well-being spectrum | 129 | 820555 | 1240205 | 126968 | 0.356 | -0.41 | 6.8E-01 | 0 |
| Monocyte chemotactic protein-1 (CCL2) | 2 | 500645 | 736100 | 265190 | 0.243 | -0.40 | 6.9E-01 | 0 |
| Total-body less head BMD | 4 | 660009 | 985103 | 324646 | 0.268 | -0.40 | 6.9E-01 | 0 |
| Diagnoses - secondary ICD10: J45 Asthma | 21 | 760445 | 1105033 | 303590 | 0.349 | -0.40 | 6.9E-01 | 0 |
| ::::X-14626 | 1 | 426800 | 426800 | 426800 | 0.143 | -0.40 | 6.9E-01 | 0 |
| Male pattern baldness (BOLT LMM non-infinitesimal mixed model) | 347 | 723280 | 1105631 | 179495 | 0.281 | -0.40 | 6.9E-01 | 0 |
| Impedance measures - Arm fat percentage (right) | 356 | 789898 | 1199016 | 156892 | 0.316 | -0.39 | 6.9E-01 | 0 |
| Openness to Experience (NEO-FFI) | 1 | 550570 | 550570 | 550570 | 0.227 | -0.39 | 7.0E-01 | 0 |
| Neuroticism (MA GWAMA) | 138 | 840680 | 1297067 | 433145 | 0.370 | -0.39 | 7.0E-01 | 0 |
| ::::X-12524 | 1 | 424935 | 424935 | 424935 | 0.133 | -0.39 | 7.0E-01 | 0 |
| Impedance measures - Arm fat mass (left) | 299 | 804745 | 1180934 | 380041 | 0.332 | -0.39 | 7.0E-01 | 0 |
| Diagnoses - secondary ICD10: Z86 Personal history of certain other diseases | 3 | 225979 | 1377815 | 131869 | 0.023 | -0.38 | 7.0E-01 | 0 |
| Body of corpus callosum axial diusivities | 3 | 647943 | 1047060 | 407927 | 0.346 | -0.38 | 7.0E-01 | 0 |
| Loneliness, isolation (LONE) | 4 | 690156 | 1055344 | 442737 | 0.381 | -0.38 | 7.0E-01 | 0 |
| Hemoglobin A1C | 9 | 645613 | 900685 | 505020 | 0.269 | -0.38 | 7.0E-01 | 0 |
| Lipid::Fatty acid, dicarboxylate::tetradecanedioate | 1 | 426800 | 426800 | 426800 | 0.143 | -0.38 | 7.0E-01 | 0 |
| Ever used hormone-replacement therapy (HRT) (female) | 5 | 475463 | 1105553 | 66235 | 0.227 | -0.38 | 7.0E-01 | 0 |
| Age-related Macular Degeneration | 30 | 667674 | 1008310 | 215463 | 0.255 | -0.38 | 7.0E-01 | 0 |
| Years since last cervical smear test (female) | 6 | 603016 | 805232 | 178825 | 0.186 | -0.38 | 7.1E-01 | 0 |
| Reaction time test - Mean time to correctly identify matches | 37 | 783943 | 1105768 | 73004 | 0.321 | -0.38 | 7.1E-01 | 0 |
| Interleukin-17 | 1 | 725510 | 725510 | 725510 | 0.409 | -0.37 | 7.1E-01 | 0 |
| ::::X-13429 | 1 | 426800 | 426800 | 426800 | 0.143 | -0.37 | 7.1E-01 | 0 |
| Impedance measures - Arm fat mass (right) | 316 | 800265 | 1193901 | 321911 | 0.325 | -0.37 | 7.1E-01 | 0 |
| Fornix (column and body of fornix) mode of anisotropy | 2 | 487471 | 713441 | 261502 | 0.237 | -0.37 | 7.1E-01 | 0 |
| Platelet distribution width (two-way meta) | 120 | 694173 | 1115410 | 158507 | 0.250 | -0.36 | 7.2E-01 | 0 |
| Granulocyte count (three-way meta) | 100 | 737218 | 1215357 | 146348 | 0.289 | -0.36 | 7.2E-01 | 0 |
| Free cholesterol in large LDL | 10 | 453360 | 937892 | 173578 | 0.168 | -0.36 | 7.2E-01 | 0 |
| Total cholesterol in very large HDL | 8 | 606231 | 1088621 | 528099 | 0.251 | -0.36 | 7.2E-01 | 0 |
| Sleep duration (mean) | 7 | 568635 | 664266 | 43752 | 0.289 | -0.36 | 7.2E-01 | 0 |

| | | | | | | | | |
|---|---|---|---|---|---|---|---|---|
| Impedance measures - Trunk fat-free mass | 632 | 750450 | 1190621 | 170107 | 0.288 | -0.36 | 7.2E-01 | 0 |
| Types of transport used (excluding work): Car/motor vehicle | 1 | 579423 | 579423 | 579423 | 0.227 | -0.35 | 7.3E-01 | 0 |
| Basophil percentage of white cells (three-way meta) | 43 | 678015 | 1041583 | 115795 | 0.275 | -0.35 | 7.3E-01 | 0 |
| Amino acid::Alanine and aspartate metabolism::asparagine | 3 | 809285 | 889586 | 803198 | 0.447 | -0.35 | 7.3E-01 | 0 |
| Coffee type: Instant coffee | 3 | 386028 | 710076 | 275317 | 0.187 | -0.35 | 7.3E-01 | 0 |
| Total body BMD | 61 | 781903 | 1203333 | 339343 | 0.326 | -0.35 | 7.3E-01 | 0 |
| Impedance measures - Arm fat-free mass (left) | 540 | 750139 | 1186552 | 122847 | 0.296 | -0.35 | 7.3E-01 | 0 |
| Vitiligo | 30 | 740455 | 1176916 | 116180 | 0.315 | -0.35 | 7.3E-01 | 0 |
| ::::X-11381 | 1 | 448098 | 448098 | 448098 | 0.167 | -0.35 | 7.3E-01 | 0 |
| Alcohol usually taken with meals | 13 | 809173 | 1017803 | 70284 | 0.385 | -0.35 | 7.3E-01 | 0 |
| ::::X-11441 | 1 | 680218 | 680218 | 680218 | 0.319 | -0.34 | 7.3E-01 | 0 |
| Total lipids in small HDL | 6 | 390464 | 1107645 | 109108 | 0.141 | -0.34 | 7.3E-01 | 0 |
| Superior longitudinal fasciculus fractional anisotropy | 4 | 498527 | 751069 | 233972 | 0.230 | -0.34 | 7.3E-01 | 0 |
| ::::X-11442 | 1 | 680218 | 680218 | 680218 | 0.319 | -0.34 | 7.3E-01 | 0 |
| Types of physical activity in last 4 weeks: Light DIY (eg: pruning, watering the lawn) | 5 | 564225 | 842960 | 76649 | 0.268 | -0.34 | 7.4E-01 | 0 |
| Coronary artery disease and total cholesterol (bivariate) | 135 | 730695 | 1120903 | 376863 | 0.275 | -0.33 | 7.4E-01 | 0 |
| Sum basophil neutrophil count (three-way meta) | 98 | 718523 | 1191007 | 107013 | 0.289 | -0.33 | 7.4E-01 | 0 |
| Estimated bone mineral density from heel ultrasounds | 517 | 736918 | 1128618 | 201271 | 0.272 | -0.33 | 7.4E-01 | 0 |
| Worry subcluster | 59 | 731810 | 1311698 | 112775 | 0.268 | -0.33 | 7.5E-01 | 0 |
| Neutrophil percentage of white cells (two-way meta) | 78 | 663046 | 1121193 | 98840 | 0.260 | -0.32 | 7.5E-01 | 0 |
| Diagnoses - secondary ICD10: E11 Type 2 diabetes mellitus | 38 | 754996 | 1094611 | 356374 | 0.301 | -0.32 | 7.5E-01 | 0 |
| Cholesterol esters in very large HDL | 7 | 576970 | 924116 | 300392 | 0.248 | -0.32 | 7.5E-01 | 0 |
| Coronary artery disease and low-density lipoprotein cholesterol (bivariate) | 133 | 730695 | 1139080 | 361015 | 0.287 | -0.31 | 7.5E-01 | 0 |
| Anterior limb of internal capsule mean diusivities | 3 | 466160 | 780749 | 249843 | 0.125 | -0.31 | 7.6E-01 | 0 |
| ApoA1 | 6 | 606613 | 1066224 | 196449 | 0.287 | -0.31 | 7.6E-01 | 0 |
| ApoB | 14 | 554768 | 970919 | 173578 | 0.176 | -0.30 | 7.6E-01 | 0 |
| External capsule axial diusivities | 1 | 466160 | 466160 | 466160 | 0.125 | -0.30 | 7.6E-01 | 0 |
| Freckles | 4 | 530375 | 666042 | 333886 | 0.193 | -0.30 | 7.6E-01 | 0 |
| Posterior thalamic radiation (include optic radiation) fractional anisotropy | 1 | 695505 | 695505 | 695505 | 0.304 | -0.30 | 7.6E-01 | 0 |
| Mouth/teeth dental problems: Dentures | 30 | 784160 | 1066246 | 208581 | 0.366 | -0.30 | 7.7E-01 | 0 |
| Cofactors and vitamins::Ascorbate and aldarate metabolism::X-11593--O-methylascorbate* | 5 | 694273 | 810723 | 646865 | 0.318 | -0.30 | 7.7E-01 | 0 |
| External capsule mean diusivities | 1 | 466160 | 466160 | 466160 | 0.125 | -0.30 | 7.7E-01 | 0 |
| Basophil percentage of granulocytes (three-way meta) | 38 | 669313 | 1004079 | 126006 | 0.271 | -0.30 | 7.7E-01 | 0 |
| Neuroticism general factor | 46 | 796220 | 1230458 | 99213 | 0.347 | -0.29 | 7.7E-01 | 0 |
| Total lipids in medium LDL | 12 | 435135 | 1050478 | 86744 | 0.153 | -0.28 | 7.8E-01 | 0 |
| Left inferior lateral ventricle | 4 | 766684 | 984178 | 446730 | 0.352 | -0.28 | 7.8E-01 | 0 |
| Lipid::Monoacylglycerol::1-stearoylglycerol (1-monostearin) | 1 | 712063 | 712063 | 712063 | 0.385 | -0.28 | 7.8E-01 | 0 |
| Symbol digit substitution test - Number of symbol digit matches attempted | 6 | 705734 | 1240339 | 455333 | 0.353 | -0.28 | 7.8E-01 | 0 |
| Snoring | 34 | 698791 | 1242741 | 111614 | 0.247 | -0.28 | 7.8E-01 | 0 |
| Femoral Neck BMD | 36 | 796618 | 1241681 | 419366 | 0.331 | -0.27 | 7.8E-01 | 0 |
| Subcutaneous adipose tissue volume | 1 | 712045 | 712045 | 712045 | 0.394 | -0.27 | 7.9E-01 | 0 |
| Lipid::Fatty acid, dicarboxylate::hexadecanedioate | 1 | 482085 | 482085 | 482085 | 0.143 | -0.27 | 7.9E-01 | 0 |
| Lipid::Lysolipid::1-eicosadienoylglycerophosphocholine* | 1 | 717513 | 717513 | 717513 | 0.349 | -0.27 | 7.9E-01 | 0 |
| Type 2 Diabetes | 409 | 767475 | 1126930 | 373483 | 0.292 | -0.27 | 7.9E-01 | 0 |
| ::::X-11538 | 1 | 482085 | 482085 | 482085 | 0.143 | -0.27 | 7.9E-01 | 0 |
| Neuroticism (univariate) | 72 | 807849 | 1108384 | 503082 | 0.319 | -0.26 | 7.9E-01 | 0 |
| Right amygdala | 6 | 709455 | 1023940 | 394565 | 0.349 | -0.26 | 8.0E-01 | 0 |
| Impedance measures - Arm predicted mass (right) | 518 | 750139 | 1164069 | 111465 | 0.294 | -0.26 | 8.0E-01 | 0 |
| Hearing aid user | 3 | 626783 | 945589 | 324752 | 0.466 | -0.26 | 8.0E-01 | 0 |
| Right entorhinal | 3 | 634860 | 931785 | 379027 | 0.235 | -0.26 | 8.0E-01 | 0 |
| Amino acid::Phenylalanine & tyrosine metabolism::3-(4-hydroxyphenyl)lactate | 1 | 786553 | 786553 | 786553 | 0.405 | -0.25 | 8.0E-01 | 0 |
| Ever had hysterectomy (womb removed) (female) | 2 | 524988 | 765437 | 284540 | 0.167 | -0.25 | 8.0E-01 | 0 |
| Adrenergics, inhalants | 40 | 751529 | 1001276 | 318350 | 0.327 | -0.25 | 8.0E-01 | 0 |
| Vascular endothelial growth factor | 2 | 816755 | 1170288 | 463223 | 0.413 | -0.25 | 8.0E-01 | 0 |
| Cingulum (cingulate gyrus) radial diusivities | 2 | 589824 | 732738 | 446909 | 0.227 | -0.25 | 8.0E-01 | 0 |
| Falls in the last year | 5 | 783183 | 893328 | 238323 | 0.385 | -0.24 | 8.1E-01 | 0 |

| | | | | | | | | |
|---|---|---|---|---|---|---|---|---|
| Age of initiation of regular smoking | 5 | 662633 | 1188550 | 465385 | 0.312 | -0.24 | 8.1E-01 | 0 |
| Gamma-glutamyl transferase | 30 | 665893 | 1161485 | 293271 | 0.277 | -0.23 | 8.2E-01 | 0 |
| Depression - Recent changes in speed/amount of moving or speaking | 2 | 411778 | 589381 | 234174 | 0.054 | -0.23 | 8.2E-01 | 0 |
| Mean corpuscular hemoglobin (three-way meta) | 143 | 738943 | 1120288 | 306305 | 0.273 | -0.23 | 8.2E-01 | 0 |
| Age at menopause (last menstrual period) (female) | 59 | 741363 | 1167133 | 120117 | 0.314 | -0.22 | 8.2E-01 | 0 |
| Right inferior lateral ventricle | 3 | 610818 | 819869 | 320824 | 0.332 | -0.22 | 8.2E-01 | 0 |
| Creatine kinase | 28 | 786454 | 1021162 | 84840 | 0.347 | -0.22 | 8.3E-01 | 0 |
| Osteoarthritis of hip | 28 | 739114 | 1045994 | 187140 | 0.351 | -0.22 | 8.3E-01 | 0 |
| Duration of light DIY | 2 | 560493 | 578058 | 542928 | 0.270 | -0.22 | 8.3E-01 | 0 |
| Concentration of medium LDL particles | 13 | 485140 | 1055113 | 106456 | 0.188 | -0.21 | 8.3E-01 | 0 |
| Drugs for peptic ulcer and gastro-oesophageal reflux disease (GORD) | 5 | 795498 | 1178155 | 66035 | 0.424 | -0.21 | 8.3E-01 | 0 |
| 1p deletion neuroblastoma | 3 | 200598 | 241084 | 121469 | 0.016 | -0.20 | 8.4E-01 | 0 |
| Lipid::Fatty acid metabolism (also BCAA metabolism)::butyrylcarnitine | 4 | 661675 | 1076433 | 267493 | 0.242 | -0.20 | 8.4E-01 | 0 |
| Free thyroxine (FT4, female) | 9 | 757165 | 821192 | 310785 | 0.367 | -0.20 | 8.4E-01 | 0 |
| Body Mass Index | 1665 | 795433 | 1181033 | 227601 | 0.312 | -0.20 | 8.4E-01 | 0 |
| Focal Epilepsy | 1 | 649165 | 649165 | 649165 | 0.249 | -0.20 | 8.4E-01 | 0 |
| Waist circumference | 371 | 802540 | 1191915 | 308986 | 0.317 | -0.20 | 8.4E-01 | 0 |
| Waist circumference (male, adjusted for BMI) | 37 | 813175 | 1190793 | 112308 | 0.322 | -0.20 | 8.4E-01 | 0 |
| Asthma (adult-onset) | 25 | 767645 | 908280 | 347450 | 0.316 | -0.19 | 8.5E-01 | 0 |
| Right ventral DC | 6 | 679235 | 1226819 | 77010 | 0.313 | -0.19 | 8.5E-01 | 0 |
| Anterior limb of internal capsule axial diusivities | 1 | 528340 | 528340 | 528340 | 0.125 | -0.19 | 8.5E-01 | 0 |
| Atrial fibrillation | 103 | 704448 | 1122404 | 342304 | 0.265 | -0.19 | 8.5E-01 | 0 |
| Chronotype | 168 | 788309 | 1169536 | 120038 | 0.304 | -0.18 | 8.5E-01 | 0 |
| Anxiety/tension factors | 13 | 782163 | 1343513 | 551685 | 0.349 | -0.18 | 8.6E-01 | 0 |
| Lipid::Carnitine metabolism::palmitoylcarnitine | 1 | 758448 | 758448 | 758448 | 0.373 | -0.18 | 8.6E-01 | 0 |
| Left postcentral | 6 | 617603 | 665156 | 322343 | 0.208 | -0.18 | 8.6E-01 | 0 |
| Impedance measures - Arm fat-free mass (right) | 521 | 755783 | 1190660 | 125680 | 0.294 | -0.18 | 8.6E-01 | 0 |
| CH2 groups to double bonds ratio | 4 | 705436 | 922774 | 496423 | 0.383 | -0.18 | 8.6E-01 | 0 |
| Non-cancer illness code, self-reported: osteoarthritis | 3 | 654168 | 848608 | 343179 | 0.435 | -0.18 | 8.6E-01 | 0 |
| Neutrophil count | 13 | 644433 | 889203 | 371408 | 0.350 | -0.18 | 8.6E-01 | 0 |
| Impedance measures - Leg fat mass (right) | 340 | 830692 | 1192355 | 251613 | 0.341 | -0.18 | 8.6E-01 | 0 |
| ::::X-11327 | 2 | 787613 | 1007905 | 567320 | 0.327 | -0.17 | 8.6E-01 | 0 |
| Worry/vulnerability factors | 13 | 782163 | 1343513 | 551685 | 0.349 | -0.17 | 8.6E-01 | 0 |
| Agents acting on the renin-angiotensin system | 145 | 741878 | 1136463 | 114163 | 0.291 | -0.17 | 8.6E-01 | 0 |
| ::::X-02269 | 3 | 671508 | 1023313 | 547845 | 0.282 | -0.17 | 8.6E-01 | 0 |
| Eosinophil count | 11 | 636658 | 958540 | 397513 | 0.264 | -0.17 | 8.6E-01 | 0 |
| Antidepressants | 1 | 769455 | 769455 | 769455 | 0.385 | -0.17 | 8.7E-01 | 0 |
| Worry too long after embarrassment | 22 | 767135 | 1021552 | 138923 | 0.295 | -0.17 | 8.7E-01 | 0 |
| ::::X-11452 | 1 | 822285 | 822285 | 822285 | 0.478 | -0.17 | 8.7E-01 | 0 |
| Nucleotide::Purine metabolism, adenine containing::N1-methyladenosine | 1 | 657820 | 657820 | 657820 | 0.237 | -0.17 | 8.7E-01 | 0 |
| Estimated glomerular filtration rate | 451 | 780430 | 1148385 | 267844 | 0.304 | -0.16 | 8.7E-01 | 0 |
| Diagnoses - main ICD10: K40 Inguinal hernia | 14 | 820809 | 1125520 | 73491 | 0.355 | -0.16 | 8.7E-01 | 0 |
| Lipid::Carnitine metabolism::oleoylcarnitine | 1 | 758448 | 758448 | 758448 | 0.373 | -0.16 | 8.7E-01 | 0 |
| Major depressive disorder (females) | 2 | 306760 | 420480 | 193040 | 0.023 | -0.16 | 8.7E-01 | 0 |
| Long sleep | 4 | 611474 | 953296 | 301139 | 0.173 | -0.16 | 8.7E-01 | 0 |
| Left pars triangularis | 1 | 669333 | 669333 | 669333 | 0.221 | -0.15 | 8.8E-01 | 0 |
| Neuroticism | 152 | 798790 | 1187466 | 226861 | 0.308 | -0.15 | 8.8E-01 | 0 |
| Illnesses of siblings: Heart disease | 3 | 851270 | 1019795 | 775553 | 0.406 | -0.15 | 8.8E-01 | 0 |
| Splenium of corpus callosum radial diusivities | 7 | 797153 | 1372359 | 717880 | 0.304 | -0.14 | 8.9E-01 | 0 |
| Concentration of chylomicrons and extremely large VLDL particles | 7 | 614910 | 976125 | 340115 | 0.129 | -0.14 | 8.9E-01 | 0 |
| Height (female) | 52 | 807424 | 1156217 | 151947 | 0.322 | -0.14 | 8.9E-01 | 0 |
| Subcutaneous adipose tissue attenuation (male) | 1 | 544410 | 544410 | 544410 | 0.103 | -0.14 | 8.9E-01 | 0 |
| Non-cancer illness code, self-reported: hypothyroidism/myxoedema | 75 | 783140 | 1139713 | 327709 | 0.332 | -0.13 | 8.9E-01 | 0 |
| Cingulum (hippocampus) fractional anisotropy | 1 | 840440 | 840440 | 840440 | 0.495 | -0.13 | 8.9E-01 | 0 |
| Impedance measures - Whole body fat-free mass | 621 | 762725 | 1191883 | 134846 | 0.298 | -0.13 | 9.0E-01 | 0 |
| Inferior fronto-occipital fasciculus fractional anisotropy | 4 | 773926 | 961300 | 624258 | 0.304 | -0.13 | 9.0E-01 | 0 |

| | | | | | | | | |
|---|---|---|---|---|---|---|---|---|
| Basophil percentage of white cells (two-way meta) | 29 | 676158 | 1020317 | 126312 | 0.271 | -0.13 | 9.0E-01 | 0 |
| Lymphocyte count (three-way meta) | 114 | 749410 | 1111608 | 171860 | 0.267 | -0.13 | 9.0E-01 | 0 |
| Frequency of friend/family visits | 10 | 677620 | 1352924 | 135529 | 0.249 | -0.13 | 9.0E-01 | 0 |
| Hemoglobin A1c | 21 | 705053 | 1112253 | 417280 | 0.275 | -0.13 | 9.0E-01 | 0 |
| Phospholipids in large LDL | 11 | 521590 | 1009733 | 240700 | 0.188 | -0.12 | 9.0E-01 | 0 |
| Glutamine | 6 | 678170 | 1395566 | 58720 | 0.220 | -0.12 | 9.0E-01 | 0 |
| Type 2 Diabetes (adjusted for BMI) | 153 | 763233 | 1077243 | 353550 | 0.290 | -0.12 | 9.0E-01 | 0 |
| Inferior fronto-occipital fasciculus mean diusivities | 2 | 573296 | 633178 | 513414 | 0.159 | -0.12 | 9.1E-01 | 0 |
| CH2 groups in fatty acids | 3 | 385130 | 439593 | 202385 | 0.079 | -0.12 | 9.1E-01 | 0 |
| Glucose levels 2 h after an oral glucose challenge | 1 | 683278 | 683278 | 683278 | 0.284 | -0.12 | 9.1E-01 | 0 |
| Estimated glomerular filtration rate based on serum creatinine | 96 | 776530 | 1103014 | 356768 | 0.296 | -0.11 | 9.1E-01 | 0 |
| Monocyte count | 25 | 767305 | 1188618 | 309813 | 0.351 | -0.11 | 9.1E-01 | 0 |
| Lipid::Long chain fatty acid::pentadecanoate (15:0) | 1 | 694215 | 694215 | 694215 | 0.209 | -0.11 | 9.1E-01 | 0 |
| Blood sugar | 14 | 582326 | 908757 | 154860 | 0.219 | -0.11 | 9.1E-01 | 0 |
| ::::X-11412 | 1 | 797550 | 797550 | 797550 | 0.324 | -0.11 | 9.2E-01 | 0 |
| Eosinophil percentage of white cells (two-way meta) | 94 | 741745 | 1221979 | 122375 | 0.259 | -0.11 | 9.2E-01 | 0 |
| Forearm BMD | 3 | 760258 | 1174644 | 578291 | 0.286 | -0.10 | 9.2E-01 | 0 |
| Albuminuria | 31 | 714893 | 1137285 | 249761 | 0.311 | -0.10 | 9.2E-01 | 0 |
| Amino acid::Valine, leucine and isoleucine metabolism::leucine | 11 | 913785 | 1076463 | 599483 | 0.451 | -0.10 | 9.2E-01 | 0 |
| Cancer (diagnosed by doctor) | 5 | 729058 | 1116113 | 718645 | 0.279 | -0.10 | 9.2E-01 | 0 |
| Treatment/medication code: simvastatin | 20 | 683355 | 873274 | 215978 | 0.304 | -0.10 | 9.2E-01 | 0 |
| Genu of corpus callosum axial diusivities | 3 | 707440 | 1132926 | 606061 | 0.253 | -0.10 | 9.2E-01 | 0 |
| High light scatter percentage of red cells (three-way meta) | 118 | 707310 | 1110043 | 119754 | 0.276 | -0.09 | 9.2E-01 | 0 |
| Uncinate fasciculus fractional anisotropy | 2 | 790013 | 994476 | 585549 | 0.354 | -0.09 | 9.3E-01 | 0 |
| Amino acid::Glycine, serine and threonine metabolism::glycine | 1 | 804745 | 804745 | 804745 | 0.312 | -0.09 | 9.3E-01 | 0 |
| Antihypertensives | 2 | 818203 | 848533 | 787873 | 0.322 | -0.09 | 9.3E-01 | 0 |
| Transport type for commuting to job workplace: Car/motor vehicle | 1 | 864035 | 864035 | 864035 | 0.469 | -0.09 | 9.3E-01 | 0 |
| Amino acid::Creatine metabolism::creatine | 1 | 804745 | 804745 | 804745 | 0.312 | -0.08 | 9.3E-01 | 0 |
| Cholesterol esters in medium LDL | 14 | 544955 | 1053568 | 173578 | 0.196 | -0.08 | 9.3E-01 | 0 |
| Impedance measures - Impedance of arm (left) | 426 | 783736 | 1189558 | 155748 | 0.305 | -0.08 | 9.3E-01 | 0 |
| Total cholesterol in LDL | 12 | 494950 | 950535 | 86744 | 0.124 | -0.08 | 9.4E-01 | 0 |
| Amino acid::Valine, leucine and isoleucine metabolism::isobutyrylcarnitine | 2 | 731771 | 992866 | 470677 | 0.313 | -0.08 | 9.4E-01 | 0 |
| HbA1c | 52 | 767241 | 1234628 | 451091 | 0.266 | -0.08 | 9.4E-01 | 0 |
| Total cholesterol in large LDL | 12 | 563180 | 1090583 | 307823 | 0.202 | -0.07 | 9.4E-01 | 0 |
| Total cholesterol in HDL | 11 | 586548 | 1042579 | 74752 | 0.221 | -0.07 | 9.4E-01 | 0 |
| Medication for cholesterol, blood pressure or diabetes: Blood pressure medication | 50 | 770760 | 1296984 | 288661 | 0.311 | -0.07 | 9.5E-01 | 0 |
| Hip circumference (female, adjusted for BMI) | 54 | 807940 | 1223229 | 189748 | 0.300 | -0.07 | 9.5E-01 | 0 |
| Frequency of depressed mood in last 2 weeks | 5 | 798015 | 810813 | 555690 | 0.314 | -0.06 | 9.5E-01 | 0 |
| LDL diameter | 5 | 805883 | 955163 | 395798 | 0.313 | -0.06 | 9.5E-01 | 0 |
| Body Mass Index (female) | 451 | 814438 | 1173691 | 369250 | 0.323 | -0.06 | 9.5E-01 | 0 |
| Mean Arterial Pressure | 11 | 600510 | 908715 | 156635 | 0.172 | -0.06 | 9.6E-01 | 0 |
| Lipid::Sterol, Steroid::cortisone | 1 | 870475 | 870475 | 870475 | 0.413 | -0.06 | 9.6E-01 | 0 |
| Triglycerides in very large HDL | 10 | 823634 | 1163668 | 650053 | 0.342 | -0.05 | 9.6E-01 | 0 |
| Concentration of very large HDL particles | 7 | 797550 | 1177365 | 584506 | 0.324 | -0.05 | 9.6E-01 | 0 |
| Hip circumference (male) | 24 | 833950 | 1196401 | 458584 | 0.367 | -0.05 | 9.6E-01 | 0 |
| Age at first live birth (female) | 14 | 673129 | 1125947 | 85269 | 0.292 | -0.05 | 9.6E-01 | 0 |
| Number of live births (female) | 7 | 734533 | 1061604 | 98554 | 0.267 | -0.05 | 9.6E-01 | 0 |
| Heart rate recovery at 20 secnds | 14 | 678653 | 1054133 | 157283 | 0.190 | -0.05 | 9.6E-01 | 0 |
| Hair colour (natural, before greying): Dark brown | 174 | 536363 | 994047 | 81878 | 0.156 | -0.05 | 9.6E-01 | 0 |
| Basophil count | 20 | 650554 | 1232640 | 265745 | 0.212 | -0.04 | 9.7E-01 | 0 |
| Superior corona radiata mean diusivities | 2 | 835703 | 905801 | 765604 | 0.345 | -0.04 | 9.7E-01 | 0 |
| Fasting glucose main effect | 16 | 762186 | 1391916 | 191450 | 0.292 | -0.04 | 9.7E-01 | 0 |
| Amino acid::Tryptophan metabolism::indolelactate | 1 | 723603 | 723603 | 723603 | 0.278 | -0.03 | 9.7E-01 | 0 |
| Bust size | 2 | 721569 | 763804 | 679333 | 0.243 | -0.03 | 9.7E-01 | 0 |
| Right precuneus | 2 | 748906 | 1094522 | 403291 | 0.305 | -0.03 | 9.8E-01 | 0 |
| Lipid::Sphingolipid::palmitoyl sphingomyelin | 1 | 737510 | 737510 | 737510 | 0.296 | -0.02 | 9.8E-01 | 0 |

| | | | | | | | | |
|---|---|---|---|---|---|---|---|---|
| Left caudal anterior cingulate | 3 | 691988 | 868646 | 358413 | 0.342 | -0.02 | 9.8E-01 | 0 |
| Epithelial ovarian cancer (high -grade and low-grade serous) | 1 | 349565 | 349565 | 349565 | 0.010 | -0.02 | 9.9E-01 | 0 |
| Number of days/week of vigorous physical activity 10+ minutes | 10 | 824791 | 1006328 | 634146 | 0.328 | -0.02 | 9.9E-01 | 0 |
| Hip circumference (adjusted for BMI) | 102 | 810285 | 1126301 | 248288 | 0.292 | -0.02 | 9.9E-01 | 0 |
| Hearing difficulty/problems with background noise | 25 | 830990 | 1001603 | 84269 | 0.363 | -0.02 | 9.9E-01 | 0 |
| Non-albumin protein | 36 | 708306 | 1201087 | 464978 | 0.256 | -0.02 | 9.9E-01 | 0 |
| Splenium of corpus callosum fractional anisotropy | 5 | 740255 | 970290 | 695505 | 0.304 | -0.01 | 9.9E-01 | 0 |
| Lipid::Carnitine metabolism::octanoylcarnitine | 3 | 852333 | 1008056 | 695191 | 0.307 | -0.01 | 9.9E-01 | 0 |
| Lipid::Sterol, Steroid::epiandrosterone sulfate | 1 | 355103 | 355103 | 355103 | 0.088 | -0.01 | 9.9E-01 | 0 |
| Retrolenticular part of internal capsule fractional anisotropy | 4 | 699931 | 1028419 | 362049 | 0.289 | -0.01 | 9.9E-01 | 0 |
| Types of transport used (excluding work): Cycle | 1 | 746725 | 746725 | 746725 | 0.238 | -0.01 | 9.9E-01 | 0 |
| Epithelial ovarian cancer (high-grade serous) | 1 | 349565 | 349565 | 349565 | 0.010 | 0.00 | 1.0E+00 | 0 |
| Diagnoses - main ICD10: K57 Diverticular disease of intestine | 12 | 732865 | 1004836 | 394734 | 0.214 | 0.00 | 1.0E+00 | 0 |
| Overall activity (conditioning sex and BMI) | 2 | 596295 | 876853 | 315738 | 0.224 | 0.00 | 1.0E+00 | 0 |
| Body of corpus callosum fractional anisotropy | 3 | 819578 | 883061 | 561786 | 0.392 | 0.00 | 1.0E+00 | 0 |
| Non-cancer illness code, self-reported: migraine | 13 | 800740 | 1061773 | 406448 | 0.316 | 0.00 | 1.0E+00 | 0 |
| Superior longitudinal fasciculus mean diusivities | 3 | 729058 | 1076166 | 516526 | 0.265 | 0.01 | 1.0E+00 | 0 |
| Sleep sedentary (conditioning sex and BMI) | 5 | 759793 | 1561180 | 701760 | 0.348 | 0.01 | 1.0E+00 | 0 |
| Superior corona radiata fractional anisotropy | 4 | 697195 | 760934 | 523477 | 0.286 | 0.01 | 9.9E-01 | 0 |
| Genetic generalized epilepsy | 1 | 903535 | 903535 | 903535 | 0.491 | 0.01 | 9.9E-01 | 0 |
| Automobile speeding propensity | 31 | 813355 | 1119875 | 538151 | 0.359 | 0.02 | 9.9E-01 | 0 |
| Retrolenticular part of internal capsule mode of anisotropy | 6 | 631640 | 953067 | 325622 | 0.252 | 0.02 | 9.9E-01 | 0 |
| Life satisfaction (MA GWAMA) | 32 | 779481 | 1198157 | 381741 | 0.265 | 0.02 | 9.9E-01 | 0 |
| Amino acid::Lysine metabolism::glutaroyl carnitine | 8 | 726981 | 1094302 | 443685 | 0.281 | 0.02 | 9.9E-01 | 0 |
| Retrolenticular part of internal capsule mean diusivities | 2 | 801855 | 855030 | 748680 | 0.292 | 0.02 | 9.8E-01 | 0 |
| Relative wall thickness | 1 | 853988 | 853988 | 853988 | 0.312 | 0.02 | 9.8E-01 | 0 |
| Nervous feelings (NERV-FEEL) | 35 | 835355 | 1175465 | 235547 | 0.316 | 0.03 | 9.8E-01 | 0 |
| Cholesterol esters in large LDL | 13 | 604770 | 1055113 | 374945 | 0.204 | 0.03 | 9.8E-01 | 0 |
| Triglycerides in medium VLDL | 8 | 628013 | 1152693 | 85903 | 0.175 | 0.03 | 9.7E-01 | 0 |
| Vitiligo (early onset) | 1 | 922435 | 922435 | 922435 | 0.449 | 0.03 | 9.7E-01 | 0 |
| Social support - Leisure/social activities: Pub or social club | 17 | 818465 | 1284278 | 506602 | 0.332 | 0.03 | 9.7E-01 | 0 |
| Lymphocyte count | 10 | 698776 | 1347217 | 201352 | 0.254 | 0.03 | 9.7E-01 | 0 |
| Thyroid-stimulating hormone (female) | 21 | 801673 | 1080473 | 576603 | 0.315 | 0.04 | 9.7E-01 | 0 |
| Impedance measures - Impedance of arm (right) | 420 | 784519 | 1167779 | 151590 | 0.302 | 0.04 | 9.7E-01 | 0 |
| Neuroticism score | 72 | 823763 | 1246051 | 393650 | 0.303 | 0.04 | 9.7E-01 | 0 |
| Genu of corpus callosum radial diusivities | 4 | 807954 | 1008323 | 699594 | 0.338 | 0.04 | 9.7E-01 | 0 |
| Energy::Krebs cycle::malate | 1 | 938865 | 938865 | 938865 | 0.413 | 0.05 | 9.6E-01 | 0 |
| Neuroticism sum score | 92 | 812364 | 1268068 | 187236 | 0.304 | 0.05 | 9.6E-01 | 0 |
| Loneliness (MTAG) | 13 | 802985 | 1398988 | 70786 | 0.330 | 0.05 | 9.6E-01 | 0 |
| Total lipids in medium HDL | 3 | 586548 | 984386 | 328255 | 0.172 | 0.05 | 9.6E-01 | 0 |
| High light scatter percentage of red cells (two-way meta) | 102 | 759251 | 1129961 | 177884 | 0.290 | 0.05 | 9.6E-01 | 0 |
| Impedance measures - Arm predicted mass (left) | 538 | 763551 | 1184503 | 142700 | 0.293 | 0.06 | 9.6E-01 | 0 |
| FEV1/FVC ratio | 478 | 757640 | 1155103 | 162547 | 0.277 | 0.06 | 9.5E-01 | 0 |
| Concentration of medium HDL particles | 3 | 586548 | 984386 | 328255 | 0.172 | 0.06 | 9.5E-01 | 0 |
| Lipid::Fatty acid metabolism (also BCAA metabolism)::propionylcarnitine | 4 | 758270 | 1254596 | 388469 | 0.287 | 0.06 | 9.5E-01 | 0 |
| Illnesses of father: Prostate cancer | 3 | 842773 | 1055566 | 566590 | 0.457 | 0.07 | 9.5E-01 | 0 |
| Basophil count (three-way meta) | 53 | 766478 | 1112530 | 96751 | 0.290 | 0.07 | 9.5E-01 | 0 |
| Heel bone mineral density | 305 | 736918 | 1117040 | 141637 | 0.271 | 0.07 | 9.5E-01 | 0 |
| Atrial Fibrillation | 103 | 740255 | 1119101 | 348511 | 0.272 | 0.07 | 9.4E-01 | 0 |
| Right hippocampus | 12 | 737163 | 905236 | 433041 | 0.290 | 0.07 | 9.4E-01 | 0 |
| Drugs affecting bone structure and mineralization | 10 | 739629 | 1139094 | 433414 | 0.257 | 0.07 | 9.4E-01 | 0 |
| Diagnoses - secondary ICD10: K21 Gastro-esophageal reflux disease | 1 | 940400 | 940400 | 940400 | 0.448 | 0.08 | 9.4E-01 | 0 |
| Eosinophil percentage of white cells (three-way meta) | 116 | 737688 | 1148856 | 122100 | 0.268 | 0.08 | 9.4E-01 | 0 |
| Age high blood pressure diagnosed | 9 | 729583 | 1372948 | 84496 | 0.289 | 0.08 | 9.4E-01 | 0 |
| Basophil percentage of granulocytes (two-way meta) | 28 | 703709 | 1128588 | 109064 | 0.277 | 0.08 | 9.4E-01 | 0 |
| Short sleep | 21 | 805503 | 1078588 | 91238 | 0.306 | 0.08 | 9.4E-01 | 0 |

| | | | | | | | | |
|---|---|---|---|---|---|---|---|---|
| Total cholesterol in medium LDL | 11 | 514640 | 831980 | 196658 | 0.118 | 0.08 | 9.4E-01 | 0 |
| Phenylalanine | 2 | 854445 | 913310 | 795580 | 0.379 | 0.09 | 9.3E-01 | 0 |
| Neutrophil percentage of granulocytes (three-way meta) | 104 | 737400 | 1165363 | 126182 | 0.268 | 0.09 | 9.3E-01 | 0 |
| Male-specific factors - Hair/balding pattern: Pattern 1 | 166 | 764678 | 1093732 | 345728 | 0.298 | 0.09 | 9.3E-01 | 0 |
| Ever had prostate specific antigen (PSA) test | 3 | 895335 | 907265 | 760590 | 0.420 | 0.09 | 9.3E-01 | 0 |
| Open-angle glaucoma (fixed-effect model) | 41 | 806553 | 1059860 | 174462 | 0.335 | 0.09 | 9.3E-01 | 0 |
| Cholesterol esters in medium HDL | 6 | 702551 | 871428 | 199109 | 0.161 | 0.09 | 9.3E-01 | 0 |
| Cannabis use | 4 | 787915 | 1245809 | 353908 | 0.297 | 0.09 | 9.3E-01 | 0 |
| Extreme Waist-hip ratio (adjusted for BMI) | 2 | 880005 | 965983 | 794028 | 0.371 | 0.09 | 9.2E-01 | 0 |
| Thyroid-stimulating hormone | 39 | 778293 | 929274 | 498726 | 0.287 | 0.09 | 9.2E-01 | 0 |
| Left ventricular internal dimension in diastole | 3 | 853988 | 1185196 | 800129 | 0.312 | 0.10 | 9.2E-01 | 0 |
| Sagittal stratum mode of anisotropy | 2 | 658305 | 969691 | 346919 | 0.259 | 0.10 | 9.2E-01 | 0 |
| Treatment/medication code: aspirin | 4 | 642720 | 938733 | 349516 | 0.172 | 0.10 | 9.2E-01 | 0 |
| Right supramarginal | 3 | 300553 | 495426 | 162765 | 0.081 | 0.10 | 9.2E-01 | 0 |
| Pericardial adipose tissue volume (male) | 1 | 958742 | 958742 | 958742 | 0.473 | 0.10 | 9.2E-01 | 0 |
| Pericardial adipose tissue volume (adjusted for height and weight, male) | 1 | 958742 | 958742 | 958742 | 0.473 | 0.10 | 9.2E-01 | 0 |
| Probable major dpressive disorder | 1 | 966988 | 966988 | 966988 | 0.411 | 0.10 | 9.2E-01 | 0 |
| Adiponectin | 6 | 499571 | 1133022 | 437864 | 0.067 | 0.11 | 9.2E-01 | 0 |
| Diagnoses - main ICD10: C44 Other and unspecified malignant neoplasm of skin | 26 | 682456 | 967618 | 200524 | 0.196 | 0.11 | 9.2E-01 | 0 |
| Lymphocyte percentage of white cells (two-way meta) | 83 | 768645 | 1144873 | 196233 | 0.258 | 0.11 | 9.1E-01 | 0 |
| Geographic atrophy | 3 | 678693 | 758206 | 552133 | 0.186 | 0.11 | 9.1E-01 | 0 |
| Cooked vegetable intake | 12 | 719039 | 981728 | 141457 | 0.275 | 0.12 | 9.1E-01 | 0 |
| Anxiety disorder (factor score) | 1 | 903160 | 903160 | 903160 | 0.316 | 0.12 | 9.1E-01 | 0 |
| Illnesses of mother: Breast cancer | 12 | 713113 | 1123981 | 72306 | 0.239 | 0.12 | 9.0E-01 | 0 |
| Non-cancer illness code, self-reported: depression | 1 | 803498 | 803498 | 803498 | 0.284 | 0.12 | 9.0E-01 | 0 |
| Immature fraction of reticulocytes (three-way meta) | 101 | 769200 | 1123655 | 238845 | 0.314 | 0.12 | 9.0E-01 | 0 |
| Stem cell factor | 2 | 896879 | 1077596 | 716162 | 0.323 | 0.13 | 9.0E-01 | 0 |
| Vitamin and mineral supplements: Multivitamins +/- minerals | 2 | 924765 | 1319770 | 529761 | 0.401 | 0.13 | 9.0E-01 | 0 |
| Heart rate variability (RMSSD) | 7 | 767923 | 1475635 | 326952 | 0.348 | 0.13 | 8.9E-01 | 0 |
| Milk type used: Soya | 4 | 736263 | 1508556 | 52620 | 0.306 | 0.13 | 8.9E-01 | 0 |
| ::::X-11799 | 1 | 691853 | 691853 | 691853 | 0.139 | 0.13 | 8.9E-01 | 0 |
| Phospholipids in medium VLDL | 10 | 666598 | 1093759 | 253186 | 0.170 | 0.13 | 8.9E-01 | 0 |
| Platelet derived growth factor BB | 3 | 836653 | 1103279 | 483783 | 0.272 | 0.14 | 8.9E-01 | 0 |
| Estimated BMD | 189 | 739313 | 1141198 | 209031 | 0.257 | 0.14 | 8.9E-01 | 0 |
| Extreme obesity (childhood) | 2 | 876113 | 939594 | 812631 | 0.332 | 0.14 | 8.9E-01 | 0 |
| Lipid::Sterol, Steroid::4-androsten-3beta,17beta-diol disulfate 2* | 2 | 802881 | 1179008 | 426753 | 0.290 | 0.14 | 8.9E-01 | 0 |
| Asthma (random effect model) | 18 | 825820 | 1147311 | 395958 | 0.327 | 0.14 | 8.9E-01 | 0 |
| Hip circumference | 435 | 810993 | 1208263 | 201310 | 0.306 | 0.15 | 8.8E-01 | 0 |
| Nervous feelings | 30 | 868378 | 1199436 | 373306 | 0.307 | 0.15 | 8.8E-01 | 0 |
| Cingulum (hippocampus) mean diusivities | 1 | 925098 | 925098 | 925098 | 0.316 | 0.15 | 8.8E-01 | 0 |
| 25-Hydroxyvitamin D level | 15 | 606380 | 922575 | 200297 | 0.189 | 0.15 | 8.8E-01 | 0 |
| Uncinate fasciculus axial diusivities | 1 | 693060 | 693060 | 693060 | 0.196 | 0.15 | 8.8E-01 | 0 |
| Waist-hip ratio (female) | 293 | 806353 | 1157218 | 379398 | 0.305 | 0.15 | 8.8E-01 | 0 |
| How are people in household related to participant: Son and/or daughter (include step-children) | 1 | 419693 | 419693 | 419693 | 0.026 | 0.15 | 8.8E-01 | 0 |
| Open-angle glaucoma (random-effect model) | 37 | 806553 | 1044898 | 168601 | 0.335 | 0.16 | 8.7E-01 | 0 |
| Inferior fronto-occipital fasciculus radial diusivities | 1 | 693060 | 693060 | 693060 | 0.196 | 0.16 | 8.7E-01 | 0 |
| HOMA-B | 4 | 826464 | 1353527 | 275448 | 0.287 | 0.16 | 8.7E-01 | 0 |
| Age at menarche | 251 | 796713 | 1153089 | 319598 | 0.291 | 0.16 | 8.7E-01 | 0 |
| Age spot | 2 | 604110 | 693733 | 514487 | 0.171 | 0.16 | 8.7E-01 | 0 |
| Right pars triangularis | 5 | 669333 | 677188 | 59909 | 0.207 | 0.17 | 8.7E-01 | 0 |
| Irritability (IRR) | 35 | 844035 | 1222873 | 187028 | 0.293 | 0.17 | 8.7E-01 | 0 |
| Male pattern baldness (BOLT LMM infinitesimal mixed model) | 326 | 757014 | 1093209 | 242427 | 0.281 | 0.17 | 8.7E-01 | 0 |
| Total lipids in medium VLDL | 8 | 651848 | 860042 | 156479 | 0.162 | 0.17 | 8.6E-01 | 0 |
| Right lateral orbitofrontal | 5 | 735818 | 1284313 | 695640 | 0.485 | 0.17 | 8.6E-01 | 0 |
| Activated partial thromboplastin time | 5 | 733210 | 972175 | 474002 | 0.263 | 0.17 | 8.6E-01 | 0 |
| Amino acid::Valine, leucine and isoleucine metabolism::hydroxyisovaleroyl carnitine | 2 | 809280 | 1142384 | 476177 | 0.272 | 0.18 | 8.6E-01 | 0 |

| | | | | | | | | |
|---|---|---|---|---|---|---|---|---|
| Visceral sdipose tissue volume (adjusted for BMI) | 4 | 799906 | 1192394 | 394419 | 0.364 | 0.18 | 8.6E-01 | 0 |
| Cancer register - Behaviour of cancer tumour: Carcinoma in situ | 1 | 840790 | 840790 | 840790 | 0.227 | 0.18 | 8.6E-01 | 0 |
| Anterior corona radiata fractional anisotropy | 6 | 756319 | 833653 | 219840 | 0.273 | 0.19 | 8.5E-01 | 0 |
| Salicylic acid and derivatives | 6 | 658126 | 995141 | 413272 | 0.156 | 0.19 | 8.5E-01 | 0 |
| Frequency of tenseness / restlessness in last 2 weeks | 8 | 739929 | 1016108 | 310178 | 0.412 | 0.19 | 8.5E-01 | 0 |
| Uncinate fasciculus radial diusivities | 1 | 703043 | 703043 | 703043 | 0.187 | 0.19 | 8.5E-01 | 0 |
| Heart rate variability (SDNN) | 6 | 751738 | 1298050 | 161926 | 0.248 | 0.19 | 8.5E-01 | 0 |
| Left supramarginal | 5 | 669333 | 690300 | 66903 | 0.221 | 0.19 | 8.5E-01 | 0 |
| Cancer register - Histology of cancer tumour: Basal cell carcinoma, NOS | 24 | 644454 | 930931 | 98940 | 0.194 | 0.19 | 8.5E-01 | 0 |
| Concentration of small LDL particles | 8 | 589284 | 877694 | 382584 | 0.124 | 0.19 | 8.5E-01 | 0 |
| Hyperthyroidism | 10 | 740721 | 1123445 | 424717 | 0.273 | 0.19 | 8.5E-01 | 0 |
| Vascular/heart problems diagnosed by doctor: High blood pressure | 189 | 776138 | 1124403 | 174418 | 0.302 | 0.20 | 8.4E-01 | 0 |
| Body fat percentage | 9 | 878838 | 1353228 | 382215 | 0.342 | 0.20 | 8.4E-01 | 0 |
| Cingulum (cingulate gyrus) axial diusivities | 4 | 557716 | 710658 | 368659 | 0.120 | 0.20 | 8.4E-01 | 0 |
| Guilty feelings (GUILT) | 12 | 854145 | 965692 | 378950 | 0.280 | 0.20 | 8.4E-01 | 0 |
| ::::X-08988 | 3 | 972713 | 992768 | 888729 | 0.460 | 0.20 | 8.4E-01 | 0 |
| Anti-inflammatory and antirheumatic products, non-steroids | 6 | 903869 | 1095183 | 247398 | 0.350 | 0.20 | 8.4E-01 | 0 |
| Right precentral | 5 | 526250 | 875192 | 29254 | 0.169 | 0.20 | 8.4E-01 | 0 |
| Concentration of small VLDL particles | 12 | 702551 | 949047 | 421839 | 0.140 | 0.21 | 8.4E-01 | 0 |
| Trunk-trunk fat ratio (male) | 15 | 822398 | 1220179 | 168480 | 0.320 | 0.21 | 8.3E-01 | 0 |
| Worry too long after embarrassment (WORR-EMB) | 18 | 861511 | 1050194 | 352567 | 0.346 | 0.21 | 8.3E-01 | 0 |
| Mean corpuscular volume (three-way meta) | 160 | 758263 | 1148619 | 188483 | 0.266 | 0.22 | 8.3E-01 | 0 |
| Central corneal thickness | 29 | 759835 | 1053798 | 422213 | 0.265 | 0.22 | 8.3E-01 | 0 |
| ::::X-13859 | 2 | 956335 | 1410120 | 502550 | 0.414 | 0.22 | 8.3E-01 | 0 |
| ::::X-03094 | 5 | 818555 | 1317188 | 83239 | 0.274 | 0.22 | 8.3E-01 | 0 |
| Number of self-reported non-cancer illnesses | 36 | 817938 | 1414458 | 363062 | 0.349 | 0.22 | 8.3E-01 | 0 |
| Family relationship satisfaction | 1 | 735878 | 735878 | 735878 | 0.190 | 0.22 | 8.2E-01 | 0 |
| Left ventricular internal dimension in systole | 3 | 853988 | 970369 | 800129 | 0.312 | 0.23 | 8.2E-01 | 0 |
| Posterior corona radiata fractional anisotropy | 2 | 922019 | 923558 | 920479 | 0.349 | 0.23 | 8.2E-01 | 0 |
| Attention deficit hyperactivity disorder | 11 | 892855 | 1212943 | 497249 | 0.426 | 0.23 | 8.2E-01 | 0 |
| Fractured/broken bones in last 5 years | 13 | 825892 | 1057395 | 434027 | 0.279 | 0.23 | 8.2E-01 | 0 |
| Mean corpuscular volume (two-way meta) | 146 | 743766 | 1151981 | 155692 | 0.278 | 0.23 | 8.2E-01 | 0 |
| Phosphorus | 6 | 606580 | 927589 | 355484 | 0.201 | 0.23 | 8.2E-01 | 0 |
| Posterior limb of internal capsule mean diusivities | 3 | 838300 | 1088341 | 628503 | 0.261 | 0.24 | 8.1E-01 | 0 |
| Lipid::Long chain fatty acid::arachidonate (20:4n6) | 1 | 978717 | 978717 | 978717 | 0.318 | 0.25 | 8.1E-01 | 0 |
| ::::X-12094 | 1 | 967390 | 967390 | 967390 | 0.366 | 0.26 | 8.0E-01 | 0 |
| Dysmenorrhea (quality of life impact) | 1 | 977578 | 977578 | 977578 | 0.334 | 0.26 | 8.0E-01 | 0 |
| Age last used hormone-replacement therapy (HRT) (female) | 1 | 473425 | 473425 | 473425 | 0.058 | 0.26 | 8.0E-01 | 0 |
| Red blood cell count | 39 | 787993 | 1056928 | 266745 | 0.262 | 0.26 | 8.0E-01 | 0 |
| ::::X-12450 | 1 | 1043453 | 1043453 | 1043453 | 0.468 | 0.26 | 8.0E-01 | 0 |
| Nucleotide::NAD metabolism::X-12095--N1-methyl-3-pyridone-4-carboxamide | 1 | 967390 | 967390 | 967390 | 0.366 | 0.26 | 7.9E-01 | 0 |
| Insomnia | 14 | 814311 | 1289623 | 64540 | 0.335 | 0.26 | 7.9E-01 | 0 |
| Total cholesterol in small LDL | 9 | 604770 | 941313 | 385130 | 0.129 | 0.27 | 7.9E-01 | 0 |
| Anxiety - Ever worried more than most people would in similar situation | 1 | 1052310 | 1052310 | 1052310 | 0.485 | 0.27 | 7.9E-01 | 0 |
| Mineral and other dietary supplements: Calcium | 4 | 749432 | 1492680 | 40017 | 0.265 | 0.27 | 7.9E-01 | 0 |
| Medication for cholesterol, blood pressure, diabetes, or take exogenous hormones: Hormone replacement therapy | 1 | 473425 | 473425 | 473425 | 0.058 | 0.27 | 7.9E-01 | 0 |
| Superior longitudinal fasciculus radial diusivities | 4 | 818631 | 1036972 | 622792 | 0.272 | 0.27 | 7.9E-01 | 0 |
| Triglycerides in small HDL | 9 | 818555 | 1118528 | 765925 | 0.221 | 0.28 | 7.8E-01 | 0 |
| Waist-hip ratio (female, adjusted for BMI) | 42 | 884941 | 1215864 | 470843 | 0.363 | 0.28 | 7.8E-01 | 0 |
| Beta blocking agents | 47 | 795763 | 1235108 | 491945 | 0.335 | 0.28 | 7.8E-01 | 0 |
| Waist-hip ratio (male, adjusted for BMI) | 9 | 937695 | 1437905 | 699138 | 0.367 | 0.28 | 7.8E-01 | 0 |
| ::::X-12844 | 4 | 824953 | 1259806 | 359058 | 0.225 | 0.28 | 7.8E-01 | 0 |
| ::::X-13435 | 2 | 948226 | 1030456 | 865997 | 0.367 | 0.28 | 7.8E-01 | 0 |
| Right inferior parietal | 6 | 973244 | 1042853 | 896782 | 0.406 | 0.28 | 7.8E-01 | 0 |
| Diagnoses - main ICD10: I25 Chronic ischemic heart disease | 25 | 755635 | 1079455 | 453870 | 0.217 | 0.28 | 7.8E-01 | 0 |
| Chloride | 11 | 801408 | 1180940 | 72265 | 0.316 | 0.29 | 7.8E-01 | 0 |

| | | | | | | | | |
|---|---|---|---|---|---|---|---|---|
| Tyrosine | 3 | 998692 | 1148823 | 528410 | 0.422 | 0.29 | 7.7E-01 | 0 |
| Lymphocyte count (two-way meta) | 110 | 785798 | 1144975 | 229143 | 0.271 | 0.29 | 7.7E-01 | 0 |
| Disc area | 27 | 673258 | 1163325 | 70832 | 0.229 | 0.29 | 7.7E-01 | 0 |
| Moderate intesity | 1 | 760235 | 760235 | 760235 | 0.174 | 0.29 | 7.7E-01 | 0 |
| Red blood cell count (three-way meta) | 129 | 811060 | 1139838 | 281423 | 0.309 | 0.29 | 7.7E-01 | 0 |
| High light scatter reticulocyte count (three-way meta) | 120 | 748144 | 1137496 | 190531 | 0.279 | 0.29 | 7.7E-01 | 0 |
| Total lipids in small LDL | 12 | 589284 | 1089553 | 296573 | 0.124 | 0.29 | 7.7E-01 | 0 |
| Types of physical activity in last 4 weeks: Heavy DIY (eg: weeding, lawn mowing, carpentry, digging) | 5 | 858305 | 1297803 | 842960 | 0.291 | 0.30 | 7.7E-01 | 0 |
| Pericardial adipose tissue volume (adjusted for height and weight, female) | 1 | 1051960 | 1051960 | 1051960 | 0.489 | 0.30 | 7.7E-01 | 0 |
| Menstrual pain medicine use | 1 | 1000075 | 1000075 | 1000075 | 0.358 | 0.30 | 7.7E-01 | 0 |
| CSF | 7 | 854105 | 914416 | 200127 | 0.307 | 0.30 | 7.7E-01 | 0 |
| Alcohol consumption (dichotomous) | 1 | 492088 | 492088 | 492088 | 0.046 | 0.30 | 7.6E-01 | 0 |
| Reason for glasses/contact lenses: For just reading/near work as you are getting older (called 'presbyopia') | 3 | 680180 | 840683 | 363239 | 0.194 | 0.30 | 7.6E-01 | 0 |
| Albumin | 9 | 671173 | 1295143 | 103940 | 0.145 | 0.31 | 7.6E-01 | 0 |
| Platelet distribution width (three-way meta) | 111 | 742700 | 1084074 | 356308 | 0.281 | 0.31 | 7.6E-01 | 0 |
| Lipid::Lysolipid::1-linoleoylglycerophosphoethanolamine* | 2 | 907378 | 916404 | 898351 | 0.289 | 0.31 | 7.6E-01 | 0 |
| Vigorous physical activity | 5 | 930240 | 1133225 | 751333 | 0.382 | 0.32 | 7.5E-01 | 0 |
| Reticulocyte count (two-way meta) | 108 | 778138 | 1094864 | 122830 | 0.294 | 0.32 | 7.5E-01 | 0 |
| Serum total cholesterol | 10 | 757501 | 1112504 | 143176 | 0.321 | 0.32 | 7.5E-01 | 0 |
| 22:6, docosahexaenoic acid (DHA) | 3 | 899838 | 1011653 | 696946 | 0.230 | 0.32 | 7.5E-01 | 0 |
| Osteoarthritis of hit or knee | 22 | 874041 | 1222210 | 167860 | 0.341 | 0.32 | 7.5E-01 | 0 |
| Cancer register - Type of cancer: ICD10: C50 Malignant neoplasm of breast | 9 | 936240 | 1069450 | 464265 | 0.397 | 0.33 | 7.4E-01 | 0 |
| Ever smoked | 41 | 869535 | 1255143 | 96500 | 0.322 | 0.33 | 7.4E-01 | 0 |
| Lipid::Lysolipid::1-arachidonoylglycerophosphocholine* | 3 | 918253 | 957725 | 903789 | 0.355 | 0.33 | 7.4E-01 | 0 |
| Sleep durataion | 9 | 832098 | 982820 | 98504 | 0.246 | 0.33 | 7.4E-01 | 0 |
| Left parahippocampal | 3 | 909942 | 1133163 | 502112 | 0.299 | 0.34 | 7.3E-01 | 0 |
| Cerebellar vermal lobules I V | 14 | 807139 | 1032873 | 105391 | 0.228 | 0.34 | 7.3E-01 | 0 |
| Lipid::Lysolipid::1-oleoylglycerophosphoethanolamine | 1 | 925430 | 925430 | 925430 | 0.222 | 0.34 | 7.3E-01 | 0 |
| Peptide::gamma-glutamyl::gamma-glutamylglutamine | 2 | 849843 | 1241605 | 458081 | 0.285 | 0.35 | 7.3E-01 | 0 |
| Total bilirubin | 8 | 738733 | 1342429 | 91881 | 0.196 | 0.35 | 7.3E-01 | 0 |
| PGC cross disorder | 4 | 799226 | 1225639 | 300538 | 0.306 | 0.35 | 7.3E-01 | 0 |
| Treatment/medication code: ramipril | 3 | 826673 | 1222518 | 610704 | 0.280 | 0.35 | 7.2E-01 | 0 |
| Lipid::Lysolipid::1-palmitoylglycerophosphoethanolamine | 1 | 925430 | 925430 | 925430 | 0.222 | 0.36 | 7.2E-01 | 0 |
| Right postcentral | 3 | 669333 | 909503 | 351999 | 0.194 | 0.36 | 7.2E-01 | 0 |
| Illnesses of siblings: Breast cancer | 3 | 951490 | 1005368 | 501975 | 0.377 | 0.36 | 7.2E-01 | 0 |
| Age stopped smoking | 1 | 1076735 | 1076735 | 1076735 | 0.452 | 0.36 | 7.2E-01 | 0 |
| ::::X-10510 | 3 | 816268 | 938858 | 810763 | 0.161 | 0.36 | 7.2E-01 | 0 |
| Non-butter spread type details: Flora Pro-Active or Benecol | 6 | 636213 | 926591 | 402315 | 0.124 | 0.37 | 7.1E-01 | 0 |
| Diuretics | 72 | 827005 | 1110203 | 341408 | 0.346 | 0.37 | 7.1E-01 | 0 |
| Corticospinal tract radial diusivities | 5 | 948305 | 1178285 | 870740 | 0.474 | 0.37 | 7.1E-01 | 0 |
| Hypothyroidism | 7 | 806485 | 1021384 | 292334 | 0.325 | 0.37 | 7.1E-01 | 0 |
| Daytime napping | 7 | 946520 | 1066675 | 399486 | 0.343 | 0.37 | 7.1E-01 | 0 |
| Total-body less head BMD and total body lean mass (bivariate meta-analysis) | 9 | 796800 | 1004520 | 666668 | 0.260 | 0.37 | 7.1E-01 | 0 |
| Focal Epilepsy, hippocampal sclerosis (HS) | 2 | 614200 | 817980 | 410421 | 0.080 | 0.37 | 7.1E-01 | 0 |
| Male-specific factors - Hair/balding pattern: Pattern 3 | 40 | 864370 | 1164823 | 210260 | 0.341 | 0.38 | 7.1E-01 | 0 |
| Diagnoses - secondary ICD10: I25 Chronic ischemic heart disease | 15 | 851270 | 1309281 | 431029 | 0.326 | 0.38 | 7.1E-01 | 0 |
| Diagnoses - secondary ICD10: I10 Essential (primary) hypertension | 82 | 831810 | 1139037 | 359159 | 0.321 | 0.38 | 7.0E-01 | 0 |
| Nucleotide::Purine metabolism, (hypo)xanthine, inosine containing::hypoxanthine | 1 | 1036650 | 1036650 | 1036650 | 0.377 | 0.39 | 7.0E-01 | 0 |
| Medication for pain relief, constipation, heartburn: Ibuprofen (e.g. Nurofen) | 7 | 912317 | 1326158 | 444606 | 0.351 | 0.39 | 7.0E-01 | 0 |
| Lipid::Sterol, Steroid::cholesterol | 1 | 946783 | 946783 | 946783 | 0.217 | 0.39 | 6.9E-01 | 0 |
| Posterior thalamic radiation (include optic radiation) axial diusivities | 2 | 938233 | 1255351 | 621114 | 0.299 | 0.40 | 6.9E-01 | 0 |
| Amino acid::Glycine, serine and threonine metabolism::N-acetylglycine | 2 | 1018876 | 1125942 | 911811 | 0.372 | 0.40 | 6.9E-01 | 0 |
| Heart rate recovery at 30 secnds | 19 | 767923 | 910101 | 102444 | 0.285 | 0.40 | 6.9E-01 | 0 |
| Guilty feelings | 13 | 881448 | 1264640 | 654675 | 0.303 | 0.40 | 6.9E-01 | 0 |
| Lipid::Carnitine metabolism::stearoylcarnitine | 1 | 1051960 | 1051960 | 1051960 | 0.372 | 0.40 | 6.9E-01 | 0 |
| Lipid::Monoacylglycerol::1-oleoylglycerol (1-monoolein) | 1 | 818555 | 818555 | 818555 | 0.129 | 0.41 | 6.8E-01 | 0 |

| | | | | | | | | |
|---|---|---|---|---|---|---|---|---|
| Sodium | 10 | 604684 | 1223986 | 71398 | 0.137 | 0.41 | 6.8E-01 | 0 |
| Mood swings | 36 | 888069 | 1397594 | 231215 | 0.323 | 0.41 | 6.8E-01 | 0 |
| Posterior corona radiata radial diusivities | 1 | 1113795 | 1113795 | 1113795 | 0.483 | 0.41 | 6.8E-01 | 0 |
| Free cholesterol in medium VLDL | 7 | 818555 | 1096754 | 297742 | 0.211 | 0.42 | 6.8E-01 | 0 |
| Standing height | 835 | 741713 | 1123048 | 140559 | 0.264 | 0.42 | 6.8E-01 | 0 |
| Glioma | 12 | 899935 | 1304187 | 617321 | 0.344 | 0.42 | 6.8E-01 | 0 |
| Coffee intake | 20 | 821479 | 1257946 | 168226 | 0.269 | 0.42 | 6.8E-01 | 0 |
| Mineral and other dietary supplements: Fish oil (including cod liver oil) | 6 | 753070 | 1034334 | 179905 | 0.239 | 0.42 | 6.8E-01 | 0 |
| Height SDS | 4 | 915130 | 1273486 | 454842 | 0.253 | 0.42 | 6.8E-01 | 0 |
| Pain type(s) experienced in last month: Back pain | 8 | 887976 | 998662 | 670678 | 0.288 | 0.42 | 6.8E-01 | 0 |
| Nucleotide::Purine metabolism, guanine containing::N2,N2-dimethylguanosine | 1 | 1060300 | 1060300 | 1060300 | 0.359 | 0.42 | 6.7E-01 | 0 |
| Bread intake | 15 | 913910 | 1714674 | 105688 | 0.388 | 0.42 | 6.7E-01 | 0 |
| Treatment/medication code: metformin | 28 | 881524 | 1168285 | 409937 | 0.290 | 0.42 | 6.7E-01 | 0 |
| Offspring birthweight (maternal) | 7 | 924228 | 1220385 | 660623 | 0.291 | 0.43 | 6.7E-01 | 0 |
| Glycine | 4 | 629152 | 1251847 | 41670 | 0.125 | 0.43 | 6.7E-01 | 0 |
| Diagnoses - secondary ICD10: K57 Diverticular disease of intestine | 10 | 849669 | 1025939 | 205592 | 0.259 | 0.43 | 6.7E-01 | 0 |
| Vascular/heart problems diagnosed by doctor: Angina | 14 | 842514 | 1116611 | 321236 | 0.272 | 0.43 | 6.7E-01 | 0 |
| Right cerebellum exterior | 19 | 713013 | 1050188 | 44579 | 0.283 | 0.44 | 6.6E-01 | 0 |
| Daytime dozing / sleeping (narcolepsy) | 26 | 897286 | 1134078 | 242213 | 0.338 | 0.44 | 6.6E-01 | 0 |
| Irritability | 33 | 873278 | 1238488 | 219206 | 0.269 | 0.44 | 6.6E-01 | 0 |
| Medication for cholesterol, blood pressure, diabetes, or take exogenous hormones: Cholesterol lowering medication | 27 | 809510 | 1178439 | 417343 | 0.268 | 0.44 | 6.6E-01 | 0 |
| Fluid intelligence test - FI7 : synonym | 1 | 961217 | 961217 | 961217 | 0.264 | 0.44 | 6.6E-01 | 0 |
| Menstrual fever | 1 | 548240 | 548240 | 548240 | 0.091 | 0.44 | 6.6E-01 | 0 |
| Never eat eggs, dairy, wheat, sugar: I eat all of the above | 11 | 924228 | 1128154 | 624635 | 0.373 | 0.44 | 6.6E-01 | 0 |
| Total cholesterol in small VLDL | 11 | 818555 | 999000 | 444793 | 0.220 | 0.44 | 6.6E-01 | 0 |
| Left insula | 4 | 627782 | 1311636 | 68113 | 0.117 | 0.45 | 6.6E-01 | 0 |
| Xenobiotics::Xanthine metabolism::3-methylxanthine | 2 | 953061 | 958687 | 947436 | 0.309 | 0.45 | 6.6E-01 | 0 |
| Intelligence | 195 | 852238 | 1176918 | 100995 | 0.321 | 0.45 | 6.6E-01 | 0 |
| Triglycerides in large VLDL | 6 | 824461 | 959667 | 532741 | 0.212 | 0.45 | 6.5E-01 | 0 |
| Number of days/week of moderate physical activity 10+ minutes | 5 | 889053 | 1369603 | 558700 | 0.279 | 0.45 | 6.5E-01 | 0 |
| High-density lipoprotein cholesterol | 98 | 764049 | 1119529 | 266303 | 0.251 | 0.45 | 6.5E-01 | 0 |
| Depressive affect subcluster | 58 | 855294 | 1307893 | 220314 | 0.326 | 0.45 | 6.5E-01 | 0 |
| Lipid::Sterol, Steroid::5alpha-androstan-3beta,17beta-diol disulfate | 3 | 640390 | 871106 | 345508 | 0.143 | 0.45 | 6.5E-01 | 0 |
| Treatment/medication code: ibuprofen | 3 | 912317 | 962444 | 467493 | 0.456 | 0.46 | 6.5E-01 | 0 |
| Pain type(s) experienced in last month: Knee pain | 8 | 942316 | 1020380 | 732324 | 0.320 | 0.46 | 6.5E-01 | 0 |
| High light scatter reticulocyte count (two-way meta) | 96 | 791799 | 1130026 | 264560 | 0.298 | 0.46 | 6.5E-01 | 0 |
| Amino acid::Alanine and aspartate metabolism::alanine | 2 | 1036141 | 1334754 | 737528 | 0.424 | 0.46 | 6.5E-01 | 0 |
| Mood swings (MOOD) | 40 | 888069 | 1370185 | 132709 | 0.323 | 0.46 | 6.4E-01 | 0 |
| Xenobiotics::Xanthine metabolism::caffeine | 1 | 1082660 | 1082660 | 1082660 | 0.369 | 0.46 | 6.4E-01 | 0 |
| Traumatic events - Felt loved as child | 3 | 976820 | 1178106 | 741361 | 0.376 | 0.47 | 6.4E-01 | 0 |
| Triglycerides in very small VLDL | 11 | 818555 | 1041381 | 296602 | 0.220 | 0.47 | 6.4E-01 | 0 |
| Total cholesterol in large HDL | 10 | 853804 | 1051374 | 596580 | 0.227 | 0.47 | 6.4E-01 | 0 |
| Free cholesterol in small VLDL | 11 | 797550 | 919060 | 423210 | 0.150 | 0.48 | 6.3E-01 | 0 |
| Mean erythrocyte cell volume | 4 | 978601 | 1075023 | 887002 | 0.356 | 0.48 | 6.3E-01 | 0 |
| Left cerebellum exterior | 11 | 763213 | 1391488 | 208020 | 0.283 | 0.48 | 6.3E-01 | 0 |
| Age first had sexual intercourse | 88 | 898039 | 1289598 | 453216 | 0.352 | 0.48 | 6.3E-01 | 0 |
| Waist-hip ratio | 464 | 826203 | 1157628 | 348876 | 0.314 | 0.48 | 6.3E-01 | 0 |
| Phospholipids in small VLDL | 9 | 818555 | 949067 | 514640 | 0.150 | 0.49 | 6.3E-01 | 0 |
| Lumbar Spine BMD | 38 | 894429 | 1223609 | 403358 | 0.339 | 0.49 | 6.2E-01 | 0 |
| Number of days/week walked 10+ minutes | 10 | 945119 | 1173691 | 89270 | 0.375 | 0.49 | 6.2E-01 | 0 |
| Impedance measures - Leg fat percentage (left) | 388 | 805615 | 1197839 | 196423 | 0.301 | 0.49 | 6.2E-01 | 0 |
| Thyroid preparations | 99 | 839208 | 1173291 | 436561 | 0.320 | 0.49 | 6.2E-01 | 0 |
| Phospholipids in IDL | 12 | 746061 | 1154155 | 95473 | 0.232 | 0.49 | 6.2E-01 | 0 |
| Alkaline phosphatase | 22 | 810458 | 1145148 | 442879 | 0.261 | 0.50 | 6.2E-01 | 0 |
| Blood clot, DVT, bronchitis, emphysema, asthma, rhinitis, eczema, allergy diagnosed by doctor: Asthma | 66 | 830396 | 1120137 | 217690 | 0.294 | 0.50 | 6.2E-01 | 0 |
| Opioids | 2 | 1048161 | 1179488 | 916834 | 0.434 | 0.50 | 6.2E-01 | 0 |

| | | | | | | | | |
|---|---|---|---|---|---|---|---|---|
| Free thyroxine (FT4) | 18 | 822717 | 1196573 | 448066 | 0.273 | 0.50 | 6.2E-01 | 0 |
| FEV1 | 406 | 826861 | 1198660 | 317436 | 0.314 | 0.50 | 6.2E-01 | 0 |
| Cingulum (cingulate gyrus) mean diusivities | 3 | 875653 | 1126300 | 670906 | 0.189 | 0.50 | 6.2E-01 | 0 |
| HDL diameter | 8 | 786835 | 1110696 | 567870 | 0.236 | 0.50 | 6.2E-01 | 0 |
| Posterior corona radiata mode of anisotropy | 3 | 988860 | 1428891 | 754460 | 0.382 | 0.51 | 6.1E-01 | 0 |
| Platelet count (three-way meta) | 166 | 775066 | 1164418 | 207387 | 0.283 | 0.51 | 6.1E-01 | 0 |
| Open-angle glaucoma | 6 | 935514 | 1277202 | 646358 | 0.367 | 0.52 | 6.0E-01 | 0 |
| Diagnoses - secondary ICD10: Z95 Presence of cardiac and vascular implants and grafts | 8 | 918658 | 1054393 | 808126 | 0.236 | 0.52 | 6.0E-01 | 0 |
| Histidine | 4 | 894400 | 1058084 | 640250 | 0.357 | 0.52 | 6.0E-01 | 0 |
| Headaches for 3+ months | 4 | 924608 | 1075645 | 649757 | 0.398 | 0.52 | 6.0E-01 | 0 |
| White blood cell count (three-way meta) | 118 | 825238 | 1215359 | 212914 | 0.303 | 0.53 | 6.0E-01 | 0 |
| Ever highly irritable/argumentative for 2 days | 3 | 886200 | 1003341 | 660144 | 0.328 | 0.53 | 6.0E-01 | 0 |
| Superior corona radiata mode of anisotropy | 3 | 982630 | 1103274 | 529679 | 0.298 | 0.53 | 6.0E-01 | 0 |
| Illnesses of mother: High blood pressure | 17 | 892065 | 1073905 | 341380 | 0.349 | 0.53 | 6.0E-01 | 0 |
| Right isthmus cingulate | 3 | 1073430 | 1189656 | 992874 | 0.356 | 0.54 | 5.9E-01 | 0 |
| Seen a psychiatrist for nerves, anxiety, tension or depression | 3 | 920673 | 1054678 | 483596 | 0.347 | 0.54 | 5.9E-01 | 0 |
| Happiness and subjective well-being - General happiness | 1 | 1181195 | 1181195 | 1181195 | 0.438 | 0.54 | 5.9E-01 | 0 |
| Illnesses of father: Alzheimer's disease/dementia | 1 | 889053 | 889053 | 889053 | 0.150 | 0.54 | 5.9E-01 | 0 |
| Lactate dehydrogenase | 5 | 1004188 | 1217050 | 617268 | 0.439 | 0.55 | 5.8E-01 | 0 |
| High-density-lipoprotein cholesterol | 22 | 700825 | 1043995 | 270208 | 0.236 | 0.55 | 5.8E-01 | 0 |
| Weight change compared with 1 year ago | 1 | 889053 | 889053 | 889053 | 0.150 | 0.56 | 5.8E-01 | 0 |
| Infantile hypertrophic pyloric stenosis | 6 | 818603 | 1039886 | 334516 | 0.215 | 0.56 | 5.7E-01 | 0 |
| Total cholesterol | 117 | 758930 | 1192875 | 395798 | 0.234 | 0.57 | 5.7E-01 | 0 |
| Non-cancer illness code, self-reported: hypertension | 151 | 788825 | 1139530 | 161005 | 0.281 | 0.57 | 5.7E-01 | 0 |
| Left superior temporal | 1 | 1191618 | 1191618 | 1191618 | 0.412 | 0.57 | 5.7E-01 | 0 |
| Illnesses of father: Bowel cancer | 3 | 1064808 | 1233548 | 842486 | 0.460 | 0.57 | 5.7E-01 | 0 |
| Heart rate recovery at 10 secnds | 16 | 776683 | 1143033 | 108092 | 0.213 | 0.58 | 5.6E-01 | 0 |
| Waist-hip ratio (adjusted for BMI) | 60 | 916700 | 1301019 | 420873 | 0.364 | 0.58 | 5.6E-01 | 0 |
| Fractional shortening | 3 | 1017888 | 1075320 | 935938 | 0.312 | 0.58 | 5.6E-01 | 0 |
| Hip circumference (male, adjusted for BMI) | 38 | 922967 | 1252582 | 141925 | 0.338 | 0.58 | 5.6E-01 | 0 |
| Age when last used oral contraceptive pill (female) | 1 | 1044205 | 1044205 | 1044205 | 0.281 | 0.58 | 5.6E-01 | 0 |
| ::::X-11440 | 1 | 622075 | 622075 | 622075 | 0.085 | 0.58 | 5.6E-01 | 0 |
| OmegaL3 fatty acids | 5 | 899838 | 1045825 | 494055 | 0.223 | 0.59 | 5.6E-01 | 0 |
| ::::X-12644 | 1 | 1045825 | 1045825 | 1045825 | 0.223 | 0.59 | 5.5E-01 | 0 |
| Concentration of large VLDL particles | 8 | 792240 | 1008001 | 391314 | 0.184 | 0.59 | 5.5E-01 | 0 |
| Mental health - Illness, injury, bereavement, stress in last 2 years: Financial difficulties | 3 | 1023008 | 1167565 | 854443 | 0.228 | 0.59 | 5.5E-01 | 0 |
| Fornix (cres) / Stria terminalis fractional anisotropy | 1 | 1138703 | 1138703 | 1138703 | 0.364 | 0.59 | 5.5E-01 | 0 |
| Amino acid::Phenylalanine & tyrosine metabolism::phenylalanine | 1 | 1148495 | 1148495 | 1148495 | 0.397 | 0.59 | 5.5E-01 | 0 |
| Lipid::Medium chain fatty acid::pelargonate (9:0) | 1 | 1151108 | 1151108 | 1151108 | 0.356 | 0.59 | 5.5E-01 | 0 |
| Alcohol dependency (full discovery samples) | 1 | 620200 | 620200 | 620200 | 0.003 | 0.60 | 5.5E-01 | 0 |
| Social support - Leisure/social activities: Sports club or gym | 9 | 948488 | 1036333 | 104020 | 0.294 | 0.60 | 5.5E-01 | 0 |
| Hematocrit (two-way meta) | 90 | 851206 | 1154568 | 232501 | 0.312 | 0.60 | 5.5E-01 | 0 |
| Anxiety disorder (case/control) | 1 | 1201733 | 1201733 | 1201733 | 0.437 | 0.60 | 5.5E-01 | 0 |
| Non-cancer illness code, self-reported: asthma | 51 | 848655 | 1136361 | 228042 | 0.293 | 0.60 | 5.5E-01 | 0 |
| Corrected insulin response | 3 | 960417 | 1036844 | 547583 | 0.275 | 0.60 | 5.5E-01 | 0 |
| Amino acid::Tryptophan metabolism::tryptophan betaine | 2 | 826492 | 979168 | 673818 | 0.144 | 0.60 | 5.5E-01 | 0 |
| Anterior corona radiata radial diusivities | 9 | 975900 | 1268678 | 466160 | 0.306 | 0.60 | 5.5E-01 | 0 |
| Age hay fever, rhinitis or eczema diagnosed | 24 | 849217 | 1202956 | 498121 | 0.254 | 0.60 | 5.5E-01 | 0 |
| Neutrophil count (two-way meta) | 65 | 810610 | 1219753 | 97941 | 0.263 | 0.60 | 5.5E-01 | 0 |
| Superior corona radiata radial diusivities | 4 | 800633 | 912428 | 524465 | 0.238 | 0.61 | 5.5E-01 | 0 |
| Total lipids in small VLDL | 11 | 818555 | 949054 | 430043 | 0.150 | 0.61 | 5.4E-01 | 0 |
| Current tobacco smoking | 11 | 957335 | 1211318 | 260396 | 0.354 | 0.61 | 5.4E-01 | 0 |
| Number of sleep episodes | 19 | 889053 | 973685 | 247575 | 0.283 | 0.61 | 5.4E-01 | 0 |
| Lipid::Long chain fatty acid::X-12442--5,8-tetradecadienoate | 1 | 1158020 | 1158020 | 1158020 | 0.386 | 0.61 | 5.4E-01 | 0 |
| Plateletcrit (two-way meta) | 125 | 786968 | 1110558 | 338498 | 0.271 | 0.61 | 5.4E-01 | 0 |
| Left ventral DC | 6 | 933798 | 1184813 | 300938 | 0.318 | 0.62 | 5.4E-01 | 0 |

| | | | | | | | | |
|---|---|---|---|---|---|---|---|---|
| Depressive symptoms (MA GWAMA) | 118 | 894703 | 1299175 | 446739 | 0.359 | 0.62 | 5.4E-01 | 0 |
| ::::X-12798 | 3 | 986395 | 1216105 | 523425 | 0.237 | 0.62 | 5.4E-01 | 0 |
| Phospholipids in chylomicrons and extremely large VLDL | 6 | 827303 | 966318 | 800503 | 0.215 | 0.62 | 5.4E-01 | 0 |
| Amino acid::Glycine, serine and threonine metabolism::betaine | 3 | 1036217 | 1223268 | 920481 | 0.312 | 0.62 | 5.3E-01 | 0 |
| Illnesses of mother: Heart disease | 2 | 1122575 | 1258228 | 986923 | 0.448 | 0.62 | 5.3E-01 | 0 |
| Mean platelet volume (two-way meta) | 141 | 771208 | 1187530 | 298253 | 0.281 | 0.62 | 5.3E-01 | 0 |
| Free cholesterol in large HDL | 11 | 889053 | 1130259 | 606613 | 0.221 | 0.62 | 5.3E-01 | 0 |
| Xenobiotics::Benzoate metabolism::4-vinylphenol sulfate | 1 | 1060360 | 1060360 | 1060360 | 0.243 | 0.63 | 5.3E-01 | 0 |
| Total protein | 22 | 762235 | 967114 | 543793 | 0.238 | 0.63 | 5.3E-01 | 0 |
| Phospholipids in very large HDL | 10 | 927767 | 1087011 | 598784 | 0.241 | 0.63 | 5.3E-01 | 0 |
| Ratio of visceral-tosubcutaneous adipose tissue volume | 3 | 1109360 | 1275428 | 607840 | 0.385 | 0.63 | 5.3E-01 | 0 |
| Job involves heavy manual or physical work | 8 | 1018508 | 1274003 | 561957 | 0.397 | 0.63 | 5.3E-01 | 0 |
| Baldness | 58 | 827924 | 1171887 | 434823 | 0.272 | 0.64 | 5.3E-01 | 0 |
| Diagnoses - secondary ICD10: Z82 Family history of certain disabilities and chronic diseases (leading to disablement) | 5 | 1077720 | 1138860 | 851270 | 0.406 | 0.64 | 5.2E-01 | 0 |
| Ever smoker | 161 | 873400 | 1234615 | 159386 | 0.340 | 0.64 | 5.2E-01 | 0 |
| Waist-hip ratio (male) | 133 | 873133 | 1210258 | 318705 | 0.354 | 0.64 | 5.2E-01 | 0 |
| Waist circumference (male) | 23 | 898940 | 1300483 | 499163 | 0.293 | 0.64 | 5.2E-01 | 0 |
| Cingulum (cingulate gyrus) fractional anisotropy | 4 | 818978 | 942934 | 579343 | 0.225 | 0.65 | 5.2E-01 | 0 |
| Isoleucine | 2 | 824168 | 1228768 | 419568 | 0.198 | 0.65 | 5.2E-01 | 0 |
| Anterior corona radiata mean diusivities | 5 | 975900 | 1147350 | 466160 | 0.286 | 0.65 | 5.2E-01 | 0 |
| Left lingual | 5 | 983585 | 1497278 | 94282 | 0.369 | 0.65 | 5.2E-01 | 0 |
| Risk taking | 16 | 982553 | 1272574 | 294044 | 0.395 | 0.65 | 5.2E-01 | 0 |
| Concentration of small HDL particles | 4 | 870363 | 1417038 | 301123 | 0.251 | 0.65 | 5.2E-01 | 0 |
| Total cholesterol in large VLDL | 11 | 765925 | 919409 | 116974 | 0.129 | 0.65 | 5.1E-01 | 0 |
| Glucose | 4 | 863451 | 1014154 | 689373 | 0.273 | 0.65 | 5.1E-01 | 0 |
| Impedance measures - Whole body water mass | 626 | 787340 | 1194653 | 143348 | 0.301 | 0.65 | 5.1E-01 | 0 |
| Amino acid::Valine, leucine and isoleucine metabolism::valine | 1 | 1226363 | 1226363 | 1226363 | 0.461 | 0.65 | 5.1E-01 | 0 |
| Amino acid::Valine, leucine and isoleucine metabolism::3-methyl-2-oxobutyrate | 1 | 1226363 | 1226363 | 1226363 | 0.461 | 0.65 | 5.1E-01 | 0 |
| Hemoglobin | 11 | 946020 | 1321773 | 266267 | 0.295 | 0.66 | 5.1E-01 | 0 |
| Ejection fraction | 2 | 970369 | 1028559 | 912178 | 0.251 | 0.66 | 5.1E-01 | 0 |
| Antimigraine preparations | 12 | 956521 | 1305876 | 419438 | 0.316 | 0.66 | 5.1E-01 | 0 |
| Ratio of bisLallylic bonds to total fatty acids in lipids | 2 | 1083444 | 1358406 | 808482 | 0.368 | 0.66 | 5.1E-01 | 0 |
| Cholesterol esters in medium VLDL | 13 | 818555 | 1048933 | 485140 | 0.150 | 0.66 | 5.1E-01 | 0 |
| Posterior cortical atrophy | 3 | 719763 | 804408 | 387647 | 0.106 | 0.66 | 5.1E-01 | 0 |
| Trail making test - Duration to complete alphanumeric path (trail #2) | 8 | 1011706 | 1550558 | 847574 | 0.373 | 0.66 | 5.1E-01 | 0 |
| Diastolic Blood Pressure (automated reading) | 194 | 796325 | 1152338 | 139206 | 0.287 | 0.66 | 5.1E-01 | 0 |
| Serum total triglycerides | 10 | 824461 | 934064 | 599834 | 0.172 | 0.66 | 5.1E-01 | 0 |
| Ever had known person concerned about, or recommended reduction of alcohol consumption | 2 | 774169 | 838752 | 709586 | 0.184 | 0.67 | 5.1E-01 | 0 |
| Reticulocyte fraction of red cells (two-way meta) | 108 | 816821 | 1102706 | 264560 | 0.298 | 0.67 | 5.0E-01 | 0 |
| Fasting insulin interaction (adjusted for BMI) | 10 | 316486 | 995656 | 37916 | 0.014 | 0.67 | 5.0E-01 | 0 |
| Mean corpuscular hemoglobin | 62 | 745123 | 1110478 | 153782 | 0.252 | 0.67 | 5.0E-01 | 0 |
| Schizophrenia vs Bipolar disorder | 75 | 873278 | 1273734 | 166938 | 0.326 | 0.67 | 5.0E-01 | 0 |
| Total lipids in large VLDL | 7 | 818555 | 979404 | 478593 | 0.129 | 0.67 | 5.0E-01 | 0 |
| Left accumbens area | 6 | 894191 | 1283032 | 204119 | 0.293 | 0.68 | 5.0E-01 | 0 |
| Plateletcrit (three-way meta) | 160 | 795856 | 1162109 | 323294 | 0.278 | 0.68 | 5.0E-01 | 0 |
| Ever unenthusiastic/disinterested for a whole week | 2 | 782587 | 1140742 | 424432 | 0.140 | 0.68 | 5.0E-01 | 0 |
| Antiglaucoma preparations and miotics | 11 | 940400 | 1241610 | 149024 | 0.239 | 0.68 | 5.0E-01 | 0 |
| Time spend outdoors in summer | 46 | 907829 | 1177597 | 101575 | 0.353 | 0.68 | 5.0E-01 | 0 |
| Diagnoses - secondary ICD10: I20 Angina pectoris | 12 | 921491 | 1185414 | 543718 | 0.303 | 0.68 | 4.9E-01 | 0 |
| Lipid::Carnitine metabolism::decanoylcarnitine | 3 | 1112685 | 1138233 | 825368 | 0.307 | 0.68 | 4.9E-01 | 0 |
| Phospholipids in large HDL | 9 | 943167 | 1141990 | 818555 | 0.233 | 0.69 | 4.9E-01 | 0 |
| Excessive hairiness | 3 | 1145635 | 1332681 | 912813 | 0.459 | 0.69 | 4.9E-01 | 0 |
| Illnesses of siblings: Diabetes | 7 | 1027155 | 1168869 | 957019 | 0.309 | 0.69 | 4.9E-01 | 0 |
| Amino acid::Butanoate metabolism::2-aminobutyrate | 2 | 1149435 | 1207090 | 1091780 | 0.434 | 0.70 | 4.9E-01 | 0 |
| Lipid::Long chain fatty acid::stearidonate (18:4n3) | 1 | 1181688 | 1181688 | 1181688 | 0.377 | 0.70 | 4.9E-01 | 0 |
| Amino acid::Guanidino and acetamido metabolism::4-acetamidobutanoate | 2 | 1078410 | 1149521 | 1007299 | 0.357 | 0.70 | 4.9E-01 | 0 |

| | | | | | | | | |
|---|---|---|---|---|---|---|---|---|
| Concentration of large HDL particles | 8 | 916110 | 1110696 | 760553 | 0.227 | 0.70 | 4.9E-01 | 0 |
| Fed-up feelings | 27 | 889275 | 1346724 | 93923 | 0.308 | 0.70 | 4.9E-01 | 0 |
| Phospholipids in very small VLDL | 14 | 761860 | 1037454 | 170611 | 0.220 | 0.70 | 4.9E-01 | 0 |
| Doctor diagnosed hayfever or allergic rhinitis | 21 | 896303 | 1174765 | 476268 | 0.289 | 0.70 | 4.9E-01 | 0 |
| Total lipids in large HDL | 8 | 916110 | 1110696 | 760553 | 0.227 | 0.70 | 4.9E-01 | 0 |
| HMG CoA reductase inhibitors | 58 | 826156 | 1142679 | 194463 | 0.292 | 0.70 | 4.8E-01 | 0 |
| Illnesses of mother: Diabetes | 19 | 889053 | 1137050 | 314279 | 0.291 | 0.70 | 4.8E-01 | 0 |
| Average total household income before tax | 28 | 835420 | 1201195 | 175231 | 0.288 | 0.70 | 4.8E-01 | 0 |
| Moderate to vigorous physical activity levels | 19 | 873448 | 1319581 | 61573 | 0.247 | 0.70 | 4.8E-01 | 0 |
| Lipid::Carnitine metabolism::3-dehydrocarnitine* | 2 | 980161 | 983278 | 977044 | 0.240 | 0.71 | 4.8E-01 | 0 |
| Extreme Height | 49 | 955428 | 1164553 | 345608 | 0.343 | 0.71 | 4.8E-01 | 0 |
| Superior longitudinal fasciculus mode of anisotropy | 1 | 1255143 | 1255143 | 1255143 | 0.416 | 0.71 | 4.8E-01 | 0 |
| Lipid::Sterol, Steroid::dehydroisoandrosterone sulfate (DHEA-S) | 2 | 729131 | 973597 | 484665 | 0.124 | 0.72 | 4.7E-01 | 0 |
| ::::X-08402 | 2 | 991992 | 1026720 | 957265 | 0.275 | 0.72 | 4.7E-01 | 0 |
| Lipid::Inositol metabolism::scyllo-inositol | 1 | 680895 | 680895 | 680895 | 0.064 | 0.73 | 4.7E-01 | 0 |
| Lipid::Sterol, Steroid::4-androsten-3beta,17beta-diol disulfate 1* | 2 | 729131 | 973597 | 484665 | 0.124 | 0.73 | 4.7E-01 | 0 |
| ::::X-09789 | 2 | 1078950 | 1149318 | 1008583 | 0.345 | 0.73 | 4.6E-01 | 0 |
| Illnesses of father: High blood pressure | 9 | 985248 | 1091518 | 479195 | 0.290 | 0.73 | 4.6E-01 | 0 |
| Age started hormone-replacement therapy (HRT) (female) | 7 | 922255 | 1408741 | 442714 | 0.316 | 0.73 | 4.6E-01 | 0 |
| Serum creatinine | 45 | 804745 | 1093425 | 214947 | 0.242 | 0.74 | 4.6E-01 | 0 |
| Lamb/mutton intake | 17 | 889053 | 1039123 | 83636 | 0.239 | 0.74 | 4.6E-01 | 0 |
| Phospholipids in very large VLDL | 7 | 836050 | 1068231 | 792240 | 0.230 | 0.74 | 4.6E-01 | 0 |
| Lipid::Lysolipid::1-arachidonoylglycerophosphoinositol* | 4 | 968436 | 1194688 | 708983 | 0.249 | 0.74 | 4.6E-01 | 0 |
| Diagnoses - main ICD10: R31 Hematuria | 4 | 841234 | 1251904 | 394397 | 0.188 | 0.74 | 4.6E-01 | 0 |
| Medication for cholesterol, blood pressure or diabetes: Cholesterol lowering medication | 33 | 833758 | 1047450 | 229329 | 0.289 | 0.74 | 4.6E-01 | 0 |
| Lipid::Medium chain fatty acid::10-undecenoate (11:1n1) | 1 | 992923 | 992923 | 992923 | 0.168 | 0.75 | 4.5E-01 | 0 |
| Total lipids in large LDL | 14 | 787651 | 1175429 | 377491 | 0.211 | 0.75 | 4.5E-01 | 0 |
| Concentration of large LDL particles | 14 | 787651 | 1175429 | 377491 | 0.211 | 0.75 | 4.5E-01 | 0 |
| Triglycerides in small VLDL | 11 | 830367 | 1137843 | 490970 | 0.194 | 0.75 | 4.5E-01 | 0 |
| Lipid::Lysolipid::1-stearoylglycerophosphoethanolamine | 1 | 1123655 | 1123655 | 1123655 | 0.217 | 0.76 | 4.5E-01 | 0 |
| Taking other prescription medications | 10 | 994278 | 1603742 | 489303 | 0.395 | 0.76 | 4.5E-01 | 0 |
| Diagnoses - secondary ICD10: E78 Disorders of lipoprotein metabolism and other lipidemias | 21 | 818555 | 1079695 | 214394 | 0.222 | 0.76 | 4.5E-01 | 0 |
| Daytime sleepiness / dozing | 1 | 701585 | 701585 | 701585 | 0.001 | 0.76 | 4.4E-01 | 0 |
| Right putamen | 9 | 1042845 | 1265200 | 73063 | 0.358 | 0.77 | 4.4E-01 | 0 |
| Pork intake | 20 | 607558 | 822370 | 88646 | 0.054 | 0.77 | 4.4E-01 | 0 |
| Frequency of walking for pleasure in last 4 weeks | 5 | 1025960 | 1286093 | 972390 | 0.409 | 0.77 | 4.4E-01 | 0 |
| Phospholipids in large VLDL | 7 | 818555 | 919409 | 314614 | 0.129 | 0.77 | 4.4E-01 | 0 |
| Arms-arm fat ratio (male) | 33 | 928230 | 1238453 | 220260 | 0.273 | 0.77 | 4.4E-01 | 0 |
| Cholesterol esters in large HDL | 9 | 889053 | 946992 | 586548 | 0.220 | 0.77 | 4.4E-01 | 0 |
| 3rd ventricle | 13 | 921338 | 1065428 | 398773 | 0.282 | 0.77 | 4.4E-01 | 0 |
| Never eat eggs, dairy, wheat, sugar: Sugar or foods/drinks containing sugar | 7 | 991630 | 1132256 | 486269 | 0.291 | 0.78 | 4.3E-01 | 0 |
| Mean arterial pressure | 20 | 867171 | 1133799 | 278778 | 0.311 | 0.78 | 4.3E-01 | 0 |
| Right handed | 1 | 1233555 | 1233555 | 1233555 | 0.387 | 0.78 | 4.3E-01 | 0 |
| Amino acid::Lysine metabolism::lysine | 1 | 1137315 | 1137315 | 1137315 | 0.273 | 0.79 | 4.3E-01 | 0 |
| Triglycerides in very large VLDL | 6 | 910661 | 1100963 | 779083 | 0.235 | 0.79 | 4.3E-01 | 0 |
| Leptin (adjusted for BMI) | 2 | 1159548 | 1169260 | 1149835 | 0.426 | 0.79 | 4.3E-01 | 0 |
| Had other major operations | 1 | 693428 | 693428 | 693428 | 0.066 | 0.79 | 4.3E-01 | 0 |
| Cigarettes per day | 19 | 922358 | 1098394 | 322805 | 0.314 | 0.79 | 4.3E-01 | 0 |
| Lumbar Spine BMD (females) | 20 | 953276 | 1213724 | 385099 | 0.336 | 0.79 | 4.3E-01 | 0 |
| Left cerebellum white matter | 11 | 977470 | 1141903 | 675613 | 0.265 | 0.79 | 4.3E-01 | 0 |
| Frequent insomnia symptoms | 40 | 970969 | 1204728 | 282681 | 0.371 | 0.80 | 4.2E-01 | 0 |
| Waist-hip ratio (adjusted for BMI, female) | 317 | 806353 | 1140565 | 265245 | 0.293 | 0.80 | 4.2E-01 | 0 |
| Eotaxin (CCL11) | 3 | 971555 | 1741565 | 500645 | 0.202 | 0.80 | 4.2E-01 | 0 |
| Diagnoses - main ICD10: K80 Cholelithiasis | 15 | 881008 | 1127658 | 80472 | 0.249 | 0.81 | 4.2E-01 | 0 |
| Rheumatoid Arthritis | 65 | 870620 | 1184573 | 431055 | 0.288 | 0.81 | 4.2E-01 | 0 |
| Lipid::Carnitine metabolism::acetylcarnitine | 1 | 1255280 | 1255280 | 1255280 | 0.364 | 0.81 | 4.2E-01 | 0 |

| | | | | | | | | |
|---|---|---|---|---|---|---|---|---|
| Total lipids in chylomicrons and extremely large VLDL | 6 | 883811 | 994572 | 631057 | 0.179 | 0.81 | 4.2E-01 | 0 |
| Other serious medical condition/disability (diagnosed by doctor) | 3 | 991492 | 1408148 | 546305 | 0.114 | 0.81 | 4.2E-01 | 0 |
| Height | 1685 | 811305 | 1161265 | 308065 | 0.301 | 0.82 | 4.1E-01 | 0 |
| Neovascular disease | 10 | 966042 | 1142473 | 580831 | 0.242 | 0.82 | 4.1E-01 | 0 |
| Birth length | 2 | 988131 | 1337664 | 638598 | 0.200 | 0.82 | 4.1E-01 | 0 |
| Non-cancer illness code, self-reported: high cholesterol | 48 | 860017 | 1209234 | 392902 | 0.283 | 0.82 | 4.1E-01 | 0 |
| ::::X-11470 | 1 | 732723 | 732723 | 732723 | 0.030 | 0.82 | 4.1E-01 | 0 |
| ::::X-18601 | 2 | 766418 | 937535 | 595300 | 0.120 | 0.82 | 4.1E-01 | 0 |
| Treatment/medication code: atenolol | 6 | 1037303 | 1204316 | 1017923 | 0.466 | 0.82 | 4.1E-01 | 0 |
| Mean corpuscular volume | 70 | 776734 | 1097958 | 356662 | 0.253 | 0.83 | 4.1E-01 | 0 |
| Total lipids in very large HDL | 9 | 946992 | 1100265 | 797550 | 0.248 | 0.83 | 4.1E-01 | 0 |
| Fractures | 16 | 976246 | 1276098 | 548429 | 0.349 | 0.83 | 4.1E-01 | 0 |
| Past tobacco smoking | 65 | 876990 | 1208848 | 87778 | 0.297 | 0.83 | 4.1E-01 | 0 |
| Xenobiotics::Xanthine metabolism::1-methylxanthine | 1 | 1165490 | 1165490 | 1165490 | 0.225 | 0.83 | 4.1E-01 | 0 |
| Asthma | 8 | 911909 | 1177654 | 502577 | 0.310 | 0.83 | 4.0E-01 | 0 |
| Left posterior cingulate | 1 | 727988 | 727988 | 727988 | 0.056 | 0.84 | 4.0E-01 | 0 |
| Lipid::Carnitine metabolism::2-tetradecenoyl carnitine | 1 | 1255280 | 1255280 | 1255280 | 0.364 | 0.84 | 4.0E-01 | 0 |
| ::::X-10395 | 1 | 1267410 | 1267410 | 1267410 | 0.370 | 0.84 | 4.0E-01 | 0 |
| ::::X-10429 | 1 | 1267410 | 1267410 | 1267410 | 0.370 | 0.84 | 4.0E-01 | 0 |
| Amino acid::Glutathione metabolism::5-oxoproline | 1 | 737328 | 737328 | 737328 | 0.086 | 0.85 | 4.0E-01 | 0 |
| Parkinson's disease age at onset | 1 | 1275658 | 1275658 | 1275658 | 0.362 | 0.85 | 4.0E-01 | 0 |
| Corticospinal tract mean diusivities | 2 | 1196910 | 1206223 | 1187598 | 0.481 | 0.86 | 3.9E-01 | 0 |
| Systemic Lupus Erythematosus | 59 | 746580 | 1215221 | 276415 | 0.226 | 0.86 | 3.9E-01 | 0 |
| Treatment/medication code: amlodipine | 15 | 1005775 | 1356610 | 769329 | 0.387 | 0.87 | 3.9E-01 | 0 |
| General risk tolerance | 24 | 978341 | 1113830 | 548938 | 0.360 | 0.87 | 3.8E-01 | 0 |
| Autism spectrum disorder | 3 | 1164070 | 1190443 | 970190 | 0.346 | 0.87 | 3.8E-01 | 0 |
| Coronary artery disease in diabetes | 1 | 1335308 | 1335308 | 1335308 | 0.487 | 0.87 | 3.8E-01 | 0 |
| Long-standing illness, disability or infirmity | 11 | 1057045 | 1610618 | 706278 | 0.396 | 0.87 | 3.8E-01 | 0 |
| Left caudate | 8 | 1049194 | 1116236 | 901826 | 0.340 | 0.88 | 3.8E-01 | 0 |
| Cheese intake | 61 | 767943 | 1156160 | 87946 | 0.222 | 0.89 | 3.7E-01 | 0 |
| Peptide::Dipeptide::X-12244--N-acetylcarnosine | 3 | 1191685 | 1234166 | 1125289 | 0.441 | 0.89 | 3.7E-01 | 0 |
| Coffee type: Ground coffee (include espresso, filter etc) | 17 | 960338 | 1291558 | 407690 | 0.334 | 0.89 | 3.7E-01 | 0 |
| Bipolar disorder | 12 | 984339 | 1212962 | 495684 | 0.357 | 0.90 | 3.7E-01 | 0 |
| Ease of getting up in the morning | 53 | 866553 | 1234990 | 187736 | 0.288 | 0.90 | 3.7E-01 | 0 |
| Heart rate increase | 15 | 997005 | 1280579 | 671900 | 0.320 | 0.91 | 3.6E-01 | 0 |
| Total cholesterol in medium VLDL | 10 | 883811 | 1099398 | 482911 | 0.170 | 0.91 | 3.6E-01 | 0 |
| Concentration of medium VLDL particles | 10 | 889717 | 1139916 | 625655 | 0.207 | 0.92 | 3.6E-01 | 0 |
| Total lipids in very small VLDL | 12 | 883811 | 1050478 | 379486 | 0.220 | 0.92 | 3.6E-01 | 0 |
| Current employment status: Doing unpaid or voluntary work | 1 | 760953 | 760953 | 760953 | 0.086 | 0.92 | 3.6E-01 | 0 |
| Hematocrit | 16 | 913133 | 1317203 | 374489 | 0.216 | 0.93 | 3.5E-01 | 0 |
| Triglycerides in chylomicrons and extremely large VLDL | 6 | 914148 | 1102706 | 303142 | 0.179 | 0.93 | 3.5E-01 | 0 |
| Aspartate aminotransferase | 16 | 894649 | 1144807 | 329670 | 0.248 | 0.93 | 3.5E-01 | 0 |
| Uric acid | 18 | 951434 | 1248551 | 83405 | 0.319 | 0.94 | 3.5E-01 | 0 |
| Amount of alcohol drunk on a typical drinking day | 6 | 999435 | 1344179 | 841460 | 0.245 | 0.94 | 3.5E-01 | 0 |
| Illnesses of father: Heart disease | 18 | 846216 | 1132386 | 321236 | 0.205 | 0.94 | 3.5E-01 | 0 |
| Length of mobile phone use | 21 | 948435 | 1247098 | 119511 | 0.312 | 0.94 | 3.4E-01 | 0 |
| Free cholesterol in large VLDL | 8 | 919409 | 1118562 | 805398 | 0.235 | 0.95 | 3.4E-01 | 0 |
| Right fusiform | 1 | 1317115 | 1317115 | 1317115 | 0.398 | 0.95 | 3.4E-01 | 0 |
| Left basal forebrain | 3 | 1065723 | 1487795 | 555382 | 0.203 | 0.96 | 3.4E-01 | 0 |
| Insomnia (female) | 8 | 948676 | 1381783 | 76377 | 0.288 | 0.96 | 3.4E-01 | 0 |
| VLDL diameter | 10 | 871420 | 1266549 | 782259 | 0.148 | 0.96 | 3.4E-01 | 0 |
| Lipid::Long chain fatty acid::nonadecanoate (19:0) | 1 | 1113115 | 1113115 | 1113115 | 0.167 | 0.97 | 3.3E-01 | 0 |
| Average weekly red wine intake | 4 | 1145219 | 1277241 | 1056739 | 0.392 | 0.97 | 3.3E-01 | 0 |
| Femoral Neck BMD (males) | 1 | 1384985 | 1384985 | 1384985 | 0.467 | 0.97 | 3.3E-01 | 0 |
| Lipid::Long chain fatty acid::margarate (17:0) | 1 | 1113115 | 1113115 | 1113115 | 0.167 | 0.97 | 3.3E-01 | 0 |
| Bread type: White | 13 | 1051725 | 1229118 | 114657 | 0.370 | 0.97 | 3.3E-01 | 0 |

| | | | | | | | | |
|---|---|---|---|---|---|---|---|---|
| Lipid::Long chain fatty acid::myristoleate (14:1n5) | 1 | 1113115 | 1113115 | 1113115 | 0.167 | 0.97 | 3.3E-01 | 0 |
| Pericardial adipose tissue volume | 2 | 1218301 | 1348081 | 1088522 | 0.407 | 0.97 | 3.3E-01 | 0 |
| Lipid::Long chain fatty acid::palmitoleate (16:1n7) | 1 | 1113115 | 1113115 | 1113115 | 0.167 | 0.97 | 3.3E-01 | 0 |
| Lipid::Medium chain fatty acid::5-dodecenoate (12:1n7) | 1 | 1113115 | 1113115 | 1113115 | 0.167 | 0.98 | 3.3E-01 | 0 |
| Lipid::Long chain fatty acid::palmitate (16:0) | 1 | 1113115 | 1113115 | 1113115 | 0.167 | 0.98 | 3.3E-01 | 0 |
| Lipid::Lysolipid::1-palmitoleoylglycerophosphocholine* | 1 | 1113115 | 1113115 | 1113115 | 0.167 | 0.98 | 3.3E-01 | 0 |
| Heart rate recovery at 40 secnds | 17 | 865542 | 1102708 | 348115 | 0.184 | 0.98 | 3.2E-01 | 0 |
| Vasodilators used in cardiac diseases | 1 | 1393880 | 1393880 | 1393880 | 0.491 | 0.99 | 3.2E-01 | 0 |
| Right inferior temporal | 2 | 1086555 | 1407824 | 765286 | 0.248 | 0.99 | 3.2E-01 | 0 |
| Maternal history of Alzheimer's disease | 2 | 1183306 | 1273576 | 1093037 | 0.324 | 1.00 | 3.2E-01 | 0 |
| Strenuous sports or other exercises | 12 | 966571 | 1341801 | 557027 | 0.305 | 1.00 | 3.2E-01 | 0 |
| Phosphatidylcholine and other cholines | 4 | 1008179 | 1067793 | 963848 | 0.205 | 1.00 | 3.2E-01 | 0 |
| Glycoprotein acetyls, mainly a1Lacid glycoprotein | 4 | 883811 | 1120143 | 617628 | 0.121 | 1.00 | 3.2E-01 | 0 |
| Weekly usage of mobile phone in last 3 months | 8 | 1059716 | 1351618 | 1000390 | 0.331 | 1.00 | 3.2E-01 | 0 |
| Right thalamus proper | 3 | 1173788 | 1253143 | 610160 | 0.328 | 1.00 | 3.2E-01 | 0 |
| Diagnoses - main ICD10: R07 Pain in throat and chest | 1 | 1246235 | 1246235 | 1246235 | 0.287 | 1.01 | 3.1E-01 | 0 |
| Calcium | 11 | 927923 | 1121278 | 621969 | 0.313 | 1.01 | 3.1E-01 | 0 |
| Loneliness | 12 | 1010251 | 1348374 | 156278 | 0.329 | 1.01 | 3.1E-01 | 0 |
| Right caudate | 6 | 1129000 | 1171759 | 1021284 | 0.400 | 1.02 | 3.1E-01 | 0 |
| Salad / raw vegetable intake | 49 | 521835 | 1066135 | 87995 | 0.009 | 1.02 | 3.1E-01 | 0 |
| Sleep sedentary | 5 | 1102738 | 1605853 | 759793 | 0.321 | 1.02 | 3.1E-01 | 0 |
| OmegaL7 and L9 and saturated fatty acids | 5 | 949067 | 1001603 | 818555 | 0.129 | 1.02 | 3.1E-01 | 0 |
| Illnesses of mother: Alzheimer's disease/dementia | 2 | 1183306 | 1273576 | 1093037 | 0.324 | 1.03 | 3.0E-01 | 0 |
| 18:2, linoleic acid (LA) | 11 | 818555 | 1073441 | 216544 | 0.129 | 1.03 | 3.0E-01 | 0 |
| Left pallidum | 10 | 1080873 | 1256461 | 1015996 | 0.354 | 1.03 | 3.0E-01 | 0 |
| Systolic Blood Pressure | 25 | 886298 | 1253125 | 394735 | 0.289 | 1.03 | 3.0E-01 | 0 |
| Cingulum (hippocampus) radial diusivities | 1 | 1347053 | 1347053 | 1347053 | 0.307 | 1.03 | 3.0E-01 | 0 |
| Concentration of very small VLDL particles | 12 | 883811 | 1070123 | 381858 | 0.216 | 1.03 | 3.0E-01 | 0 |
| Insulin sensitivity index (adjusted for age, sex. Bmi) | 1 | 1353228 | 1353228 | 1353228 | 0.352 | 1.03 | 3.0E-01 | 0 |
| Mental distress - Ever suffered mental distress preventing usual activities | 4 | 915492 | 1171415 | 523001 | 0.231 | 1.03 | 3.0E-01 | 0 |
| Heart rate recovery at 50 secnds | 17 | 865542 | 1102708 | 348115 | 0.191 | 1.03 | 3.0E-01 | 0 |
| Total lipids in very large VLDL | 7 | 949067 | 1071718 | 486283 | 0.129 | 1.03 | 3.0E-01 | 0 |
| 4th ventricle | 13 | 918940 | 1192940 | 158949 | 0.202 | 1.04 | 3.0E-01 | 0 |
| Low-density-lipoprotein cholesterol | 13 | 831683 | 1074380 | 454363 | 0.120 | 1.04 | 3.0E-01 | 0 |
| Paternal history of Alzheimer's disease | 2 | 1126449 | 1245147 | 1007751 | 0.284 | 1.04 | 3.0E-01 | 0 |
| Type 1 Diabetes | 74 | 894158 | 1187693 | 357298 | 0.285 | 1.04 | 3.0E-01 | 0 |
| Antithrombotic agents | 9 | 833758 | 1082570 | 385130 | 0.129 | 1.05 | 2.9E-01 | 0 |
| Lipid::Medium chain fatty acid::heptanoate (7:0) | 2 | 1267661 | 1356518 | 1178804 | 0.439 | 1.05 | 2.9E-01 | 0 |
| Other polyunsaturated fatty acids than 18:2 | 6 | 957164 | 1027002 | 636089 | 0.205 | 1.06 | 2.9E-01 | 0 |
| Milk type used: Full cream | 4 | 1148098 | 1407964 | 742005 | 0.384 | 1.06 | 2.9E-01 | 0 |
| Fluid intelligence test - Number of fluid intelligence questions attempted within time limit | 9 | 1137883 | 1319015 | 160094 | 0.424 | 1.07 | 2.9E-01 | 0 |
| ::::X-13671 | 1 | 1432335 | 1432335 | 1432335 | 0.457 | 1.07 | 2.9E-01 | 0 |
| Loneliness, isolation | 4 | 1141716 | 1363014 | 813379 | 0.381 | 1.07 | 2.8E-01 | 0 |
| Illnesses of father: Diabetes | 19 | 975063 | 1239001 | 472826 | 0.291 | 1.08 | 2.8E-01 | 0 |
| Lipid::Sterol, Steroid::androsterone sulfate | 2 | 914850 | 1052080 | 777620 | 0.149 | 1.08 | 2.8E-01 | 0 |
| Serum Urate | 23 | 1015240 | 1329810 | 453078 | 0.363 | 1.09 | 2.8E-01 | 0 |
| Free cholesterol | 9 | 899838 | 1045825 | 374945 | 0.201 | 1.09 | 2.8E-01 | 0 |
| Diagnoses - main ICD10: R10 Abdominal and pelvic pain | 1 | 845510 | 845510 | 845510 | 0.062 | 1.09 | 2.8E-01 | 0 |
| Generalised epilepsy | 10 | 1098946 | 1325679 | 772659 | 0.346 | 1.09 | 2.8E-01 | 0 |
| Nucleotide::Purine metabolism, urate metabolism::urate | 1 | 1294678 | 1294678 | 1294678 | 0.215 | 1.09 | 2.8E-01 | 0 |
| Amino acid::Phenylalanine & tyrosine metabolism::phenyllactate (PLA) | 1 | 841233 | 841233 | 841233 | 0.098 | 1.10 | 2.7E-01 | 0 |
| Non-cancer illness code, self-reported: eczema/dermatitis | 12 | 1063461 | 1186424 | 731673 | 0.409 | 1.10 | 2.7E-01 | 0 |
| Triglyceride | 24 | 924130 | 1057793 | 230395 | 0.260 | 1.10 | 2.7E-01 | 0 |
| Lipid::Long chain fatty acid::10-nonadecenoate (19:1n9) | 1 | 1301655 | 1301655 | 1301655 | 0.203 | 1.10 | 2.7E-01 | 0 |
| Noisy workplace | 1 | 1164618 | 1164618 | 1164618 | 0.124 | 1.10 | 2.7E-01 | 0 |
| White matter | 8 | 1017278 | 1261486 | 429354 | 0.246 | 1.11 | 2.7E-01 | 0 |

| | | | | | | | | |
|---|---|---|---|---|---|---|---|---|
| Energy::Krebs cycle::citrate | 4 | 1108011 | 1261512 | 879573 | 0.383 | 1.11 | 2.7E-01 | 0 |
| Frequency of tiredness / lethargy in last 2 weeks | 21 | 1011308 | 1222053 | 398010 | 0.336 | 1.11 | 2.7E-01 | 0 |
| Lipid::Long chain fatty acid::oleate (18:1n9) | 1 | 1301655 | 1301655 | 1301655 | 0.203 | 1.11 | 2.7E-01 | 0 |
| Free cholesterol in very large HDL | 10 | 997120 | 1266549 | 669396 | 0.235 | 1.11 | 2.7E-01 | 0 |
| Calcium channel blockers | 82 | 858356 | 1221251 | 317506 | 0.286 | 1.12 | 2.6E-01 | 0 |
| Sagittal stratum fractional anisotropy | 6 | 1070869 | 1154893 | 876299 | 0.317 | 1.12 | 2.6E-01 | 0 |
| Symbol digit substitution test - Number of symbol digit matches made correctly | 4 | 1003465 | 1162194 | 775033 | 0.172 | 1.12 | 2.6E-01 | 0 |
| Right pericalcarine | 3 | 1220193 | 1407921 | 1160199 | 0.336 | 1.12 | 2.6E-01 | 0 |
| Skin colour | 96 | 843245 | 1119064 | 117471 | 0.306 | 1.13 | 2.6E-01 | 0 |
| Uncinate fasciculus mode of anisotropy | 1 | 1409495 | 1409495 | 1409495 | 0.366 | 1.13 | 2.6E-01 | 0 |
| Esterified cholesterol | 9 | 970533 | 1048933 | 385130 | 0.223 | 1.13 | 2.6E-01 | 0 |
| Acetoacetate | 2 | 1016169 | 1114976 | 917362 | 0.116 | 1.13 | 2.6E-01 | 0 |
| Fasting glucose interaction (adjusted for BMI) | 5 | 527945 | 538765 | 512633 | 0.006 | 1.13 | 2.6E-01 | 0 |
| Sum neutrophil eosinophil count (two-way meta) | 74 | 875973 | 1216620 | 187466 | 0.290 | 1.14 | 2.5E-01 | 0 |
| Cereal intake | 27 | 889053 | 1246224 | 83385 | 0.252 | 1.14 | 2.5E-01 | 0 |
| Concentration of IDL particles | 12 | 831980 | 1050478 | 304479 | 0.202 | 1.15 | 2.5E-01 | 0 |
| Amino acid::Valine, leucine and isoleucine metabolism::2-hydroxyisobutyrate | 3 | 1233350 | 1380466 | 1152186 | 0.308 | 1.15 | 2.5E-01 | 0 |
| Alcohol intake versus 10 years previously | 10 | 1101795 | 1591109 | 794321 | 0.385 | 1.15 | 2.5E-01 | 0 |
| Myeloid white cell count (two-way meta) | 80 | 845603 | 1208599 | 103018 | 0.275 | 1.15 | 2.5E-01 | 0 |
| Posterior corona radiata mean diusivities | 1 | 1423275 | 1423275 | 1423275 | 0.328 | 1.15 | 2.5E-01 | 0 |
| Amino acid::Phenylalanine & tyrosine metabolism::tyrosine | 3 | 1217793 | 1366889 | 864193 | 0.290 | 1.16 | 2.5E-01 | 0 |
| Microalbuminuria | 1 | 1187663 | 1187663 | 1187663 | 0.149 | 1.16 | 2.5E-01 | 0 |
| Duration of walks | 5 | 1121600 | 1341578 | 757313 | 0.400 | 1.16 | 2.5E-01 | 0 |
| Urinary albumin-to-creatinine ratio | 1 | 1187663 | 1187663 | 1187663 | 0.149 | 1.16 | 2.5E-01 | 0 |
| Drinks per day | 28 | 952556 | 1272131 | 87686 | 0.306 | 1.16 | 2.4E-01 | 0 |
| Average weekly champagne plus white wine intake | 7 | 1134608 | 1405525 | 571003 | 0.373 | 1.17 | 2.4E-01 | 0 |
| Fraction of accelerations | 2 | 1315089 | 1408074 | 1222103 | 0.477 | 1.17 | 2.4E-01 | 0 |
| Low-density lipoprotein cholesterol | 77 | 833758 | 1278630 | 471527 | 0.230 | 1.17 | 2.4E-01 | 0 |
| Amino acid::Urea cycle; arginine-, proline-, metabolism::proline | 2 | 1151220 | 1207369 | 1095071 | 0.226 | 1.18 | 2.4E-01 | 0 |
| Concentration of very large VLDL particles | 7 | 949067 | 1071718 | 669135 | 0.129 | 1.18 | 2.4E-01 | 0 |
| Cannabis use - Ever taken cannabis | 7 | 904135 | 1237786 | 168857 | 0.126 | 1.19 | 2.4E-01 | 0 |
| Insulin sensitivity index (combined influence of the genotype effect adjusted for BMI and the interaction effect between the genotype and BMI on ISI) | 11 | 472168 | 593669 | 31838 | 0.027 | 1.19 | 2.3E-01 | 0 |
| Triglycerides in IDL | 12 | 976042 | 1106157 | 705657 | 0.216 | 1.19 | 2.3E-01 | 0 |
| Tense / 'highly strung' (TENSE) | 20 | 976814 | 1202053 | 345118 | 0.299 | 1.20 | 2.3E-01 | 0 |
| Illnesses of father: Chronic bronchitis/emphysema | 4 | 1026484 | 1485056 | 431966 | 0.288 | 1.20 | 2.3E-01 | 0 |
| Xenobiotics::Xanthine metabolism::1-methylurate | 1 | 1346083 | 1346083 | 1346083 | 0.222 | 1.20 | 2.3E-01 | 0 |
| Hemoglobin concentration (two-way meta) | 83 | 896078 | 1147904 | 461254 | 0.273 | 1.20 | 2.3E-01 | 0 |
| Total fatty acids | 9 | 970533 | 1097950 | 889053 | 0.188 | 1.20 | 2.3E-01 | 0 |
| Smoking status: Never | 65 | 963830 | 1307260 | 123532 | 0.345 | 1.21 | 2.3E-01 | 0 |
| Birth weight | 159 | 850303 | 1151315 | 286344 | 0.271 | 1.21 | 2.3E-01 | 0 |
| Medication for cholesterol, blood pressure, diabetes, or take exogenous hormones: Blood pressure medication | 76 | 912392 | 1185896 | 201241 | 0.342 | 1.21 | 2.3E-01 | 0 |
| Sum basophil neutrophil count (two-way meta) | 68 | 878646 | 1232085 | 103315 | 0.284 | 1.21 | 2.3E-01 | 0 |
| ::::X-14625 | 1 | 892930 | 892930 | 892930 | 0.086 | 1.22 | 2.2E-01 | 0 |
| Lipid::Carnitine metabolism::hexanoylcarnitine | 4 | 1200301 | 1352965 | 830523 | 0.305 | 1.22 | 2.2E-01 | 0 |
| Total phosphoglycerides | 6 | 1008179 | 1111728 | 950479 | 0.205 | 1.22 | 2.2E-01 | 0 |
| Processed meat intake | 34 | 799271 | 1117726 | 61463 | 0.184 | 1.22 | 2.2E-01 | 0 |
| ::::X-12556 | 1 | 1361353 | 1361353 | 1361353 | 0.288 | 1.22 | 2.2E-01 | 0 |
| Right pallidum | 12 | 1080873 | 1471642 | 990671 | 0.323 | 1.23 | 2.2E-01 | 0 |
| Waist-hip ratio (female > 50 yrs, adjusted for BMI) | 13 | 1109360 | 1305485 | 743133 | 0.385 | 1.23 | 2.2E-01 | 0 |
| Superior fronto-occipital fasciculus radial diusivities | 2 | 1332534 | 1489222 | 1175846 | 0.445 | 1.23 | 2.2E-01 | 0 |
| MonoLunsaturated fatty acids | 6 | 883811 | 988469 | 250732 | 0.121 | 1.24 | 2.2E-01 | 0 |
| Waist-hip ratio (female <= 50 yrs, adjusted for BMI) | 9 | 1131123 | 1348585 | 792000 | 0.358 | 1.24 | 2.2E-01 | 0 |
| Right accumbens area | 6 | 1055123 | 1240547 | 763986 | 0.261 | 1.24 | 2.1E-01 | 0 |
| Diagnoses - main ICD10: H26 Other cataract | 6 | 1048938 | 1397449 | 342022 | 0.259 | 1.25 | 2.1E-01 | 0 |
| Leg pain on walking | 1 | 1525045 | 1525045 | 1525045 | 0.492 | 1.25 | 2.1E-01 | 0 |
| Coronary artery disease and triglyceride (bivariate) | 125 | 850163 | 1055668 | 388948 | 0.278 | 1.26 | 2.1E-01 | 0 |

| | | | | | | | | |
|---|---|---|---|---|---|---|---|---|
| Types of physical activity in last 4 weeks: Strenuous sports | 4 | 1107429 | 1337814 | 688202 | 0.327 | 1.26 | 2.1E-01 | 0 |
| Neck/shoulder pain for 3+ months | 1 | 1477533 | 1477533 | 1477533 | 0.389 | 1.27 | 2.1E-01 | 0 |
| Anilides | 7 | 1088348 | 1271423 | 553504 | 0.228 | 1.27 | 2.0E-01 | 0 |
| Alcohol consumption | 1 | 1484388 | 1484388 | 1484388 | 0.379 | 1.28 | 2.0E-01 | 0 |
| Lipid::Medium chain fatty acid::laurate (12:0) | 2 | 1276010 | 1437755 | 1114265 | 0.381 | 1.29 | 2.0E-01 | 0 |
| Superior fronto-occipital fasciculus axial diusivities | 1 | 1381228 | 1381228 | 1381228 | 0.275 | 1.29 | 2.0E-01 | 0 |
| Triglycerides cholesterol | 61 | 895060 | 1138703 | 417360 | 0.283 | 1.29 | 2.0E-01 | 0 |
| Ratio of visceral-tosubcutaneous adipose tissue volume (female) | 1 | 1505643 | 1505643 | 1505643 | 0.317 | 1.29 | 2.0E-01 | 0 |
| Mean corpuscular hemoglobin concentration (two-way meta) | 42 | 934730 | 1227079 | 537324 | 0.292 | 1.29 | 2.0E-01 | 0 |
| Lumbar Spine BMD (males) | 3 | 1356720 | 1525356 | 936653 | 0.451 | 1.29 | 2.0E-01 | 0 |
| Granulocyte count (two-way meta) | 74 | 892709 | 1233725 | 267406 | 0.303 | 1.31 | 1.9E-01 | 0 |
| Mother's age | 2 | 1362870 | 1561574 | 1164166 | 0.457 | 1.31 | 1.9E-01 | 0 |
| Drugs used in diabetes | 45 | 922742 | 1007783 | 353550 | 0.259 | 1.32 | 1.9E-01 | 0 |
| Cerebellar vermal lobules VIII X | 23 | 1063038 | 1361818 | 611216 | 0.366 | 1.32 | 1.9E-01 | 0 |
| Parkinson disease of sibling pairs (tier 1) | 17 | 1047335 | 1358525 | 557648 | 0.315 | 1.32 | 1.9E-01 | 0 |
| Valine | 3 | 1264745 | 1382799 | 1153795 | 0.427 | 1.32 | 1.9E-01 | 0 |
| Transport type for commuting to job workplace: Cycle | 1 | 1559753 | 1559753 | 1559753 | 0.484 | 1.32 | 1.9E-01 | 0 |
| Amino acid::Cysteine, methionine, SAM, taurine metabolism::methionine | 1 | 1270505 | 1270505 | 1270505 | 0.132 | 1.33 | 1.8E-01 | 0 |
| Sleeplessness / insomnia | 28 | 1022165 | 1090000 | 490857 | 0.320 | 1.34 | 1.8E-01 | 0 |
| Sensitivity / hurt feelings | 30 | 984788 | 1155241 | 102087 | 0.280 | 1.34 | 1.8E-01 | 0 |
| Alanine aminotransferase | 20 | 910055 | 1116002 | 415264 | 0.202 | 1.35 | 1.8E-01 | 0 |
| OmegaL6 fatty acids | 7 | 1001603 | 1073441 | 677831 | 0.188 | 1.36 | 1.7E-01 | 0 |
| Lipid::Bile acid metabolism::deoxycholate | 1 | 1529150 | 1529150 | 1529150 | 0.345 | 1.36 | 1.7E-01 | 0 |
| Sum eosinophil basophil count (two-way meta) | 102 | 868940 | 1120523 | 195566 | 0.293 | 1.36 | 1.7E-01 | 0 |
| Body Mass Index (Dominance deviation model) | 1 | 1527753 | 1527753 | 1527753 | 0.311 | 1.36 | 1.7E-01 | 0 |
| Platelet count (two-way meta) | 138 | 824285 | 1138923 | 150059 | 0.262 | 1.37 | 1.7E-01 | 0 |
| Tense / 'highly strung' | 21 | 1039700 | 1240205 | 773518 | 0.315 | 1.37 | 1.7E-01 | 0 |
| Medication for pain relief, constipation, heartburn: Aspirin | 3 | 1054003 | 1068286 | 719566 | 0.119 | 1.37 | 1.7E-01 | 0 |
| Right insula | 5 | 1176590 | 1426355 | 854958 | 0.308 | 1.38 | 1.7E-01 | 0 |
| Diverticular disease | 53 | 948725 | 1175743 | 463995 | 0.262 | 1.41 | 1.6E-01 | 0 |
| Posterior limb of internal capsule radial diusivities | 8 | 1050655 | 1283801 | 532716 | 0.283 | 1.41 | 1.6E-01 | 0 |
| Frequency of unenthusiasm / disinterest in last 2 weeks | 4 | 1168816 | 1387924 | 801984 | 0.363 | 1.41 | 1.6E-01 | 0 |
| Sensitivity / hurt feelings (HURT) | 26 | 1034370 | 1155241 | 311621 | 0.347 | 1.41 | 1.6E-01 | 0 |
| Left putamen | 8 | 1178478 | 1253103 | 927500 | 0.355 | 1.41 | 1.6E-01 | 0 |
| Mean platelet volume (three-way meta) | 152 | 822603 | 1151882 | 267160 | 0.268 | 1.42 | 1.5E-01 | 0 |
| Childhood Absence Epilepsy | 2 | 1201125 | 1328860 | 1073390 | 0.227 | 1.43 | 1.5E-01 | 0 |
| Amino acid::Tryptophan metabolism::kynurenine | 2 | 1332175 | 1733535 | 930815 | 0.363 | 1.43 | 1.5E-01 | 0 |
| Illnesses of siblings: High blood pressure | 15 | 1035558 | 1242953 | 388766 | 0.289 | 1.44 | 1.5E-01 | 0 |
| Osteoarthritis (self reported) | 1 | 1573255 | 1573255 | 1573255 | 0.366 | 1.44 | 1.5E-01 | 0 |
| Left rostral anterior cingulate | 4 | 922554 | 1338835 | 389791 | 0.110 | 1.44 | 1.5E-01 | 0 |
| Amyotrophic lateral sclerosis (logistic) | 3 | 1263588 | 1334358 | 675340 | 0.232 | 1.46 | 1.5E-01 | 0 |
| Height (male) | 38 | 981094 | 1174986 | 303788 | 0.266 | 1.46 | 1.4E-01 | 0 |
| Age at first birth | 8 | 1209933 | 1261298 | 869150 | 0.375 | 1.46 | 1.4E-01 | 0 |
| White blood cell count (two-way meta) | 98 | 872283 | 1252935 | 319749 | 0.262 | 1.46 | 1.4E-01 | 0 |
| Left inferior parietal | 4 | 1214493 | 1386809 | 799920 | 0.273 | 1.46 | 1.4E-01 | 0 |
| Late pubertal growth | 1 | 1588765 | 1588765 | 1588765 | 0.322 | 1.47 | 1.4E-01 | 0 |
| Medication for pain relief, constipation, heartburn: Paracetamol | 9 | 1193768 | 1440695 | 1061773 | 0.346 | 1.47 | 1.4E-01 | 0 |
| Hemoglobin concentration (three-way meta) | 93 | 920392 | 1316713 | 318140 | 0.295 | 1.47 | 1.4E-01 | 0 |
| Mania - Ever had period extreme irritability | 1 | 1360448 | 1360448 | 1360448 | 0.167 | 1.48 | 1.4E-01 | 0 |
| Free cholesterol in IDL | 11 | 996508 | 1132234 | 84490 | 0.217 | 1.48 | 1.4E-01 | 0 |
| Lipid::Long chain fatty acid::stearate (18:0) | 2 | 1207385 | 1254520 | 1160250 | 0.185 | 1.48 | 1.4E-01 | 0 |
| Treatment/medication code: paracetamol | 4 | 1243300 | 1426584 | 1081704 | 0.271 | 1.50 | 1.3E-01 | 0 |
| Pubertal growth | 1 | 1642628 | 1642628 | 1642628 | 0.484 | 1.51 | 1.3E-01 | 0 |
| Tanner scale | 1 | 1613123 | 1613123 | 1613123 | 0.336 | 1.51 | 1.3E-01 | 0 |
| Amyotrophic lateral sclerosis | 6 | 935901 | 1369743 | 171442 | 0.151 | 1.51 | 1.3E-01 | 0 |
| Breastfed as a baby | 1 | 1504390 | 1504390 | 1504390 | 0.281 | 1.51 | 1.3E-01 | 0 |

| | | | | | | | | |
|---|---|---|---|---|---|---|---|---|
| Mental distress - Ever sought or received professional help for mental distress | 2 | 1129539 | 1134594 | 1124483 | 0.255 | 1.52 | 1.3E-01 | 0 |
| Red blood cell count (two-way meta) | 119 | 925328 | 1217070 | 466781 | 0.300 | 1.52 | 1.3E-01 | 0 |
| Fornix (cres) / Stria terminalis radial diusivities | 2 | 1420088 | 1560780 | 1279395 | 0.414 | 1.52 | 1.3E-01 | 0 |
| Amino acid::Valine, leucine and isoleucine metabolism::4-methyl-2-oxopentanoate | 2 | 1364501 | 1433571 | 1295432 | 0.332 | 1.52 | 1.3E-01 | 0 |
| Alzheimer's disease | 11 | 1109820 | 1196119 | 576701 | 0.303 | 1.52 | 1.3E-01 | 0 |
| Left entorhinal | 4 | 1068055 | 1685108 | 392550 | 0.240 | 1.53 | 1.3E-01 | 0 |
| Able to confide | 8 | 1230328 | 1433348 | 842774 | 0.374 | 1.53 | 1.3E-01 | 0 |
| Diastolic Blood Pressure | 23 | 984975 | 1269851 | 465290 | 0.288 | 1.53 | 1.3E-01 | 0 |
| Antisocial behavior (female) | 2 | 830614 | 1216454 | 444773 | 0.064 | 1.53 | 1.3E-01 | 0 |
| Sleep duration | 104 | 946296 | 1238599 | 343410 | 0.315 | 1.54 | 1.2E-01 | 0 |
| Depressive symptoms | 10 | 1172510 | 1239854 | 907187 | 0.390 | 1.54 | 1.2E-01 | 0 |
| Carbohydrate::Fructose, mannose, galactose, starch, and sucrose metabolism::mannose | 1 | 1633368 | 1633368 | 1633368 | 0.389 | 1.55 | 1.2E-01 | 0 |
| Total lipids in IDL | 13 | 970533 | 1055113 | 385130 | 0.217 | 1.55 | 1.2E-01 | 0 |
| Types of physical activity in last 4 weeks: Walking for pleasure (not as a means of transport) | 3 | 1400850 | 1478315 | 752347 | 0.354 | 1.56 | 1.2E-01 | 0 |
| Number of cigarettes previously smoked daily | 2 | 1347998 | 1502146 | 1193849 | 0.271 | 1.56 | 1.2E-01 | 0 |
| Pyruvate | 1 | 1633368 | 1633368 | 1633368 | 0.389 | 1.56 | 1.2E-01 | 0 |
| Diagnoses - secondary ICD10: Z72 Problems related to lifestyle | 1 | 1529998 | 1529998 | 1529998 | 0.222 | 1.57 | 1.2E-01 | 0 |
| Waist circumference (female, adjusted for BMI) | 36 | 1069156 | 1223924 | 760473 | 0.357 | 1.58 | 1.2E-01 | 0 |
| Amino acid::Valine, leucine and isoleucine metabolism::3-methyl-2-oxovalerate | 2 | 1429865 | 1531616 | 1328114 | 0.425 | 1.59 | 1.1E-01 | 0 |
| Diagnoses - secondary ICD10: Z92 Personal history of medical treatment | 3 | 1053010 | 1791330 | 1049563 | 0.099 | 1.60 | 1.1E-01 | 0 |
| Adiponectin level | 11 | 1111730 | 1281746 | 589694 | 0.297 | 1.61 | 1.1E-01 | 0 |
| Left inferior temporal | 2 | 1438299 | 1493834 | 1382763 | 0.389 | 1.61 | 1.1E-01 | 0 |
| Peptide::gamma-glutamyl::gamma-glutamylvaline | 1 | 1565210 | 1565210 | 1565210 | 0.276 | 1.61 | 1.1E-01 | 0 |
| Total cholesterol in medium HDL | 5 | 1249520 | 1391218 | 889053 | 0.179 | 1.61 | 1.1E-01 | 0 |
| Treatment/medication code: atorvastatin | 15 | 1074650 | 1165868 | 664050 | 0.288 | 1.61 | 1.1E-01 | 0 |
| ::::X-12696 | 2 | 1265263 | 1684281 | 846244 | 0.218 | 1.62 | 1.1E-01 | 0 |
| Total body BMD (45-60 years old) | 16 | 1143158 | 1458020 | 525667 | 0.340 | 1.63 | 1.0E-01 | 0 |
| Hematocrit (three-way meta) | 81 | 963245 | 1242993 | 322390 | 0.318 | 1.63 | 1.0E-01 | 0 |
| Waist-hip ratio (adjusted for BMI, male) | 128 | 954580 | 1197550 | 273060 | 0.356 | 1.65 | 9.9E-02 | 0 |
| Epilepsy | 4 | 1357533 | 1549868 | 1062998 | 0.354 | 1.65 | 9.9E-02 | 0 |
| Hair colour (natural, before greying): Blonde | 124 | 844196 | 1116346 | 253876 | 0.275 | 1.66 | 9.7E-02 | 0 |
| Light smokers, at least 100 smokes in lifetime | 2 | 1328129 | 1415968 | 1240289 | 0.263 | 1.69 | 9.0E-02 | 0 |
| Overall activity | 2 | 1395661 | 1403164 | 1388158 | 0.313 | 1.70 | 9.0E-02 | 0 |
| Amyotrophic lateral sclerosis (linear mixed model) | 5 | 1224425 | 1263588 | 1174275 | 0.242 | 1.71 | 8.7E-02 | 0 |
| Job involves mainly walking or standing | 10 | 1245224 | 1470354 | 973370 | 0.415 | 1.71 | 8.7E-02 | 0 |
| Diagnoses - secondary ICD10: K44 Diaphragmatic hernia | 3 | 1398910 | 1434606 | 1106865 | 0.314 | 1.71 | 8.7E-02 | 0 |
| Major depressive disorder (ICD-coded) | 1 | 1604963 | 1604963 | 1604963 | 0.255 | 1.72 | 8.6E-02 | 0 |
| Juvenile Myoclonic Epilepsy (JME) | 1 | 1725310 | 1725310 | 1725310 | 0.355 | 1.75 | 8.0E-02 | 0 |
| Mean corpuscular hemoglobin concentration (three-way meta) | 60 | 934730 | 1243933 | 550698 | 0.265 | 1.76 | 7.8E-02 | 0 |
| Eczema | 9 | 1250975 | 1392105 | 451063 | 0.332 | 1.76 | 7.8E-02 | 0 |
| Birth weight of first child (female) | 39 | 1007783 | 1305894 | 133390 | 0.314 | 1.77 | 7.7E-02 | 0 |
| Mean corpuscular hemoglobin concentration | 33 | 971610 | 1222290 | 245163 | 0.268 | 1.77 | 7.7E-02 | 0 |
| Proxy and clinically diagnosed Alzheimer's disease | 32 | 1001873 | 1288531 | 427361 | 0.295 | 1.78 | 7.6E-02 | 0 |
| Time spent outdoors in winter | 11 | 1181195 | 1235116 | 555412 | 0.269 | 1.83 | 6.8E-02 | 0 |
| Right middle temporal | 1 | 1772910 | 1772910 | 1772910 | 0.394 | 1.83 | 6.7E-02 | 0 |
| ::::X-12850 | 2 | 1267595 | 1350481 | 1184709 | 0.144 | 1.84 | 6.6E-02 | 0 |
| Poultry intake | 12 | 1226236 | 1249541 | 393457 | 0.359 | 1.84 | 6.6E-02 | 0 |
| Fed-up feelings (FED_UP) | 28 | 1072941 | 1317129 | 510283 | 0.305 | 1.88 | 6.0E-02 | 0 |
| Treatment/medication code: glucosamine product | 1 | 1558458 | 1558458 | 1558458 | 0.189 | 1.89 | 5.9E-02 | 0 |
| Red cell distribution width (two-way meta) | 99 | 910498 | 1213903 | 301390 | 0.282 | 1.92 | 5.4E-02 | 0 |
| Pain type(s) experienced in last month: Headache | 31 | 1061773 | 1216263 | 200157 | 0.315 | 1.92 | 5.4E-02 | 0 |
| Lipid::Inositol metabolism::myo-inositol | 2 | 1247600 | 1248508 | 1246693 | 0.209 | 1.93 | 5.3E-02 | 0 |
| C-reactive protein | 12 | 1049074 | 1229494 | 805354 | 0.159 | 1.96 | 5.0E-02 | 0 |
| Amyotrophic lateral sclerosis (LMM) | 4 | 1334358 | 1863639 | 969464 | 0.237 | 1.96 | 4.9E-02 | 0 |
| Carbohydrate::Aminosugars metabolism::erythronate* | 3 | 1516068 | 1629749 | 1335585 | 0.251 | 1.98 | 4.8E-02 | 0 |
| Diagnoses - main ICD10: M23 Internal derangement of knee | 1 | 1884003 | 1884003 | 1884003 | 0.411 | 1.99 | 4.7E-02 | 0 |

| | | | | | | | | |
|---|---|---|---|---|---|---|---|---|
| Genu of corpus callosum fractional anisotropy | 5 | 1377788 | 1381305 | 577610 | 0.257 | 1.99 | 4.6E-02 | 0 |
| Lactate | 2 | 1566465 | 1599916 | 1533014 | 0.378 | 1.99 | 4.6E-02 | 0 |
| Family history of Alzheimer's disease | 6 | 1365889 | 1602587 | 1348568 | 0.331 | 2.01 | 4.4E-02 | 0 |
| Fasting insulin main effect | 3 | 1382225 | 1433306 | 1214708 | 0.172 | 2.03 | 4.2E-02 | 0 |
| Number of pregnancy terminations (female) | 4 | 790909 | 995873 | 469390 | 0.001 | 2.04 | 4.1E-02 | 0 |
| Salt added to food | 71 | 1019300 | 1258454 | 526048 | 0.310 | 2.07 | 3.9E-02 | 0 |
| Platelet count | 67 | 924842 | 1167686 | 532243 | 0.257 | 2.11 | 3.5E-02 | 0 |
| High blood pressure | 235 | 858542 | 1126979 | 237043 | 0.276 | 2.12 | 3.4E-02 | 0 |
| Amino acid::Histidine metabolism::histidine | 1 | 1843255 | 1843255 | 1843255 | 0.204 | 2.17 | 3.0E-02 | 0 |
| Red cell distribution width (three-way meta) | 112 | 913854 | 1185386 | 322062 | 0.282 | 2.22 | 2.6E-02 | 0 |
| Femoral Neck BMD (females) | 10 | 1349605 | 1404806 | 992534 | 0.338 | 2.23 | 2.6E-02 | 0 |
| Hair colour (natural, before greying): Black | 134 | 776128 | 1175388 | 106876 | 0.176 | 2.24 | 2.5E-02 | 0 |
| Alanine | 4 | 1476583 | 1625281 | 1015272 | 0.409 | 2.27 | 2.3E-02 | 0 |
| Amino acid::Amino fatty acid::X-12510--2-aminooctanoic acid | 2 | 1378240 | 1449574 | 1306906 | 0.244 | 2.28 | 2.3E-02 | 0 |
| Free cholesterol in medium HDL | 4 | 1386433 | 1415448 | 1183306 | 0.249 | 2.29 | 2.2E-02 | 0 |
| Superior fronto-occipital fasciculus mean diusivities | 2 | 1571661 | 1638739 | 1504583 | 0.311 | 2.31 | 2.1E-02 | 0 |
| ::::X-12063 | 4 | 1313896 | 1412121 | 926203 | 0.152 | 2.31 | 2.1E-02 | 0 |
| Why stopped smoking: Health precaution | 2 | 1648120 | 2260548 | 1035693 | 0.385 | 2.32 | 2.1E-02 | 0 |
| Phospholipids in medium HDL | 4 | 1363576 | 1384329 | 1155333 | 0.189 | 2.32 | 2.0E-02 | 0 |
| Waist circumference (adjusted for BMI) | 89 | 1036333 | 1250248 | 550950 | 0.325 | 2.34 | 1.9E-02 | 0 |
| Lean body mass (whole body) | 1 | 1408560 | 1408560 | 1408560 | 0.034 | 2.37 | 1.8E-02 | 0 |
| Fasting insulin main effect (adjusted for BMI) | 9 | 1391218 | 1518550 | 936215 | 0.260 | 2.43 | 1.5E-02 | 0 |
| Amino acid::Urea cycle; arginine-, proline-, metabolism::dimethylarginine (SDMA + ADMA) | 1 | 1443173 | 1443173 | 1443173 | 0.100 | 2.47 | 1.3E-02 | 0 |
| Getting up in morning | 52 | 1038246 | 1241453 | 185757 | 0.261 | 2.55 | 1.1E-02 | 0 |
| Ease of skin tanning | 319 | 628330 | 1088799 | 76151 | 0.086 | 3.01 | 2.6E-03 | 0 |
| Ulcerative colitis | 213 | 595308 | 934892 | 225410 | 0.024 | 3.45 | 5.7E-04 | 0 |
| Age at last live birth (female) | 1 | 2660000 | 2660000 | 2660000 | 0.488 | 3.58 | 3.4E-04 | 0 |
