## Supplementary Table 3 for "Mapping the genetic evolutionary timeline of human neural and cognitive traits"

| **Brain structure** | **Phenotype** | **N SNPs** |
|---|---|---|
| Amygdala | IDP SWI T2star left amygdala | 1 |
| Amygdala | IDP SWI T2star right amygdala | 14 |
| Amygdala | IDP T1 FAST ROIs L amygdala | 3 |
| Amygdala | IDP T1 FAST ROIs R amygdala | 1 |
| Amygdala | IDP tfMRI 90th percentile BOLD faces shapes amygdala | 8 |
| Amygdala | IDP tfMRI 90th percentile zstat faces shapes amygdala | 1 |
| Amygdala | IDP tfMRI median BOLD faces shapes amygdala | 2 |
| Amygdala | IDP tfMRI median zstat faces shapes amygdala | 9 |
| Caudate nucleus | IDP SWI T2star left caudate | 3 |
| Caudate nucleus | IDP SWI T2star right caudate | 1 |
| Caudate nucleus | IDP T1 FAST ROIs L caudate | 6 |
| Caudate nucleus | IDP T1 FAST ROIs R caudate | 4 |
| Caudate nucleus | IDP T1 FIRST left caudate volume | 7 |
| Caudate nucleus | IDP T1 FIRST right caudate volume | 3 |
| Caudate nucleus | Volume Right Caudate | 1 |
| Cerebellum | IDP T1 FAST ROIs L cerebellum I IV | 4 |
| Cerebellum | IDP T1 FAST ROIs L cerebellum IX | 6 |
| Cerebellum | IDP T1 FAST ROIs L cerebellum V | 4 |
| Cerebellum | IDP T1 FAST ROIs L cerebellum VI | 3 |
| Cerebellum | IDP T1 FAST ROIs L cerebellum VIIIa | 6 |
| Cerebellum | IDP T1 FAST ROIs L cerebellum VIIIb | 2 |
| Cerebellum | IDP T1 FAST ROIs L cerebellum VIIb | 5 |
| Cerebellum | IDP T1 FAST ROIs L cerebellum X | 2 |
| Cerebellum | IDP T1 FAST ROIs L cerebellum crus I | 10 |

| | | |
|---|---|---|
| Cerebellum | IDP T1 FAST ROIs L cerebellum crus II | 10 |
| Cerebellum | IDP T1 FAST ROIs R cerebellum I IV | 4 |
| Cerebellum | IDP T1 FAST ROIs R cerebellum IX | 2 |
| Cerebellum | IDP T1 FAST ROIs R cerebellum V | 3 |
| Cerebellum | IDP T1 FAST ROIs R cerebellum VI | 3 |
| Cerebellum | IDP T1 FAST ROIs R cerebellum VIIIa | 3 |
| Cerebellum | IDP T1 FAST ROIs R cerebellum VIIIb | 2 |
| Cerebellum | IDP T1 FAST ROIs R cerebellum VIIb | 7 |
| Cerebellum | IDP T1 FAST ROIs R cerebellum X | 3 |
| Cerebellum | IDP T1 FAST ROIs R cerebellum crus I | 10 |
| Cerebellum | IDP T1 FAST ROIs R cerebellum crus II | 12 |
| Cerebellum | IDP T1 FAST ROIs V cerebellum IX | 4 |
| Cerebellum | IDP T1 FAST ROIs V cerebellum VI | 9 |
| Cerebellum | IDP T1 FAST ROIs V cerebellum VIIIa | 6 |
| Cerebellum | IDP T1 FAST ROIs V cerebellum VIIIb | 10 |
| Cerebellum | IDP T1 FAST ROIs V cerebellum X | 5 |
| Cerebellum | IDP T1 FAST ROIs V cerebellum crus II | 8 |
| Cerebellum | Volume Right Cerebellum White Matter | 2 |
| Hippocampus | IDP SWI T2star left hippocampus | 8 |
| Hippocampus | IDP SWI T2star right hippocampus | 7 |
| Hippocampus | IDP T1 FAST ROIs L hippocampus | 4 |
| Hippocampus | IDP T1 FAST ROIs R hippocampus | 4 |
| Hippocampus | IDP T1 FIRST left hippocampus volume | 4 |
| Hippocampus | IDP T1 FIRST right hippocampus volume | 2 |
| Neocortex | DKTatlas lh lateralorbitofrontal area | 1 |

| | | |
|---|---|---|
| Neocortex | DKTatlas lh parahippocampal area | 2 |
| Neocortex | DKTatlas lh pericalcarine area | 1 |
| Neocortex | DKTatlas lh pericalcarine thickness | 1 |
| Neocortex | DKTatlas lh precentral thickness | 1 |
| Neocortex | DKTatlas rh caudalanteriorcingulate thickness | 1 |
| Neocortex | DKTatlas rh entorhinal area | 1 |
| Neocortex | DKTatlas rh inferiortemporal area | 2 |
| Neocortex | DKTatlas rh lingual area | 1 |
| Neocortex | DKTatlas rh medialorbitofrontal area | 1 |
| Neocortex | DKTatlas rh paracentral thickness | 1 |
| Neocortex | DKTatlas rh precuneus thickness | 2 |
| Neocortex | IDP T1 FAST ROIs L cuneal cortex | 1 |
| Neocortex | IDP T1 FAST ROIs L front med cortex | 1 |
| Neocortex | IDP T1 FAST ROIs L front operc cortex | 3 |
| Neocortex | IDP T1 FAST ROIs L front orb cortex | 3 |
| Neocortex | IDP T1 FAST ROIs L insular cortex | 3 |
| Neocortex | IDP T1 FAST ROIs L intracalc cortex | 14 |
| Neocortex | IDP T1 FAST ROIs L latocc cortex inf | 1 |
| Neocortex | IDP T1 FAST ROIs L latocc cortex sup | 1 |
| Neocortex | IDP T1 FAST ROIs L precun cortex | 1 |
| Neocortex | IDP T1 FAST ROIs L subcallosal cortex | 4 |
| Neocortex | IDP T1 FAST ROIs L supracalc cortex | 4 |
| Neocortex | IDP T1 FAST ROIs L temp fusif cortex post | 2 |
| Neocortex | IDP T1 FAST ROIs L temp occ fusif cortex | 1 |
| Neocortex | IDP T1 FAST ROIs R cuneal cortex | 1 |

| | | |
|---|---|---|
| Neocortex | IDP T1 FAST ROIs R front operc cortex | 1 |
| Neocortex | IDP T1 FAST ROIs R front orb cortex | 3 |
| Neocortex | IDP T1 FAST ROIs R insular cortex | 6 |
| Neocortex | IDP T1 FAST ROIs R intracalc cortex | 9 |
| Neocortex | IDP T1 FAST ROIs R latocc cortex sup | 1 |
| Neocortex | IDP T1 FAST ROIs R parietal operc cortex | 3 |
| Neocortex | IDP T1 FAST ROIs R subcallosal cortex | 5 |
| Neocortex | IDP T1 FAST ROIs R supracalc cortex | 1 |
| Neocortex | IDP T1 FAST ROIs R temp fusif cortex ant | 1 |
| Neocortex | IDP T1 SIENAX grey normalised volume | 2 |
| Neocortex | IDP T1 SIENAX peripheral grey normalised volume | 4 |
| Neocortex | IDP T1 SIENAX peripheral grey unnormalised volume | 1 |
| Neocortex | a2009s lh G oc temp med Parahip thickness | 1 |
| Neocortex | a2009s lh G occipital middle area | 1 |
| Neocortex | a2009s lh G orbital area | 1 |
| Neocortex | a2009s lh G temp sup G T transv thickness | 1 |
| Neocortex | a2009s lh G temporal inf thickness | 1 |
| Neocortex | a2009s lh G temporal middle thickness | 1 |
| Neocortex | a2009s lh G&S frontomargin area | 1 |
| Neocortex | a2009s lh Lat Fis ant Vertical thickness | 1 |
| Neocortex | a2009s lh S central area | 1 |
| Neocortex | a2009s lh S cingul Marginalis area | 1 |
| Neocortex | a2009s lh S circular insula sup thickness | 1 |
| Neocortex | a2009s lh S front middle area | 1 |
| Neocortex | a2009s lh S front middle thickness | 1 |

| | | |
|---|---|---|
| Neocortex | a2009s lh S intrapariet&P trans area | 1 |
| Neocortex | a2009s lh S orbital H Shaped area | 1 |
| Neocortex | a2009s lh S orbital lateral area | 1 |
| Neocortex | a2009s lh S parieto occipital area | 1 |
| Neocortex | a2009s lh S precentral sup part area | 1 |
| Neocortex | a2009s lh S suborbital thickness | 1 |
| Neocortex | a2009s lh S temporal inf thickness | 1 |
| Neocortex | a2009s rh G Ins lg&S cent ins thickness | 1 |
| Neocortex | a2009s rh G oc temp med Parahip thickness | 3 |
| Neocortex | a2009s rh G orbital area | 1 |
| Neocortex | a2009s rh G pariet inf Angular thickness | 1 |
| Neocortex | a2009s rh G subcallosal area | 1 |
| Neocortex | a2009s rh G subcallosal thickness | 2 |
| Neocortex | a2009s rh G temp sup Lateral area | 1 |
| Neocortex | a2009s rh G temporal inf thickness | 1 |
| Neocortex | a2009s rh G&S frontomargin thickness | 1 |
| Neocortex | a2009s rh Lat Fis ant Vertical area | 1 |
| Neocortex | a2009s rh S calcarine area | 1 |
| Neocortex | a2009s rh S circular insula ant area | 1 |
| Neocortex | a2009s rh S oc middle&Lunatus thickness | 1 |
| Neocortex | a2009s rh S oc temp med&Lingual thickness | 1 |
| Neocortex | a2009s rh S orbital lateral thickness | 1 |
| Neocortex | a2009s rh S precentral sup part area | 1 |
| Nucleus accumbens | IDP SWI T2star left accumbens | 4 |
| Nucleus accumbens | IDP SWI T2star right accumbens | 5 |

| | | |
|---|---|---|
| Nucleus accumbens | IDP T1 FIRST left accumbens volume | 2 |
| Pallidum | IDP SWI T2star left pallidum | 2 |
| Pallidum | IDP SWI T2star right pallidum | 8 |
| Pallidum | IDP T1 FAST ROIs L pallidum | 1 |
| Pallidum | IDP T1 FAST ROIs R pallidum | 1 |
| Pallidum | IDP T1 FIRST left pallidum volume | 7 |
| Pallidum | IDP T1 FIRST right pallidum volume | 2 |
| Thalamus | IDP SWI T2star left thalamus | 2 |
| Thalamus | IDP SWI T2star right thalamus | 1 |
| Thalamus | IDP T1 FAST ROIs L thalamus | 8 |
| Thalamus | IDP T1 FAST ROIs R thalamus | 4 |
| Thalamus | IDP T1 FIRST left thalamus volume | 4 |
| White matter | IDP T1 SIENAX white normalised volume | 5 |
| White matter | IDP T2 FLAIR BIANCA WMH volume | 2 |
| White matter | IDP dMRI ProbtrackX FA ar r | 1 |
| White matter | IDP dMRI ProbtrackX FA atr l | 1 |
| White matter | IDP dMRI ProbtrackX FA cgc r | 1 |
| White matter | IDP dMRI ProbtrackX FA cgh l | 1 |
| White matter | IDP dMRI ProbtrackX FA cst r | 2 |
| White matter | IDP dMRI ProbtrackX FA fma | 2 |
| White matter | IDP dMRI ProbtrackX FA ifo r | 1 |
| White matter | IDP dMRI ProbtrackX FA ilf l | 1 |
| White matter | IDP dMRI ProbtrackX FA ml r | 1 |
| White matter | IDP dMRI ProbtrackX FA ptr r | 2 |
| White matter | IDP dMRI ProbtrackX FA str l | 1 |

| | | |
|---|---|---|
| White matter | IDP dMRI ProbtrackX FA str r | 2 |
| White matter | IDP dMRI ProbtrackX FA unc r | 2 |
| White matter | IDP dMRI ProbtrackX ICVF ar r | 4 |
| White matter | IDP dMRI ProbtrackX ICVF atr l | 2 |
| White matter | IDP dMRI ProbtrackX ICVF cgc r | 3 |
| White matter | IDP dMRI ProbtrackX ICVF cgh l | 1 |
| White matter | IDP dMRI ProbtrackX ICVF cgh r | 1 |
| White matter | IDP dMRI ProbtrackX ICVF cst l | 1 |
| White matter | IDP dMRI ProbtrackX ICVF cst r | 1 |
| White matter | IDP dMRI ProbtrackX ICVF fma | 2 |
| White matter | IDP dMRI ProbtrackX ICVF fmi | 3 |
| White matter | IDP dMRI ProbtrackX ICVF ifo l | 3 |
| White matter | IDP dMRI ProbtrackX ICVF ifo r | 1 |
| White matter | IDP dMRI ProbtrackX ICVF mcp | 1 |
| White matter | IDP dMRI ProbtrackX ICVF ptr r | 2 |
| White matter | IDP dMRI ProbtrackX ICVF str r | 1 |
| White matter | IDP dMRI ProbtrackX ISOVF cst r | 2 |
| White matter | IDP dMRI ProbtrackX ISOVF fmi | 1 |
| White matter | IDP dMRI ProbtrackX ISOVF ilf r | 1 |
| White matter | IDP dMRI ProbtrackX ISOVF ml l | 1 |
| White matter | IDP dMRI ProbtrackX ISOVF ptr l | 3 |
| White matter | IDP dMRI ProbtrackX ISOVF slf l | 1 |
| White matter | IDP dMRI ProbtrackX ISOVF str l | 1 |
| White matter | IDP dMRI ProbtrackX ISOVF str r | 1 |
| White matter | IDP dMRI ProbtrackX ISOVF unc r | 2 |

| | | |
|---|---|---|
| White matter | IDP dMRI ProbtrackX L1 ar r | 1 |
| White matter | IDP dMRI ProbtrackX L1 atr l | 5 |
| White matter | IDP dMRI ProbtrackX L1 atr r | 1 |
| White matter | IDP dMRI ProbtrackX L1 fma | 1 |
| White matter | IDP dMRI ProbtrackX L1 ml l | 1 |
| White matter | IDP dMRI ProbtrackX L1 ml r | 2 |
| White matter | IDP dMRI ProbtrackX L1 ptr l | 1 |
| White matter | IDP dMRI ProbtrackX L1 ptr r | 1 |
| White matter | IDP dMRI ProbtrackX L1 slf l | 2 |
| White matter | IDP dMRI ProbtrackX L1 unc r | 2 |
| White matter | IDP dMRI ProbtrackX L2 ar l | 1 |
| White matter | IDP dMRI ProbtrackX L2 cgc l | 3 |
| White matter | IDP dMRI ProbtrackX L2 cgc r | 1 |
| White matter | IDP dMRI ProbtrackX L2 cgh l | 1 |
| White matter | IDP dMRI ProbtrackX L2 cgh r | 1 |
| White matter | IDP dMRI ProbtrackX L2 cst r | 7 |
| White matter | IDP dMRI ProbtrackX L2 fma | 1 |
| White matter | IDP dMRI ProbtrackX L2 ifo r | 2 |
| White matter | IDP dMRI ProbtrackX L2 slf r | 1 |
| White matter | IDP dMRI ProbtrackX L2 str r | 2 |
| White matter | IDP dMRI ProbtrackX L3 cgc r | 2 |
| White matter | IDP dMRI ProbtrackX L3 cgh l | 2 |
| White matter | IDP dMRI ProbtrackX L3 cst r | 2 |
| White matter | IDP dMRI ProbtrackX L3 ifo l | 1 |
| White matter | IDP dMRI ProbtrackX L3 ifo r | 1 |

| | | |
|---|---|---|
| White matter | IDP dMRI ProbtrackX L3 ilf l | 1 |
| White matter | IDP dMRI ProbtrackX L3 ilf r | 1 |
| White matter | IDP dMRI ProbtrackX L3 slf l | 1 |
| White matter | IDP dMRI ProbtrackX L3 slf r | 1 |
| White matter | IDP dMRI ProbtrackX L3 str l | 2 |
| White matter | IDP dMRI ProbtrackX L3 str r | 1 |
| White matter | IDP dMRI ProbtrackX L3 unc l | 3 |
| White matter | IDP dMRI ProbtrackX MD ar l | 2 |
| White matter | IDP dMRI ProbtrackX MD ar r | 1 |
| White matter | IDP dMRI ProbtrackX MD atr r | 4 |
| White matter | IDP dMRI ProbtrackX MD cgc l | 3 |
| White matter | IDP dMRI ProbtrackX MD cgc r | 3 |
| White matter | IDP dMRI ProbtrackX MD cgh r | 2 |
| White matter | IDP dMRI ProbtrackX MD cst l | 1 |
| White matter | IDP dMRI ProbtrackX MD fmi | 1 |
| White matter | IDP dMRI ProbtrackX MD ifo l | 1 |
| White matter | IDP dMRI ProbtrackX MD ifo r | 4 |
| White matter | IDP dMRI ProbtrackX MD ilf r | 5 |
| White matter | IDP dMRI ProbtrackX MD mcp | 2 |
| White matter | IDP dMRI ProbtrackX MD ml r | 1 |
| White matter | IDP dMRI ProbtrackX MD ptr l | 2 |
| White matter | IDP dMRI ProbtrackX MD ptr r | 1 |
| White matter | IDP dMRI ProbtrackX MD slf l | 2 |
| White matter | IDP dMRI ProbtrackX MD slf r | 10 |
| White matter | IDP dMRI ProbtrackX MD str l | 2 |

| | | |
|---|---|---|
| White matter | IDP dMRI ProbtrackX MD str r | 1 |
| White matter | IDP dMRI ProbtrackX MD unc l | 2 |
| White matter | IDP dMRI ProbtrackX MD unc r | 1 |
| White matter | IDP dMRI ProbtrackX MO ar l | 1 |
| White matter | IDP dMRI ProbtrackX MO ar r | 1 |
| White matter | IDP dMRI ProbtrackX MO atr r | 1 |
| White matter | IDP dMRI ProbtrackX MO cgc l | 4 |
| White matter | IDP dMRI ProbtrackX MO cgh r | 1 |
| White matter | IDP dMRI ProbtrackX MO cst l | 1 |
| White matter | IDP dMRI ProbtrackX MO cst r | 1 |
| White matter | IDP dMRI ProbtrackX MO fma | 2 |
| White matter | IDP dMRI ProbtrackX MO fmi | 2 |
| White matter | IDP dMRI ProbtrackX MO ifo l | 3 |
| White matter | IDP dMRI ProbtrackX MO ilf r | 1 |
| White matter | IDP dMRI ProbtrackX MO ptr r | 4 |
| White matter | IDP dMRI ProbtrackX MO slf r | 3 |
| White matter | IDP dMRI ProbtrackX MO str l | 2 |
| White matter | IDP dMRI ProbtrackX MO str r | 3 |
| White matter | IDP dMRI ProbtrackX MO unc r | 1 |
| White matter | IDP dMRI ProbtrackX OD ar r | 1 |
| White matter | IDP dMRI ProbtrackX OD atr l | 1 |
| White matter | IDP dMRI ProbtrackX OD cgh r | 1 |
| White matter | IDP dMRI ProbtrackX OD ifo r | 2 |
| White matter | IDP dMRI ProbtrackX OD ilf r | 1 |
| White matter | IDP dMRI ProbtrackX OD ml r | 1 |

| | | |
|---|---|---|
| White matter | IDP dMRI ProbtrackX OD ptr l | 2 |
| White matter | IDP dMRI ProbtrackX OD ptr r | 8 |
| White matter | IDP dMRI ProbtrackX OD str l | 4 |
| White matter | IDP dMRI ProbtrackX OD unc r | 1 |
| White matter | IDP dMRI TBSS FA Anterior corona radiata L | 2 |
| White matter | IDP dMRI TBSS FA Anterior corona radiata R | 2 |
| White matter | IDP dMRI TBSS FA Anterior limb of internal capsule L | 2 |
| White matter | IDP dMRI TBSS FA Anterior limb of internal capsule R | 4 |
| White matter | IDP dMRI TBSS FA Body of corpus callosum | 6 |
| White matter | IDP dMRI TBSS FA Cerebral peduncle L | 11 |
| White matter | IDP dMRI TBSS FA Cerebral peduncle R | 6 |
| White matter | IDP dMRI TBSS FA Cingulum cingulate gyrus L | 2 |
| White matter | IDP dMRI TBSS FA Cingulum hippocampus L | 4 |
| White matter | IDP dMRI TBSS FA Corticospinal tract L | 1 |
| White matter | IDP dMRI TBSS FA Corticospinal tract R | 7 |
| White matter | IDP dMRI TBSS FA Fornix | 2 |
| White matter | IDP dMRI TBSS FA Genu of corpus callosum | 5 |
| White matter | IDP dMRI TBSS FA Inferior cerebellar peduncle L | 5 |
| White matter | IDP dMRI TBSS FA Medial lemniscus L | 1 |
| White matter | IDP dMRI TBSS FA Medial lemniscus R | 6 |
| White matter | IDP dMRI TBSS FA Middle cerebellar peduncle | 8 |
| White matter | IDP dMRI TBSS FA Pontine crossing tract | 5 |
| White matter | IDP dMRI TBSS FA Posterior corona radiata L | 1 |
| White matter | IDP dMRI TBSS FA Posterior corona radiata R | 1 |
| White matter | IDP dMRI TBSS FA Posterior limb of internal capsule L | 3 |

| | | |
|---|---|---|
| White matter | IDP dMRI TBSS FA Posterior limb of internal capsule R | 3 |
| White matter | IDP dMRI TBSS FA Posterior thalamic radiation R | 2 |
| White matter | IDP dMRI TBSS FA Retrolenticular part of internal capsule L | 1 |
| White matter | IDP dMRI TBSS FA Retrolenticular part of internal capsule R | 6 |
| White matter | IDP dMRI TBSS FA Sagittal stratum R | 1 |
| White matter | IDP dMRI TBSS FA Splenium of corpus callosum | 1 |
| White matter | IDP dMRI TBSS FA Superior cerebellar peduncle R | 6 |
| White matter | IDP dMRI TBSS FA Superior corona radiata L | 2 |
| White matter | IDP dMRI TBSS FA Superior corona radiata R | 1 |
| White matter | IDP dMRI TBSS FA Superior longitudinal fasciculus L | 1 |
| White matter | IDP dMRI TBSS FA Superior longitudinal fasciculus R | 1 |
| White matter | IDP dMRI TBSS FA Tapetum L | 4 |
| White matter | IDP dMRI TBSS ICVF Anterior corona radiata R | 1 |
| White matter | IDP dMRI TBSS ICVF Cerebral peduncle L | 4 |
| White matter | IDP dMRI TBSS ICVF Cingulum cingulate gyrus L | 2 |
| White matter | IDP dMRI TBSS ICVF Cingulum hippocampus L | 2 |
| White matter | IDP dMRI TBSS ICVF Fornix cres and Stria terminalis L | 1 |
| White matter | IDP dMRI TBSS ICVF Fornix cres and Stria terminalis R | 2 |
| White matter | IDP dMRI TBSS ICVF Genu of corpus callosum | 1 |
| White matter | IDP dMRI TBSS ICVF Inferior cerebellar peduncle L | 2 |
| White matter | IDP dMRI TBSS ICVF Medial lemniscus L | 2 |
| White matter | IDP dMRI TBSS ICVF Pontine crossing tract | 1 |
| White matter | IDP dMRI TBSS ICVF Posterior corona radiata R | 1 |
| White matter | IDP dMRI TBSS ICVF Superior corona radiata L | 2 |
| White matter | IDP dMRI TBSS ICVF Uncinate fasciculus L | 1 |

| | | |
|---|---|---|
| White matter | IDP dMRI TBSS ISOVF Anterior limb of internal capsule R | 1 |
| White matter | IDP dMRI TBSS ISOVF Cingulum hippocampus L | 4 |
| White matter | IDP dMRI TBSS ISOVF Fornix | 1 |
| White matter | IDP dMRI TBSS ISOVF Fornix cres+Stria terminalis L | 1 |
| White matter | IDP dMRI TBSS ISOVF Fornix cres+Stria terminalis R | 2 |
| White matter | IDP dMRI TBSS ISOVF Genu of corpus callosum | 1 |
| White matter | IDP dMRI TBSS ISOVF Inferior cerebellar peduncle R | 1 |
| White matter | IDP dMRI TBSS ISOVF Posterior limb of internal capsule L | 1 |
| White matter | IDP dMRI TBSS ISOVF Posterior limb of internal capsule R | 1 |
| White matter | IDP dMRI TBSS ISOVF Sagittal stratum L | 1 |
| White matter | IDP dMRI TBSS ISOVF Splenium of corpus callosum | 1 |
| White matter | IDP dMRI TBSS ISOVF Superior corona radiata L | 1 |
| White matter | IDP dMRI TBSS ISOVF Superior fronto occipital fasciculus L | 1 |
| White matter | IDP dMRI TBSS ISOVF Superior fronto occipital fasciculus R | 1 |
| White matter | IDP dMRI TBSS L1 Anterior corona radiata L | 9 |
| White matter | IDP dMRI TBSS L1 Anterior corona radiata R | 2 |
| White matter | IDP dMRI TBSS L1 Anterior limb of internal capsule L | 2 |
| White matter | IDP dMRI TBSS L1 Body of corpus callosum | 7 |
| White matter | IDP dMRI TBSS L1 Cerebral peduncle L | 6 |
| White matter | IDP dMRI TBSS L1 Cerebral peduncle R | 1 |
| White matter | IDP dMRI TBSS L1 Cingulum cingulate gyrus L | 1 |
| White matter | IDP dMRI TBSS L1 Cingulum hippocampus L | 1 |
| White matter | IDP dMRI TBSS L1 Cingulum hippocampus R | 1 |
| White matter | IDP dMRI TBSS L1 Corticospinal tract L | 7 |
| White matter | IDP dMRI TBSS L1 External capsule L | 1 |

| | | |
|---|---|---|
| White matter | IDP dMRI TBSS L1 External capsule R | 1 |
| White matter | IDP dMRI TBSS L1 Genu of corpus callosum | 3 |
| White matter | IDP dMRI TBSS L1 Middle cerebellar peduncle | 16 |
| White matter | IDP dMRI TBSS L1 Pontine crossing tract | 1 |
| White matter | IDP dMRI TBSS L1 Posterior corona radiata L | 1 |
| White matter | IDP dMRI TBSS L1 Posterior corona radiata R | 1 |
| White matter | IDP dMRI TBSS L1 Posterior limb of internal capsule L | 1 |
| White matter | IDP dMRI TBSS L1 Posterior thalamic radiation L | 1 |
| White matter | IDP dMRI TBSS L1 Retrolenticular part of internal capsule L | 1 |
| White matter | IDP dMRI TBSS L1 Retrolenticular part of internal capsule R | 3 |
| White matter | IDP dMRI TBSS L1 Sagittal stratum L | 5 |
| White matter | IDP dMRI TBSS L1 Superior cerebellar peduncle L | 2 |
| White matter | IDP dMRI TBSS L1 Superior corona radiata R | 2 |
| White matter | IDP dMRI TBSS L1 Superior fronto occipital fasciculus L | 3 |
| White matter | IDP dMRI TBSS L1 Superior longitudinal fasciculus L | 1 |
| White matter | IDP dMRI TBSS L1 Superior longitudinal fasciculus R | 3 |
| White matter | IDP dMRI TBSS L1 Tapetum L | 1 |
| White matter | IDP dMRI TBSS L1 Uncinate fasciculus L | 2 |
| White matter | IDP dMRI TBSS L2 Anterior corona radiata L | 1 |
| White matter | IDP dMRI TBSS L2 Anterior corona radiata R | 1 |
| White matter | IDP dMRI TBSS L2 Anterior limb of internal capsule R | 1 |
| White matter | IDP dMRI TBSS L2 Body of corpus callosum | 1 |
| White matter | IDP dMRI TBSS L2 Cerebral peduncle R | 2 |
| White matter | IDP dMRI TBSS L2 Cingulum cingulate gyrus L | 1 |
| White matter | IDP dMRI TBSS L2 Cingulum cingulate gyrus R | 1 |

| | | |
|---|---|---|
| White matter | IDP dMRI TBSS L2 Cingulum hippocampus L | 1 |
| White matter | IDP dMRI TBSS L2 Cingulum hippocampus R | 1 |
| White matter | IDP dMRI TBSS L2 Corticospinal tract L | 1 |
| White matter | IDP dMRI TBSS L2 External capsule L | 2 |
| White matter | IDP dMRI TBSS L2 External capsule R | 3 |
| White matter | IDP dMRI TBSS L2 Genu of corpus callosum | 1 |
| White matter | IDP dMRI TBSS L2 Medial lemniscus L | 1 |
| White matter | IDP dMRI TBSS L2 Medial lemniscus R | 10 |
| White matter | IDP dMRI TBSS L2 Middle cerebellar peduncle | 1 |
| White matter | IDP dMRI TBSS L2 Pontine crossing tract | 1 |
| White matter | IDP dMRI TBSS L2 Posterior limb of internal capsule R | 2 |
| White matter | IDP dMRI TBSS L2 Posterior thalamic radiation L | 1 |
| White matter | IDP dMRI TBSS L2 Posterior thalamic radiation R | 1 |
| White matter | IDP dMRI TBSS L2 Retrolenticular part of internal capsule L | 1 |
| White matter | IDP dMRI TBSS L2 Sagittal stratum L | 3 |
| White matter | IDP dMRI TBSS L2 Superior cerebellar peduncle R | 3 |
| White matter | IDP dMRI TBSS L2 Superior fronto occipital fasciculus R | 1 |
| White matter | IDP dMRI TBSS L2 Superior longitudinal fasciculus L | 1 |
| White matter | IDP dMRI TBSS L2 Tapetum L | 2 |
| White matter | IDP dMRI TBSS L2 Tapetum R | 3 |
| White matter | IDP dMRI TBSS L2 Uncinate fasciculus L | 1 |
| White matter | IDP dMRI TBSS L3 Anterior corona radiata R | 1 |
| White matter | IDP dMRI TBSS L3 Body of corpus callosum | 2 |
| White matter | IDP dMRI TBSS L3 Cerebral peduncle R | 1 |
| White matter | IDP dMRI TBSS L3 Cingulum cingulate gyrus L | 1 |

| | | |
|---|---|---|
| White matter | IDP dMRI TBSS L3 Cingulum hippocampus L | 1 |
| White matter | IDP dMRI TBSS L3 External capsule R | 2 |
| White matter | IDP dMRI TBSS L3 Genu of corpus callosum | 1 |
| White matter | IDP dMRI TBSS L3 Inferior cerebellar peduncle L | 1 |
| White matter | IDP dMRI TBSS L3 Pontine crossing tract | 1 |
| White matter | IDP dMRI TBSS L3 Posterior corona radiata R | 1 |
| White matter | IDP dMRI TBSS L3 Posterior limb of internal capsule R | 1 |
| White matter | IDP dMRI TBSS L3 Posterior thalamic radiation R | 1 |
| White matter | IDP dMRI TBSS L3 Retrolenticular part of internal capsule R | 1 |
| White matter | IDP dMRI TBSS L3 Splenium of corpus callosum | 3 |
| White matter | IDP dMRI TBSS L3 Superior corona radiata L | 1 |
| White matter | IDP dMRI TBSS L3 Superior fronto occipital fasciculus R | 1 |
| White matter | IDP dMRI TBSS L3 Superior longitudinal fasciculus R | 1 |
| White matter | IDP dMRI TBSS L3 Uncinate fasciculus R | 1 |
| White matter | IDP dMRI TBSS MD Anterior corona radiata L | 1 |
| White matter | IDP dMRI TBSS MD Anterior limb of internal capsule L | 1 |
| White matter | IDP dMRI TBSS MD Anterior limb of internal capsule R | 1 |
| White matter | IDP dMRI TBSS MD Body of corpus callosum | 1 |
| White matter | IDP dMRI TBSS MD Cerebral peduncle L | 1 |
| White matter | IDP dMRI TBSS MD Cingulum cingulate gyrus L | 1 |
| White matter | IDP dMRI TBSS MD Cingulum cingulate gyrus R | 2 |
| White matter | IDP dMRI TBSS MD Cingulum hippocampus R | 2 |
| White matter | IDP dMRI TBSS MD Corticospinal tract R | 1 |
| White matter | IDP dMRI TBSS MD External capsule L | 2 |
| White matter | IDP dMRI TBSS MD External capsule R | 3 |

| | | |
|---|---|---|
| White matter | IDP dMRI TBSS MD Fornix | 3 |
| White matter | IDP dMRI TBSS MD Fornix cres and Stria terminalis R | 1 |
| White matter | IDP dMRI TBSS MD Genu of corpus callosum | 2 |
| White matter | IDP dMRI TBSS MD Inferior cerebellar peduncle L | 3 |
| White matter | IDP dMRI TBSS MD Inferior cerebellar peduncle R | 1 |
| White matter | IDP dMRI TBSS MD Medial lemniscus L | 1 |
| White matter | IDP dMRI TBSS MD Middle cerebellar peduncle | 4 |
| White matter | IDP dMRI TBSS MD Pontine crossing tract | 3 |
| White matter | IDP dMRI TBSS MD Posterior limb of internal capsule L | 2 |
| White matter | IDP dMRI TBSS MD Posterior limb of internal capsule R | 2 |
| White matter | IDP dMRI TBSS MD Posterior thalamic radiation L | 1 |
| White matter | IDP dMRI TBSS MD Posterior thalamic radiation R | 1 |
| White matter | IDP dMRI TBSS MD Retrolenticular part of internal capsule L | 2 |
| White matter | IDP dMRI TBSS MD Sagittal stratum R | 2 |
| White matter | IDP dMRI TBSS MD Superior cerebellar peduncle L | 3 |
| White matter | IDP dMRI TBSS MD Superior corona radiata R | 1 |
| White matter | IDP dMRI TBSS MD Superior longitudinal fasciculus L | 3 |
| White matter | IDP dMRI TBSS MD Superior longitudinal fasciculus R | 2 |
| White matter | IDP dMRI TBSS MD Tapetum L | 1 |
| White matter | IDP dMRI TBSS MD Uncinate fasciculus L | 1 |
| White matter | IDP dMRI TBSS MD Uncinate fasciculus R | 1 |
| White matter | IDP dMRI TBSS MO Anterior corona radiata L | 3 |
| White matter | IDP dMRI TBSS MO Anterior limb of internal capsule R | 1 |
| White matter | IDP dMRI TBSS MO Body of corpus callosum | 1 |
| White matter | IDP dMRI TBSS MO Cerebral peduncle L | 2 |

| | | |
|---|---|---|
| White matter | IDP dMRI TBSS MO Cerebral peduncle R | 1 |
| White matter | IDP dMRI TBSS MO Cingulum hippocampus L | 2 |
| White matter | IDP dMRI TBSS MO Cingulum hippocampus R | 1 |
| White matter | IDP dMRI TBSS MO Corticospinal tract L | 1 |
| White matter | IDP dMRI TBSS MO Corticospinal tract R | 1 |
| White matter | IDP dMRI TBSS MO External capsule L | 1 |
| White matter | IDP dMRI TBSS MO External capsule R | 2 |
| White matter | IDP dMRI TBSS MO Fornix | 3 |
| White matter | IDP dMRI TBSS MO Fornix cres and Stria terminalis L | 3 |
| White matter | IDP dMRI TBSS MO Genu of corpus callosum | 1 |
| White matter | IDP dMRI TBSS MO Inferior cerebellar peduncle L | 2 |
| White matter | IDP dMRI TBSS MO Inferior cerebellar peduncle R | 1 |
| White matter | IDP dMRI TBSS MO Medial lemniscus L | 2 |
| White matter | IDP dMRI TBSS MO Medial lemniscus R | 3 |
| White matter | IDP dMRI TBSS MO Pontine crossing tract | 2 |
| White matter | IDP dMRI TBSS MO Posterior corona radiata R | 1 |
| White matter | IDP dMRI TBSS MO Posterior limb of internal capsule R | 2 |
| White matter | IDP dMRI TBSS MO Posterior thalamic radiation L | 2 |
| White matter | IDP dMRI TBSS MO Posterior thalamic radiation R | 2 |
| White matter | IDP dMRI TBSS MO Retrolenticular part of internal capsule L | 1 |
| White matter | IDP dMRI TBSS MO Sagittal stratum R | 2 |
| White matter | IDP dMRI TBSS MO Splenium of corpus callosum | 1 |
| White matter | IDP dMRI TBSS MO Superior cerebellar peduncle L | 1 |
| White matter | IDP dMRI TBSS MO Superior corona radiata L | 2 |
| White matter | IDP dMRI TBSS MO Superior fronto occipital fasciculus L | 3 |

| | | |
|---|---|---|
| White matter | IDP dMRI TBSS MO Superior fronto occipital fasciculus R | 2 |
| White matter | IDP dMRI TBSS MO Tapetum L | 3 |
| White matter | IDP dMRI TBSS MO Tapetum R | 1 |
| White matter | IDP dMRI TBSS MO Uncinate fasciculus L | 1 |
| White matter | IDP dMRI TBSS MO Uncinate fasciculus R | 4 |
| White matter | IDP dMRI TBSS OD Anterior corona radiata L | 1 |
| White matter | IDP dMRI TBSS OD Anterior limb of internal capsule L | 2 |
| White matter | IDP dMRI TBSS OD Anterior limb of internal capsule R | 2 |
| White matter | IDP dMRI TBSS OD Cingulum cingulate gyrus R | 2 |
| White matter | IDP dMRI TBSS OD External capsule R | 2 |
| White matter | IDP dMRI TBSS OD Fornix cres and Stria terminalis R | 1 |
| White matter | IDP dMRI TBSS OD Inferior cerebellar peduncle L | 3 |
| White matter | IDP dMRI TBSS OD Medial lemniscus L | 1 |
| White matter | IDP dMRI TBSS OD Medial lemniscus R | 2 |
| White matter | IDP dMRI TBSS OD Posterior thalamic radiation L | 2 |
| White matter | IDP dMRI TBSS OD Posterior thalamic radiation R | 1 |
| White matter | IDP dMRI TBSS OD Retrolenticular part of internal capsule L | 1 |
| White matter | IDP dMRI TBSS OD Sagittal stratum L | 1 |
| White matter | IDP dMRI TBSS OD Sagittal stratum R | 1 |
| White matter | IDP dMRI TBSS OD Superior corona radiata R | 1 |
| White matter | IDP dMRI TBSS OD Superior longitudinal fasciculus R | 1 |
| White matter | Volume Cerebral White Matter | 1 |
